# Decoding protein glycosylation by an integrative mass spectrometry-based *de novo* sequencing strategy

**DOI:** 10.1101/2024.08.23.609376

**Authors:** Jing Gao, Hongxu Chen, Hongrui Yin, Xin Chen, Zhicheng Yang, Yuqiu Wang, Jianhong Wu, Yinping Tian, Hong Shao, Liuqing Wen, Hu Zhou

## Abstract

Glycoproteins, representing over 50% of human proteins and most biopharmaceuticals, are crucial for regulating various biological processes. The complexity of multiple glycosylation sites, often leading to incomplete sequence coverage and ambiguous glycan modification profiles. Here, we developed an integrative mass spectrometry-based approach for decoding unknown glycoproteins, which is featured with the combination of deglycosylation-mediated *de novo* sequencing with glycosylation site characterization. We utilized enzymatic deglycosylation of *N*-/ *O*-glycan to achieve comprehensive sequence coverage. Additionally, EThcD fragmentation enables the identification of high-quality long peptides, facilitating precise protein assembly. We subsequently applied this method to *de novo* sequencing of the highly glycosylated therapeutic fusion protein Etanercept (Enbrel^®^). We also sequenced three new tumor necrosis factor receptor (TNFR): Fc-fusion biologics with largely unknown sequences, unveiling subtle distinctions in the primary sequences. Furthermore, we characterized *N*- and *O*-glycosylation modifications of these proteins at subunit, glycopeptide, and glycan levels. This strategy bridges the gap between the *de novo* sequencing and glycosylation modification, providing complete information of the primary structure and glycosylation modifications for glycoproteins. Notably, our method could be a robust solution for accurate sequencing of the glycoproteins and has practical value not only in basic research but also in the biopharmaceutical industry.

## Introduction

Protein glycosylation, as one of the most widespread and complex post-translational modifications (PTMs), plays fundamental roles in many biological processes ^1, 2^. A comprehensive analysis of the primary sequences of glycoproteins, including the identification of glycosylation sites and associated glycan structures, forms the cornerstone of glycoprotein functional research ^3, 4^. It also serves as a critical quality attribute (cQA) for biopharmaceuticals, offering insights into protein structure, function, disease associations, and biotechnological applications.

Liquid chromatography-tandem mass spectrometry (LC-MS/MS) has emerged as a powerful tool for glycoprotein identification, spanning from glycoprotein characterization to glycoproteomic ^5, 6, 7^. Diverging from conventional database searching based protein detection strategies, *de novo* protein sequencing is a novel strategy that focuses on the ‘black-box’ protein without prior knowledge of DNA/amino acid sequence ^8, 9, 10^. In this approach, overlapping peptides are generated by multiple proteases digestion, then analyzed by LC-MS/MS, identified by *de novo* algorithms, and assembled into full-length proteins using a combination of software and manual interpretation. In-depth sequencing of peptides is essential to generate arrays of adjacent overlapping peptides that reveal the full coverage of protein sequence. However, owing to the suppression effects of glycan moieties on glycopeptide sequencing efficiency ^11^, it is hard to get informative glycopeptide fragmentation spectra to support *de novo* sequencing. Moreover, the complexity of spectra comprising both the peptide and glycan fragment, makes it hard to *de novo* interpretation of glycopeptide fragmentation spectra for highly accurate glycopeptide sequencing ^12^. As a result, extensive glycosylation often results in gaps in certain regions, limiting the application of *de novo* sequencing to monoclonal antibodies (mAbs) with consistent structures and known *N*-linked glycosylation sites ^8, 9, 10, 13^.

In this study, we employed a hybridization approach to decode glycosylated proteins, combining deglycosylation-mediated *de novo* sequencing with glycosylation site characterization. The *de novo* sequencing involves enzymatic deglycosylation of *N*-/ *O*-glycans to significantly reduce the difficulty of MS/MS fragmentation and the complexity of glycopeptide-derived fragmentation spectra, thereby achieving full sequence coverage. EThcD fragmentation enables the identification of high-quality long peptides to facilitate precise protein assembly. To test the above integrative workflow, we used a group of highly glycosylated therapeutic recombinant TNFR: Fc-fusion proteins, Etanercept and three sequence unknown TNFR: Fc-fusion biologics as model proteins. These proteins contain numerous and diverse *N*-/ *O*-glycosylation sites as well as an artificial hinge domain, demonstrating the applicability of our integrated approach in some of the most challenging scenarios. Furthermore, we performed characterization of glycosylation by multi-level analysis, including *N*-glycosylation sites identification, released glycan analysis, glycoprotein subunit analysis and glycopeptide analysis, providing comprehensive information on both *N*- and *O*-glycosylation patterns. This method is a powerful strategy for sequencing glycosylation proteins, and is likely to find applications in various biopharmaceuticals as well as biotechnological fields.

## Results and Discussion

### Development of *de novo* sequencing method for glycoproteins

The complex spectrum of glycopeptide containing fragment ions from both the peptide and glycan, makes it difficult to obtain high-quality *de novo* peptide sequencing results, leading to gaps and errors in the sequence coverage. To simplify the mass spectrometry analysis and the peptide sequence determination, we employed glycosidases for sequential hydrolysis of monosaccharides and obtained the primary sequence from the deglycosylated protein by *de novo* sequencing. Subsequently, the glycosylation characterization was carried out on multi-levels, including released glycan analysis, glycoprotein subunit analysis, and glycopeptide analysis, and to pinpoint the *N*-/ *O*-glycosylation (Fig. 1a).

**Figure 1.**
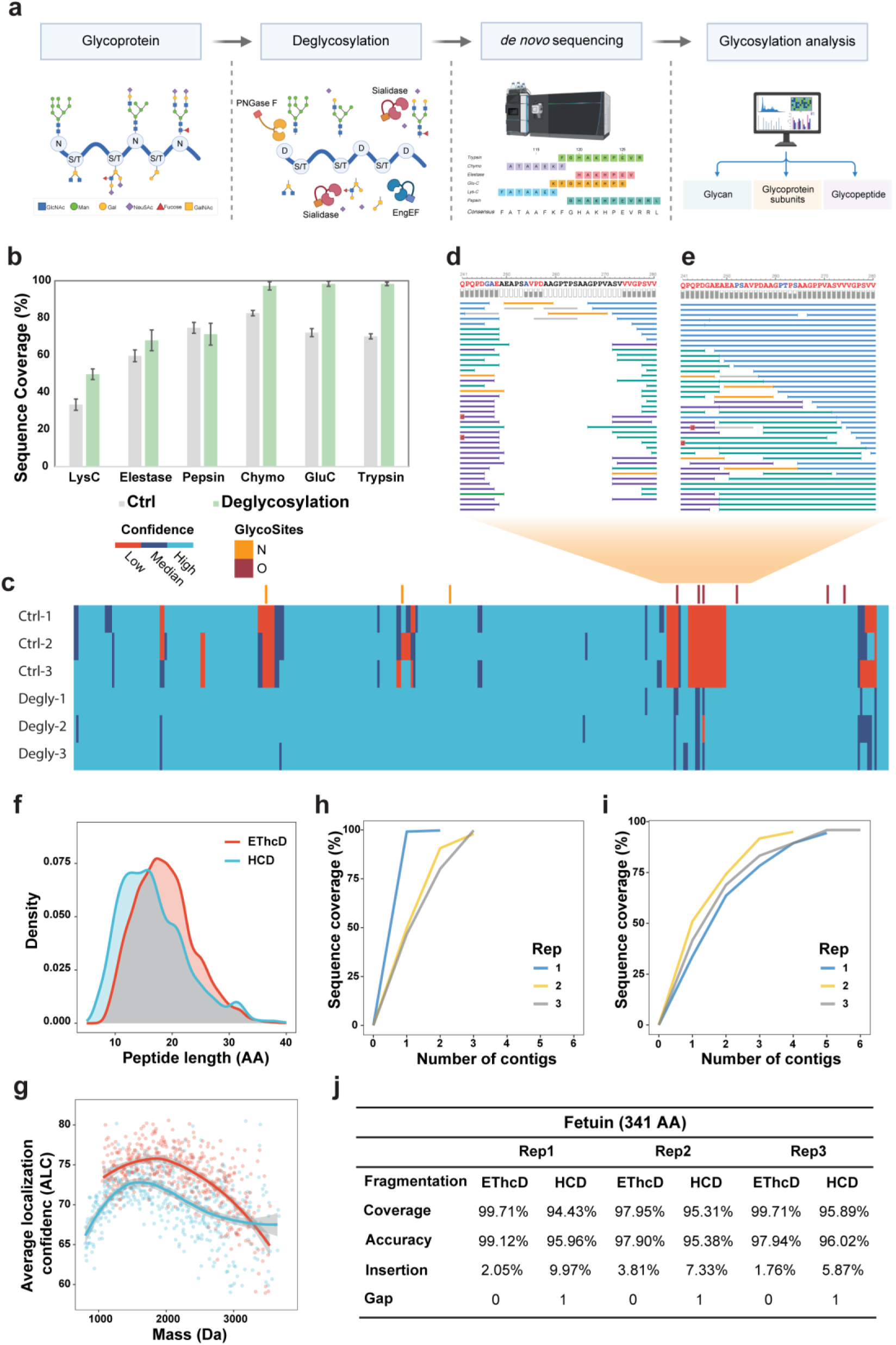
Develop of the de novo sequencing method for glycoprotein. (a) Workflow of glycoprotein de novo sequencing and glycosylation characterization. The glycoprotein containing sialytated N-glycans and O-glycans was treated with deglycosylating enzymes (PNGase F, Sialidase and Eng EF) to sequential hydrolysis of monosaccharides. Then the deglycosylated protein was sent to mass spectrometry based de novo sequencing by multiple proteases digestion to generate overlapping peptides and then assembling peptides into full-length protein followed by correcting I/L and N/D conversions. Finally, on basis of the primary sequence output by de novo sequencing, the glycosylation characterization was carried out on multi-levels, including released glycan analysis, glycoprotein subunit analysis and glycopeptide analysis, to pinpoint the N-/ O-glycosylation. (b) Sequence coverage of Fetuin A digested with different endo-proteinase with/ without deglycosylation treatment. (c) Comparing the impact of deglycosylation on the confidence of sequence identification through the integration of results from multiple proteases (LysC, Elestase, Pepsin, Chymotrypsin, GluC, Trypsin). (d) Zoomed peptide mapping for amino acids 241-280 of Fetuin without deglycosylation using the integration of results from multiple proteases. (e) Zoomed peptide mapping for amino acids 241-280 of Fetuin A following deglycosylation using the integration of results from multiple proteases. (f) Density plot of peptide length distribution. (g) Average localization confidence (ALC) of de novo peptide candidates. Assembly performance of EThcD datasets (h) and stepped HCD datasets (i). It shows the full-length protein sequence coverage (y-axis) when certain numbers of contigs were used (x-axis). (j) Assembly performance comparison between EThcD and stepped HCD using de novo sequenced peptides.

We employed Fetuin A (Uniport Accession No. P12763), a commonly used model glycoprotein containing sialylated *N*-glycans and *O*-glycans (Supplementary Fig. 1), to develop the *de novo* sequencing strategy. Both the PNGase F and *O*-glycosidase EngEF were applied to remove *N*-glycans and *O*-glycans, respectively. As the EngEF is only active on unsubstituted *O*-glycan cores, the digest was performed in combination with NanA. The deglycosylation effect was confirmed by intact mass analysis (Supplementary Fig. 2). As expected, the sequence coverage improved significantly in most of the enzymatic digestion groups after the removal of glycans (Fig. 1b). For Chymotrypsin, Trypsin and Glu-C digestion groups, over 90% sequence coverage were achieved, laying foundation for starting *de novo* glycoprotein sequencing. Considering the complementary of different enzymatic methods, we tried to integrate them with the assumption that the robustness would be improved. Largely like the single enzyme digest results, the integrated results also confirmed the higher confidence of sequence identification after deglycosylation (Fig. 1c). Indeed, the glycopeptide backbone identification was more confident after glycan removal (Supplementary Fig. 3), allowing the in-depth identification of the glycosylation sites containing sequence domain (Fig. 1d, e).

Extensive fragmentation of peptide ions is essential to generate arrays of adjacent fragments that reveal the amino acid sequence. Given the EThcD fragmentation significantly increases the number of fragment ions containing more sequence information in MS/MS scans, which is beneficial for inferring the peptide sequence and post-translational modification ^14^, we assessed the *de novo* peptide sequencing performance of the EThcD fragmentation method comparing with the commonly used stepped HCD fragmentation. Our results revealed that despite the total number of *de novo* peptide candidates was fewer than that of the HCD method (Supplementary Fig. 4), EThcD methods outperformed the HCD methods in terms of peptide sequence length (Fig. 1f and Supplementary Fig. 5a) and spectrum quality (Fig.1g and Supplementary Fig. 5b). Moreover, the longer sequencing peptide fragment of EThcD spectrum facilitates the assembly of protein. The top 3 contigs with overlapped sequences of EThcD covered nearly complete of the full length (Fig. 1h), while the HCD method generated ∼5 contigs covering the ∼95% sequence (Fig. 1i). Importantly, considering the sequencing accuracy, EThcD achieved an overall average sequence accuracy of 98.3% (Fig. 1j and Supplementary Fig. 6), while the stepped HCD method resulted in 95.8% overall average sequence accuracy as well as 1 gap (Fig. 1j, Supplementary Fig. 7).

Inspection of the assembles, the most error frequency occurred on Ile-Leu (I-L) and Asn-Asp (N-D) conversion. The three N-D conversions are in good agreement with the *N*-glycosylation sites, on which the attached *N*-glycan was released by PNGase F (Supplementary Fig. 6), converting glycosylated Asn residues into Asp ^15^. This result indicated that both the HCD and EThcD methods can faithfully reveal the actual amino acids of the peptide mixture. The Asn residues that are *N*-glycosylated can be determined by ^18^O labeling based peptide mapping method as well as Glycosidase based method (see later). Notably, EThcD method showed its advantage in reliable Leu/Ile determination. With the diagnostic w’ -ion in EThcD spectrum (characteristic loss of C_3_H_7_ (−43.05 Da) for Leu or C_2_H_5_ (−29.04 Da) for Ile from the side chain of particular z-ions) ^16^, we reliably identified 12 out of 13 I-L conversion occur in HCD datasets (data not shown, peptide DIEIDTLETTCH as an example, Supplementary Fig. 8).

Consequently, EThcD mass acquisition simplifies the protein assemble and enabled higher accuracy for protein *de novo* sequencing. In combine with the glycan removal and EThcD mass spectrometry acquisition, we developed a robust method for glycoprotein *de novo* sequencing, enabling precise sequence coverage in the glycosylation region.

### *De novo* sequencing of highly glycosylated TNFR: Fc-fusion pharmaceuticals

To test the robustness of our *de novo* strategies and demonstrate their application for highly glycosylated biopharmaceuticals, we applied the developed method to sequencing a group of Fc-fusion proteins including one marketed therapeutic Fc-fusion protein original drug Etanercept and three new TNFR: Fc biologics (Supplementary Table 1) whose sequences were unknown. As one of the most complex highly glycosylated Fc-fusion pharmaceutical proteins, Etanercept possesses at least 3 *N*- and 13 *O*-glycosylation sites, a soluble fusion protein of the tumor necrosis factor receptor extracellular domain, linked to a Fc part of IgG1 ^17^. We confirmed the deglycosylation efficacy for Etanercept by SDS-PAGE (Supplementary Fig. 9) and intact mass spectrometry measurement (Supplementary Fig. 10). A wide range of glycan structures and attachment sites contributed to the heterogeneity, leading to the failure of intact mass measurement under the denature condition (Supplementary Fig. 10a, c). After releasing the glycans, the heterogeneity was significantly reduced, enabling the precise measurement of intact protein mass (Supplementary Fig. 10b, d). Following the same approach, we successfully deglycosylation and evaluated the effects of 3 new TNFR:Fc biologics (Supplementary Fig. 11-14). Notably, the intact mass results revealed the subtle distinctions in the primary sequences of TNFR:Fc 2 and TNFR:Fc 3 to Etanercept, while the intact mass of TNFR:Fc 1 is as same as Etanercept.

The MS/MS spectra were obtained by EThcD fragmentation for the following *de novo* protein sequencing. Owing to the deep sequence coverage achieved by deglycosylation treatment, we can assemble the entire protein sequence of Etanercept as one contig (Supplementary Fig. 15). The three replicates reached 98.93% coverage and 99.57∼100% accuracy (Supplementary Table 2). While the conventional strategy failed to assemble probably because of the ambiguous gaps rising from the complex MS/MS spectra of glycosylated peptide which could not be solved by *de novo* sequencing algorithm and multi-enzyme digestion strategy (Supplementary Fig. 16). Indeed, the gap is located in the Hinge region, which is absent in the human proteome database and reported highly *O*-glycosylated to impede the mass sequencing of peptide, indicating that it is impossible to obtain the primary sequence with conventional database searching approach. In summary, our method enables *de novo* sequencing of glycoproteins and achieves deep coverage of glycosylation modification regions.

### Comparison of sequence similarity between Etanercept with TNFR:Fc biologics

Inspire of sequence similarity between the samples reveals subtle differences between TNFR:Fc 2/ TNFR:Fc 3 and Etanercept at three positions, specifically: M174R ( in TNFR domain), E376D and M378L (both in Fc domain) (Fig. 2a and Supplementary Fig. 17). The MS/MS spectra corresponding to the mutated sites demonstrate good quality and accurate localization (Fig. 2b-e and Supplementary Fig. 18, 19). Moreover, the differences in protein sequences are aligned with the intact protein mass (Figure 2f, g, Supplementary Fig. 20 and Supplementary Table 3). The sequence variant on M174R is a polymorphism in the coding region of human TNFR2, and seems to be associated with polycystic ovary syndrome ^18^, hyperandrogenism and systemic lupus erythematosus ^19^. Similarly, E376D and M378L variations on Fc domain were also reported in commercially available TNF receptor2-Fc fusion protein products ^20^.

**Figure 2.**
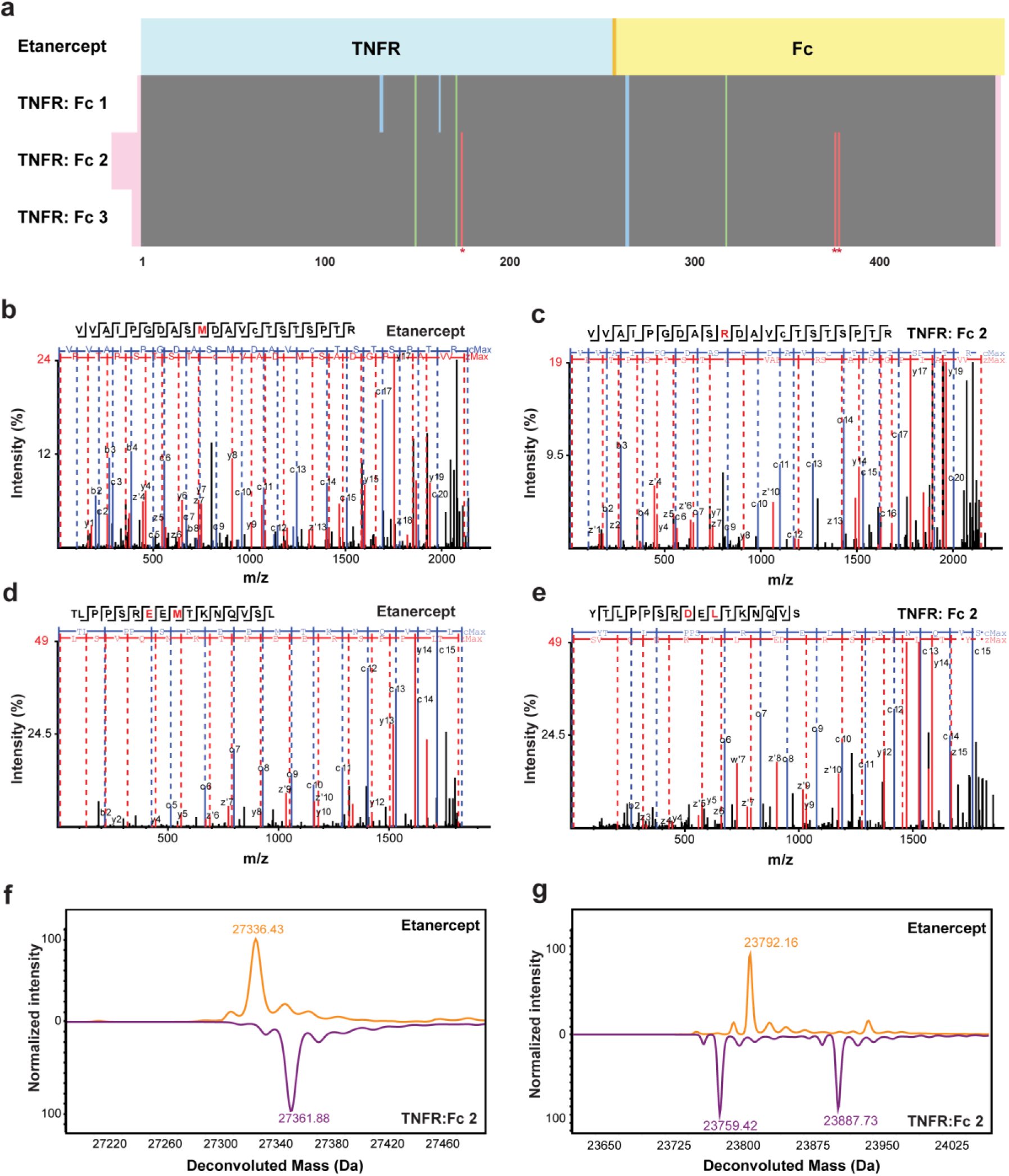
*De novo* sequencing of TNFR: Fc-fusion biopharmaceuticals. (a) Sequence similarity of de novo sequencing results of Etanercept and three new TNFR: Fc biologics. (b) MS/MS spectra of original peptide containing Met174 in Etanercept. (c) MS/MS spectra of variant peptide containing M174R in TNFR: Fc 2 biologics. (d) MS/MS spectra of original peptide containing E376 and M378 in Etanercept. (e) MS/MS spectra of variant peptide containing E376D, M378L in TNFR: Fc 2 biologics. (f) Mirror plot of deconvoluted mass spectra of TNFR fragment of Etanercept and TNFR: Fc 2 biologics. (g) Mirror plot of deconvoluted mass spectra of Fc fragment of Etanercept and TNFR: Fc 2 biologics.

We validated three TNFR:Fc biologics protein sequences by searching against the sequence determined by *de novo* sequencing using the sequence validation module of the PEAKS AB software. ∼99% sequence had high identification confidence (Fragment ion confidence > 85%), with multiple spectra supporting each tryptic peptide (Supplementary Fig. 21). Therefore, the results confirmed the high accuracy of the *de novo* sequencing results.

### Analysis of *N*- and *O*-glycosylation of TNFR:Fc-fusion pharmaceutical proteins

Since glycosylation can influence the biological function of glycoprotein, especially for the efficacy and safety of the biopharmaceutical ^21, 22^, a thorough assessment of glycosylation is as important as that of amino-acid sequence to decoding the detailed primary structure of glycoproteins. Here, we employed multi-level analysis, including *N*-glycosylation sites identification, released glycan analysis, glycoprotein subunit analysis and glycopeptide analysis, to pinpoint the *N*- and *O*-glycosylation.

We initially conducted quantitative PNGase F-catalyzed glycosylation site ^18^O stable isotopic labeling analysis to determine the *N*-glycosylation sites. PNGase F specifically deamidated the *N*-glycan linked Asn to Asp labeled with ^18^O-water and resulted in the mass addition of 2.98 Da on Asn which is glycosylated (Asn to Asp conversion causes an increment of +0.98 Da and ^18^O labeling causes an increment of +2.00 Da, Fig. 3a, Supplementary Fig. 22). By calculating the peptide intensity with ^18^O labeled sites, three *N*-glycosylation sites (N149, N171 and N317) were screened out in all four TNFR: Fc fusion protein samples (Ratio cutoff > 2, ^18^O-labeling sample/ the control sample labeled in ^16^O-water) (Fig. 3b). The three *N*-glycosylation sites were exactly matched with known *N*-glycosylation sites in Etanercept. Furthermore, we confirmed the *N*-glycosylation site identification results by partially cleaving *N*-glycan by Endo H to generate a fixed modification with ‘GlcNAc’ on the *N*-glycosylation ^23^ (Fig. 3a, c and Supplementary Fig. 23). Indeed, the results are highly consistent with the ^18^O labeling results, all three sites can be identified as *N*-glycosylation (Fig. 3d). In particular, these data can be used to correct the Asn to Asp conversion in *de novo* sequencing results.

**Figure 3.**
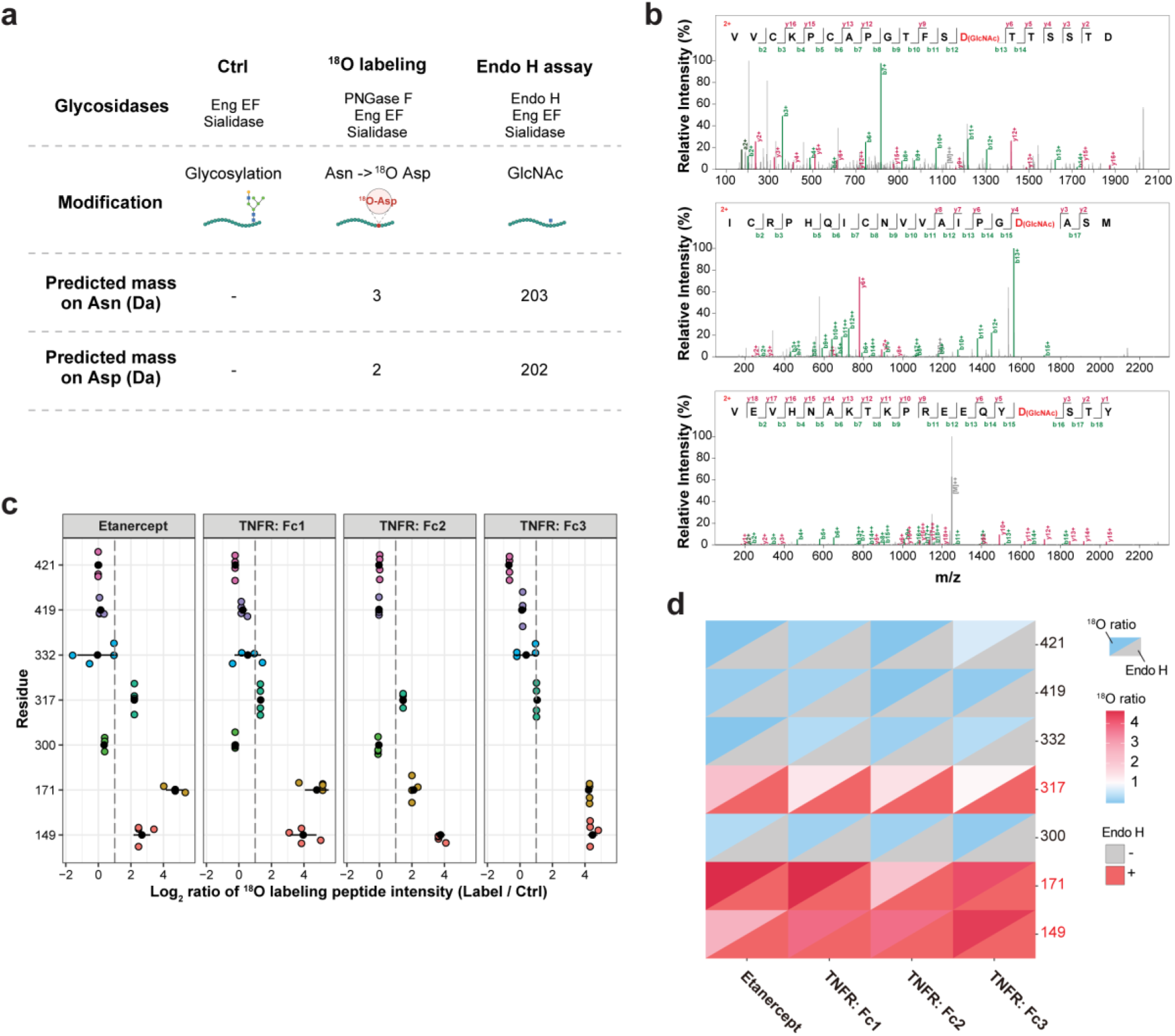
Identification of N-glycosylation sites. (a) Schematic overview of the protocol with mass signatures for peptide glycosylation sites that with ^18^O labeling and Endo H treatment. (b) ^18^O labeling based quantitative screening of N-glycosylation sites. We compared the intensity peak area of ^18^O labeled peptides from labeled to control samples to quantify the relative distribution of the N-glycosylation state at each glycosylation site detected. (c) Exemplary MS/MS spectra in support of the assigned N-glycosylation sites on Etanercept. The peptide sequence and fragment coverage are indicated in the top-left of each spectrum spectra, with b ions indicated in green and y ions in magenta. The same color annotation is used for peaks in the spectra, with additional peaks such as intact/charge reduced precursors, neutral losses, and immonium ions indicated in grey. The *de novo* sequencing output sequences, with potential Asn to Asp conversions, was used for database searching. GlcNAc (D +202.10 Da) modification sites are indicated in red. (d) The Consistency evaluation of the N-glycosylation site identification by ^18^O labeling based quantitative screening and Endo H treatment.

To pinpoint the content and structure of *N*-glycans on the whole protein level, enzymatically released *N*-glycans are derivatized with 2-AB labeling to allow for fluorescence and mass spectrometry detection by UPLC-HILIC-FLR-MS. The most abundant peaks of *N*-glycans released from Etanercept were F(6)G2F, biantennary, core-fucosylated *N*-glycans with two terminal galactose, followed by the F(6)G0F, biantennary, core-fucosylated with no terminal galactose residues, and G2, biantennary *N*-glycans with two terminal galactose residues, and F(6)G1F, biantennary, core-fucosylated with one terminal galactose residues (Fig. 4a). The results of three TNFR: Fc biologics were similar to Etanercept (Supplementary Fig. 24).

**Figure 4.**
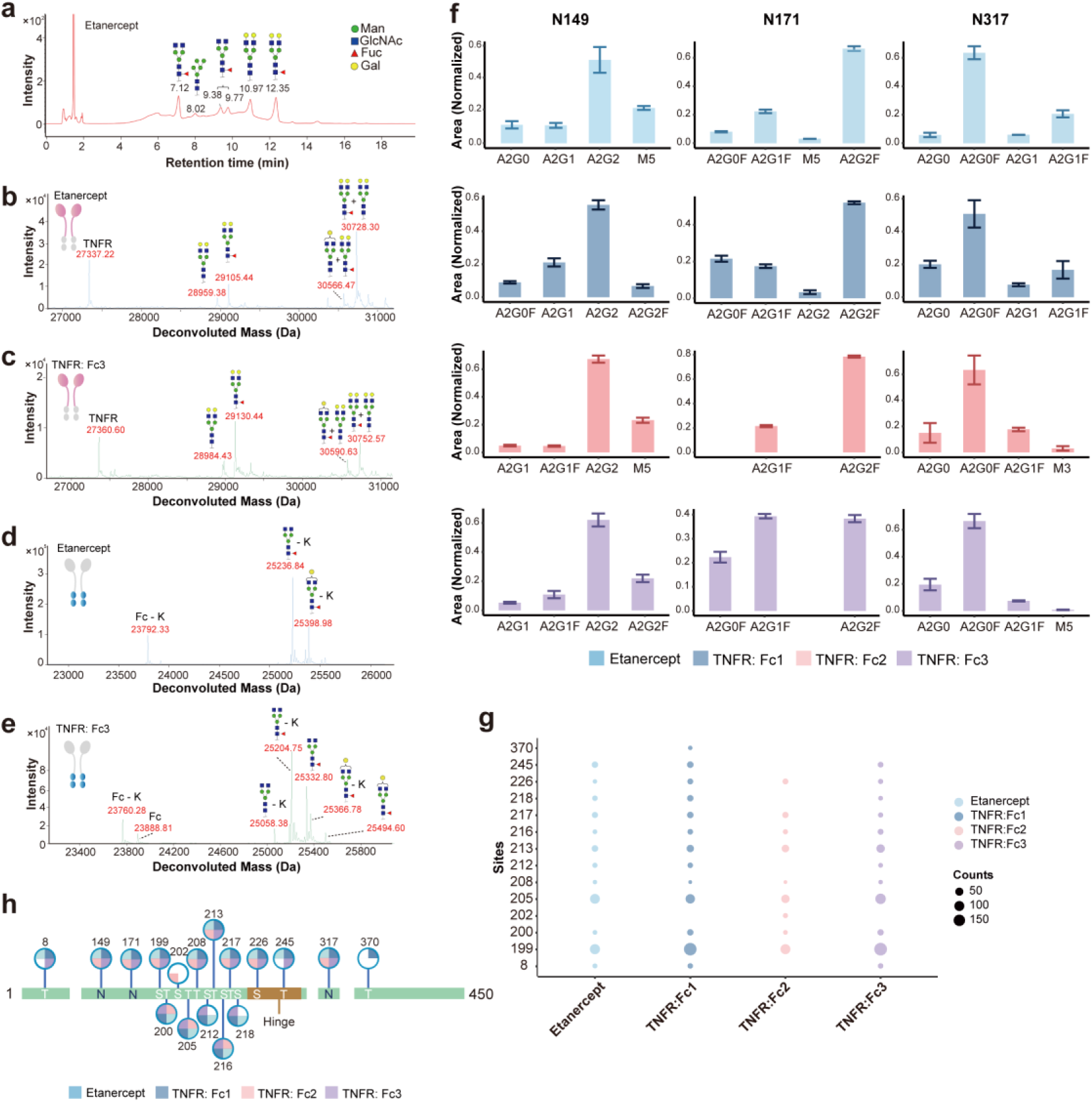
N- and O-glycosylation analysis of TNFR :Fc-fusion pharmasutical proteins (a) HILIC-UPLC profile of the total N-linked glycans released from Etanercept after desialylation. Deconvoluted spectrum of subunit analysis for Etanercept TNFR domain (b), TNFR: Fc 3 TNFR domain (c), Etanercept Fc domain (d) and TNFR: Fc 3 Fc domain (e). (f) Glycopeptide profiles of Asn147, Asn179 and Asn317. (g) Site-specific statistics of O-glycosylation glycoforms. (h) Summary plot of N-/ O-glycosylation modification on Etanercept and three biologics.

Furthermore, we assessed the *N*-glycan variants at the subunit level after removing *O*-glycans as well as the sialic acid. Slight differences were observed in the samples (Supplementary Fig. 25). The top abundant glycoforms at TNFR domain of Etanercept/ TNFR:Fc1 carries two different *N*-glycans, namely G2 and G2F (Fig. 4b and Supplementary Fig. 25). Whereas TNFR domain of the TNFR:Fc3 also contained a number of the G1F glycoform, in addition to the major glycoforms G2 and G2F (Fig. 4c and Supplementary Fig. 25). On the other side, the most prominent glycoforms at Fc domain were assigned to G0F and G1F constantly in four samples, despite a subtle varying degrees of Lysine-loss modification was observed (Fig. 4d, e and Supplementary Fig. 25).

Moving down site-specific *N*-glycopeptide level (Fig. 4f, Supplementary Table 4), the results are in consistent with the assignment of glycoforms based on subunit level, verifying the coherence of data integration at different digestion levels. Indeed, the *N*-glycopeptides provide higher resolution of multiple glycovariants, namely the N149, N171 and N317 glycosylation site is mainly occupied by G2, G2F and G0F, respectively. The N171 is the site that contributed to the difference of glycoforms for TNFR:Fc3, containing a considerable amount of the G1F glycoform.

We identified 14 *O*-glycosylation sites on Etanercept and three TNFR: Fc biologics (Fig. 4g, Supplementary Table 5). Consistent with previous reports^17^, Etanercept displayed 12 *O*-glycosylation sites. Similar *O*-glycosylation patterns were observed in the TNFR: Fc biologics. Furthermore, a comprehensive assessment of glycosylation sites revealed the highest level of glycosylation in the hinge region (Fig. 4h).

Despite structural similarities in their backbones, Etanercept and the TNFR:Fc biologics exhibited variability in glycoform profiles attributed to distinct manufacturing processes and cell sources. Notably, while TNFR: Fc closely resembled Etanercept in glycoform distribution across *N*-glycosylation sites, TNFR: Fc2 and TNFR: Fc3 exhibited more similar profiles to each other. These findings highlight potential physicochemical implications that warrant further investigation and contribute to the comprehensive characterization of Etanercept and its biosimilars.

## Discussion

High-quality MS/MS fragmentation spectra and complete coverage are fundamental for protein *de novo* sequencing. To address the adverse effects of glycosylation modifications on sample heterogeneity, the efficacy of MS/MS fragmentation, and the complexity of fragmentation patterns, we utilized deglycosylating enzymes to release glycans, thereby focusing the analysis on the peptide backbone, leading to improved sequence coverage. Furthermore, the use of deglycosylating enzymes led to the specific conversion of Asn to Asp during *N*-glycan release, as supported by *de novo* sequencing results, which could be rectified through *N*-glycosylation site determination strategies. Other commonly used chemical based deglycosylation methods, such as hydrazinolysis for *N*-glycan release and beta-elimination reaction for *O*-glycan, should be carefully optimized of reaction conditions to ensure efficient deglycosylation and minimize protein degradation or modification. However, the heterogeneity of glycans attached to glycoproteins can affect the accessibility of glycosidase enzymes to their target sites, leading to variability in deglycosylation efficiency. In this study, we optimized glycosidase enzymes for glycosylation modifications in mammals, particularly for biopharmaceuticals from CHO cells. Given the similarity of the major classes of glycoconjugates present in mammals, these glycosidases theoretically can be applied to sequentially hydrolyze monosaccharides from most human and mammalian glycosylated proteins. For proteins originating from other sources, such as fungi or plants, glycan analysis should be conducted first to optimize glycosidase enzymes for proper deglycosylation efficiency.

Inspection of the *de novo* sequencing results of Fetuin and Etanercept, challenges remain in sequencing the N- and C-terminal regions of proteins, where sequencing errors in terms of substitution and insertion of amino acids frequently occurred. This is consistent with the CD147 sequencing results reported by Mai et al ^24^, despite the different assemble software used. We speculated two reasons were account for it. Firstly, post-translational modifications (PTMs) occur at the N- and C-terminal residues of proteins, leading to peptide fragmentation pattern alterations and complicating *de novo* sequencing. Actually, C-terminal Lysine is usually partially removed during mammalian cell culture ^25^. Especially for the antibody, the first major modification is the removal of C-terminal Lysine ^26, 27^. We also observed varying degrees of Lysine-loss modification in our intact mass and subunit analysis for Etanercept and three biologics. Lys-loss accounts for approximately 90% of Etanercept, which may lead to the inability to accurately determine the terminal ‘K’. Additionally, the availability of overlapping peptides derived from terminal regions may be reduced, thus limiting the contextual information provided by adjacent amino acids. In our Etanercept sequencing dataset, we found that the C-terminal peptide data only came from Glu-C and Pepsin digested samples. Considering the enzyme-specificity, the Deepnovo algorithm mistakenly interpreted the sequence ‘LSPG’ as ending with the acidic amino acid E, outputting the sequence ‘PKE’. Here, ‘LSPG’ and ‘PKE’ have nearly identical monoisotopic peak masses 372.20 Da. Although commercial software with built-in antibody structural frameworks can reduce error probabilities for antibody sequencing, strategies are needed to enhance the accuracy and sensitivity of *de novo* sequencing for various proteins lacking uniform structural frameworks. Additionally, the top-down mass spectrometry can be used to enhance the accuracy of protein terminal sequencing. Liu et al reported an algorithm named TBNovo, in which a top-down tandem mass spectrum is utilized as a scaffold, and bottom-up tandem mass spectra are aligned to the scaffold to increase sequence coverage ^28^. Although there remain numerous challenges particularly for the robust characterization of proteins larger than ∼30 kDa, top-down MS exhibits effective sequencing capability for protein terminals ^29, 30^, particularly C-terminal sequences. In conclusion, the challenges inherent in accurately sequencing the N- and C-terminal regions of proteins persist despite advancements in sequencing technologies and software. Addressing these challenges will be essential for advancing our understanding of novel proteins and their functional implications in biological systems.

## Conclusion

Mass spectrometry-based protein *de novo* sequencing is a powerful tool to reveal the amino acid sequence. Combined with glycosidase enzymatic dissection, EThcD-based protein sequencing and glycosylation characterization, the novel strategy opens up new avenues in the insight of heavily glycosylated biopharmaceuticals. Our analysis of Etanercept demonstrates that the method provides comprehensive information not only unravels the amino acid sequence but also pinpoints the glycosylation sites, elucidating the complex landscape of glycoform heterogeneity. Given the commonality of *N*-/*O*-glycosylation in protein expressed in CHO cell lines, which comprise the majority of protein therapeutics, the study lays the framework for successful analysis of various biopharmaceuticals, which are instrumental in the treatment of a wide range of diseases.

## Methods

### Chemicals and materials

Sialidase from *Streptococcus pneumoniae* (NanA) was prepared as reported previously^31, 32^. PNGase F from *Flavobacterium meningosepticum* and endoglycosidase H from *Streptomyces plicatus* (Endo H) were prepared as reported previously^33^. Endo-α-GalNAcases from *Enterococcus faecalis* (EngEF)^34^ was cloned into the pET-28a vector with six histidines tag (6xHis tag) for purification by Ni-affinity chromatography. Etanercept was obtained from Pfizer (Enbrel^®^, Rijksweg, Belgium, Lot-No: GA5942). Three new therapeutic recombinant human tumor necrosis factor receptor fusion protein biologics were obtained from Sunshine Guojian Pharmaceutical (Shanghai, China, Lot-No: 301-C2031), BioRay Biopharmaceutical (Zhejiang, China, Lot-No: 0500222005), Celgen Biopharma (Shanghai, China, Lot-No: A201222U), respectively. LC-MS grade acetonitrile and water were purchased from Thermo Fisher Scientific (Nepean, ON, Canada). Fetuin, sodium deoxycholate (SDC), Tris(2-carboxyethyl)phosphine (TCEP), chloracetamide (CAA), dithiothreitol (DTT), 2-AB dye, formic acid, acetic acid, ammonium bicarbonate (ABC), ammonium formate, guanidine hydrochloride, Tris-HCl and Pepsin were purchased from Sigma-Aldrich (St. Louis, MO). MS sequencing grade Trypsin, Elastase, AspN, Chymotrypsin and IdeZ were purchased from Promega (Madison, WI). MS sequencing grade Glu-C was purchased from Roche (Basel, Switzerland). MS sequencing grade Lys-C was purchased from FUJIFILM Wako Chemicals (Osaka, Japan). ^18^O-water (98 atom% of ^18^O) was purchased from Shanghai Engineering Research Center of Stable Isotope (Shanghai, China).

### Sample preparation for *de novo* sequencing and glycopeptide analysis

30 μg of protein was incubated in 30 μL of 2% SDC, 50 mM Tris–HCl, 10 mM TCEP and 40 mM CAA (pH 8.0) at 95 °C for 10 min to accomplish denature, reduction and alkylation in one step. The sample was then diluted with 30 μL of PBS buffer (pH 7.6). For *de novo* sequencing, *N*-glycans and *O*-glycans were released by adding a cocktail of 2U of NanA, 2U of EngEF and 1U of PNGase F followed by incubating at 37°C for 2h while shaking at 1,200 rpm. The sample was then diluted with 60 μL of 50 mM Tris–HCl (pH=8.0) and digested by one of the following proteases: Trypsin, Chymotrypsin, Lys-C, Glu-C, Pepsin and Elastase at a 1:50 ratio (w/w) for 16 h. The digestion was performed at 37 °C, except for Chymotrypsin reaction work at 25°C. For Pepsin digestion, samples were acidified to pH 3.5 using 10% formic acid before adding the enzyme. After digestion, SDC was removed by adding 8 μL of pure formic acid and centrifuging at 14,000*g* for 5 min at 4 °C. The supernatant containing the peptides was then collected for desalting with homemade C18 tips.

The *N*-glycopeptide sample preparation was conducted for glycosylation characterization, with modifications involving NanA and EngEF instead of a cocktail of three endoglycosidases and utilizing Chymotrypsin, Pepsin and Trypsin for digestion. Similarly, the *O*-glycopeptide sample preparation was performed, with modifications utilizing PNGase F and NanA instead of a cocktail of three endoglycosidases and utilizing Trypsin for digestion.

### Sample preparation for *N*-glycosylation site identification

30 μg of pepsin-treated *O*-glycan removed peptides (treated with NanA and EngEF) were subjected to ^18^O labeling quantitative analysis for glycosylation site identification. Equal amounts of both *N*- and *O*-glycan (treated with NanA, EngEF, and PNGase F) released peptides were set as the control. The peptides were thoroughly dried overnight in a speed Vac and resuspended in 30 μl of ABC buffer (ammonium bicarbonate prepared in H_2_^18^O). Asn-linked *N*-glycans were removed by 1μL of PNGase F (8U). The sample was incubated for 3 h at 37 °C while shaking at 1,200 rpm. The peptides were then collected for desalting with homemade C18 tips.

The Endo H-treated peptide sample preparation was conducted as sample preparation for *de novo* sequencing, except for replacing the PNGase F with Endo H in a 1:10 ratio (w/w).

### Sample preparation for intact and subunits mass analysis

For release of N-glycans from protein sample, 80 μg of protein was digested with 4 U of PNGase F in PBS buffer (pH 7.6) in a final volume of 80 μL. The protein was incubated for 2 h at 37 °C while shaking at 1,200 rpm.

For release of O-glycans from protein sample, 80 μg of protein was digested with a combination of 4U of sialidase and 4U of EngEF in PBS buffer (pH 7.6) in a final volume of 80 μL. The sample was incubated for 1 h at 25 °C followed by 16 h at 37 °C while shaking at 1,200 rpm.

For reduce the disulfide bonds, the sample was reduced with a final concentration of 20 mM DTT in 3M guanidine hydrochloride at 57°C for 45 minutes. To prepare for subunits, 50 μg of protein was digested with 1μL of IdeZ in PBS buffer (pH 7.6) in a final volume of 50 μL. The sample was incubated for 1 h at 37 °C while shaking at 1,200rpm.

### LC-MS/MS for peptide *de novo* sequencing

0.5 μg of peptides were separated by online reversed-phase chromatography on a Vanquish™ Neo UHPLC system (self-packed column 75μm × 150mm; 3 μm ReproSil-Pur C18 beads, 120 Å, Dr.Maisch GmbH, Ammerbuch, Germany) coupled to an Orbitrap Eclipse mass spectrometer (Thermo Fisher Scientific, Waltham, MA). Samples were eluted over a 60 min gradient at a flow rate of 300 nL/min. Mobile phase A consisted of 0.1% (v/v) formic acid in H_2_O and Mobile phase B consisted of 0.1% (v/v) formic acid in 80% acetonitrile. The gradient was set as follows, 3%-12%B in 1 min; 12%-30% B in 45 min; 30%-80% B in 5 min; 80%-99% B in 3 min; 99% B for 5 min; 99%-0% B in 1 min.

Peptides were analyzed with a resolution setting of 60,000 in MS1 scans. MS1 scans were obtained with a standard automatic gain control (AGC) target of 400,000, a maximum injection time of 50 ms, and a scan range of 350-2000 m/z. The precursors were selected with a 2 m/z window and fragmented by stepped high-energy collision dissociation (HCD) or electron-transfer high-energy collision dissociation (EThcD) in a duty cycle of 3 seconds. The stepped HCD fragmentation included steps of 27%, 35% and 40% normalized collision energies (NCE). EThcD fragmentation was performed with calibrated charge-dependent electron-transfer dissociation parameters and 30% NCE supplemental activation. For both fragmentation types, MS2 scans were acquired at a 30,000 resolution, a 400,000 AGC target and a 250 ms maximum injection time. The charge state(s) was set as 3-7 and 2-7 for EThcD and HCD fragmentation, respectively. Dynamic Exclusion was set for 30 seconds.

### LC-MS/MS for *N*-glycosylation site identification

0.5 μg of peptides were separated by online reversed-phase chromatography on an EASY-nLC1000 system (self-packed column 75μm × 150mm; 3 μm ReproSil-Pur C18 beads, 120 Å, Dr.Maisch GmbH, Ammerbuch, Germany) coupled to a Q Exactive HF mass spectrometer (Thermo Fisher Scientific, Waltham, MA). Peptides were eluted over a 60 min gradient at a flow rate of 300 nL/ min. Mobile phase A consisted of 0.1% (v/v) formic acid in H_2_O and Mobile phase B consisted of 0.1% (v/v) formic acid in acetonitrile. The gradient was set as follows, 1%-4%B in 1 min; 4%-26% B in 43 min; 26%-32% B in 5 min; 32%-90% B in 2 min; 90% B for 9 min.

Peptides were analyzed with a resolution setting of 60,000 in MS1 scans. MS1 scans were obtained with an AGC target of 3,000,000, a maximum injection time of 20 ms, and a scan range of 150-1700 m/z. The top 20 precursors were selected with a 1.2 m/z window and fragmented by HCD with 27% NCE. MS2 scans were acquired at a 15,000 resolution, with an AGC target of 100,000 and a maximum injection time of 100 ms. Dynamic Exclusion was set for 30 seconds.

### LC-MS/MS for *N*-glycopeptide analysis

*N*-glycopeptide analysis was performed as described previously with slight modifications^35^. 0.5 μg of peptides were separated by online reversed-phase chromatography on an EASY-nLC1200 system (self-packed column 75μm × 150mm; 3 μm ReproSil-Pur C18 beads, 120 Å, Dr.Maisch GmbH, Ammerbuch, Germany) coupled to an Orbitrap Fusion mass spectrometer (Thermo Fisher Scientific, Waltham, MA). Samples were eluted over a 60 min gradient at a flow rate of 300 nL/min. Mobile phase A consisted of 0.1% (v/v) formic acid in H_2_O and Mobile phase B consisted of 0.1% (v/v) formic acid in 80% acetonitrile. The gradient was set as follows, 2%-8%B in 1 min; 12%-30% B in 45 min; 30%-50% B in 5 min; 50%-100% B in 3 min; 100% B for 5 min.

Orbitrap spectra (AGC target 400,000, maximum injection time of 50 ms) were collected from 375-2,000 m/z at a resolution of 120,000 followed by oxonium ions triggered data-dependent HCD MS/MS at a resolution of 30,000 using an isolation width of 1 m/z for 20% collision energy and 2 m/z for 33% collision energy. Charge state screening was enabled to reject unassigned and singly charged ions. A dynamic exclusion time of 30 seconds was used to discriminate against previously selected ions.

### LC-MS/MS for *O*-glycopeptide analysis

0.5 μg of glycopeptides were separated by online reversed-phase chromatography on a Vanquish™ Neo UHPLC system coupled to an Orbitrap Eclipse mass spectrometer (Thermo Fisher Scientific, Waltham, MA). The LC-MS/MS paraments were the same as those for peptide *de novo* sequencing with EThcD fragmentation.

### Intact mass analysis

Intact proteins were analyzed using an Agilent 1290 Infinity II LC coupled with a 6545 QTOF mass spectrometer (Agilent, Santa Clara, CA). 1 μg of protein was injected for each LC/MS run. Protein samples were separated with an Agilent PLRP-S column (1.0 × 50 mm, 5 μm, 1000Å). Protein was eluted over a 12 min gradient (hold at 5% B for 5 min, 5-95% B for 5 min, 95% hold for 2min) at a flow rate of 0.300 mL/min. Mobile phase A was made up of water with 0.1% formic acid, while Mobile phase B was made up of acetonitrile with 0.1% formic acid. The mass spectrometry instrument parameters were set as the following: the dry gas flow rate was set 10.0 L/min at 325 °C, the nebulizer was set at 50 psig, the capillary voltage was set at 4.5 kV and the scan range was from 500 to 3000 m/z at 1 Hz. The capillary voltage was set at 4.5 kV and the scan range was from 500 to 3200 m/z at 1 Hz for fetuin, reduced or middle-down TNFR:Fc fusion protein samples digested by IdeZ and 500 to 6000 m/z at 1 Hz for native TNFR:Fc fusion protein samples. The LC/MS raw data was processed using MassHunter BioConfirm (Version 10.0, Agilent, Santa Clara, CA, USA) to deconvolute the protein average masses.

### Released *N*-glycans analysis

The labeled released N-glycans were analyzed on a Waters UPLC-Xevo G2-S QTOF system (Waters, Milford, MA). The separation of N-glycans was conducted at 60 °C using a Glycan BEH Amide column (2.1 ×150 mm, 1.7 µm, 130 Å, Waters, Milford, MA). Mobile phase A was 50 mM ammonium formate in water (pH 4.4), while Mobile phase B was 100% acetonitrile. The gradient was set as 30%-47% A over 33 min at a flow rate of 0.4 mL/ min, 47%-80% A in 2 min and maintained at 80% A for 3min at a flow rate of 0.25 mL/ min, and 80%-30% A in 1min and maintained at 30% A for 5.00 min at a flow rate of 0.4 mL/ min for reequilibration. The FLR detector was normalized and then set at 330 nm excitation wavelength and 420 nm emission wavelength. The MS settings were set as follows: scan range, 200-2000 m/z; capillary voltage, 3.0kV; cone voltage, 40 V; desolvation temperature, 350 °C; source temperature, 120 °C. Acquired MS data of the glycans were automatically processed using the UNIFI 1.9.3 Scientific Information System, and a Waters Glycan GU library was used for glycan identification.

### Data processing for peptide *de novo* sequencing and protein assemble

Automated peptide *de novo* sequencing was performed with PEAKS AB (version 2.0, Bioinformatics Solutions Inc.) ^13^. Scans were not filtered or merged, and precursors were corrected by mass only. Precursor and product mass tolerance were set to 10 ppm and 0.02 Da, respectively. Enzymes were specified on a sample-by-sample basis. Carbamidomethylation (Cys +57.02 Da) was set as a fixed modification, with Oxidation (Met +15.99 Da) as variable modifications. The peptide candidates with average local confidence (ALC) scores above 50 were used for protein assemble. The peptide candidates were subjected to the protein assembly process, performed with Multiple Contigs & Scaffolding (MuCS) algorithm using the default parameters^24^.

### Sequence validation for protein *de novo* sequencing

Sequence validation was performed with PEAKS AB (version 2.0, Bioinformatics Solutions Inc.) using the *de novo* sequencing output sequences as reference sequences with default paraments.

### Data process for *N*-glycosylation sites identification

Quantitative ^18^O labeling N-glycosylation sites analysis was performed with Biopharma Finder (version 4.0, Thermo Fisher Scientific, Waltham, MA) to search against the *de novo* sequencing output sequences, with potential Asn to Asp conversions caused by the deglycosylation of the Asn-linked glycan by PNGase F. The parameters for peptide mapping included: enzyme, partial pepsin; missed cleavages allowed, two; fixed modification, Carbamidomethylation (Cys +57.02 Da); variable modifications, Asp-Asn substitute (Asp -0.98 Da), ^18^O labeling (Asp +2.00 Da), and Oxidation (Met +15.99 Da); peptide tolerance 10 ppm; and MS/MS tolerance 0.02 Da.

The Endo H treated Mass spectrometric data were analyzed using pFind (version 3.2.0) ^36^ to search against the *de novo* sequencing output sequences, with potential Asn to Asp conversions. The parameters included: enzyme, partial pepsin; missed cleavages allowed, two; fixed modification, Carbamidomethylation (Cys +57.02 Da); variable modifications, Asp-Asn substitute (Asp -0.98 Da), GlcNAc (Asp +202.10 Da), and Oxidation (Met +15.99 Da); precursor tolerance 20 ppm; and fragment tolerance 20 ppm.

### Data process for *N*-glycopeptide identification

*N*-glycopeptide identification was performed with Biopharma Finder (version 5.2, Thermo Fisher Scientific, Waltham, MA) to search against the *de novo* sequencing output sequences with corrected Asn-Asp conversion. The following parameters were used: carbamidomethyl (Cys +57.02 Da) as the fixed modification and oxidation (Met +15.99Da); Lys loss on C terminal (−128.095 Da) and the built-in N-glycan repertoire for Chinese hamster ovary cell lines as the variable modification; precursor tolerance 20 ppm; and fragment tolerance 20 ppm.

### Data process for *O*-glycopeptide identification

For *O*-glycosylation site identification, *O*-glycan data were analyzed by pGlyco (version 3.1)^37^. The parameters of pGlyco included semi-specific enzyme on a sample-by-sample basis, allowing for up to five missed cleavages. The fixed modification was carbamidomethyl (Cys +57.02 Da) and the variable modification was oxidation (Met +15.99 Da). Peptide length ranges from 6 to 40 amino acids, with a precursor tolerance of 10 ppm and a fragment tolerance of 20 ppm. The glycopeptide FDR was set at 0.01. The *O*-glycan database consisted predominantly of *O*-glycans expressed in CHO cells, with the inclusion of reported *O*-glycans specific to etanercept. The glycan score threshold was set to ≥ 5.

## Data availability

The mass spectrometry proteomics data have been deposited to the ProteomeXchange Consortium via the PRIDE^38^ partner repository (https://www.ebi.ac.uk/pride/archive/) with the dataset identifier XXX.

## Acknowledgments

This work is supported by the Strategic Priority Research Program of the Chinese Academy of sciences, Grant No. XDB0850000. We thank Dr. Zhibiao Mai and Prof. Gong Zhang, for the instruction in the use of MuCS software and protein assemble.

## Author contributions

J. Gao, H.X. Chen, H.R. Yin, L.Q. Wen, and H. Zhou designed the study, prepared and revised the manuscript. J. Gao, H.X. Chen, H.R. Yin, X. Chen, Z.C. Yang, Y.Q. Wang, Y.P. Tian performed the experiments and analyzed the experimental data. J.H. Wu provided technical and material support. L.Q. Wen, and H. Zhou supervised the studies. All authors have read and approved the article.

