## Supplementary for "Decoding protein glycosylation by an integrative mass spectrometry-based *de novo* sequencing strategy"

```
1   MKSFVLLFCLAQLWGCHSIPLDPVAGYKEPACDDPDTEQAALAAVDYINKHLPRGYKHTL
61  NQIDSVKVWPRRPTGEVYDIEIDTLETTCHVLDPTPLANCSVRQQTQHAVEGDCDIHVLK
121 QDGQFSVLFTKCDSSPDSAEDVRKLCPDCPLLAPLNDSRVVHAVEVALATFNAESNGSYL
181 QLVEISRAQFVPLPVSVSVEFAVAATDCIAKEVVDPTKCNLLAEKQYGFCKGSVIQKALG
241 GEDVRVTCTLFQTQPVIPQPQPDGAEAEAPSAVPDAAGPTPSAAGPPVASVVVGPSVVAV
301 PLPLHRAHYDLRHTFSGVASVESSSGEAFHVGKTPIVGQPSIPGGPVRLCPGRIRYFKI
```

**Supplementary Fig. 1.** Amino acid sequence of Fetuin A. *N*-glycosylation sites are indicated in green; *O*-glycosylation sites of serine and threonine are shown in orange and blue, respectively. The *N*-terminal signal peptide, which is in the grey box, is absent in the most abundant proteoform.

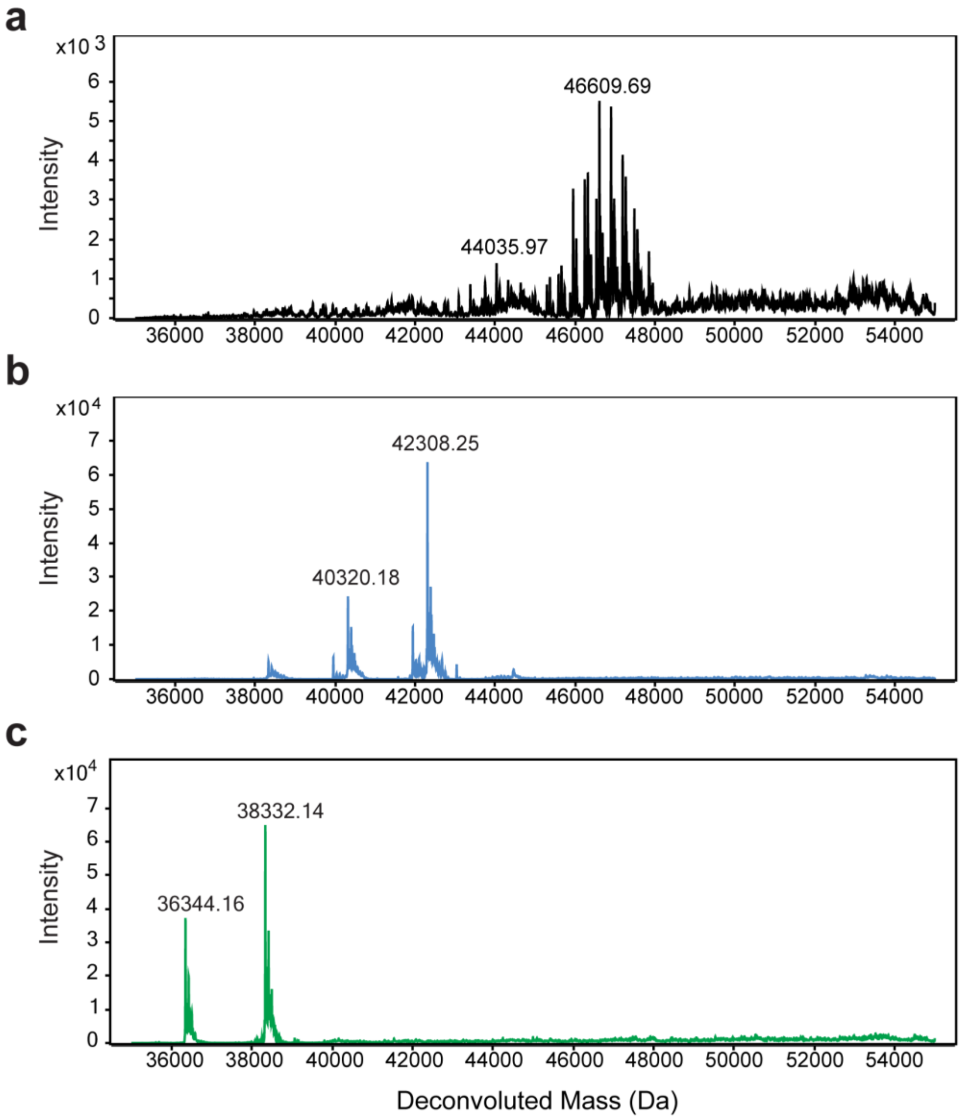

**Supplementary Fig. 2**. Deconvoluted ESI Q-TOF sepctra of Fetuin. (a) Intact mass of untreated Fetuin control. (b) Intact mass of Fetuin incubated with sialidase and EngEF. (c) Intact mass of Fetuin incubated with three endoglycosidases mixture of PNGase F, EngEF and sialidase.

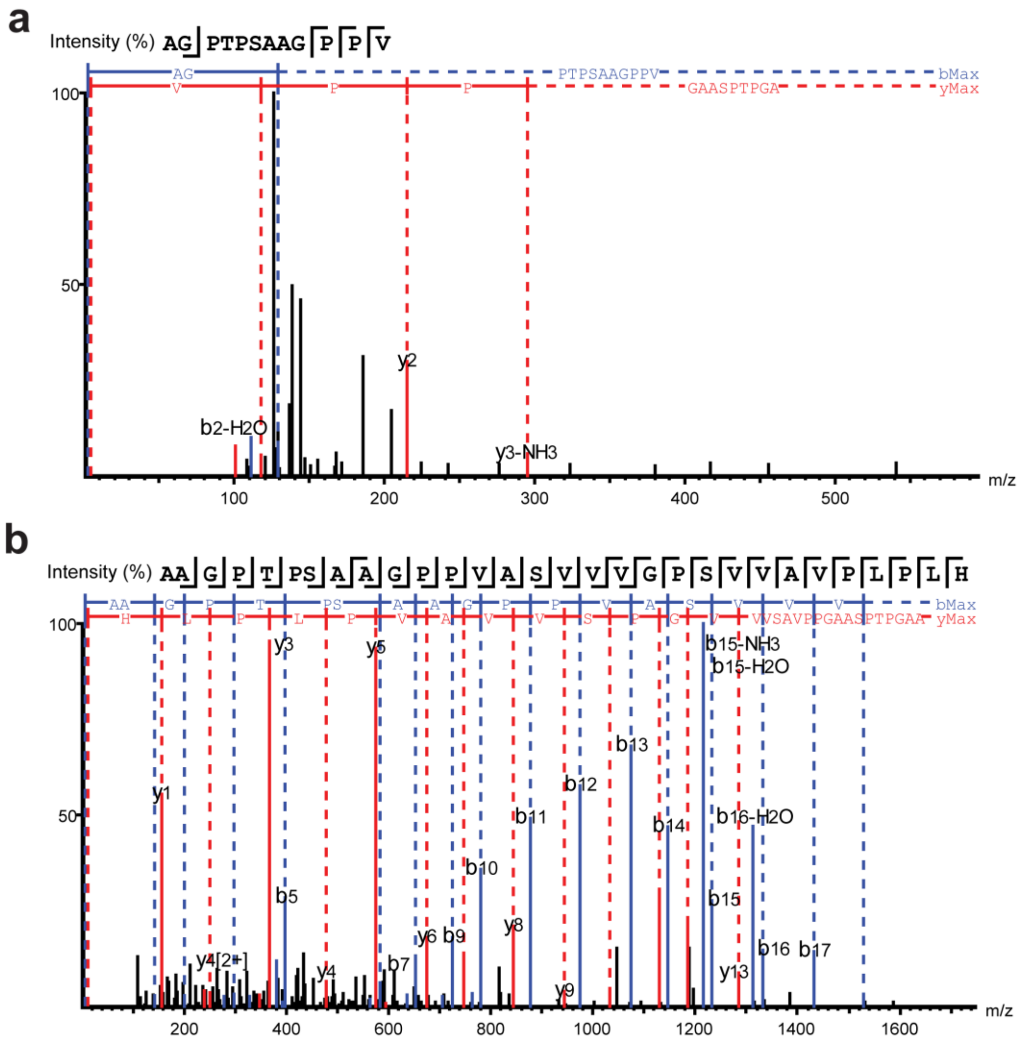

**Supplementary Fig. 3**. Release of *N*-glycans and *O*-glycans aid in the MS/MS determination of the peptide sequence. (a) A sample tandem mass spectrum of an identified peptide (AGPTPSAAGPPV) within amino acids region 241-280 of Fetuin A without deglycosylation. (b) A sample tandem mass spectrum of an identified peptide (AAGPTPSAAGPPVASVVVGPSVVAVPLPLH) within amino acids region 241-280 of Fetuin A following deglycosylation.

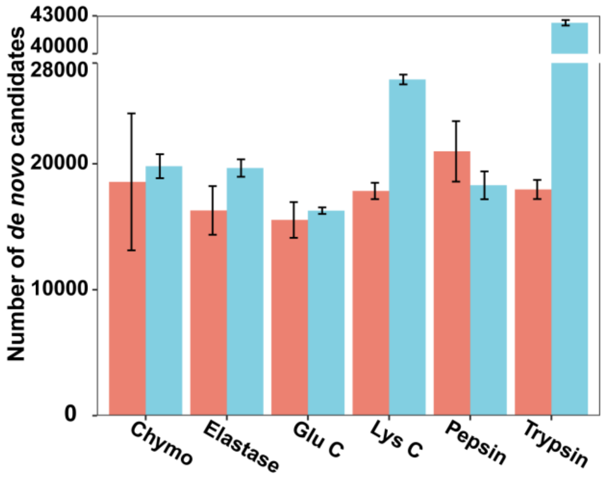

**Supplementary Fig. 4**. Comparison of number of *de novo* candidates acquired under EThcD fragmentation (Red) and stepped HCD fragmentation (Blue). Each experiment was performed 3 replications.

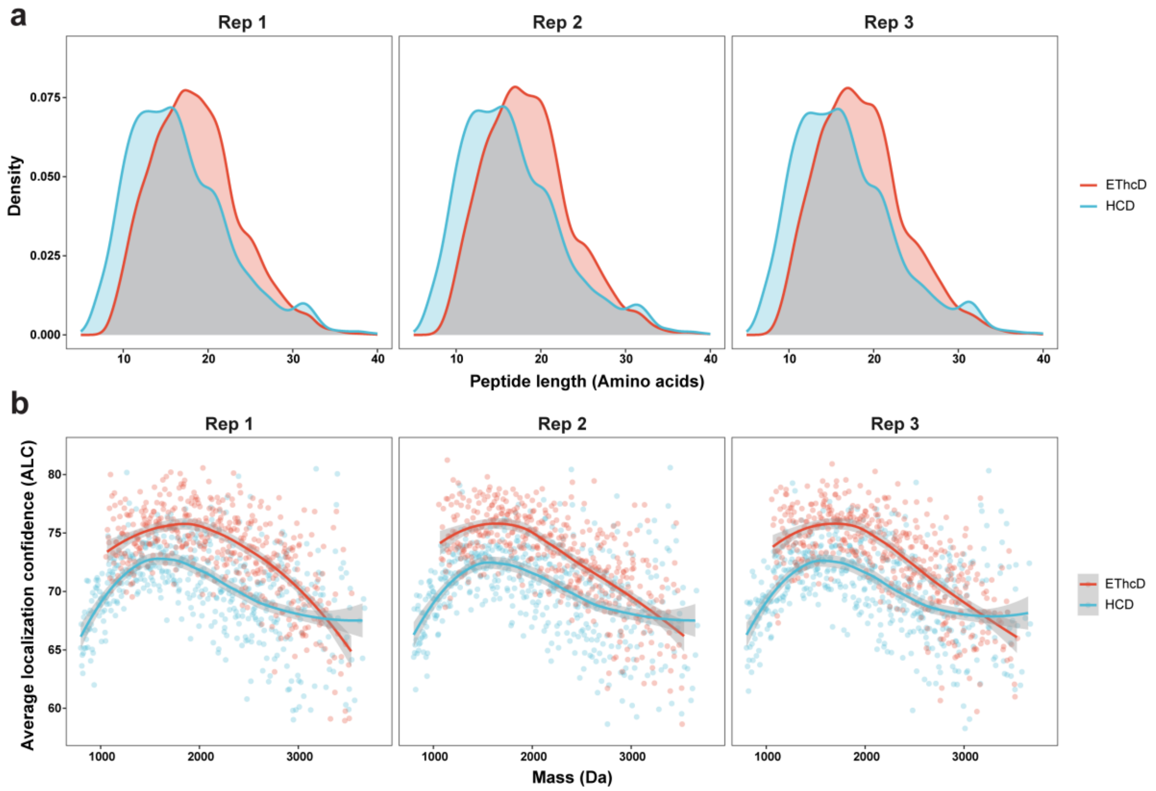

**Supplementary Fig. 5**. Comparison of *de novo* peptide sequencing performance of EThcD fragmentation versus HCD fragmentation. (a) Density plot of peptide length. (b) Scatter plot of average localization confidence (ALC) of *de novo* peptide candidates. Each experiment was performed 3 replications.

| | |
|---|---|
| *FETUA* | ------IPLDPVAGYKEPACDDPDTEQAALAAVDYINKHLPRGYKHTLNQIDSVKVWPRRPTGEVYDIEIDTLETTCHVLDPTPLANCSVRQQTQHAVEG |
| *Rep1* | HAVEGDLPLDPVAGYKEPACDDPDTEQAALAAVDYINKHLPRGYKHTLNQIDSVKVWPRRPTGEVYDIEIDTLETTCHVLDPTPLADCSVRQQTQHAVEG |
| *Rep2* | HAVEGDLPLDPVAGYKEPACDDPDTEQAALAAVDYINKHLPRGYKHTLNQIDSVKVWPRPRTGEVYDIEIDTLETTCHVLDPTPLADCSVRQQTQHAVEG |
| *Rep3* | -AVEGDLPLDPVAGYKEPACDDPDTEQAALAAVDYINKHLPRGYKHTLNQIDSVKVWPRPRTGEVYDIEIDTLETTCHVLDPTPLADCSVRQQTQHAVEG |
| *FETUA* | DCDIHVLKQDGQFSVLFTKCDSSPDSAEDVRKLCPDCPLLAPLNDSRVVHAVEVALATFNAESNGSYLQLVEISRAQFVPLPVSVSVEFAVAATDCIAKE |
| *Rep1* | DCDIHVLKQDGQFSVLFTKCDSSPDSAEDVRKLCPDCPLLAPLDDSRVVHAVEVALATFNAESDGSYLQLVEISRAQFVPPLVSVSVEFAVAATDCIAKE |
| *Rep2* | DCDIHVLKQDGQFSVLFTKCDSSPDSAEDVRKLACSVHVLAPLDDSRVVHAVEVALATFNAESDGSYLQLVEISRAQFVPLPVSVSVEFAVAATDCIAKE |
| *Rep3* | DCDIHVLKQDGQFSVLFTKCDSSPDSAEDVRKLCPDCPLLAPLDDSRVVHAVEVALATFNAESDGSYLQLVEISRAQFVPPLVSVSVEFAVAATDCIAKE |
| *FETUA* | VVDPTKCNLLAEKQYGFCKGSVIQKALGGEDVRVTCTLFQTQPVIPQPQPDGAEAEAPSAVPDAAGPTPSAAGPPVASVVVGPSVVAVPLPLHRAHYDLR |
| *Rep1* | VVDPTKCNLLAEKQYGFCKGSVIQKALGGEDVRVTCTLFQTQPVIPQPQPDGAEAEAPSAVPDAAGPTPSAAGPPVASVVVGPSVVAVPLPLHRAHYDLR |
| *Rep2* | VVDPTKCNLLAEKQYGFCKGSVIQKALGGEDVRVTCTLFQTQPVIPQPQPDGAEAEAPSAVPDAAGPTPSAAGPPVASVVVGPSVVAVPLPLHRAHYDLR |
| *Rep3* | VVDPTKCNLLAEKQYGFCKGSVIQKALGGEDVRVTCTLFQTQPVIPQPQPDGAEAEAPSAVPDAAGPTPSAAGPPVASVVVGPSVVAVPLPLHRAHYDLR |
| *FETUA* | HTFSGVASVESSSGEAFHVGKTPIVGQPSIPGGPVRLCPGRIRYFKI |
| *Rep1* | HTFSGVASVESSSGEAFHVGKTPIVGQPSIPGGPVRLCPGRIRYFKK |
| *Rep2* | HTFSR-ASVESSSGEAFHVGKTPIVGQPSIPGGPVRLPCGRIRYFKK |
| *Rep3* | HTFSR-ASVESSSGEAFHVGKTPIVGQPSIPGGPVRLCPGRIRYFKK |

**Supplementary Fig. 6**. Assemble results of deglycosylated Fetuin sequenced by EThcD fragmentation method. Integration of sequencing results from multiple proteases (LysC, Elestase, Pepsin, Chymotrypsin, GluC, Trypsin) digestion were assembled into scaffolds by MuCS. The output sequences are aligned to the known sequence. Substitution of amino acids and residue swap are highlighted in blue and involved in the accuracy calculations. N-D conversions by PNGase F mediate *N*-Glycan releasing were highlighted in green. Insertion of amino acids are highlighted in red.

```
FETUA         ------IPLDPVAGYKEPACDDPDTEQAALAAVDYINKHLPRGYKHTLNQIDSVKVWPRRPTGEVYDIEIDTLETTCHVLDPTPLANCSVRQQTQHAVEG
Rep1_contig1  TCHVLDLPLDPVAGYKEPACDDPDTEQAALAAVDYLNKHLPRGYKHTLNQLDSVKVWPRRPTGEVYDDELDTLETTCHVLDPTPLADCSVRQQTQHAVEG
Rep1_contig2  ----------------------------------------------------------------------------------------------------
Rep2_contig1  TCHVLDLPLDPVAGYKEPACDDPDTEQAALAAVDYLNKHLPRGYKHTLNQLDSVKVWPRRPTGEVYDLELDTLETTCHVLDPTPLADCSVRQQTQHAVEG
Rep2_contig2  ----------------------------------------------------------------------------------------------------
Rep3_contig1  TCHVLDLPLDPVAGYKEPACDDPDTEQAALAAVDYLNKHLPRGYKHTLNQLDSVKVWPRRQ-QQVYDLELDTLETTCHVLDPTPLADCSVRQQTQHAVEG
Rep3_contig2  ----------------------------------------------------------------------------------------------------

FETUA         DCDIHVLKQDGQFSVLFTKCDSSPDSAEDVRKLCPDCPLLAPLNDSRVVHAVEVALATFNAESNGSYLQLVEISRAQFVPLPVSVSVEFAVAATDCIAKE
Rep1_contig1  DCDLHVLKQDGQFSVLFTKCDSSPDSAEDVRKLCPDCPLLAPLDDSRVVHAVEVALATFNAESDGSYLQLVELSRAQFVPLPVSVSVEFAVAATDCLAKE
Rep1_contig2  ----------------------------------------------------------------------------------------------------
Rep2_contig1  DCDLHVLKQDGQFSVLFTKCDSSPDSAEDVRKLCPDCPLLAPLDDSRVVHAVEVALATFNAESDGSYLQLVELSRAQFVPLPVSVSVEFAVAATDCLAKE
Rep2_contig2  ----------------------------------------------------------------------------------------------------
Rep3_contig1  DCDLHVLKQDGQFSVLFTKCDSSPDSAEDVRKLCPDCPLLAPLDDSRVVHAVEVALATFNAESDGSYLQLVELSRAQFVPLPVSVSVEFAVAATDCLAKE
Rep3_contig2  ----------------------------------------------------------------------------------------------------

FETUA         VVDPTKCNLLAEKQYGFCKGSVIQKA-LGGEDVRVTCTLFQTQPVIPQPQPDGAEAEAPSAVPDAAGPTPSAAGPPVASVVVGPSVVAVPLPLHRAHYDL
Rep1_contig1  VVDPTKCNLLAEKQYGFCKGSVLQKA-LGGEDVRVTCTLFQTQPVLPQPQPDGAEARSQ--PLVPLFQGPG-----------------------------
Rep1_contig2  ------------------------------------WPYYVAGVPVP--KP-GV---VVSAV--PPGAASPAAGPPVASVVVGPSVVAVPLPLHRAHYDL
Rep2_contig1  VVDPTKCARLAEKQYGFCKGSVLGAGKALGWDVRVTCTLFQTQPVLPQPQPDGAEATNNY--LVPQTQFPVLPQ--------------------------
Rep2_contig2  ----------------------------------DPTDEQHKQPVLPQEPP-NQEAEAPSAVPDAAGPTPSAAGPPVASVVVGPSVVAVPLPLHRAHYDL
Rep3_contig1  VVDPTKCNLLAEKQYGFCKGSVLQKA-LGGEDVRVTCTLFQTQPVLPQPQPDGAEAEADKQPLVPA--GTAW----------------------------
Rep3_contig2  ------------------------------------WWWYKTQPVLQPPQP-DQQAEAPSAVPDAAGPTPSAAGPPVASVVVGPSVVAVPLPLHRAHYDL

FETUA         RHTFSGVASVESSSGEAFHVGKTPIVGQPSIPGGPVRLCPGRIRYFKI-----------
Rep1_contig1  -----------------------------------------------------------
Rep1_contig2  RHTFSGVASVESSSGEAFHVGKTPLVGQPSLPGGPVRLCKPPAAATDTAADAHDKRHQK
Rep2_contig1  -----------------------------------------------------------
Rep2_contig2  RHTFSGVASVESSSEGAFHVGKTPLVGQPSLPGGPVRLCKPAAAPPSGLK---------
Rep3_contig1  -----------------------------------------------------------
Rep3_contig2  RHTFSGVASVESSSEGAFHVGKTPLVGQPSLPGGPVRLCKFLKKSGSGK----------
```

**Supplementary Fig. 7**. Assemble results of deglycosylated Fetuin sequenced by HCD fragmentation method. Integration of sequencing results from multiple proteases (LysC, Elestase, Pepsin, Chymotrypsin, GluC, Trypsin) digestion were assembled into scaffolds by MuCS. The output sequences are aligned to the known sequence. Substitution of amino acids and residue swap are highlighted in blue and involved in the accuracy calculations. N-D conversions by PNGase F mediate *N*-Glycan releasing were highlighted in green. Insertion of amino acids are highlighted in red.

a

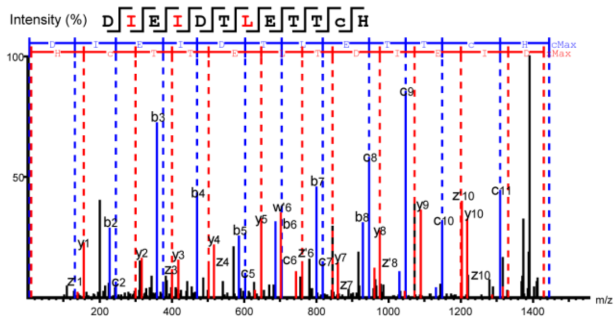

c d e

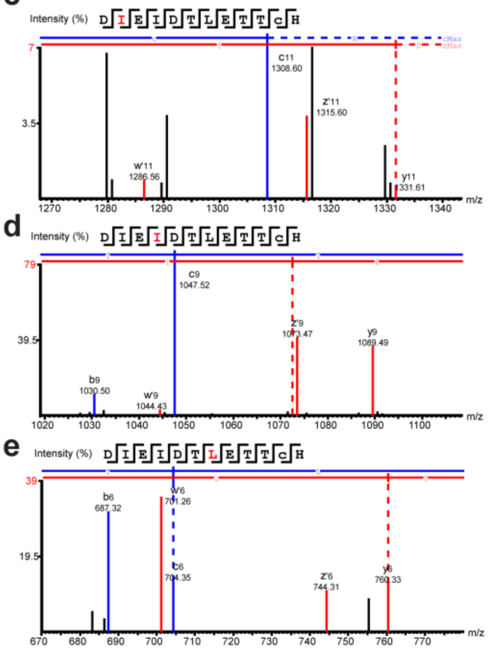

b

| # | b | c | c-H | Seq | y | z | z' | w | w' | # |
|---|---|---|---|---|---|---|---|---|---|---|
| 1 | 116.03 | 133.06 | 132.05 | D | | | | | | 12 |
| 2 | 229.12 | 246.15 | 245.14 | I | 1331.61 | 1314.59 | 1315.60 | 1285.55 | 1286.56 | 11 |
| 3 | 358.16 | 375.19 | 374.18 | E | 1218.53 | 1201.52 | 1202.52 | | | 10 |
| 4 | 471.25 | 488.27 | 487.26 | I | 1089.49 | 1072.48 | 1073.47 | 1043.42 | 1044.43 | 9 |
| 5 | 586.27 | 603.30 | 602.29 | D | 976.41 | 959.38 | 960.39 | | | 8 |
| 6 | 687.32 | 704.35 | 703.34 | T | 861.38 | 844.36 | 845.36 | | | 7 |
| 7 | 800.41 | 817.43 | 816.42 | L | 760.33 | 743.30 | 744.31 | 700.25 | 701.26 | 6 |
| 8 | 929.45 | 946.48 | 945.47 | E | 647.25 | 630.22 | 631.23 | | | 5 |
| 9 | 1030.50 | 1047.52 | 1046.51 | T | 518.21 | 501.18 | 502.18 | | | 4 |
| 10 | 1131.55 | 1148.57 | 1147.56 | T | 417.16 | 400.13 | 401.14 | | | 3 |
| 11 | 1291.57 | 1308.60 | 1307.59 | C(+57.02) | 316.11 | 299.08 | 300.09 | | | 2 |
| 12 | | | | H | 156.08 | 139.05 | 140.06 | | | 1 |

**Supplementary Fig. 8**. EThcD fragmentation enables Leu/Ile determination. (a) EThcD MS/MS spectra of peptide DIEIDTLETTCH. (b) Product ion list map of peptide DIEIDTLETTCH. The product ions matching the theoretical m/z values were highlighted in blue and red. (c) Zoomed-in MS/MS spectra contained diagnostic ion z'11+ ions and the diagnostic w'-ions resulting from neutral loss of $C_2H_5$ (-29 Da) which aids in identifying the presence of Ile at position 2 in the peptide. (d) Zoomed-in MS/MS spectra contained diagnostic ion z'9+ ions and the diagnostic w'-ions resulting from neutral loss of $C_2H_5$ (-29 Da) which aids in identifying the presence of Ile at position 4 in the peptide. (e) Zoomed-in MS/MS spectra contained diagnostic ion z'6+ ions and the diagnostic w'-ions resulting from neutral loss of $C_3H_7$ (-43 Da). This observation conclusively identifies residue 7 in the peptide as Leu.

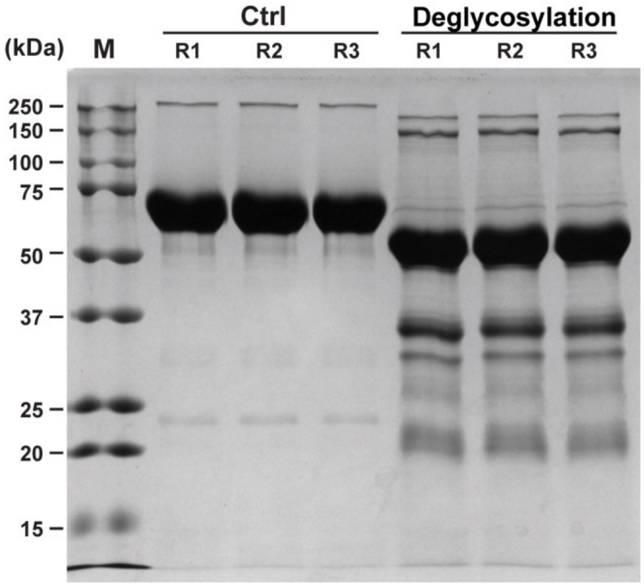

**Supplementary Fig. 9**. SDS-PAGE gel of Etanercept deglycosylation effect. lane M is the molecular weight standards. The control lane (Ctrl) with three replications represents the intact Etanercept control that without deglycosylation. The deglycosylation lane (Deglyco) with three replications represents Etanercept released *N*-/ *O*-glycan with a mixture of PNGase F, EngEF and sialidase.

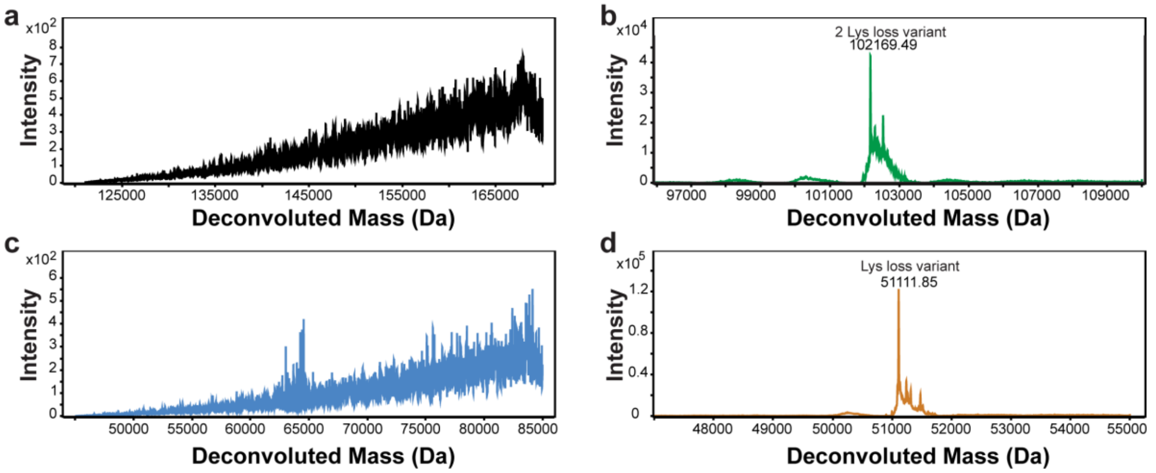

**Supplementary Fig. 10**. Deconvoluted ESI-QTOF mass spectrum of Etanercept. (a) Intact Etanercept without deglycosylation. (b) Etanercept after deglycosylation. (c) Etanercept without deglycosylation after reduction. (d) Etanercept after deglycosylation and reduction.

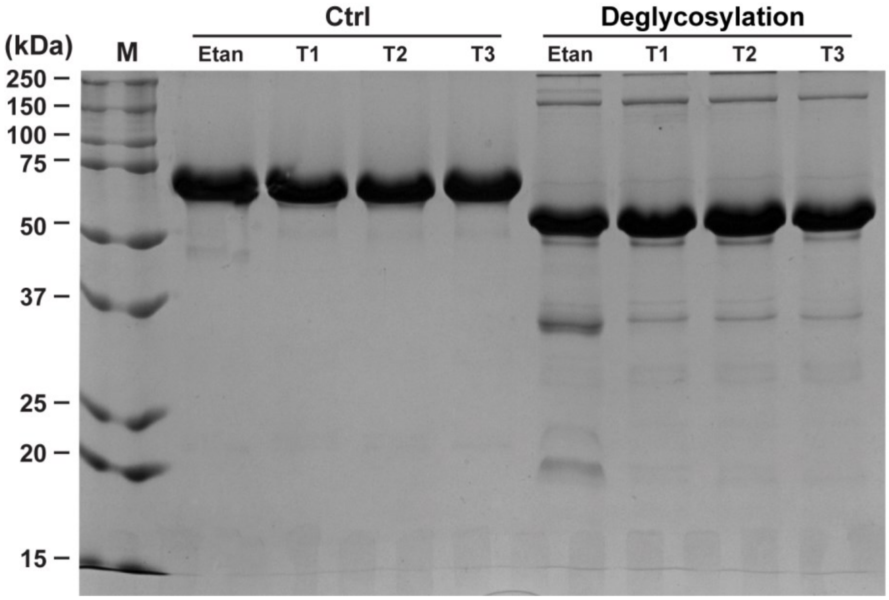

**Supplementary Fig. 11**. SDS-PAGE gel of TNFR: Fc-fusion proteins deglycosylation effect. lane M is the molecular weight standards. The control lane (Ctrl) represents four untreated controls, including Etanercept, T1 (TNFR: Fc 1), T2 (TNFR: Fc 2) and T3 (TNFR: Fc 3). The deglycosylation lane (Deglyco) represents deglycosylated samples, including Etanercept, T1 (TNFR: Fc 1), T2 (TNFR: Fc 2) and T3 (TNFR: Fc 3).

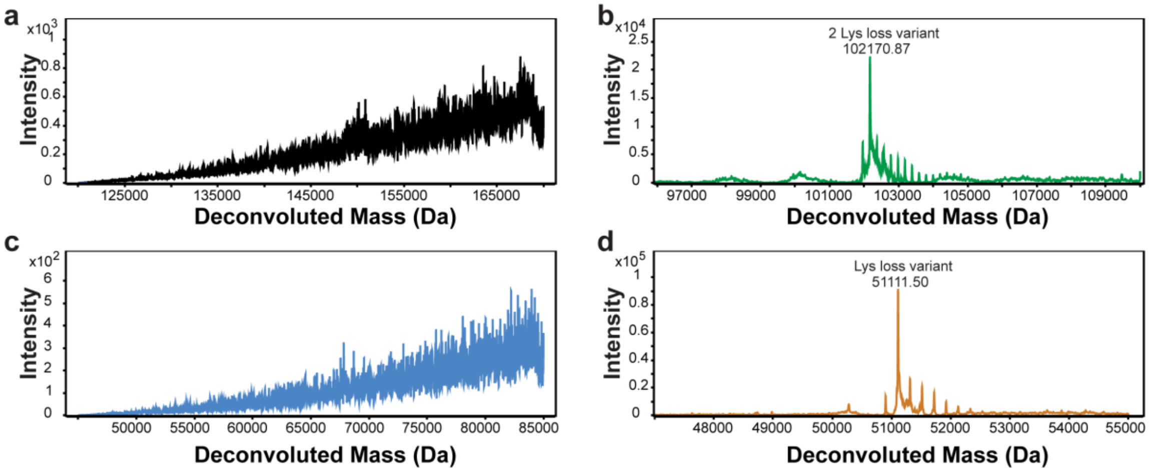

**Supplementary Fig. 12**. Deconvoluted ESI-QTOF mass spectrum of TNFR: Fc 1 fusion protein. (a) Intact TNFR: Fc 1 without deglycosylation. (b) TNFR: Fc 1 after deglycosylation. (c) TNFR: Fc 1 after reduction without deglycosylation. (d) TNFR: Fc 1 after deglycosylation and reduction.

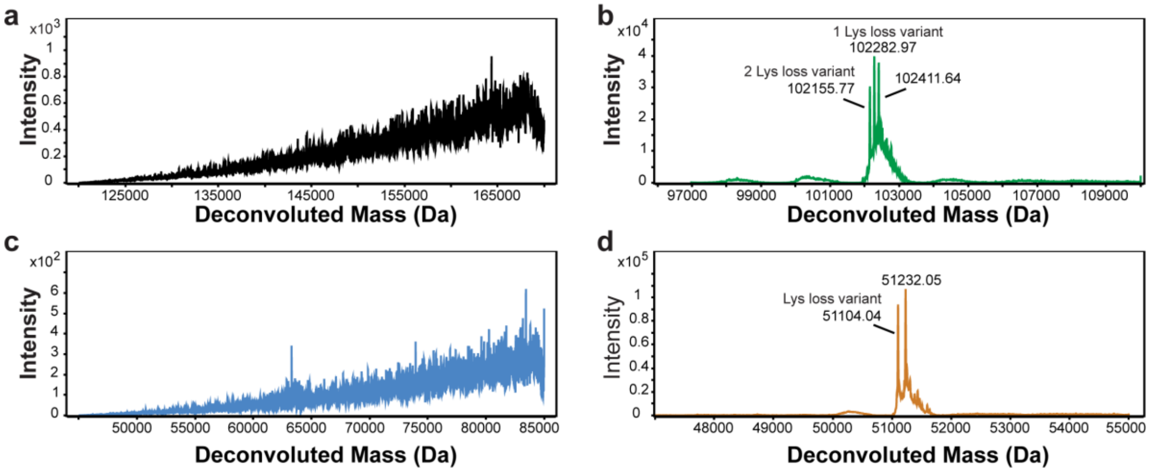

**Supplementary Fig. 13**. Deconvoluted ESI-QTOF mass spectrum of TNFR: Fc 2 fusion protein. (a) Intact TNFR: Fc 2 without deglycosylation. (b) TNFR: Fc 2 after deglycosylation. (c) TNFR: Fc 2 after reduction without deglycosylation. (d) TNFR: Fc 2 after deglycosylation and reduction.

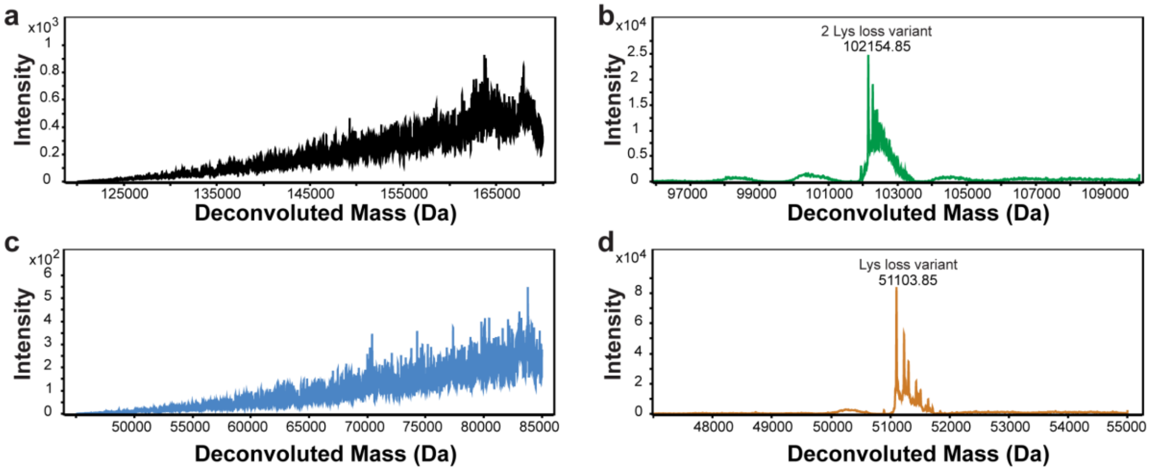

**Supplementary Fig. 14**. Deconvoluted ESI-QTOF mass spectrum of TNFR: Fc 3 fusion protein. (a) Intact TNFR: Fc 3 without deglycosylation. (b) TNFR: Fc 3 after deglycosylation. (c) TNFR: Fc 3 after reduction without deglycosylation. (d) TNFR: Fc 3 after deglycosylation and reduction.

```
Etanercept   -----LPAQVAFTPYAPEPGSTCRLREYYDQTAQMCCSKCSPGQHAKVFCTKTSDTVCDSCEDSTYTQLWNWVPECLSCGSRCSSDQVETQACTREQNRI
Original_1   -----LPAQVAFTPYAPEPGSTCRLREYYDQTAQMCCSKCSPGQHAKVFCTKTSDTVCDSCEDSTYTQLWNWVPECLSCGSRCSSDQVETQACTREQNRI
Original_2   VSHEDLPAQVAFTPYAPEPGSTCRLREYYDQTAQMCCSKCSPGQHAKVFCTKTSDTVCDSCEDSTYTQLWNWVPECLSCGSRCSSDQVETQACTREQNRI
Original_3   -EYYDLPAQVAFTPYAPEPGSTCRLREYYDQTAQMCCSKCSPGQHAKVFCTKTSDTVCDSCEDSTYTQLWNWVPECLSCGSRCSSDQVETQACTREQNRI

Etanercept   CTCRPGWYCALSKQEGCRLCAPLRKCRPGFGVARPGTETSDVVCKPCAPGTFSNTTSSTDICRPHQICNVVAIPGNASMDAVCTSTSPTRSMAPGAVHLP
Original_1   CTCRPGWYCALSKQEGCRLCAPLRKCRPGFGVARGPTETSDVVCKPCAPGTFSDTTSSTDICRPHQICNVVAIPGDASMDAVCTSTSPTRSMAPGAVHLP
Original_2   CTCRPGWYCALSKQEGCRLCAPLRKCRPGFGVARPGTETSDVVCKPCAPGTFSDTTSSTDICRPHQICNVVAIPGDASMDAVCTSTSPTRSMAPGAVHLP
Original_3   CTCRPGWYCALSKQEGCRLCAPLRKCRPGFGVARPPTETSDVVCKPCAPGTFSDTTSSTDICRPHQICNVVAIPGDASMDAVCTSTSPTRSMAPGAVHLP

Etanercept   QPVSTRSQHTQPTPEPSTAPSTSFLLPMGPSPPAEGSTGDEPKSCDKTHTCPPCPAPELLGGPSVFLFPPKPKDTLMISRTPEVTCVVVDVSHEDPEVKF
Original_1   QPVSTRSQHTQPTPEPSTAPSTSFLLPMGPSPPAEGSTGDEPKSCDKTHTCPPCPAPELLGGPSVFLFPPKPKDTLMISRTPEVTCVVVDVSHEDPEVKF
Original_2   QPVSTRSQHTQPTPEPSTAPSTSFLLPMGPSPPAEGSTGDEPKSCDKTHTCPPCPAPELLGGPSVFLFPPKPKDTLMISRTPEVTCVVVDVSHEDPEVKF
Original_3   QPVSTRSQHTQPTPEPSTAPSTSFLLPMGPSPPAEGSTGDEPKSCDKTHTCPPCPAPELLGGPSVFLFPPKPKDTLMISRTPEVTCVVVDVSHEDPEVKF

Etanercept   NWYVDGVEVHNAKTKPREEQYNSTYRVVSVLTVLHQDWLNGKEYKCKVSNKALPAPIEKTISKAKGQPREPQVYTLPPSREEMTKNQVSLTCLVKGFYPS
Original_1   NWYVDGVEVHNAKTKPREEQYDSTYRVVSVLTVLHQDWLNGKEYKCKVSNKALPAPIEKTISKAKGQPREPQVYTLPPSREEMTKNQVSLTCLVKGFYPS
Original_2   NWYVDGVEVHNAKTKPREEQYDSTYRVVSVLTVLHQDWLNGKEYKCKVSNKALPAPIEKTISKAKGQPREPQVYTLPPSREEMTKNQVSLTCLVKGFYPS
Original_3   NWYVDGVEVHNAKTKPREEQYDSTYRVVSVLTVLHQDWLNGKEYKCKVSNKALPAPIEKTISKAKGQPREPQVYTLPPSREEMTKNQVSLTCLVKGFYPS

Etanercept   DIAVEWESNGQPENNYKTTPPVLDSDGSFFLYSKLTVDKSRWQQGNVFSCSVMHEALHNHYTQKSLSLSPGK
Original_1   DIAVEWESNGQPENNYKTTPPVLDSDGSFFLYSKLTVDKSRWQQGNVFSCSVMHEALHNHYTQKSLS-----
Original_2   DIAVEWESNGQPENNYKTTPPVLDSDGSFFLYSKLTVDKSRWQQGNVFSCSVMHEALHNHYTQKSLSPKE--
Original_3   DIAVEWESNGQPENNYKTTPPVLDSDGSFFLYSKLTVDKSRWQQGNVFSCSVMHEALHNHYTQKSLSPKE--
```

**Supplementary Fig. 15**. Assemble results of deglycosylated Etanercept. Integration of sequencing results from multiple proteases (LysC, Elestase, Pepsin, Chymotrypsin, GluC, Trypsin) digestion were assembled into scaffolds by MuCS. The output sequences are aligned to the known sequence. Substitution of amino acids and residue swap are highlighted in blue and involved in the accuracy calculations. N-D conversions by PNGase F mediate *N*-Glycan releasing were highlighted in green. Insertion of amino acids are highlighted in red.

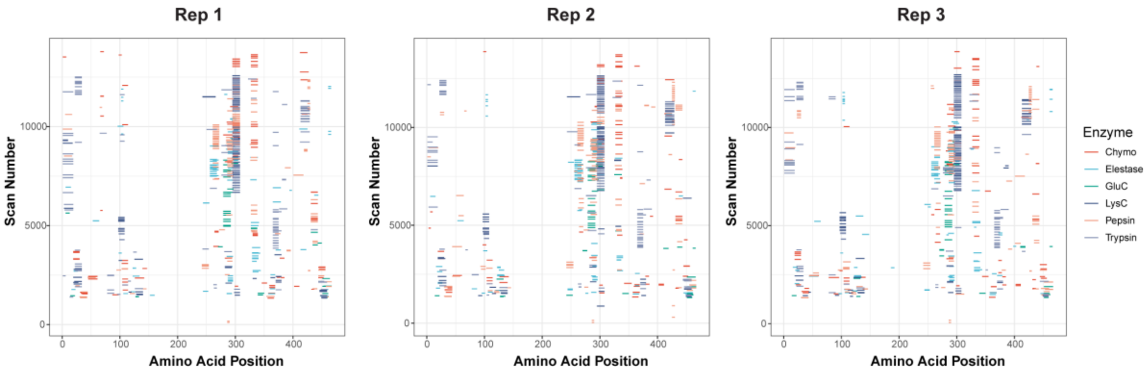

**Supplementary Fig. 16**. Peptide mapping coverage of Etanercept without deglycosylation.

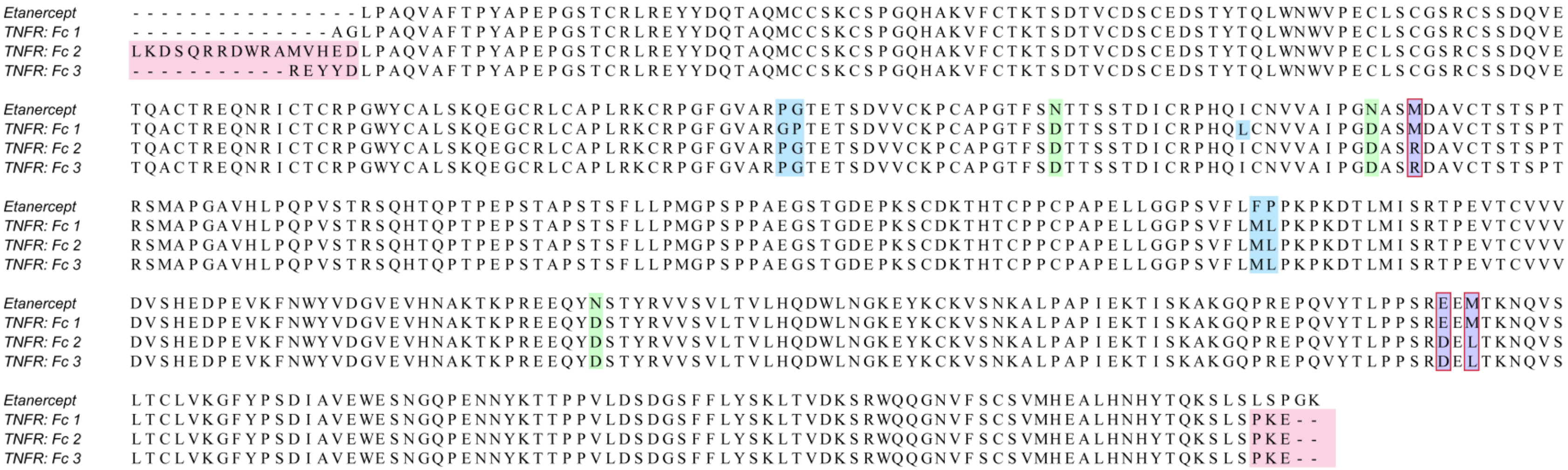

**Supplementary Fig. 17**. Assemble results of deglycosylated TNFR: Fc-fusion pharmasutical proteins. Integration of sequencing results from multiple proteases (LysC, Elestase, Pepsin, Chymotrypsin, GluC, Trypsin) digestion were assembled into scaffolds by MuCS. The output sequences are aligned to the known sequence of Etanercept. Variant of amino acids are highlighted in purple. Substitution of amino acids and residue swap are highlighted in blue. N-D conversions by PNGase F mediate *N*-Glycan releasing were highlighted in green. Insertion of amino acids are highlighted in red.

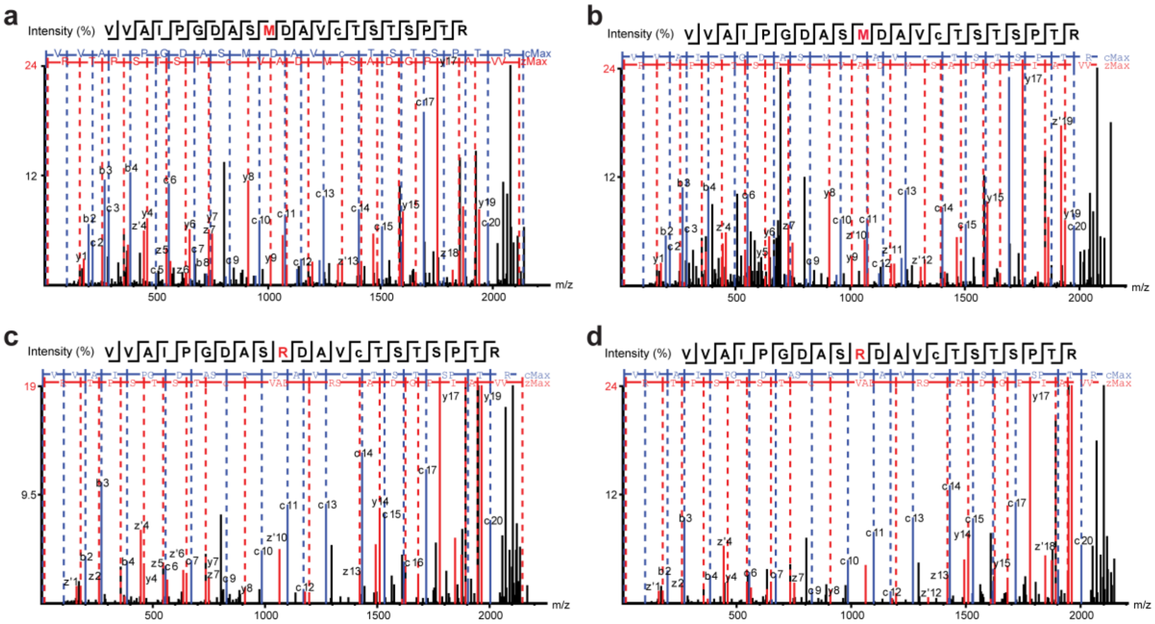

**Supplementary Fig. 18**. MS/MS spectra of variant peptide containing M174R. (a) Etanercept; (b) TNFR: Fc 1; (c) TNFR: Fc 2; (d) TNFR: Fc 3.

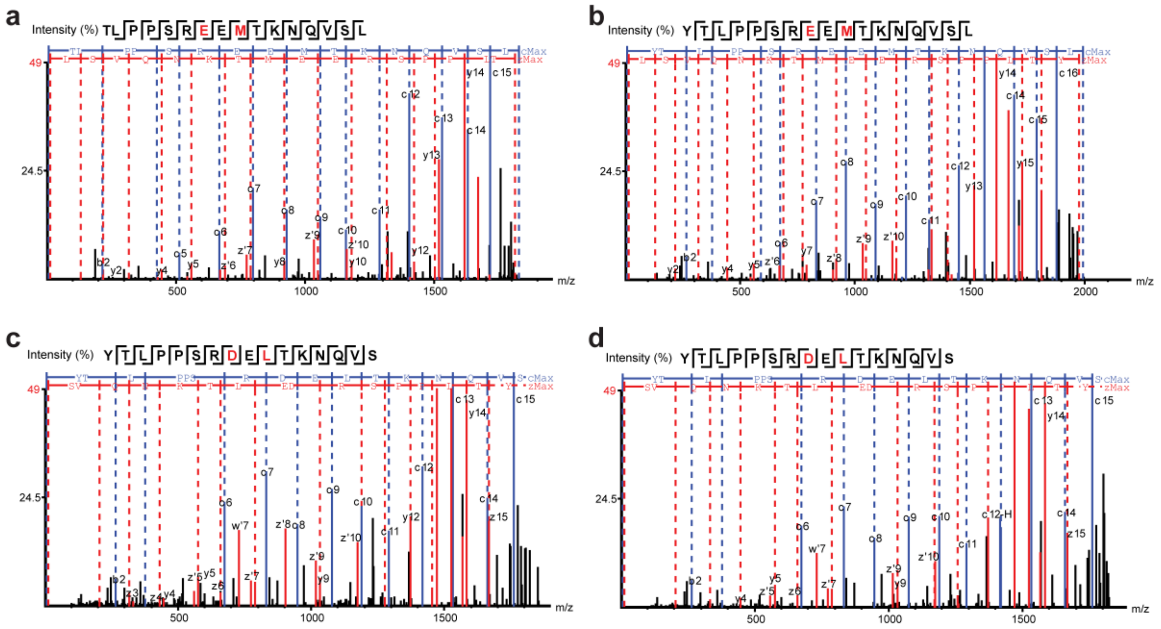

**Supplementary Fig. 19**. MS/MS spectra of variant peptide containing E376D, M378L. (a) Etanercept; (b) TNFR: Fc 1; (c) TNFR: Fc 2; (d) TNFR: Fc 3.

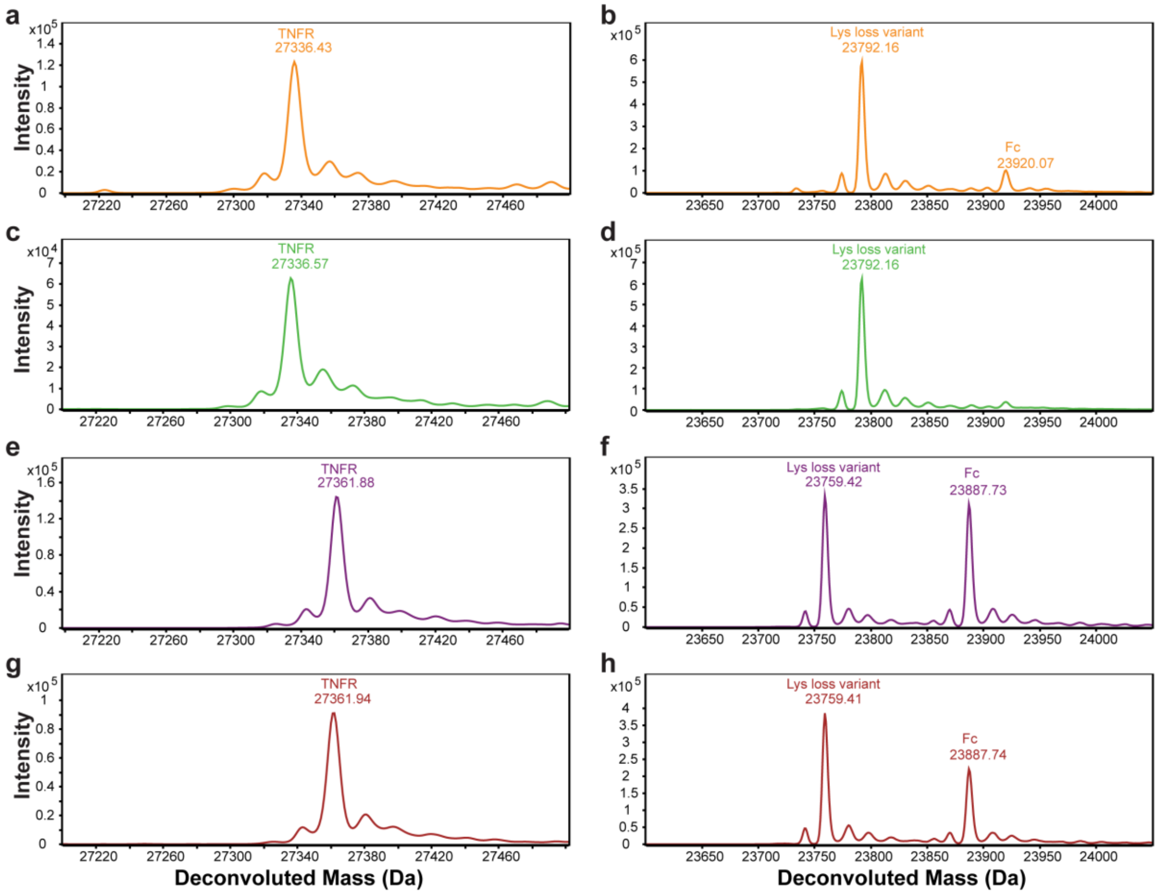

**Supplementary Fig. 20**. Deconvoluted mass spectra of deglycosylated TNFR and Fc domains. (a, b) Etanercept; (c, d) TNFR: Fc 1; (e, f) TNFR: Fc 2; (g, h) TNFR: Fc 3.

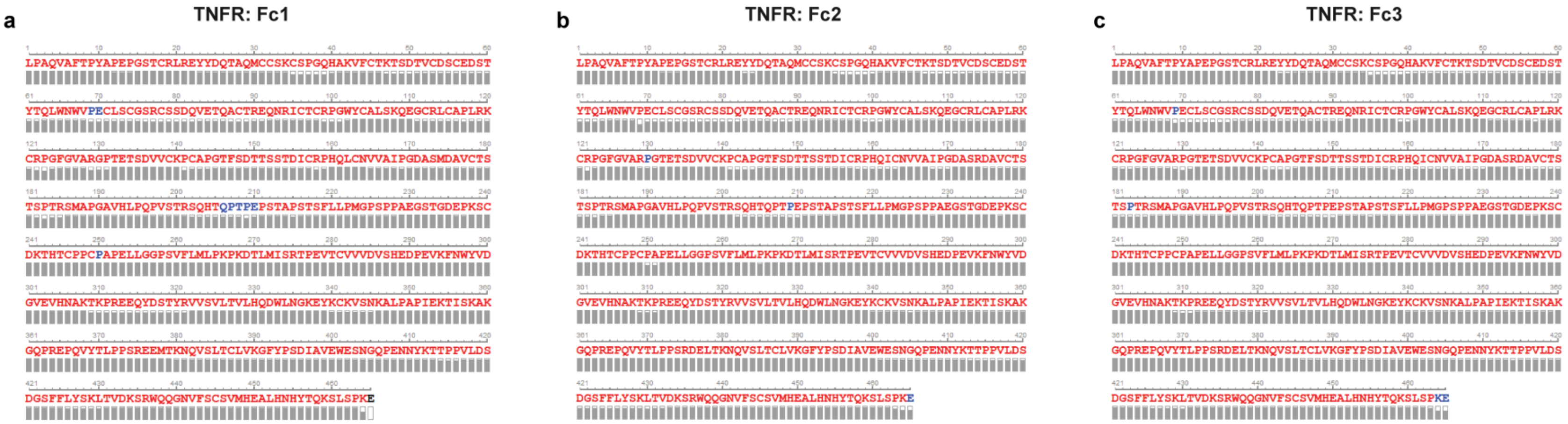

**Supplementary Fig. 21**. Sequence validation of deglycosylated TNFR: Fc-fusion pharmasutical proteins mapping with *de novo* sequencing output protein sequences. The fragment ion confident of each amino acid is indicated by colors, namely red for confidence>95%, blue for confidence > 85%, and black is for confidence <= 85%. The gray bars below the amino acid is the indicator of the number of matched MS/MS spectrum.

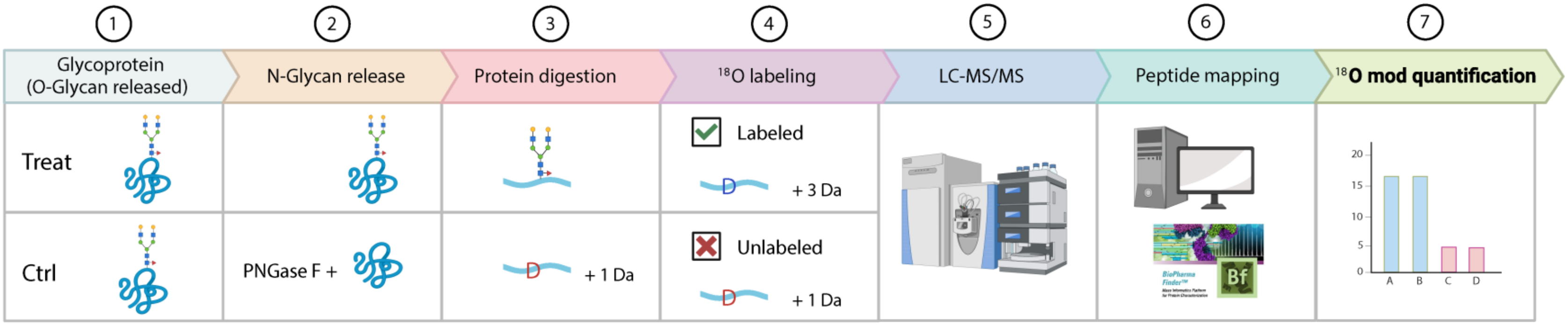

**Supplementary Fig. 22**. Workflow of quantitative $^{18}O$ labeling peptide mapping analysis for *N*-glycosylation site screening. *O*-glycans removed (treated with sialidase and EngEF) glycoprotein (Treat) was subjected to glycosylation site identification. Equal amount of glycoportein releasing both *N*- and *O*-glycans (treated with sialidase, EngEF and PNGase F) was set as a control (Ctrl). The proteins were digested into peptides with pepsin and thoroughly dried overnight. The peptides were labeled with $^{18}O$ by PNGase F mediated Asn linked *N*-glycans removing, resulting in 3 Da mass increasement in the Treat group while 1 Da mass increasement in the Ctrl group which only contain the Asn-Asp conversion without the $^{18}O$ labeling. Each experiment was performed 4 replications. The peptides were identified and quantified by LC-MS/MS in conjugated with peptide mapping data analysis work flow in Biopharma Finder software. The sites with $^{18}O$ modification ratio (Treat vs Ctrl) $\geq 2$ were considered as *N*-glycosylated candidates.

Etanercept_N149

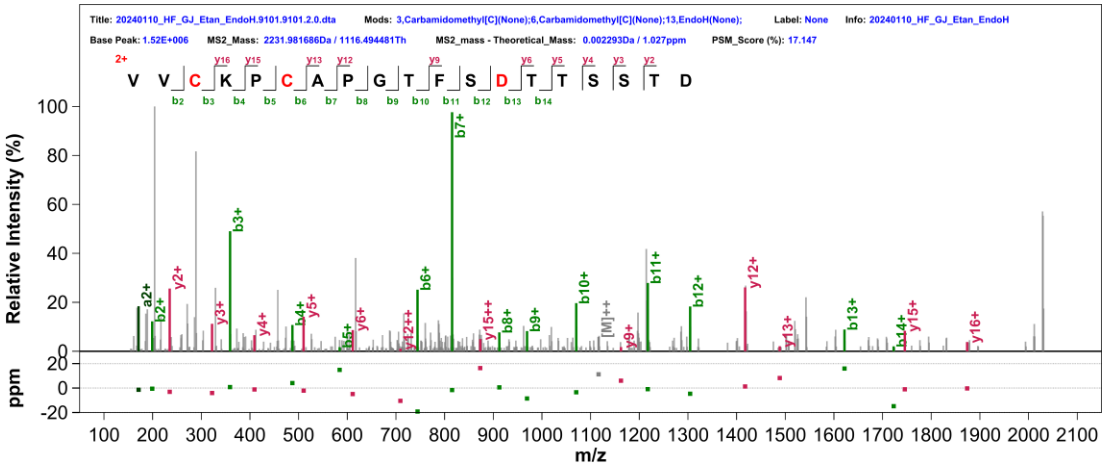

Etanercept_N171

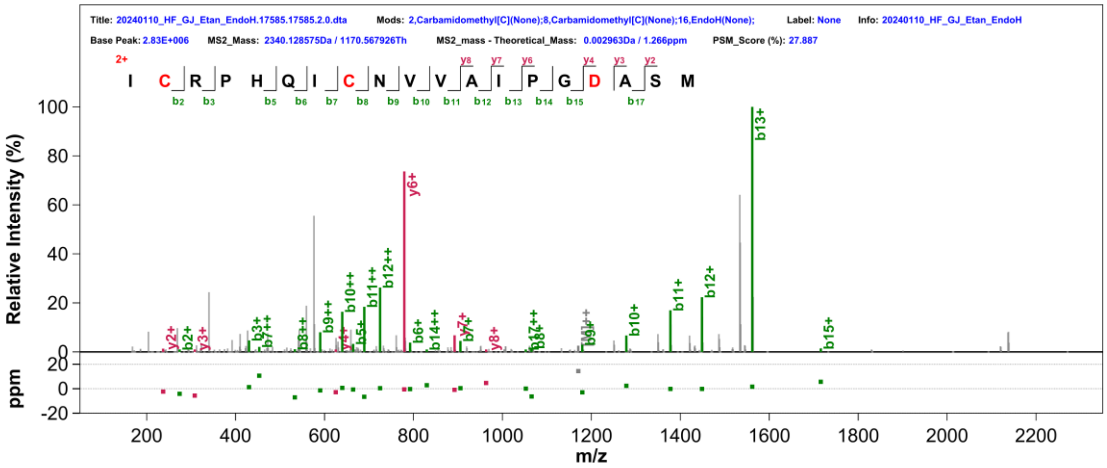

Etanercept_N317

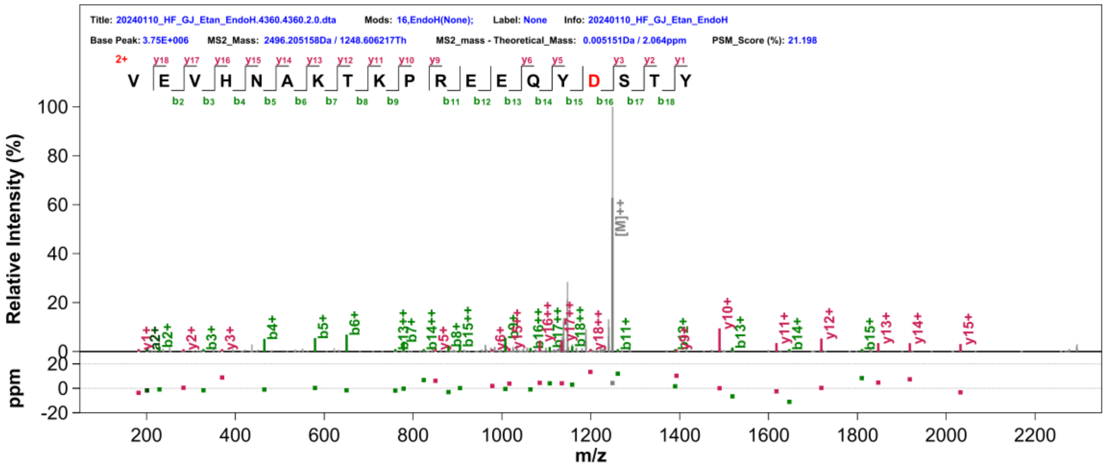

TNFR: Fc1_N149

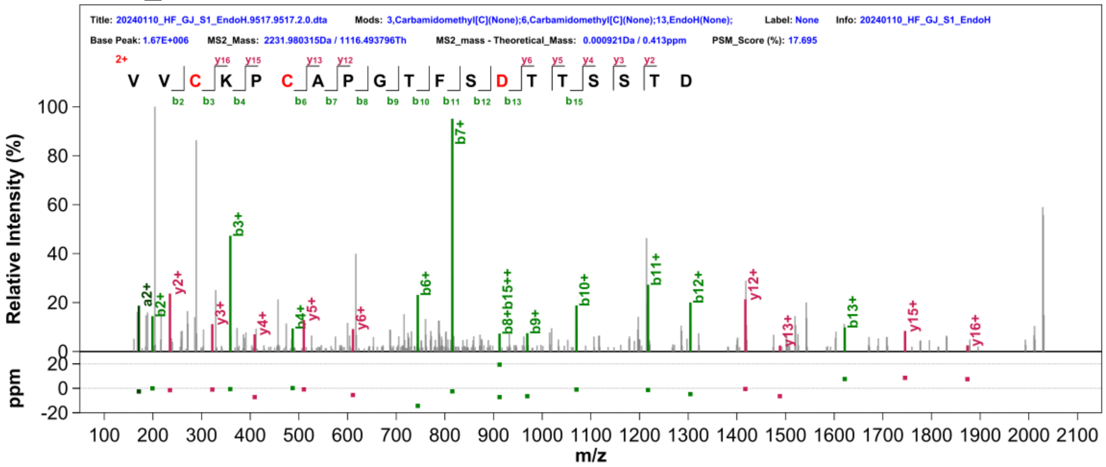

TNFR: Fc1_N171

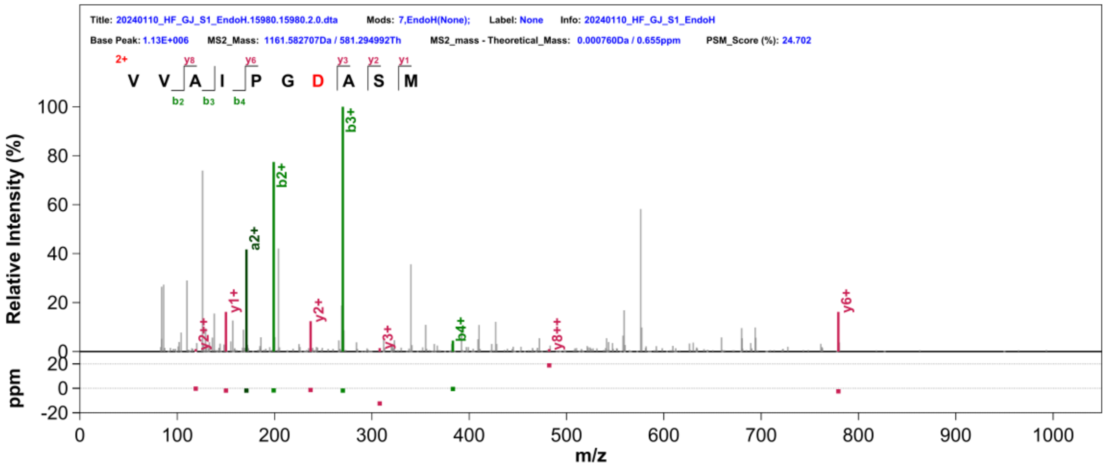

TNFR: Fc1_N317

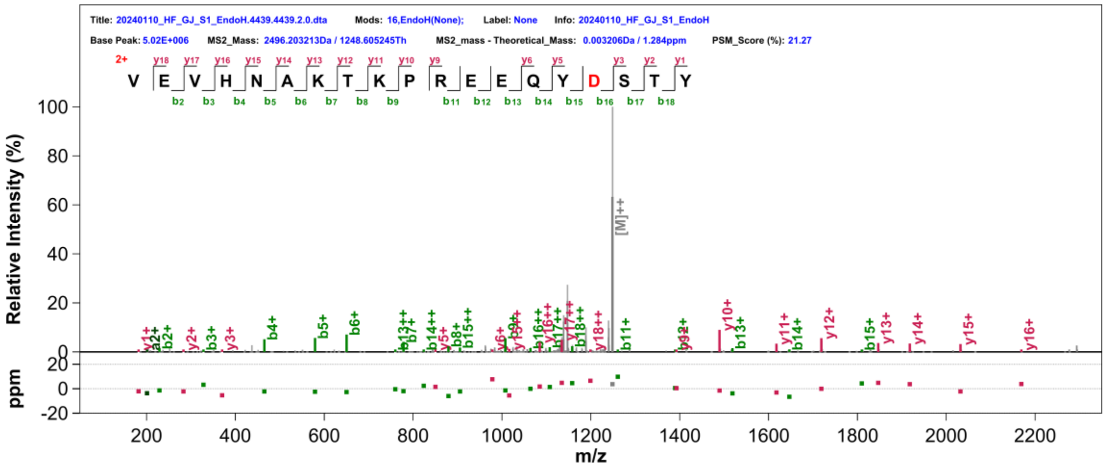

TNFR: Fc2_N149

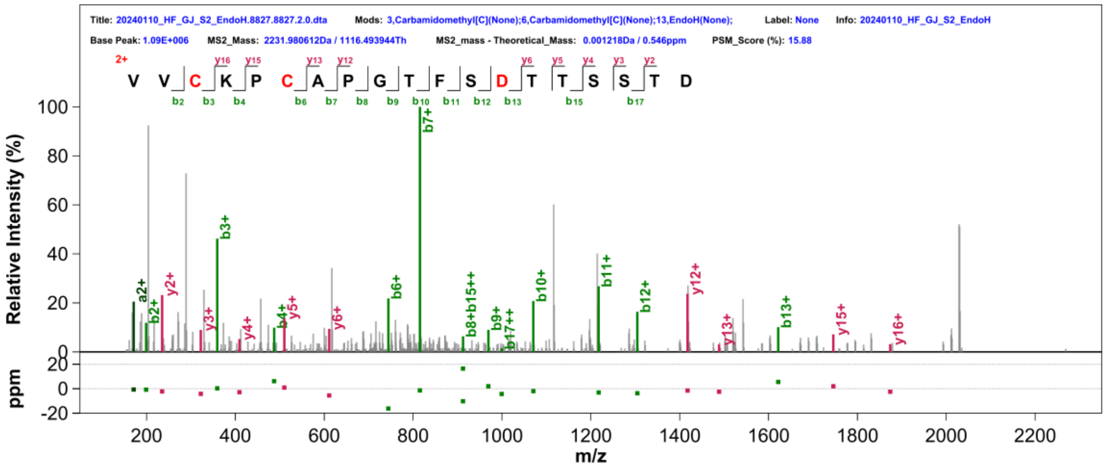

TNFR: Fc2_N171

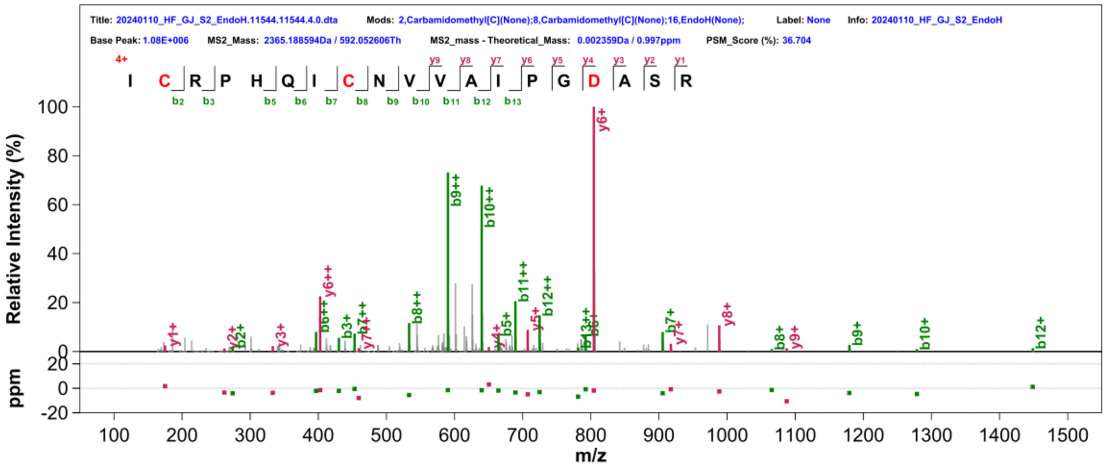

TNFR: Fc2_N317

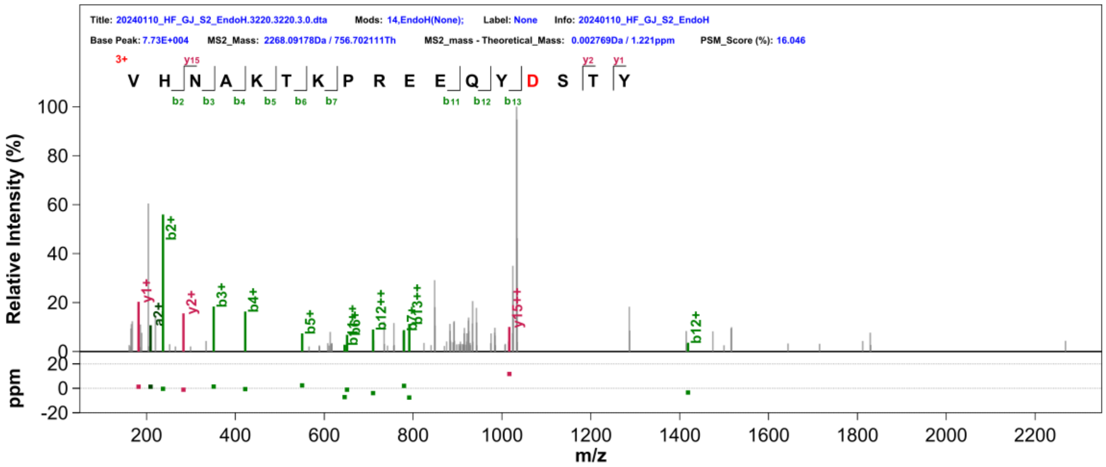

TNFR: Fc3_N149

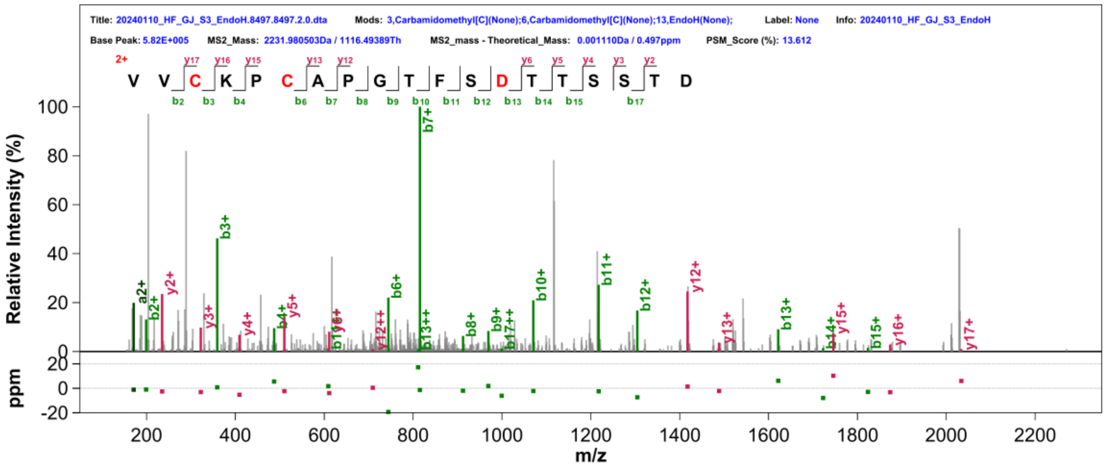

TNFR: Fc3_N171

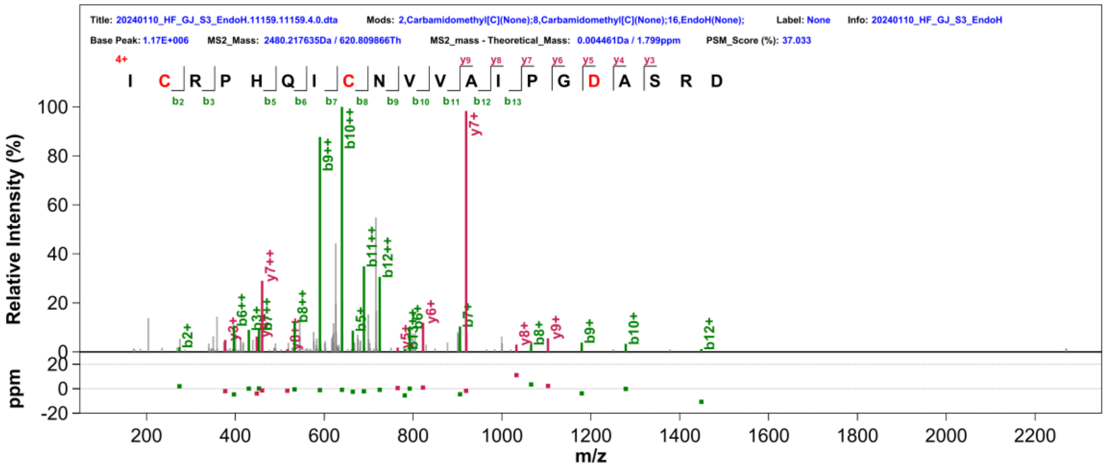

TNFR: Fc3_N317

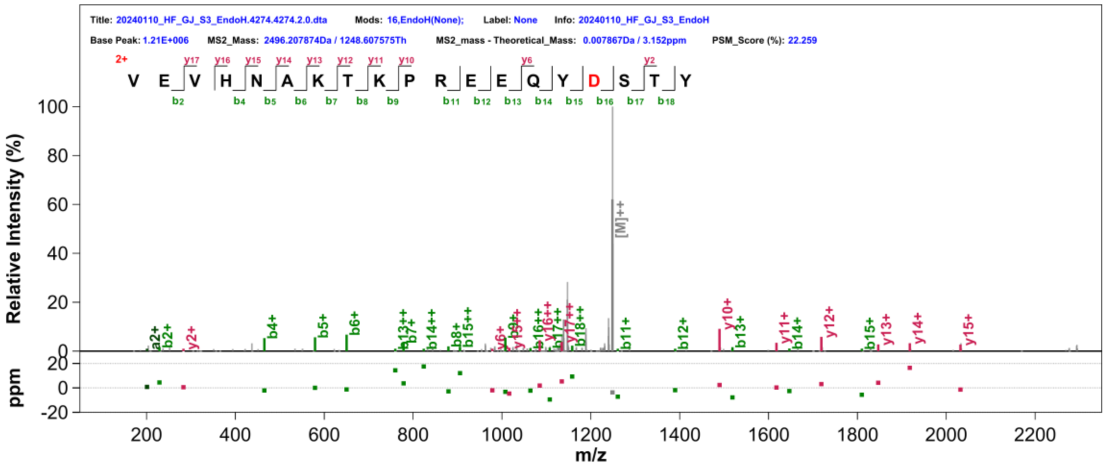

**Supplementary Fig. 23.** MS/MS spectra in support of the assigned *N*-glycosylation sites by Endo H treatment. The peptide sequence and fragment coverage are indicated in the top- left of each spectrum spectra, with b ions indicated in green, y ions indicated in magenta. The same color annotation is used for peaks in the spectra, with additional peaks such as intact/charge reduced precursors, neutral losses, and immonium ions indicated in grey. The *de novo* sequencing output sequences, with potential Asn (N) to Asp (D) conversions, was used for database searching. GlcNAc (D +202.10 Da) modification sites are indicated in red.

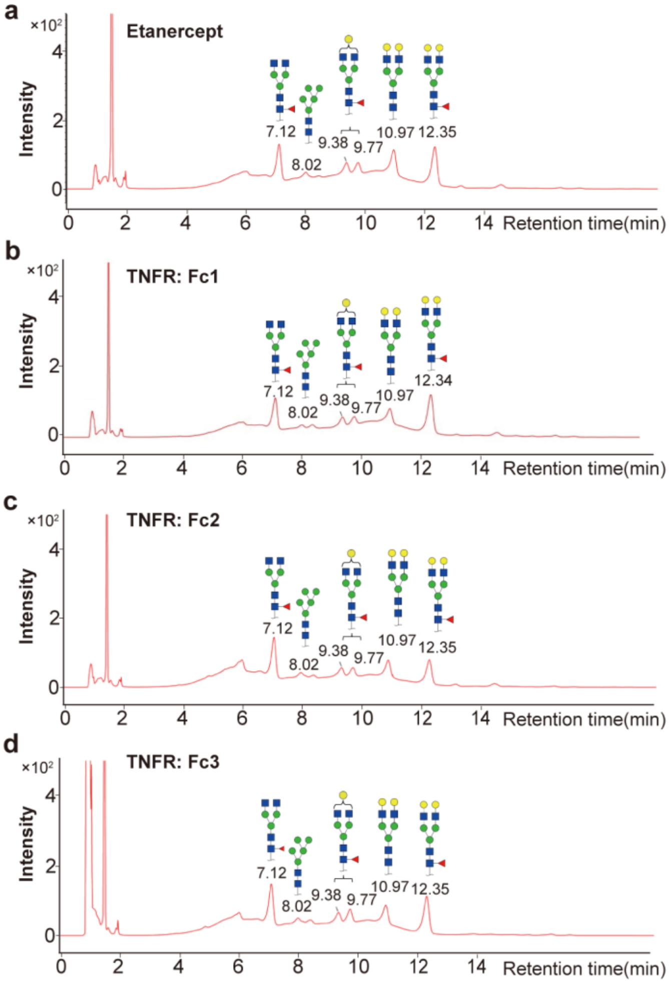

**Supplementary Fig. 24**. HILIC-UPLC profile of the total *N*-linked glycans released from TNFR: Fc-fusion pharmasutical proteins after desialylation. (a) Etanercept, (b) TNFR: Fc 1, (c) TNFR: Fc 2, (d) TNFR: Fc 3.

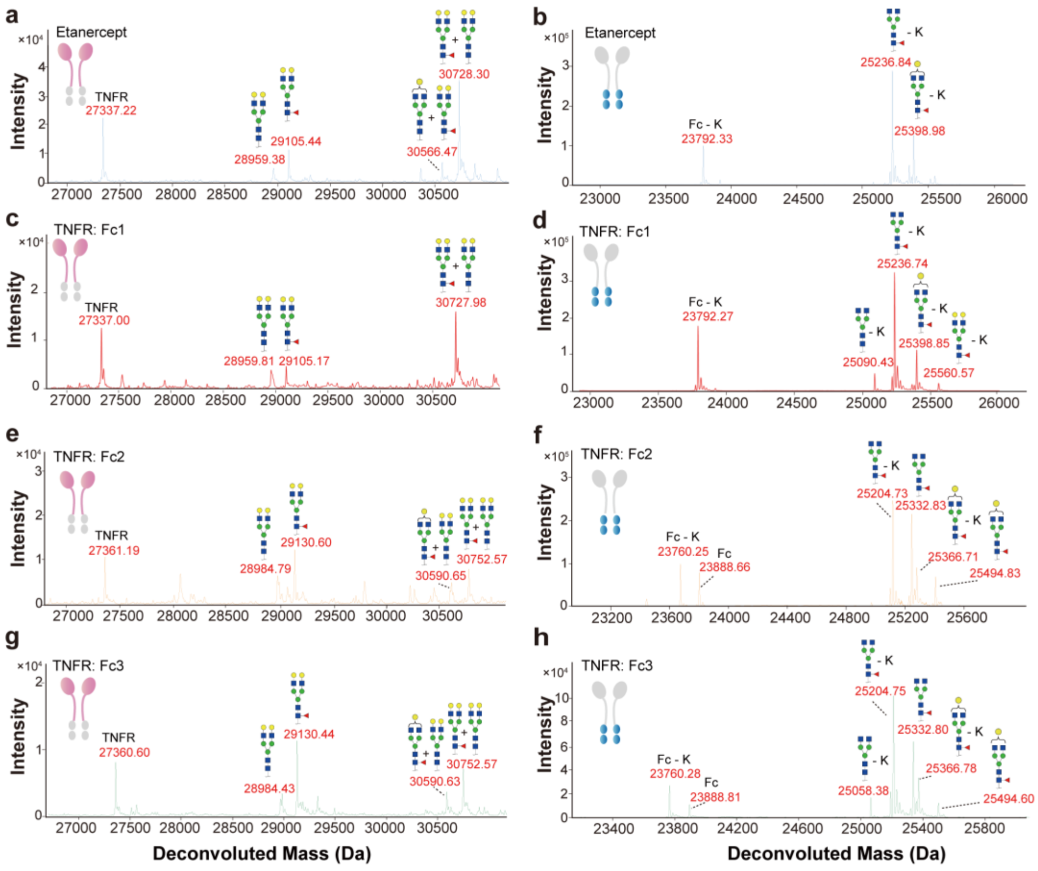

**Supplementary Fig. 25**. *N*-glycosylation TNFR: Fc-fusion pharmasutical proteins TNFR and Fc domains. Deconvoluted spectrum of Etanercept TNFR domain (a), Etanercept Fc domain (b), TNFR: Fc 1 TNFR domain (c), TNFR: Fc 1 Fc domain (d), TNFR: Fc 2 TNFR domain (e), TNFR: Fc 2 Fc domain (f), TNFR: Fc 3 TNFR domain (g) and TNFR: Fc 3 Fc domain (h). The glycoprotein protein was digested with sialidase and EngEF to remove sialic acid and *O*-glycan. The most probable *N*-glycoforms and *C*-terminal lysine loss variants are annotated.

**Supplementary Table 1**. Summary of TNFR: Fc-fusion biopharmaceuticals.

| Sample name | Product | Drug name | Manufacturers | Lot number | Notes |
|---|---|---|---|---|---|
| **Etanercept** | Enbrel® | Etanercept | Pfizer | GA5942 | original drug |
| **TNFR: Fc1** | YISAIPU® | Recombinant human type II tumor necrosis factor receptor-antibody fusion protein | Sunshine Guojian Pharmaceutical | 301-C2031 | biologics |
| **TNFR: Fc2** | Anbainuo® | Recombinant human type II tumor necrosis factor receptor-antibody fusion protein | BioRay Pharmaceutical | 500222005 | biologics |
| **TNFR: Fc3** | Qiangke® | Recombinant human TNF receptor-Ig fusion protein for injection | Celgen Biopharma | A201222U | biologics |

**Supplementary Table 2**. Assembly performance of Etanercept using *de novo* sequenced peptides.

| | Replication 1 | Replication 2 | Replication 3 |
|---|---|---|---|
| **Coverage** | 98.93% | 98.93% | 98.93% |
| **Accuracy** | 99.57% | 100.00% | 99.78% |
| **Insertion** | 0.00% | 1.71% | 1.50% |
| **Gap** | 0 | 0 | 0 |

**Supplementary Table 3**. Comparison summary of the theoretical molecular mass and experimental mass as determined by subunit mass analysis of deglycosylated TNFR and Fc domains.

| | **Domain** | **Molecular Weight (Da)** | **Amino acide variant** | **Theoretical Δ MW (Da)** | **Measured Δ MW (Da)** | **Δ mass (Da)** |
|---|---|---|---|---|---|---|
| **Etanercept** | TNFR | 27336.43 | - | - | - | - |
| | Fc | 23792.16 | - | - | - | - |
| **TNFR: Fc 1** | TNFR | 27336.57 | - | 0 | 0.14 | 0.14 |
| | Fc | 23792.16 | - | 0 | 0 | 0 |
| **TNFR:Fc 2** | TNFR | 27361.88 | M174R | 24.9996 | 25.45 | 0.4504 |
| | Fc | 23759.42 | E376D，M378L | -32.0651 | -32.74 | -0.6749 |
| **TNFR:Fc 3** | TNFR | 27361.94 | M174R | 24.9996 | 25.51 | 0.5104 |
| | Fc | 23759.41 | E376D，M378L | -32.0651 | -32.75 | -0.6849 |

**Supplementary Table 4**. *N*- glycopeptide identification by StrucGP.

| **Sample** | **Digestion** | **Modification** | **Site** | **Fractional abundance [%]： Rep2** | **Fractional abundance [%]： Rep2** | **Fractional abundance [%]： Rep3** |
|---|---|---|---|---|---|---|
| **Etanercept** | Chymotrypsin | A2G0, (Carbamidomethylation) | N149, (C157) | 11.37% | 8.87% | 13.25% |
| **Etanercept** | Chymotrypsin | A2G1, (Carbamidomethylation) | N149, (C157) | 12.51% | 9.64% | 10.30% |
| **Etanercept** | Chymotrypsin | A2G2, (Carbamidomethylation) | N149, (C157) | 45.05% | 59.35% | 46.87% |
| **Etanercept** | Chymotrypsin | A2G2F, (Carbamidomethylation) | N149, (C157) | 9.03% | 0.15% | 9.63% |
| **Etanercept** | Chymotrypsin | M5, (Carbamidomethylation) | N149, (C157) | 22.04% | 22.00% | 19.96% |
| **Etanercept** | Pepsin | A2G2F | N171 | 68.20% | 66.17% | 64.94% |
| **Etanercept** | Pepsin | A2G1F | N171 | 20.95% | 22.91% | 23.29% |
| **Etanercept** | Pepsin | M5 | N171 | 3.02% | 3.07% | 3.23% |
| **Etanercept** | Pepsin | A2G0F | N171 | 7.83% | 7.85% | 8.54% |
| **Etanercept** | Trypsin | A2G0 | N317 | 6.74% | 6.69% | 4.02% |
| **Etanercept** | Trypsin | A2G0F | N317 | 63.02% | 59.18% | 68.27% |
| **Etanercept** | Trypsin | A2G1 | N317 | 5.86% | 6.04% | 5.83% |
| **Etanercept** | Trypsin | A2G1F | N317 | 20.13% | 23.36% | 18.43% |
| **Etanercept** | Trypsin | A2G2F | N317 | 2.37% | 2.76% | 1.89% |
| **Etanercept** | Trypsin | M5 | N317 | 1.87% | 1.98% | 1.56% |
| **TNFR:FC1** | Chymotrypsin | A2G0F, (Carbamidomethylation) | N149, (C157) | 8.13% | 9.49% | 9.40% |
| **TNFR:FC1** | Chymotrypsin | A2G1, (Carbamidomethylation) | N149, (C157) | 23.91% | 19.25% | 20.07% |
| **TNFR:FC1** | Chymotrypsin | A2G1F, (Carbamidomethylation) | N149, (C157) | 4.19% | 4.40% | 3.75% |
| **TNFR:FC1** | Chymotrypsin | A2G2, (Carbamidomethylation) | N149, (C157) | 52.71% | 57.51% | 57.74% |
| **TNFR:FC1** | Chymotrypsin | A2G2F, (Carbamidomethylation) | N149, (C157) | 8.08% | 6.21% | 5.97% |
| **TNFR:FC1** | Chymotrypsin | M5, (Carbamidomethylation) | N149, (C157) | 2.98% | 3.15% | 3.07% |
| **TNFR:FC1** | Pepsin | A2G0F | N171 | 20.51% | 20.71% | 23.47% |
| **TNFR:FC1** | Pepsin | A2G1 | N171 | 2.50% | 2.93% | 3.13% |
| **TNFR:FC1** | Pepsin | A2G1F | N171 | 16.90% | 18.59% | 16.86% |
| **TNFR:FC1** | Pepsin | A2G2 | N171 | 4.40% | 3.04% | 2.40% |
| **TNFR:FC1** | Pepsin | A2G2F | N171 | 52.86% | 51.88% | 51.35% |
| **TNFR:FC1** | Pepsin | M5 | N171 | 2.83% | 2.86% | 2.79% |
| **TNFR:FC1** | Trypsin | A2G0 | N317 | 22.12% | 17.74% | 20.23% |
| **TNFR:FC1** | Trypsin | A2G0F | N317 | 41.30% | 53.97% | 56.98% |
| **TNFR:FC1** | Trypsin | A2G1 | N317 | 8.62% | 6.90% | 7.34% |
| **TNFR:FC1** | Trypsin | A2G1F | N317 | 22.13% | 16.47% | 11.41% |
| **TNFR:FC1** | Trypsin | A2G2F | N317 | 3.01% | 2.55% | 1.52% |
| **TNFR:FC1** | Trypsin | M5 | N317 | 2.83% | 2.37% | 2.52% |
| **TNFR:FC2** | Chymotrypsin | A2G1, (Carbamidomethylation) | N149, (C157) | 5.10% | 5.37% | 4.25% |

| Sample | Digestion | Modification | Site | Fractional abundance [%]: Rep2 | Fractional abundance [%]: Rep2 | Fractional abundance [%]: Rep3 |
|---|---|---|---|---|---|---|
| **TNFR:FC2** | Chymotrypsin | A2G1F, (Carbamidomethylation) | N149, (C157) | 5.14% | 4.54% | 4.13% |
| **TNFR:FC2** | Chymotrypsin | A2G2, (Carbamidomethylation) | N149, (C157) | 64.42% | 67.87% | 69.53% |
| **TNFR:FC2** | Chymotrypsin | M5, (Carbamidomethylation) | N149, (C157) | 25.34% | 22.22% | 22.09% |
| **TNFR:FC2** | Pepsin | A2G2F, (Carbamidomethylation) | N171, (C178) | 79.16% | 78.79% | 77.46% |
| **TNFR:FC2** | Pepsin | A2G1F, (Carbamidomethylation) | N171, (C178) | 20.84% | 21.21% | 22.54% |
| **TNFR:FC2** | Trypsin | A2G0 | N317 | 10.99% | 10.30% | 23.76% |
| **TNFR:FC2** | Trypsin | A2G0F | N317 | 70.03% | 69.54% | 50.59% |
| **TNFR:FC2** | Trypsin | A2G1F | N317 | 16.12% | 18.15% | 18.73% |
| **TNFR:FC2** | Trypsin | M3 | N317 | 2.08% | 2.01% | 5.02% |
| **TNFR:FC2** | Trypsin | M5 | N317 | 0.79% | 0.00% | 1.89% |
| **TNFR:FC3** | Chymotrypsin | A2G1, (Carbamidomethylation) | N149, (C157) | 5.39% | 5.79% | 4.55% |
| **TNFR:FC3** | Chymotrypsin | A2G1F, (Carbamidomethylation) | N149, (C157) | 8.03% | 12.84% | 11.63% |
| **TNFR:FC3** | Chymotrypsin | A2G2, (Carbamidomethylation) | N149, (C157) | 67.17% | 59.57% | 59.28% |
| **TNFR:FC3** | Chymotrypsin | A2G2F, (Carbamidomethylation) | N149, (C157) | 19.41% | 21.80% | 24.53% |
| **TNFR:FC3** | Pepsin | A2G0F, (Carbamidomethylation) | N171, (C178) | 24.84% | 20.67% | 21.45% |
| **TNFR:FC3** | Pepsin | A2G1F, (Carbamidomethylation) | N171, (C178) | 38.34% | 39.50% | 40.19% |
| **TNFR:FC3** | Pepsin | A2G2F, (Carbamidomethylation) | N171, (C178) | 36.82% | 39.83% | 38.37% |
| **TNFR:FC3** | Trypsin | A2G0 | N317 | 22.32% | 21.44% | 14.51% |
| **TNFR:FC3** | Trypsin | A2G0F | N317 | 63.31% | 63.43% | 72.48% |
| **TNFR:FC3** | Trypsin | A2G1 | N317 | 6.08% | 7.16% | 4.32% |
| **TNFR:FC3** | Trypsin | A2G1F | N317 | 7.69% | 6.92% | 7.49% |
| **TNFR:FC3** | Trypsin | A2G2F | N317 | 0.02% | 0.03% | 0.41% |
| **TNFR:FC3** | Trypsin | M5 | N317 | 0.58% | 1.02% | 0.79% |

**Supplementary Table 5**. *O*- glycopeptide identification by pGlyco3.1.

| Sample | Peptide | TotalScore | PepScore | GlyScore | ProSites |
|---|---|---|---|---|---|
| **Etanercept** | SMAPGAVHLPQPVSTR | 48.69302 | 69.60077 | 9.86435 | 199 |
| **Etanercept** | SMAPGAVHLPQPVSTR | 36.46027 | 50.76124 | 9.90131 | 199 |
| **Etanercept** | APGAVHLPQPVSTR | 38.40356 | 56.12473 | 5.49283 | 199 |
| **Etanercept** | APGAVHLPQPVSTR | 36.78503 | 53.60856 | 5.54135 | 199 |
| **Etanercept** | APGAVHLPQPVSTR | 43.71388 | 64.2148 | 5.64073 | 199 |
| **Etanercept** | SMAPGAVHLPQPVSTR | 52.81863 | 78.14735 | 5.77958 | 199 |
| **Etanercept** | PGAVHLPQPVSTR | 28.66586 | 38.32222 | 10.73261 | 199 |
| **Etanercept** | SMAPGAVHLPQPVSTR | 72.97516 | 108.8096 | 6.42553 | 199 |
| **Etanercept** | SMAPGAVHLPQPVSTR | 62.70363 | 93.12933 | 6.19874 | 199 |
| **Etanercept** | SMAPGAVHLPQPVSTR | 56.32771 | 83.4203 | 6.01289 | 199 |
| **Etanercept** | SMAPGAVHLPQPVSTR | 52.9847 | 78.32633 | 5.92167 | 199 |
| **Etanercept** | SMAPGAVHLPQPVSTR | 41.25636 | 60.42682 | 5.65409 | 199 |
| **Etanercept** | SMAPGAVHLPQPVSTR | 39.06281 | 54.51271 | 10.37013 | 199 |
| **Etanercept** | SMAPGAVHLPQPVSTR | 39.37343 | 54.97705 | 10.3953 | 199 |
| **Etanercept** | SMAPGAVHLPQPVSTR | 37.30314 | 52.01068 | 9.98913 | 199 |
| **Etanercept** | SMAPGAVHLPQPVSTR | 30.66547 | 47.17765 | 0 | 200 |
| **Etanercept** | SMAPGAVHLPQPVSTR | 42.41149 | 62.23747 | 5.59182 | 199 |
| **Etanercept** | SMAPGAVHLPQPVSTR | 43.89936 | 64.53654 | 5.57319 | 199 |
| **Etanercept** | SMAPGAVHLPQPVSTR | 43.78018 | 64.44484 | 5.40294 | 199 |
| **Etanercept** | APGAVHLPQPVSTR | 63.25064 | 91.13565 | 11.46418 | 199 |
| **Etanercept** | APGAVHLPQPVSTR | 58.00358 | 83.62055 | 10.4292 | 199 |
| **Etanercept** | SMAPGAVHLPQPVSTR | 7.62251 | 11.72694 | 0 | 199 |
| **Etanercept** | SMAPGAVHLPQPVSTR | 82.09141 | 122.5908 | 6.87835 | 199 |
| **Etanercept** | SMAPGAVHLPQPVSTR | 37.91932 | 52.16682 | 11.45968 | 199 |
| **Etanercept** | SMAPGAVHLPQPVSTR | 27.96535 | 36.98278 | 11.21871 | 199 |
| **Etanercept** | SMAPGAVHLPQPVSTR | 28.92575 | 38.38853 | 11.35201 | 199 |
| **Etanercept** | SMAPGAVHLPQPVSTR | 62.63771 | 93.20116 | 5.87702 | 199 |
| **Etanercept** | SMAPGAVHLPQPVSTR | 47.3158 | 69.96392 | 5.255 | 199 |
| **Etanercept** | SMAPGAVHLPQPVSTR | 14.61371 | 17.50422 | 9.24562 | 199 |
| **Etanercept** | SMAPGAVHLPQPVSTR | 38.77409 | 56.99196 | 4.94089 | 199 |
| **Etanercept** | SQHTQPTPEPSTAPSTSFLLPMGPSPPAEGSTGDEPK | 67.56348 | 101.2296 | 5.04068 | 216 |
| **Etanercept** | SQHTQPTPEPSTAPSTSFLLPMGPSPPAEGSTGDEPK | 79.20463 | 115.3236 | 12.12658 | 217 |
| **Etanercept** | SQHTQPTPEPSTAPSTSFLLPMGPSPPAEGSTGDEPK | 90.86454 | 133.2669 | 12.11733 | 217 |
| **Etanercept** | SQHTQPTPEPSTAPSTSFLLPMGPSPPAEGSTGDEPK | 75.28544 | 110.8768 | 9.18724 | 216 |
| **Etanercept** | SQHTQPTPEPSTAPSTSFLLPMGPSPPAEGSTGDEPK | 96.91629 | 145.8815 | 5.98095 | 213 |
| **Etanercept** | SQHTQPTPEPSTAPSTSFLLPMGPSPPAEGSTGDEPK | 70.47919 | 103.9817 | 8.26018 | 213 |
| **Etanercept** | SQHTQPTPEPSTAPSTSFLLPMGPSPPAEGSTGDEPK | 76.79597 | 111.7125 | 11.95104 | 205 |
| **Etanercept** | SQHTQPTPEPSTAPSTSFLLPMGPSPPAEGSTGDEPK | 55.31879 | 85.10583 | 0 | 205 |
| **Etanercept** | SQHTQPTPEPSTAPSTSFLLPMGPSPPAEGSTGDEPK | 88.65918 | 133.3206 | 5.71662 | 205 |
| **Etanercept** | SQHTQPTPEPSTAPSTSFLLPMGPSPPAEGSTGDEPK | 66.04549 | 98.49779 | 5.77694 | 205 |
| **Etanercept** | SQHTQPTPEPSTAPSTSFLLPMGPSPPAEGSTGDEPK | 85.27414 | 127.9485 | 6.02177 | 205 |

| Sample | Peptide | TotalScore | PepScore | GlyScore | ProSites |
|---|---|---|---|---|---|
| **Etanercept** | SQHTQPTPEPSTAPSTSFLLPMGPSPPAEGSTGDEPK | 81.89171 | 122.85 | 5.82642 | 205 |
| **Etanercept** | SQHTQPTPEPSTAPSTSFLLPMGPSPPAEGSTGDEPK | 65.23917 | 97.10889 | 6.05254 | 205 |
| **Etanercept** | SQHTQPTPEPSTAPSTSFLLPMGPSPPAEGSTGDEPK | 77.25456 | 115.5676 | 6.10186 | 213 |
| **Etanercept** | SQHTQPTPEPSTAPSTSFLLPMGPSPPAEGSTGDEPK | 75.06811 | 109.5159 | 11.09364 | 205 |
| **Etanercept** | SQHTQPTPEPSTAPSTSFLLPMGPSPPAEGSTGDEPK | 54.72168 | 84.1872 | 0 | 205 |
| **Etanercept** | LPAQVAFTPYAPEPGSTCR | 93.03266 | 139.9202 | 5.95579 | 8 |
| **Etanercept** | SQHTQPTPEPSTAPSTSFLLPMGPSPPAEGSTGDEPK | 55.8086 | 83.13453 | 5.06044 | 205 |
| **Etanercept** | SQHTQPTPEPSTAPSTSFLLPMGPSPPAEGSTGDEPK | 61.29918 | 91.25234 | 5.67188 | 205 |
| **Etanercept** | SQHTQPTPEPSTAPSTSFLLPMGPSPPAEGSTGDEPK | 67.84285 | 99.23463 | 9.54381 | 205 |
| **Etanercept** | SQHTQPTPEPSTAPSTSFLLPMGPSPPAEGSTGDEPK | 64.89619 | 96.70979 | 5.81378 | 205 |
| **Etanercept** | SQHTQPTPEPSTAPSTSFLLPMGPSPPAEGSTGDEPK | 58.76913 | 87.6429 | 5.1464 | 205 |
| **Etanercept** | SQHTQPTPEPSTAPSTSFLLPMGPSPPAEGSTGDEPK | 73.93381 | 108.7541 | 9.26752 | 205 |
| **Etanercept** | SQHTQPTPEPSTAPSTSFLLPMGPSPPAEGSTGDEPK | 72.75766 | 109.2305 | 5.0224 | 205 |
| **Etanercept** | SQHTQPTPEPSTAPSTSFLLPMGPSPPAEGSTGDEPK | 35.03742 | 51.38513 | 4.67739 | 213 |
| **Etanercept** | SQHTQPTPEPSTAPSTSFLLPMGPSPPAEGSTGDEPK | 82.88939 | 124.5101 | 5.59369 | 205 |
| **Etanercept** | SQHTQPTPEPSTAPSTSFLLPMGPSPPAEGSTGDEPK | 54.06005 | 80.59578 | 4.77941 | 213 |
| **Etanercept** | SQHTQPTPEPSTAPSTSFLLPMGPSPPAEGSTGDEP | 2.47303 | 3.80466 | 0 | 208 |
| **Etanercept** | SQHTQPTPEPSTAPSTSFLLPMGPSPPAEGSTGDEPK | 30.80187 | 47.38749 | 0 | 213 |
| **Etanercept** | SQHTQPTPEPSTAPSTSFLLPMGPSPPAEGSTGDEP | 4.06419 | 6.2526 | 0 | 226 |
| **Etanercept** | SQHTQPTPEPSTAPSTSFLLPMGPSPPAEGSTGDEPK | 74.27063 | 111.4248 | 5.27003 | 205 |
| **Etanercept** | SQHTQPTPEPSTAPSTSFLLPMGPSPPAEGSTGDEPK | 105.9894 | 156.9706 | 11.30997 | 205 |
| **Etanercept** | SCDKTHTCPPCPAPELLGGPSVF | 28.13768 | 42.57009 | 1.33464 | 245 |
| **Etanercept** | SCDKTHTCPPCPAPELLGGPSVF | 14.96575 | 22.32951 | 1.2902 | 245 |
| **Etanercept** | SQHTQPTPEPSTAPSTSFLLPMGPSPPAEGSTGDEPK | 98.23392 | 144.0255 | 13.19236 | 218 |
| **Etanercept** | SQHTQPTPEPSTAPSTSFLLPMGPSPPAEGSTGDEPK | 63.22303 | 94.56535 | 5.01586 | 213 |
| **Etanercept** | SQHTQPTPEPSTAPSTSFLLPMGPSPPAEGSTGDEPK | 66.15872 | 99.24753 | 4.70808 | 216 |
| **Etanercept** | SQHTQPTPEPSTAPSTSFLLPMGPSPPAEGSTGDEPK | 31.88316 | 49.05102 | 0 | 217 |
| **Etanercept** | SQHTQPTPEPSTAPSTSFLLPMGPSPPAEGSTGDEPK | 59.25432 | 89.1714 | 3.69403 | 205 |
| **Etanercept** | SQHTQPTPEPSTAPSTSFLLPMGPSPPAEGSTGDEPK | 90.85748 | 136.1035 | 6.82912 | 212 |
| **Etanercept** | SQHTQPTPEPSTAPSTSFLLPMGPSPPAEGSTGDEPK | 43.97594 | 67.65529 | 0 | 205 |
| **Etanercept** | SQHTQPTPEPSTAPSTSFLLPMGPSPPAEGSTGDEPK | 85.51478 | 128.4944 | 5.69541 | 213 |
| **Etanercept** | SQHTQPTPEPSTAPSTSFLLPMGPSPPAEGSTGDEPK | 97.62218 | 146.8811 | 6.14138 | 213 |
| **Etanercept** | HTQPTPEPSTAPSTSFLLPMGPSPPAEGSTGDEPK | 10.75617 | 15.86332 | 1.27145 | 216 |
| **Etanercept** | SQHTQPTPEPSTAPSTSFLLPMGPSPPAEGSTGDEPK | 115.8654 | 174.3129 | 7.32014 | 205 |
| **Etanercept** | SQHTQPTPEPSTAPSTSFLLPMGPSPPAEGSTGDEPK | 91.43102 | 136.8437 | 7.0933 | 205 |
| **Etanercept** | SQHTQPTPEPSTAPSTSFLLPMGPSPPAEGSTGDEPK | 49.05969 | 71.95621 | 6.53758 | 205 |
| **Etanercept** | SQHTQPTPEPSTAPSTSFLLPMGPSPPAEGSTGDEPK | 88.56004 | 133.0726 | 5.89387 | 205 |
| **Etanercept** | SQHTQPTPEPSTAPSTSFLLPMGPSPPAEGSTGDEPK | 79.90455 | 119.7831 | 5.84433 | 205 |
| **Etanercept** | SQHTQPTPEPSTAPSTSFLLPMGPSPPAEGSTGDEPK | 82.73229 | 124.156 | 5.80252 | 205 |
| **Etanercept** | SQHTQPTPEPSTAPSTSFLLPMGPSPPAEGSTGDEPK | 72.10295 | 107.8677 | 5.68269 | 205 |
| **Etanercept** | HTQPTPEPSTAPSTSFLLPMGPSPPAEGSTGDEPK | 42.74564 | 65.76252 | 0 | 205 |
| **Etanercept** | SQHTQPTPEPSTAPSTSFLLPMGPSPPAEGSTGDEPK | 69.61479 | 104.4377 | 4.94362 | 213 |

| Sample | Peptide | TotalScore | PepScore | GlyScore | ProSites |
|---|---|---|---|---|---|
| **Etanercept** | HTQPTPEPSTAPSTSFLLPMGPSPPAEGSTGDEPK | 42.14134 | 62.30906 | 4.68702 | 205 |
| **Etanercept** | SQHTQPTPEPSTAPSTSFLLPMGPSPPAEGSTGDEPK | 86.76252 | 130.7896 | 4.99797 | 205 |
| **Etanercept** | SRWQQGNVFSCSVMHEALHNHYTQKSLSLS | 3.78691 | 5.82602 | 0 | 457 |
| **Etanercept** | SQHTQPTPEPSTAPSTSFLLPMGPSPPAEGSTGDEPK | 46.31379 | 71.25198 | 0 | 205 |
| **Etanercept** | SQHTQPTPEPSTAPSTSFLLPMGPSPPAEGSTGDEPK | 51.89989 | 79.84598 | 0 | 205 |
| **Etanercept** | THTCPPCPAPELLGGPSVFLFPPKPK | 53.52667 | 72.73973 | 17.84528 | 245 |
| **Etanercept** | THTCPPCPAPELLGGPSVFLFPPKPK | 51.71703 | 69.71676 | 18.28897 | 245 |
| **Etanercept** | THTCPPCPAPELLGGPSVFLFPPKPK | 15.94191 | 21.46761 | 5.6799 | 245 |
| **Etanercept** | THTCPPCPAPELLGGPSVFLFPPKPK | 52.34508 | 70.01439 | 19.53065 | 245 |
| **Etanercept** | THTCPPCPAPELLGGPSVFLFPPKPK | 67.56348 | 102.4049 | 2.85801 | 245 |
| **Etanercept** | THTCPPCPAPELLGGPSVFLFPPKPK | 47.35797 | 71.34365 | 2.81314 | 245 |
| **Etanercept** | REQNRICTCRPGWYCALSK | 2.6499 | 4.07676 | 0 | 97 |
| **Etanercept** | THTCPPCPAPELLGGPSVFLFPPK | 37.103 | 56.36416 | 1.33228 | 245 |
| **Etanercept** | DTVCDSCEDSTYTQLWNWVPECLSCGSR | 5.58331 | 8.58971 | 0 | 62 |
| **Etanercept** | SMAPGAVHLPQPVSTR | 18.21446 | 28.02224 | 0 | 199 |
| **Etanercept** | SMAPGAVHLPQPVSTR | 16.2275 | 23.20679 | 3.26596 | 199 |
| **Etanercept** | SQHTQPTPEPSTAPSTSFLLPMGPSPPAEGSTGDEPK | 3.77152 | 3.98294 | 3.37889 | 202 |
| **Etanercept** | SMAPGAVHLPQPVSTR | 7.6633 | 10.15433 | 3.03709 | 199 |
| **Etanercept** | SMAPGAVHLPQPVSTR | 42.02317 | 59.23765 | 10.05343 | 199 |
| **Etanercept** | APGAVHLPQPVSTR | 39.97135 | 58.75239 | 5.09227 | 199 |
| **Etanercept** | SMAPGAVHLPQPVSTR | 40.2582 | 56.5568 | 9.98938 | 199 |
| **Etanercept** | APGAVHLPQPVSTR | 44.01287 | 64.73853 | 5.52236 | 199 |
| **Etanercept** | APGAVHLPQPVSTR | 42.77384 | 62.80563 | 5.57195 | 199 |
| **Etanercept** | APGAVHLPQPVSTR | 45.08967 | 66.58575 | 5.16839 | 199 |
| **Etanercept** | SMAPGAVHLPQPVSTR | 54.00604 | 80.088 | 5.56813 | 199 |
| **Etanercept** | SMAPGAVHLPQPVSTR | 47.96867 | 70.67253 | 5.80437 | 199 |
| **Etanercept** | SMAPGAVHLPQPVSTR | 98.56455 | 148.0383 | 6.68473 | 199 |
| **Etanercept** | SMAPGAVHLPQPVSTR | 83.29558 | 124.6811 | 6.43667 | 199 |
| **Etanercept** | SMAPGAVHLPQPVSTR | 45.10617 | 66.15074 | 6.02339 | 199 |
| **Etanercept** | SMAPGAVHLPQPVSTR | 46.60688 | 68.48804 | 5.97043 | 199 |
| **Etanercept** | SMAPGAVHLPQPVSTR | 42.74733 | 60.02959 | 10.6517 | 199 |
| **Etanercept** | SMAPGAVHLPQPVSTR | 42.24815 | 59.14514 | 10.86802 | 199 |
| **Etanercept** | MAPGAVHLPQPVSTR | 3.5132 | 5.40492 | 0 | 200 |
| **Etanercept** | SMAPGAVHLPQPVSTR | 49.51008 | 73.27893 | 5.36792 | 199 |
| **Etanercept** | SMAPGAVHLPQPVSTR | 31.23611 | 45.36021 | 5.00563 | 200 |
| **Etanercept** | SMAPGAVHLPQPVSTR | 48.41498 | 71.45198 | 5.63198 | 199 |
| **Etanercept** | SMAPGAVHLPQPVSTR | 38.40821 | 56.20901 | 5.34958 | 199 |
| **Etanercept** | SMAPGAVHLPQPVSTR | 54.05059 | 80.00011 | 5.85861 | 199 |
| **Etanercept** | SMAPGAVHLPQPVSTR | 47.43015 | 69.94924 | 5.60897 | 199 |
| **Etanercept** | APGAVHLPQPVSTR | 58.01744 | 83.1666 | 11.31186 | 199 |
| **Etanercept** | SMAPGAVHLPQPVSTR | 77.97716 | 116.2964 | 6.8129 | 199 |
| **Etanercept** | SMAPGAVHLPQPVSTR | 32.86925 | 44.39391 | 11.46631 | 199 |

| Sample | Peptide | TotalScore | PepScore | GlyScore | ProSites |
|---|---|---|---|---|---|
| **Etanercept** | SMAPGAVHLPQPVSTR | 37.38892 | 51.35557 | 11.45085 | 199 |
| **Etanercept** | SMAPGAVHLPQPVSTR | 60.451 | 89.77049 | 6.00053 | 199 |
| **Etanercept** | QQGNVFSCSVMHEALHNHYTQK | 29.73945 | 43.61982 | 3.96161 | 444 |
| **Etanercept** | SMAPGAVHLPQPVSTR | 52.94282 | 78.40764 | 5.65103 | 199 |
| **Etanercept** | SMAPGAVHLPQPVSTR | 61.94776 | 92.27344 | 5.62864 | 199 |
| **Etanercept** | SMAPGAVHLPQPVSTR | 24.27923 | 37.35267 | 0 | 199 |
| **Etanercept** | SMAPGAVHLPQPVSTR | 46.55353 | 68.86578 | 5.11651 | 199 |
| **Etanercept** | SMAPGAVHLPQPVSTR | 20.18598 | 28.69757 | 4.37874 | 199 |
| **Etanercept** | SMAPGAVHLPQPVSTR | 24.85304 | 38.23545 | 0 | 199 |
| **Etanercept** | SQHTQPTPEPSTAPSTSFLLPMGPSPPAEGSTGDEPK | 32.64172 | 48.11774 | 3.90054 | 205 |
| **Etanercept** | SQHTQPTPEPSTAPSTSFLLPMGPSPPAEGSTGDEPK | 68.72414 | 102.8514 | 5.34495 | 217 |
| **Etanercept** | SQHTQPTPEPSTAPSTSFLLPMGPSPPAEGSTGDEPK | 82.80885 | 120.4414 | 12.91988 | 217 |
| **Etanercept** | SQHTQPTPEPSTAPSTSFLLPMGPSPPAEGSTGDEPK | 70.13363 | 102.5015 | 10.02189 | 217 |
| **Etanercept** | QPTPEPSTAPSTSFLLPMGPSPPAEGSTGDEPK | 10.51986 | 16.1844 | 0 | 208 |
| **Etanercept** | SQHTQPTPEPSTAPSTSFLLPMGPSPPAEGSTGDEPK | 82.17826 | 123.0302 | 6.31034 | 213 |
| **Etanercept** | SQHTQPTPEPSTAPSTSFLLPMGPSPPAEGSTGDEPK | 53.96662 | 80.12867 | 5.37996 | 213 |
| **Etanercept** | PQVYTLPPSREEMTK | 3.58409 | 4.99741 | 0.95936 | 370 |
| **Etanercept** | SQHTQPTPEPSTAPSTSFLLPMGPSPPAEGSTGDEPK | 62.91412 | 96.79095 | 0 | 205 |
| **Etanercept** | SQHTQPTPEPSTAPSTSFLLPMGPSPPAEGSTGDEPK | 107.004 | 161.1239 | 6.49539 | 205 |
| **Etanercept** | SQHTQPTPEPSTAPSTSFLLPMGPSPPAEGSTGDEPK | 87.77541 | 131.9315 | 5.77132 | 205 |
| **Etanercept** | SQHTQPTPEPSTAPSTSFLLPMGPSPPAEGSTGDEPK | 70.95659 | 105.9313 | 6.00364 | 205 |
| **Etanercept** | SQHTQPTPEPSTAPSTSFLLPMGPSPPAEGSTGDEPK | 75.47073 | 112.9788 | 5.81284 | 205 |
| **Etanercept** | SQHTQPTPEPSTAPSTSFLLPMGPSPPAEGSTGDEPK | 79.51651 | 119.2253 | 5.77167 | 205 |
| **Etanercept** | SQHTQPTPEPSTAPSTSFLLPMGPSPPAEGSTGDEPK | 73.97025 | 110.514 | 6.10324 | 205 |
| **Etanercept** | SQHTQPTPEPSTAPSTSFLLPMGPSPPAEGSTGDEPK | 59.04454 | 87.80792 | 5.62682 | 205 |
| **Etanercept** | SQHTQPTPEPSTAPSTSFLLPMGPSPPAEGSTGDEPK | 85.68365 | 128.522 | 6.12681 | 213 |
| **Etanercept** | LPAQVAFTPYAPEPGSTCR | 67.43233 | 100.6064 | 5.82333 | 8 |
| **Etanercept** | LPAQVAFTPYAPEPGSTCR | 91.03752 | 136.8034 | 6.04373 | 8 |
| **Etanercept** | SQHTQPTPEPSTAPSTSFLLPMGPSPPAEGSTGDEPK | 66.15741 | 97.28703 | 8.34527 | 205 |
| **Etanercept** | SQHTQPTPEPSTAPSTSFLLPMGPSPPAEGSTGDEPK | 78.94255 | 115.4369 | 11.16738 | 213 |
| **Etanercept** | SQHTQPTPEPSTAPSTSFLLPMGPSPPAEGSTGDEPK | 49.30154 | 73.39967 | 4.54788 | 213 |
| **Etanercept** | SQHTQPTPEPSTAPSTSFLLPMGPSPPAEGSTGDEPK | 75.54918 | 113.098 | 5.81565 | 213 |
| **Etanercept** | SQHTQPTPEPSTAPSTSFLLPMGPSPPAEGSTGDEPK | 67.24618 | 100.7001 | 5.11748 | 217 |
| **Etanercept** | SQHTQPTPEPSTAPSTSFLLPMGPSPPAEGSTGDEPK | 65.26796 | 97.74343 | 4.95638 | 205 |
| **Etanercept** | SQHTQPTPEPSTAPSTSFLLPMGPSPPAEGSTGDEPK | 63.47755 | 95.06042 | 4.82364 | 205 |
| **Etanercept** | SQHTQPTPEPSTAPSTSFLLPMGPSPPAEGSTGDEPK | 69.87736 | 104.3508 | 5.85524 | 213 |
| **Etanercept** | SQHTQPTPEPSTAPSTSFLLPMGPSPPAEGSTGDEPK | 36.24048 | 53.28485 | 4.58666 | 212 |
| **Etanercept** | SQHTQPTPEPSTAPSTSFLLPMGPSPPAEGSTGDEPK | 70.83339 | 106.3525 | 4.86938 | 205 |
| **Etanercept** | SQHTQPTPEPSTAPSTSFLLPMGPSPPAEGSTGDEPK | 20.23751 | 28.65359 | 4.60765 | 213 |
| **Etanercept** | SCDKTHTCPPCPAPELLGGPSVF | 33.60453 | 46.1814 | 10.24749 | 245 |
| **Etanercept** | SCDKTHTCPPCPAPELLGGPSVF | 17.76012 | 26.62347 | 1.29962 | 245 |
| **Etanercept** | SQHTQPTPEPSTAPSTSFLLPMGPSPPAEGSTGDEPK | 70.41874 | 103.5304 | 8.92564 | 205 |

| Sample | Peptide | TotalScore | PepScore | GlyScore | ProSites |
|---|---|---|---|---|---|
| **Etanercept** | SQHTQPTPEPSTAPSTSFLLPMGPSPPAEGSTGDEPK | 89.18807 | 130.2741 | 12.88553 | 218 |
| **Etanercept** | SQHTQPTPEPSTAPSTSFLLPMGPSPPAEGSTGDEPK | 72.58233 | 106.7032 | 9.2151 | 205 |
| **Etanercept** | SQHTQPTPEPSTAPSTSFLLPMGPSPPAEGSTGDEPK | 68.03512 | 101.573 | 5.75054 | 205 |
| **Etanercept** | SQHTQPTPEPSTAPSTSFLLPMGPSPPAEGSTGDEPK | 66.0382 | 98.5249 | 5.70575 | 213 |
| **Etanercept** | SQHTQPTPEPSTAPSTSFLLPMGPSPPAEGSTGDEPK | 72.2467 | 108.0295 | 5.79292 | 218 |
| **Etanercept** | SQHTQPTPEPSTAPSTSFLLPMGPSPPAEGSTGDEPK | 69.50191 | 103.8873 | 5.64325 | 218 |
| **Etanercept** | SQHTQPTPEPSTAPSTSFLLPMGPSPPAEGSTGDEPK | 62.74894 | 93.79472 | 5.09248 | 205 |
| **Etanercept** | SQHTQPTPEPSTAPSTSFLLPMGPSPPAEGSTGDEPK | 120.0795 | 181.0622 | 6.826 | 217 |
| **Etanercept** | TVDKSRWQQGNVFSCSVMHEALHNHYTQK | 3.00236 | 4.21026 | 0.75913 | 457 |
| **Etanercept** | SQHTQPTPEPSTAPSTSFLLPMGPSPPAEGSTGDEPK | 80.2641 | 120.0797 | 6.32084 | 212 |
| **Etanercept** | SQHTQPTPEPSTAPSTSFLLPMGPSPPAEGSTGDEPK | 82.5169 | 123.4403 | 6.51631 | 212 |
| **Etanercept** | SQHTQPTPEPSTAPSTSFLLPMGPSPPAEGSTGDEPK | 5.83915 | 7.53409 | 2.69141 | 226 |
| **Etanercept** | SQHTQPTPEPSTAPSTSFLLPMGPSPPAEGSTGDEPK | 68.18065 | 100.3829 | 8.37642 | 205 |
| **Etanercept** | SQHTQPTPEPSTAPSTS | 7.01235 | 10.78823 | 0 | 216 |
| **Etanercept** | HTQPTPEPSTAPSTSFLLPMGPSPPAEGSTGDEPK | 33.01226 | 49.41651 | 2.54722 | 226 |
| **Etanercept** | SQHTQPTPEPSTAPSTSFLLPMGPSPPAEGSTGDEPK | 61.22128 | 89.48258 | 8.73601 | 205 |
| **Etanercept** | HTQPTPEPSTAPSTSFLLPMGPSPPAEGSTGDEPK | 33.64556 | 51.00692 | 1.40303 | 226 |
| **Etanercept** | SQHTQPTPEPSTAPSTSFLLPMGPSPPAEGSTGDEPK | 102.791 | 154.3103 | 7.11231 | 205 |
| **Etanercept** | SQHTQPTPEPSTAPSTSFLLPMGPSPPAEGSTGDEPK | 92.87336 | 139.0218 | 7.16913 | 205 |
| **Etanercept** | SQHTQPTPEPSTAPSTSFLLPMGPSPPAEGSTGDEPK | 48.96605 | 72.58386 | 5.10442 | 205 |
| **Etanercept** | SQHTQPTPEPSTAPSTSFLLPMGPSPPAEGSTGDEPK | 122.0209 | 183.7718 | 7.34052 | 205 |
| **Etanercept** | SQHTQPTPEPSTAPSTSFLLPMGPSPPAEGSTGDEPK | 96.45338 | 148.3898 | 0 | 205 |
| **Etanercept** | HTQPTPEPSTAPSTSFLLPMGPSPPAEGSTGDEPK | 10.16775 | 15.64269 | 0 | 212 |
| **Etanercept** | SQHTQPTPEPSTAPSTSFLLPMGPSPPAEGSTGDEPK | 75.57658 | 110.3136 | 11.06501 | 205 |
| **Etanercept** | SQHTQPTPEPSTAPSTSFLLPMGPSPPAEGSTGDEPK | 71.95874 | 107.6695 | 5.63872 | 205 |
| **Etanercept** | EEMTKNQVSLTCLVKGFYPSDIAVEWESNGQ | 2.3859 | 3.67062 | 0 | 379 |
| **Etanercept** | SQHTQPTPEPSTAPSTSFLLPMGPSPPAEGSTGDEPK | 72.7272 | 108.8225 | 5.693 | 208 |
| **Etanercept** | SQHTQPTPEPSTAPSTSFLLPMGPSPPAEGSTGDEPK | 50.55418 | 77.77566 | 0 | 205 |
| **Etanercept** | SQHTQPTPEPSTAPSTSFLLPMGPSPPAEGSTGDEPK | 85.2272 | 128.3991 | 5.05086 | 205 |
| **Etanercept** | SQHTQPTPEPSTAPSTSFLLPMGPSPPAEGSTGDEPK | 77.20261 | 112.8496 | 11.00113 | 205 |
| **Etanercept** | SQHTQPTPEPSTAPSTSFLLPMGPSPPAEGSTGDEPK | 77.72941 | 116.5746 | 5.5883 | 217 |
| **Etanercept** | SQHTQPTPEPSTAPSTSFLLPMGPSPPAEGSTGDEPK | 75.31375 | 113.2632 | 4.83618 | 208 |
| **Etanercept** | HTQPTPEPSTAPSTSFLLPMGPSPPAEGSTGDEPK | 30.97653 | 47.65619 | 0 | 205 |
| **Etanercept** | SQHTQPTPEPSTAPSTSFLLPMGPSPPAEGSTGDEPK | 75.15795 | 112.9524 | 4.96834 | 205 |
| **Etanercept** | HTQPTPEPSTAPSTSFLLPMGPSPPAEGSTGDEPK | 10.96503 | 16.21806 | 1.20941 | 205 |
| **Etanercept** | HTQPTPEPSTAPSTSFLLPMGPSPPAEGSTGDEPK | 26.95028 | 41.46196 | 0 | 205 |
| **Etanercept** | SQHTQPTPEPSTAPSTSFLLPMGPSPPAEGSTGDEPK | 57.00444 | 85.07677 | 4.8701 | 205 |
| **Etanercept** | QHTQPTPEPSTAPSTSFLLPMGPSPPAEGSTGDEPK | 9.44005 | 14.52316 | 0 | 217 |
| **Etanercept** | THTCPPCPAPELLGGPSVFLFPPKPK | 44.24812 | 58.59736 | 17.59954 | 245 |
| **Etanercept** | THTCPPCPAPELLGGPSVFLFPPKPK | 49.45815 | 66.75034 | 17.34407 | 245 |
| **Etanercept** | THTCPPCPAPELLGGPSVFLFPPKPK | 28.23809 | 40.32015 | 5.79997 | 245 |
| **Etanercept** | THTCPPCPAPELLGGPSVFLFPPKPK | 31.39221 | 39.04943 | 17.17166 | 245 |

| Sample | Peptide | TotalScore | PepScore | GlyScore | ProSites |
|---|---|---|---|---|---|
| **Etanercept** | THTCPPCPAPELLGGPSVFLFPPK | 55.44385 | 80.01947 | 9.80341 | 245 |
| **Etanercept** | SMAPGAVHLPQPVSTR | 13.18594 | 18.33531 | 3.62281 | 199 |
| **Etanercept** | SQHTQPTPEPSTAPSTSFLLPMGPSPPAEGSTGDEPK | 3.43242 | 5.28065 | 0 | 205 |
| **Etanercept** | SQHTQPTPEPSTAPSTSFLLPMGPSPPAEGSTGDEPK | 2.40315 | 3.69715 | 0 | 202 |
| **Etanercept** | SMAPGAVHLPQPVSTR | 41.87201 | 58.99017 | 10.08113 | 199 |
| **Etanercept** | SMAPGAVHLPQPVSTR | 43.77818 | 61.8893 | 10.14324 | 199 |
| **Etanercept** | SMAPGAVHLPQPVSTR | 38.10781 | 53.05535 | 10.3481 | 199 |
| **Etanercept** | APGAVHLPQPVSTR | 44.98824 | 66.34115 | 5.33284 | 199 |
| **Etanercept** | APGAVHLPQPVSTR | 46.39193 | 68.41443 | 5.493 | 199 |
| **Etanercept** | APGAVHLPQPVSTR | 42.58894 | 62.59915 | 5.42712 | 199 |
| **Etanercept** | APGAVHLPQPVSTR | 49.84618 | 73.93055 | 5.11808 | 199 |
| **Etanercept** | SMAPGAVHLPQPVSTR | 46.00703 | 67.64877 | 5.81524 | 199 |
| **Etanercept** | SMAPGAVHLPQPVSTR | 58.83536 | 87.27026 | 6.02769 | 199 |
| **Etanercept** | SMAPGAVHLPQPVSTR | 73.2152 | 109.2526 | 6.28856 | 199 |
| **Etanercept** | SMAPGAVHLPQPVSTR | 60.7945 | 87.21853 | 11.7213 | 200 |
| **Etanercept** | SMAPGAVHLPQPVSTR | 51.98191 | 76.72166 | 6.03667 | 199 |
| **Etanercept** | SMAPGAVHLPQPVSTR | 4.71929 | 7.26044 | 0 | 199 |
| **Etanercept** | SMAPGAVHLPQPVSTR | 37.10117 | 57.07872 | 0 | 199 |
| **Etanercept** | SMAPGAVHLPQPVSTR | 41.20242 | 60.39855 | 5.55247 | 199 |
| **Etanercept** | SMAPGAVHLPQPVSTR | 48.76475 | 71.97903 | 5.65252 | 199 |
| **Etanercept** | SMAPGAVHLPQPVSTR | 39.19978 | 54.99906 | 9.85827 | 199 |
| **Etanercept** | SMAPGAVHLPQPVSTR | 40.92281 | 57.088 | 10.90173 | 199 |
| **Etanercept** | SMAPGAVHLPQPVSTR | 44.6114 | 65.63224 | 5.57269 | 199 |
| **Etanercept** | SMAPGAVHLPQPVSTR | 28.39612 | 40.88203 | 5.208 | 200 |
| **Etanercept** | SMAPGAVHLPQPVSTR | 52.25921 | 77.39351 | 5.58123 | 199 |
| **Etanercept** | SMAPGAVHLPQPVSTR | 44.45569 | 65.73529 | 4.93645 | 199 |
| **Etanercept** | SMAPGAVHLPQPVSTR | 53.44586 | 79.31844 | 5.39678 | 199 |
| **Etanercept** | SMAPGAVHLPQPVSTR | 57.29775 | 85.05857 | 5.74194 | 199 |
| **Etanercept** | APGAVHLPQPVSTR | 65.45757 | 97.43101 | 6.07834 | 199 |
| **Etanercept** | APGAVHLPQPVSTR | 61.89677 | 89.05823 | 11.45407 | 199 |
| **Etanercept** | SMAPGAVHLPQPVSTR | 17.06403 | 26.25235 | 0 | 199 |
| **Etanercept** | APGAVHLPQPVSTR | 66.83031 | 96.8027 | 11.16729 | 199 |
| **Etanercept** | SMAPGAVHLPQPVSTR | 82.71854 | 123.4394 | 7.09404 | 199 |
| **Etanercept** | SMAPGAVHLPQPVSTR | 29.576 | 39.43237 | 11.27131 | 199 |
| **Etanercept** | SMAPGAVHLPQPVSTR | 64.40627 | 95.85446 | 6.0025 | 199 |
| **Etanercept** | SMAPGAVHLPQPVSTR | 32.31791 | 43.69064 | 11.19712 | 199 |
| **Etanercept** | SMAPGAVHLPQPVSTR | 56.0741 | 83.27374 | 5.56048 | 199 |
| **Etanercept** | SMAPGAVHLPQPVSTR | 53.65624 | 79.75321 | 5.19045 | 199 |
| **Etanercept** | SMAPGAVHLPQPVSTR | 37.47424 | 54.99776 | 4.93054 | 199 |
| **Etanercept** | REPQVYTLPPSR | 3.28878 | 4.6327 | 0.79293 | 374 |
| **Etanercept** | SMAPGAVHLPQPVSTR | 38.35119 | 56.37345 | 4.88128 | 199 |
| **Etanercept** | SQHTQPTPEPSTAPSTSFLLPMGPSPPAEGSTGDEPK | 17.05882 | 23.74137 | 4.64835 | 205 |

| Sample | Peptide | TotalScore | PepScore | GlyScore | ProSites |
|---|---|---|---|---|---|
| **Etanercept** | SQHTQPTPEPSTAPSTSFLLPMGPSPPAEGSTGDEPK | 86.3562 | 125.9333 | 12.85582 | 217 |
| **Etanercept** | SQHTQPTPEPSTAPSTSFLLPMGPSPPAEGSTGDEPK | 65.26661 | 97.52811 | 5.35238 | 217 |
| **Etanercept** | QPTPEPSTAPSTSFLLPMGPSPPAEGSTGDEPK | 15.32824 | 23.5819 | 0 | 212 |
| **Etanercept** | LPMGPSPPAEGSTGDEPK | 6.35925 | 9.35436 | 0.79689 | 226 |
| **Etanercept** | SQHTQPTPEPSTAPSTSFLLPMGPSPPAEGSTGDEPK | 19.68029 | 24.77738 | 10.21427 | 208 |
| **Etanercept** | SQHTQPTPEPSTAPSTSFLLPMGPSPPAEGSTGDEPK | 92.42299 | 138.7833 | 6.32523 | 217 |
| **Etanercept** | SQHTQPTPEPSTAPSTSFLLPMGPSPPAEGSTGDEPK | 52.51543 | 77.85775 | 5.45113 | 213 |
| **Etanercept** | QPTPEPSTAPSTSFLLPMGPSPPAEGSTGDEPK | 9.06064 | 13.93945 | 0 | 208 |
| **Etanercept** | SQHTQPTPEPSTAPSTSFLLPMGPSPPAEGSTGDEPK | 72.32004 | 107.7978 | 6.4327 | 205 |
| **Etanercept** | SQHTQPTPEPSTAPSTSFLLPMGPSPPAEGSTGDEPK | 75.62965 | 112.8453 | 6.51481 | 205 |
| **Etanercept** | SQHTQPTPEPSTAPSTSFLLPMGPSPPAEGSTGDEPK | 71.16426 | 109.4835 | 0 | 205 |
| **Etanercept** | SQHTQPTPEPSTAPSTSFLLPMGPSPPAEGSTGDEPK | 84.86444 | 123.9392 | 12.29697 | 205 |
| **Etanercept** | PQVYTLPPSREEMTK | 2.22522 | 3.42341 | 0 | 370 |
| **Etanercept** | SQHTQPTPEPSTAPSTSFLLPMGPSPPAEGSTGDEPK | 86.80999 | 127.9543 | 10.39909 | 205 |
| **Etanercept** | SQHTQPTPEPSTAPSTSFLLPMGPSPPAEGSTGDEPK | 80.73392 | 120.8786 | 6.17953 | 205 |
| **Etanercept** | SQHTQPTPEPSTAPSTSFLLPMGPSPPAEGSTGDEPK | 79.39003 | 119.0073 | 5.81507 | 205 |
| **Etanercept** | SQHTQPTPEPSTAPSTSFLLPMGPSPPAEGSTGDEPK | 77.29543 | 115.6503 | 6.0649 | 205 |
| **Etanercept** | SQHTQPTPEPSTAPSTSFLLPMGPSPPAEGSTGDEPK | 74.51258 | 111.587 | 5.6602 | 205 |
| **Etanercept** | SQHTQPTPEPSTAPSTSFLLPMGPSPPAEGSTGDEPK | 71.05944 | 106.3702 | 5.48227 | 205 |
| **Etanercept** | SQHTQPTPEPSTAPSTSFLLPMGPSPPAEGSTGDEPK | 39.34841 | 59.04271 | 2.77329 | 226 |
| **Etanercept** | SQHTQPTPEPSTAPST | 11.52598 | 17.73228 | 0 | 217 |
| **Etanercept** | SQHTQPTPEPSTAPSTSFLLPMGPSPPAEGSTGDEPK | 72.96007 | 108.9357 | 6.14823 | 205 |
| **Etanercept** | SQHTQPTPEPSTAPSTSFLLPMGPSPPAEGSTGDEPK | 82.16898 | 120.0427 | 11.83213 | 205 |
| **Etanercept** | LPAQVAFTPYAPEPGSTCR | 98.45328 | 148.2446 | 5.98361 | 8 |
| **Etanercept** | SQHTQPTPEPSTAPSTSFLLPMGPSPPAEGSTGDEPK | 41.15351 | 58.15063 | 9.58743 | 205 |
| **Etanercept** | SQHTQPTPEPSTAPSTSFLLPMGPSPPAEGSTGDEPK | 50.10632 | 74.37168 | 5.04207 | 205 |
| **Etanercept** | SQHTQPTPEPSTAPSTSFLLPMGPSPPAEGSTGDEPK | 64.95664 | 96.77702 | 5.86164 | 205 |
| **Etanercept** | SQHTQPTPEPSTAPSTSFLLPMGPSPPAEGSTGDEPK | 64.44352 | 95.92122 | 5.98492 | 217 |
| **Etanercept** | SQHTQPTPEPSTAPSTSFLLPMGPSPPAEGSTGDEPK | 81.72915 | 119.718 | 11.17852 | 205 |
| **Etanercept** | SQHTQPTPEPSTAPSTSFLLPMGPSPPAEGSTGDEPK | 64.07178 | 95.83396 | 5.08489 | 205 |
| **Etanercept** | HTQPTPEPSTAPSTSFLLPMGPSPPAEGSTGDEPK | 15.3078 | 23.55046 | 0 | 212 |
| **Etanercept** | SQHTQPTPEPSTAPSTSFLLPMGPSPPAEGSTGDEPK | 51.34727 | 76.39314 | 4.83351 | 205 |
| **Etanercept** | SQHTQPTPEPSTAPSTSFLLPMGPSPPAEGSTGDEPK | 61.92962 | 92.18434 | 5.74228 | 205 |
| **Etanercept** | SQHTQPTPEPSTAPSTSFLLPMGPSPPAEGSTGDEPK | 27.92359 | 42.95937 | 0 | 213 |
| **Etanercept** | SQHTQPTPEPSTAPSTSFLLPMGPSPPAEGSTGDEPK | 70.15645 | 105.2087 | 5.05938 | 213 |
| **Etanercept** | SQHTQPTPEPSTAPSTSFLLPMGPSPPAEGSTGDEPK | 13.11429 | 18.52837 | 3.05959 | 216 |
| **Etanercept** | SQHTQPTPEPSTAPSTSFLLPMGPSPPAEGSTGDEPK | 97.74074 | 143.5224 | 12.71766 | 218 |
| **Etanercept** | SQHTQPTPEPSTAPSTSFLLPMGPSPPAEGSTGDEPK | 105.826 | 155.5285 | 13.52135 | 226 |
| **Etanercept** | SQHTQPTPEPSTAPSTSFLLPMGPSPPAEGSTGDEPK | 72.49756 | 108.2906 | 6.02476 | 218 |
| **Etanercept** | SQHTQPTPEPSTAPSTSFLLPMGPSPPAEGSTGDEPK | 90.17512 | 133.091 | 10.47418 | 218 |
| **Etanercept** | SQHTQPTPEPSTAPSTSFLLPMGPSPPAEGSTGDEPK | 84.79137 | 127.3916 | 5.67674 | 216 |
| **Etanercept** | SQHTQPTPEPSTAPSTSFLLPMGPSPPAEGSTGDEPK | 66.23907 | 99.15808 | 5.10375 | 205 |

| Sample | Peptide | TotalScore | PepScore | GlyScore | ProSites |
|---|---|---|---|---|---|
| **Etanercept** | SQHTQPTPEPSTAPSTSFLLPMGPSPPAEGSTGDEPK | 63.97294 | 95.72391 | 5.00685 | 205 |
| **Etanercept** | SQHTQPTPEPSTAPSTSFLLPMGPSPPAEGSTGDEPK | 105.5254 | 155.3822 | 12.93416 | 213 |
| **Etanercept** | SQHTQPTPEPSTAPSTSFLLPMGPSPPAEGSTGDEPK | 27.03098 | 41.58612 | 0 | 213 |
| **Etanercept** | SQHTQPTPEPSTAPSTSFLLPMGPSPPAEGSTGDEPK | 86.78043 | 129.8556 | 6.78362 | 212 |
| **Etanercept** | SQHTQPTPEPSTAPSTSFLLPMGPSPPAEGSTGDEPK | 73.46662 | 109.436 | 6.66625 | 212 |
| **Etanercept** | SQHTQPTPEPSTAPSTSFLLPMGPSPPAEGSTGDEPK | 5.26227 | 6.70048 | 2.59132 | 226 |
| **Etanercept** | HTQPTPEPSTAPSTSFLLPMGPSPPAEGSTGDEPK | 36.10574 | 50.39746 | 9.56397 | 226 |
| **Etanercept** | HTQPTPEPSTAPSTSFLLPMGPSPPAEGSTGDEPK | 36.69833 | 55.71819 | 1.37575 | 226 |
| **Etanercept** | SQHTQPTPEPSTAPSTSFLLPMGPSPPAEGSTGDEPK | 82.33361 | 126.6671 | 0 | 212 |
| **Etanercept** | QHTQPTPEPSTAPSTSFLLPMGPSPPAEGSTGDEPK | 12.5503 | 19.30815 | 0 | 217 |
| **Etanercept** | SQHTQPTPEPSTAPSTSFLLPMGPSPPAEGSTGDEPK | 56.97573 | 84.63978 | 5.59962 | 205 |
| **Etanercept** | SQHTQPTPEPSTAPSTSFLLPMGPSPPAEGSTGDEPK | 56.41061 | 86.78556 | 0 | 205 |
| **Etanercept** | HTQPTPEPSTAPSTSFLLPMGPSPPAEGSTGDEPK | 25.6543 | 39.46815 | 0 | 216 |
| **Etanercept** | SQHTQPTPEPSTAPSTSFLLPMGPSPPAEGSTGDEPK | 107.1067 | 160.9451 | 7.12115 | 205 |
| **Etanercept** | SQHTQPTPEPSTAPSTSFLLPMGPSPPAEGSTGDEPK | 123.8559 | 186.602 | 7.32732 | 205 |
| **Etanercept** | SQHTQPTPEPSTAPSTSFLLPMGPSPPAEGSTGDEPK | 75.82516 | 113.9942 | 4.93979 | 205 |
| **Etanercept** | SQHTQPTPEPSTAPSTSFLLPMGPSPPAEGSTGDEPK | 86.36902 | 129.1895 | 6.84522 | 205 |
| **Etanercept** | NQVSLTCLVKGFYPSDIAVEWESNGQPENNY | 2.65877 | 3.42862 | 1.22905 | 395 |
| **Etanercept** | SQHTQPTPEPSTAPSTSFLLPMGPSPPAEGSTGDEPK | 89.56022 | 134.1524 | 6.74612 | 205 |
| **Etanercept** | SQHTQPTPEPSTAPSTSFLLPMGPSPPAEGSTGDEPK | 81.248 | 119.3761 | 10.43875 | 205 |
| **Etanercept** | NQVSLTCLVKGFYPSDIAVEWESNGQPENNY | 2.22846 | 3.4284 | 0 | 395 |
| **Etanercept** | SQHTQPTPEPSTAPSTSFLLPMGPSPPAEGSTGDEPK | 58.10109 | 86.40929 | 5.52872 | 205 |
| **Etanercept** | SQHTQPTPEPSTAPSTSFLLPMGPSPPAEGSTGDEPK | 49.4252 | 73.1573 | 5.35131 | 213 |
| **Etanercept** | SQHTQPTPEPSTAPSTSFLLPMGPSPPAEGSTGDEPK | 55.91278 | 83.09565 | 5.4303 | 205 |
| **Etanercept** | SQHTQPTPEPSTAPSTSFLLPMGPSPPAEGSTGDEPK | 101.8588 | 153.7352 | 5.51682 | 205 |
| **Etanercept** | SQHTQPTPEPSTAPSTSFLLPMGPSPPAEGSTGDEPK | 52.40535 | 77.82319 | 5.2008 | 205 |
| **Etanercept** | THTCPPCPAPELLGGPSVFLFPPKPK | 53.42978 | 72.47051 | 18.06842 | 245 |
| **Etanercept** | THTCPPCPAPELLGGPSVFLFPPKPK | 73.35436 | 111.4112 | 2.67741 | 245 |
| **Etanercept** | THTCPPCPAPELLGGPSVFLFPPKPK | 59.40779 | 89.92647 | 2.73025 | 245 |
| **Etanercept** | THTCPPCPAPELLGGPSVFLFPPKPK | 60.45839 | 91.51588 | 2.78018 | 245 |
| **Etanercept** | THTCPPCPAPELLGGPSVFLFPPKPK | 42.26576 | 61.7946 | 5.99791 | 245 |
| **Etanercept** | THTCPPCPAPELLGGPSVFLFPPKPK | 57.03828 | 81.18318 | 12.19774 | 245 |
| **Etanercept** | THTCPPCPAPELLGGPSVFLFPPKPK | 51.09814 | 68.64524 | 18.51068 | 245 |
| **Etanercept** | THTCPPCPAPELLGGPSVFLFPPKPK | 77.44022 | 108.665 | 19.45143 | 245 |
| **Etanercept** | TQPTPEPSTAPSTSFLLPMGPSPPAEGSTGDEPK | 6.56964 | 9.67461 | 0.80328 | 208 |
| **Etanercept** | SQHTQPTPEPSTAPSTSFLLPMGPSPPAEG | 7.2496 | 10.53357 | 1.1508 | 226 |
| **Etanercept** | STAPSTSFLLPMGPSPPAEGSTGDEPK | 3.4248 | 5.26892 | 0 | 226 |
| **Etanercept** | THTCPPCPAPELLGGPSVFLFPPK | 32.26453 | 48.96943 | 1.24114 | 245 |
| **Etanercept** | THTCPPCPAPELLGGPSVFLFPPK | 37.86561 | 57.55724 | 1.29545 | 245 |
| **Etanercept** | SMAPGAVHLPQPVSTR | 15.31799 | 21.58391 | 3.68129 | 199 |
| **Etanercept** | SMAPGAVHLPQPVSTR | 12.34414 | 18.99098 | 0 | 199 |
| **TNFR: Fc1** | AVHLPQPVSTR | 41.99521 | 59.81475 | 8.90178 | 199 |

| Sample | Peptide | TotalScore | PepScore | GlyScore | ProSites |
|---|---|---|---|---|---|
| **TNFR: Fc1** | AVHLPQPVSTR | 35.50301 | 49.33938 | 9.80691 | 199 |
| **TNFR: Fc1** | GAVHLPQPVSTR | 39.7245 | 56.09266 | 9.3265 | 199 |
| **TNFR: Fc1** | APGAVHLPQPVSTR | 48.99558 | 67.05738 | 15.45224 | 199 |
| **TNFR: Fc1** | SMAPGAVHLPQPVSTR | 50.64899 | 71.98206 | 11.03043 | 199 |
| **TNFR: Fc1** | APGAVHLPQPVSTR | 46.46543 | 68.57961 | 5.39625 | 199 |
| **TNFR: Fc1** | APGAVHLPQPVSTR | 55.17563 | 81.76176 | 5.8014 | 199 |
| **TNFR: Fc1** | SMAPGAVHLPQPVSTR | 35.7118 | 52.02615 | 5.41372 | 200 |
| **TNFR: Fc1** | SMAPGAVHLPQPVSTR | 41.8339 | 61.41897 | 5.46163 | 200 |
| **TNFR: Fc1** | APGAVHLPQPVSTR | 69.80128 | 104.2663 | 5.79487 | 199 |
| **TNFR: Fc1** | APGAVHLPQPVSTR | 77.70608 | 116.2246 | 6.17171 | 199 |
| **TNFR: Fc1** | SMAPGAVHLPQPVSTR | 22.8647 | 29.3948 | 10.73737 | 199 |
| **TNFR: Fc1** | SMAPGAVHLPQPVSTR | 37.30295 | 51.49794 | 10.94083 | 199 |
| **TNFR: Fc1** | SQHTQPTPEPSTAPSTSF | 31.04542 | 40.5708 | 13.35541 | 212 |
| **TNFR: Fc1** | SMAPGAVHLPQPVSTR | 41.11102 | 57.17922 | 11.27008 | 199 |
| **TNFR: Fc1** | SMAPGAVHLPQPVSTR | 38.45284 | 53.21558 | 11.03631 | 199 |
| **TNFR: Fc1** | SMAPGAVHLPQPVSTR | 47.40159 | 69.85774 | 5.69732 | 199 |
| **TNFR: Fc1** | SMAPGAVHLPQPVSTR | 54.57452 | 80.68504 | 6.08355 | 199 |
| **TNFR: Fc1** | SMAPGAVHLPQPVSTR | 87.61097 | 131.1952 | 6.6688 | 199 |
| **TNFR: Fc1** | SMAPGAVHLPQPVSTR | 62.21315 | 92.23632 | 6.45582 | 199 |
| **TNFR: Fc1** | SMAPGAVHLPQPVSTR | 19.61217 | 24.69981 | 10.1637 | 199 |
| **TNFR: Fc1** | SMAPGAVHLPQPVSTR | 41.94471 | 64.53033 | 0 | 199 |
| **TNFR: Fc1** | SMAPGAVHLPQPVSTR | 49.13234 | 72.21377 | 6.26681 | 199 |
| **TNFR: Fc1** | SMAPGAVHLPQPVSTR | 42.06621 | 64.71724 | 0 | 199 |
| **TNFR: Fc1** | SMAPGAVHLPQPVSTR | 36.38072 | 55.97034 | 0 | 199 |
| **TNFR: Fc1** | SMAPGAVHLPQPVSTR | 54.95023 | 81.28858 | 6.03616 | 199 |
| **TNFR: Fc1** | SMAPGAVHLPQPVSTR | 52.32583 | 77.32431 | 5.90009 | 199 |
| **TNFR: Fc1** | SMAPGAVHLPQPVSTR | 47.61778 | 70.09074 | 5.88229 | 199 |
| **TNFR: Fc1** | SMAPGAVHLPQPVSTR | 55.4067 | 78.80713 | 11.94875 | 199 |
| **TNFR: Fc1** | PGAVHLPQPVSTR | 42.65614 | 53.02095 | 23.40721 | 199 |
| **TNFR: Fc1** | SMAPGAVHLPQPVSTR | 63.43232 | 89.70756 | 14.63543 | 199 |
| **TNFR: Fc1** | APGAVHLPQPVSTR | 62.53554 | 84.25465 | 22.20005 | 199 |
| **TNFR: Fc1** | MAPGAVHLPQPVSTR | 23.74168 | 36.52566 | 0 | 199 |
| **TNFR: Fc1** | PGAVHLPQPVSTR | 64.99917 | 83.64925 | 30.36329 | 199 |
| **TNFR: Fc1** | SMAPGAVHLPQPVSTR | 72.96162 | 111.2049 | 1.9384 | 199 |
| **TNFR: Fc1** | SMAPGAVHLPQPVSTR | 52.84884 | 78.23174 | 5.70917 | 200 |
| **TNFR: Fc1** | SMAPGAVHLPQPVSTR | 52.04132 | 76.79506 | 6.07009 | 199 |
| **TNFR: Fc1** | SMAPGAVHLPQPVSTR | 50.7575 | 74.87973 | 5.95906 | 199 |
| **TNFR: Fc1** | SMAPGAVHLPQPVSTR | 53.29845 | 78.88533 | 5.77996 | 199 |
| **TNFR: Fc1** | SMAPGAVHLPQPVSTR | 51.70723 | 72.8724 | 12.4005 | 199 |
| **TNFR: Fc1** | SMAPGAVHLPQPVSTR | 46.07118 | 64.19143 | 12.4193 | 199 |
| **TNFR: Fc1** | SMAPGAVHLPQPVSTR | 51.42002 | 72.75982 | 11.78895 | 199 |
| **TNFR: Fc1** | SMAPGAVHLPQPVSTR | 53.62361 | 79.21588 | 6.0951 | 199 |

| Sample | Peptide | TotalScore | PepScore | GlyScore | ProSites |
|---|---|---|---|---|---|
| **TNFR: Fc1** | SMAPGAVHLPQPVSTR | 52.54307 | 75.61905 | 9.68767 | 199 |
| **TNFR: Fc1** | SMAPGAVHLPQPVSTR | 59.92874 | 89.04293 | 5.85952 | 199 |
| **TNFR: Fc1** | SMAPGAVHLPQPVSTR | 43.29328 | 66.60504 | 0 | 199 |
| **TNFR: Fc1** | APGAVHLPQPVSTR | 61.71738 | 88.34465 | 12.26675 | 199 |
| **TNFR: Fc1** | APGAVHLPQPVSTR | 66.11164 | 95.00767 | 12.44758 | 199 |
| **TNFR: Fc1** | PGAVHLPQPVSTR | 49.68496 | 69.40968 | 13.05332 | 199 |
| **TNFR: Fc1** | PGAVHLPQPVSTR | 52.33974 | 76.5583 | 7.36241 | 199 |
| **TNFR: Fc1** | SMAPGAVHLPQPVSTR | 60.03673 | 89.26573 | 5.75431 | 199 |
| **TNFR: Fc1** | SMAPGAVHLPQPVSTR | 49.71234 | 67.89097 | 15.95204 | 199 |
| **TNFR: Fc1** | SMAPGAVHLPQPVSTR | 57.21659 | 84.88543 | 5.83158 | 199 |
| **TNFR: Fc1** | SMAPGAVHLPQPVSTR | 91.99518 | 137.4655 | 7.5504 | 199 |
| **TNFR: Fc1** | SMAPGAVHLPQPVSTR | 45.37212 | 68.01546 | 3.3202 | 199 |
| **TNFR: Fc1** | SMAPGAVHLPQPVSTR | 86.79104 | 129.7431 | 7.02293 | 199 |
| **TNFR: Fc1** | SMAPGAVHLPQPVSTR | 39.37611 | 57.97852 | 4.82878 | 199 |
| **TNFR: Fc1** | SMAPGAVHLPQPVSTR | 94.90368 | 142.4023 | 6.69192 | 199 |
| **TNFR: Fc1** | SMAPGAVHLPQPVSTR | 31.17643 | 47.96374 | 0 | 199 |
| **TNFR: Fc1** | REPQVYTLPPSR | 5.45703 | 7.69463 | 1.30148 | 370 |
| **TNFR: Fc1** | SMAPGAVHLPQPVSTR | 74.58571 | 111.2851 | 6.42973 | 199 |
| **TNFR: Fc1** | REPQVYTLPPSR | 5.01639 | 7.054 | 1.23225 | 370 |
| **TNFR: Fc1** | SMAPGAVHLPQPVSTR | 76.34284 | 114.178 | 6.07762 | 199 |
| **TNFR: Fc1** | REPQVYTLPPSR | 4.24434 | 6.52975 | 0 | 370 |
| **TNFR: Fc1** | SMAPGAVHLPQPVSTR | 73.52994 | 109.6781 | 6.39759 | 199 |
| **TNFR: Fc1** | SMAPGAVHLPQPVSTR | 64.45861 | 95.87001 | 6.12315 | 199 |
| **TNFR: Fc1** | SMAPGAVHLPQPVSTR | 61.35501 | 91.18556 | 5.95542 | 199 |
| **TNFR: Fc1** | SMAPGAVHLPQPVSTR | 52.80454 | 78.08451 | 5.85602 | 199 |
| **TNFR: Fc1** | SMAPGAVHLPQPVSTR | 52.69973 | 77.98366 | 5.74387 | 199 |
| **TNFR: Fc1** | EPQVYTLPPSREEMTK | 18.85613 | 20.73462 | 15.36751 | 370 |
| **TNFR: Fc1** | SMAPGAVHLPQPVSTR | 26.80985 | 38.59965 | 4.9145 | 199 |
| **TNFR: Fc1** | SMAPGAVHLPQPVSTR | 50.63397 | 74.89366 | 5.58027 | 199 |
| **TNFR: Fc1** | SMAPGAVHLPQPVSTR | 53.37308 | 79.25801 | 5.30106 | 199 |
| **TNFR: Fc1** | SMAPGAVHLPQPVSTR | 47.41424 | 70.09449 | 5.29378 | 199 |
| **TNFR: Fc1** | SMAPGAVHLPQPVSTR | 47.79731 | 70.6583 | 5.34119 | 199 |
| **TNFR: Fc1** | SMAPGAVHLPQPVSTR | 48.28767 | 71.42964 | 5.30974 | 199 |
| **TNFR: Fc1** | AQVAFTPYAPEPGSTCR | 10.17442 | 12.90204 | 5.10882 | 8 |
| **TNFR: Fc1** | SQHTQPTPEPSTAPSTSFLLPMGPSPPAEGSTGDEPK | 42.66031 | 63.06119 | 4.77294 | 205 |
| **TNFR: Fc1** | SQHTQPTPEPSTAPSTSFLLPMGPSPPAEGSTGDEPK | 76.68389 | 112.2288 | 10.67195 | 217 |
| **TNFR: Fc1** | SQHTQPTPEPSTAPSTSFLLPMGPSPPAEGSTGDEPK | 86.62118 | 129.9306 | 6.18933 | 217 |
| **TNFR: Fc1** | QPTPEPSTAPSTSFLLPMGPSPPAEGSTGDEPK | 14.53631 | 22.36355 | 0 | 208 |
| **TNFR: Fc1** | SQHTQPTPEPSTAPSTSFLLPMGPSPPAEGSTGDEPK | 72.66392 | 108.7943 | 5.56465 | 213 |
| **TNFR: Fc1** | SQHTQPTPEPSTAPSTSFLLPMGPSPPAEGSTGDEPK | 87.08469 | 133.9765 | 0 | 213 |
| **TNFR: Fc1** | SQHTQPTPEPSTAPSTSFLLPMGPSPPAEGSTGDEPK | 97.47796 | 143.2223 | 12.52419 | 205 |
| **TNFR: Fc1** | SQHTQPTPEPSTAPSTSFLLPMGPSPPAEGSTGDEPK | 99.37975 | 149.3294 | 6.61613 | 205 |

| Sample | Peptide | TotalScore | PepScore | GlyScore | ProSites |
|---|---|---|---|---|---|
| **TNFR: Fc1** | SQHTQPTPEPSTAPSTSFLLPMGPSPPAEGSTGDEPK | 88.50986 | 136.169 | 0 | 205 |
| **TNFR: Fc1** | SQHTQPTPEPSTAPSTSFLLPMGPSPPAEGSTGDEPK | 77.42204 | 115.8164 | 6.11828 | 205 |
| **TNFR: Fc1** | SQHTQPTPEPSTAPSTSFLLPMGPSPPAEGSTGDEPK | 76.46603 | 111.1344 | 12.08195 | 205 |
| **TNFR: Fc1** | SQHTQPTPEPSTAPSTSFLLPMGPSPPAEGSTGDEPK | 61.53593 | 91.32217 | 6.21863 | 217 |
| **TNFR: Fc1** | SQHTQPTPEPSTAPSTSFLLPMGPSPPAEGSTGDEPK | 91.36854 | 137.0258 | 6.57651 | 213 |
| **TNFR: Fc1** | SQHTQPTPEPSTAPSTSFLLPMGPSPPAEGSTGDEPK | 68.80101 | 100.0794 | 10.71252 | 213 |
| **TNFR: Fc1** | SQHTQPTPEPSTAPSTSFLLPMGPSPPAEGSTGDEPK | 32.06128 | 49.32504 | 0 | 205 |
| **TNFR: Fc1** | SQHTQPTPEPSTAPSTSFLLPMGPSPPAEGSTGDEPK | 82.57268 | 120.7474 | 11.6768 | 205 |
| **TNFR: Fc1** | LPAQVAFTPYAPEPGSTCR | 112.1472 | 169.0612 | 6.44989 | 8 |
| **TNFR: Fc1** | LPAQVAFTPYAPEPGSTCR | 134.9558 | 204.0949 | 6.55451 | 8 |
| **TNFR: Fc1** | LPAQVAFTPYAPEPGSTCR | 40.15855 | 55.07343 | 12.45949 | 8 |
| **TNFR: Fc1** | SQHTQPTPEPSTAPSTSFLLPMGPSPPAEGSTGDEPK | 66.9923 | 99.89751 | 5.88262 | 205 |
| **TNFR: Fc1** | SQHTQPTPEPSTAPSTSFLLPMGPSPPAEGSTGDEPK | 66.5899 | 99.55063 | 5.37711 | 213 |
| **TNFR: Fc1** | SQHTQPTPEPSTAPSTSFLLPMGPSPPAEGSTGDEPK | 54.94439 | 84.52983 | 0 | 217 |
| **TNFR: Fc1** | SQHTQPTPEPSTAPSTSFLLPMGPSPPAEGSTGDEPK | 62.16883 | 92.55824 | 5.73136 | 205 |
| **TNFR: Fc1** | SQHTQPTPEPSTAPSTSFLLPMGPSPPAEGSTGDEPK | 66.43681 | 96.82968 | 9.99292 | 205 |
| **TNFR: Fc1** | SQHTQPTPEPSTAPSTSFLLPMGPSPPAEGSTGDEPK | 55.42865 | 82.33916 | 5.45199 | 205 |
| **TNFR: Fc1** | SQHTQPTPEPSTAPSTSFLLPMGPSPPAEGSTGDEPK | 62.79998 | 91.24877 | 9.96652 | 205 |
| **TNFR: Fc1** | SQHTQPTPEPSTAPSTSFLLPMGPSPPAEGSTGDEPK | 66.86525 | 99.70733 | 5.87281 | 213 |
| **TNFR: Fc1** | SQHTQPTPEPSTAPSTSFLLPMGPSPPAEGSTGDEPK | 67.81099 | 101.3927 | 5.44505 | 205 |
| **TNFR: Fc1** | SQHTQPTPEPSTAPSTSFLLPMGPSPPAEGSTGDEPK | 73.51264 | 110.0725 | 5.61576 | 208 |
| **TNFR: Fc1** | SQHTQPTPEPSTAPSTSFLLPMGPSPPAEGSTGDEPK | 66.56681 | 99.58516 | 5.24703 | 205 |
| **TNFR: Fc1** | SQHTQPTPEPSTAPSTSFLLPMGPSPPAEGSTGDEPK | 38.81531 | 54.31766 | 10.02523 | 213 |
| **TNFR: Fc1** | SQHTQPTPEPSTAPSTSFLLPMGPSPPAEGSTGDEPK | 56.44651 | 81.75834 | 9.43882 | 205 |
| **TNFR: Fc1** | SQHTQPTPEPSTAPSTSFLLPMGPSPPAEGSTGDEPK | 58.49447 | 86.84358 | 5.84614 | 216 |
| **TNFR: Fc1** | SQHTQPTPEPSTAPSTSFLLPMGPSPPAEGSTGDEPK | 85.08949 | 123.9301 | 12.95689 | 226 |
| **TNFR: Fc1** | SQHTQPTPEPSTAPSTSFLLPMGPSPPAEGSTGDEPK | 74.48036 | 111.3355 | 6.03506 | 218 |
| **TNFR: Fc1** | SQHTQPTPEPSTAPSTSFLLPMGPSPPAEGSTGDEPK | 69.67224 | 103.9796 | 5.95856 | 216 |
| **TNFR: Fc1** | SQHTQPTPEPSTAPSTSFLLPMGPSPPAEGSTGDEPK | 68.88076 | 103.1903 | 5.16298 | 205 |
| **TNFR: Fc1** | SQHTQPTPEPSTAPSTSFLLPMGPSPPAEGSTGDEPK | 68.24748 | 104.9961 | 0 | 213 |
| **TNFR: Fc1** | SQHTQPTPEPSTAPSTSFLLPMGPSPPAEGSTGDEPK | 114.0878 | 171.8259 | 6.85989 | 217 |
| **TNFR: Fc1** | SQHTQPTPEPSTAPSTSFLLPMGPSPPAEGSTGDEPK | 78.06002 | 116.5517 | 6.5754 | 213 |
| **TNFR: Fc1** | SQHTQPTPEPSTAPSTSFLLPMGPSPPAEGSTGDEPK | 59.40645 | 88.57457 | 5.23707 | 205 |
| **TNFR: Fc1** | SQHTQPTPEPSTAPSTSFLLPMGPSPPAEGSTGDEPK | 102.1118 | 153.3872 | 6.88606 | 212 |
| **TNFR: Fc1** | SQHTQPTPEPSTAPSTSFLLPMGPSPPAEGSTGDEPK | 75.67635 | 116.4252 | 0 | 212 |
| **TNFR: Fc1** | SQHTQPTPEPSTAPSTSFLLPMGPSPPAEGSTGDEPK | 127.8386 | 192.9237 | 6.96629 | 205 |
| **TNFR: Fc1** | SQHTQPTPEPSTAPSTSFLLPMGPSPPAEGSTGDEPK | 64.98878 | 97.12385 | 5.30936 | 205 |
| **TNFR: Fc1** | SQHTQPTPEPSTAPSTSFLLPMGPSPPAEGSTGDEPK | 78.2213 | 120.3405 | 0 | 205 |
| **TNFR: Fc1** | HTQPTPEPSTAPSTSFLLPMGPSPPAEGSTGDEPK | 12.17328 | 18.19802 | 0.98449 | 205 |
| **TNFR: Fc1** | QHTQPTPEPSTAPSTSFLLPMGPSPPAEGSTGDEPK | 15.59081 | 23.98586 | 0 | 217 |
| **TNFR: Fc1** | SQHTQPTPEPSTAPSTSFLLPMGPSPPAEGSTGDEPK | 86.77076 | 129.7364 | 6.97737 | 205 |
| **TNFR: Fc1** | QHTQPTPEPSTAPSTSFLLPMGPSPPAEGSTGDEPK | 14.1991 | 21.84477 | 0 | 217 |

| Sample | Peptide | TotalScore | PepScore | GlyScore | ProSites |
|---|---|---|---|---|---|
| **TNFR: Fc1** | SQHTQPTPEPSTAPSTSFLLPMGPSPPAEGSTGDEPK | 87.40503 | 130.8527 | 6.71656 | 218 |
| **TNFR: Fc1** | SQHTQPTPEPSTAPSTSFLLPMGPSPPAEGSTGDEPK | 79.78567 | 115.6719 | 13.13991 | 226 |
| **TNFR: Fc1** | SQHTQPTPEPSTAPSTSFLLPMGPSPPAEGSTGDEPK | 80.44719 | 117.1892 | 12.21208 | 226 |
| **TNFR: Fc1** | SQHTQPTPEPSTAPSTSFLLPMGPSPPAEGSTGDEPK | 81.21451 | 118.4976 | 11.97445 | 218 |
| **TNFR: Fc1** | SQHTQPTPEPSTAPSTSFLLPMGPSPPAEGSTGDEPK | 65.75284 | 97.96889 | 5.92304 | 217 |
| **TNFR: Fc1** | SQHTQPTPEPSTAPSTSFLLPMGPSPPAEGSTGDEPK | 67.29167 | 100.3857 | 5.83142 | 205 |
| **TNFR: Fc1** | SQHTQPTPEPSTAPSTSFLLPMGPSPPAEGSTGDEPK | 70.96419 | 106.0748 | 5.75882 | 218 |
| **TNFR: Fc1** | SQHTQPTPEPSTAPSTSFLLPMGPSPPAEGSTGDEPK | 68.35467 | 102.1481 | 5.59547 | 218 |
| **TNFR: Fc1** | SQHTQPTPEPSTAPSTSFLLPMGPSPPAEGSTGDEPK | 86.0115 | 129.197 | 5.80985 | 205 |
| **TNFR: Fc1** | HTQPTPEPSTAPSTSFLLPMGPSPPAEGSTGDEPK | 21.75034 | 30.78491 | 4.97187 | 205 |
| **TNFR: Fc1** | HTQPTPEPSTAPSTSFLLPMGPSPPAEGSTGDEPK | 36.94107 | 54.25659 | 4.78367 | 205 |
| **TNFR: Fc1** | SQHTQPTPEPSTAPSTSFLLPMGPSPPAEGSTGDEPK | 52.33575 | 77.56867 | 5.47461 | 205 |
| **TNFR: Fc1** | SQHTQPTPEPSTAPSTSFLLPMGPSPPAEGSTGDEPK | 57.12987 | 87.89211 | 0 | 205 |
| **TNFR: Fc1** | HTQPTPEPSTAPSTSFLLPMGPSPPAEGSTGDEPK | 24.80995 | 38.16915 | 0 | 205 |
| **TNFR: Fc1** | SQHTQPTPEPSTAPSTSFLLPMGPSPPAEGSTGDEPK | 53.76396 | 79.84692 | 5.32418 | 205 |
| **TNFR: Fc1** | SQHTQPTPEPSTAPSTSFLLPMGPSPPAEGSTGDEPK | 15.09638 | 23.22521 | 0 | 205 |
| **TNFR: Fc1** | HTQPTPEPSTAPSTSFLLPMGPSPPAEGSTGDEPK | 7.97124 | 12.26344 | 0 | 205 |
| **TNFR: Fc1** | SQHTQPTPEPSTAPSTSFLLPMGPSPPAEGSTGDEPK | 52.53524 | 80.82345 | 0 | 205 |
| **TNFR: Fc1** | THTCPPCPAPELLGGPSVFLFPPKPK | 50.62336 | 68.56464 | 17.30385 | 245 |
| **TNFR: Fc1** | THTCPPCPAPELLGGPSVFLFPPKPK | 24.30605 | 36.00229 | 2.58445 | 245 |
| **TNFR: Fc1** | THTCPPCPAPELLGGPSVFLFPPKPK | 48.92005 | 69.10561 | 11.43258 | 245 |
| **TNFR: Fc1** | THTCPPCPAPELLGGPSVFLFPPKPK | 97.82277 | 144.5827 | 10.98294 | 245 |
| **TNFR: Fc1** | THTCPPCPAPELLGGPSVFLFPPKPK | 90.73278 | 133.6438 | 11.04081 | 245 |
| **TNFR: Fc1** | TQPTPEPSTAPSTSFLLPMGPSPPAEGSTGDEPK | 18.20364 | 28.00559 | 0 | 205 |
| **TNFR: Fc1** | SQHTQPTPEPSTAPSTSFLLPMGPSPPAEGSTGDEPK | 6.72116 | 10.34025 | 0 | 202 |
| **TNFR: Fc1** | SMAPGAVHLPQPVSTR | 33.67808 | 49.66057 | 3.99631 | 199 |
| **TNFR: Fc1** | SMAPGAVHLPQPVSTR | 6.34777 | 7.44132 | 4.3169 | 199 |
| **TNFR: Fc1** | SMAPGAVHLPQPVSTR | 38.87059 | 57.56175 | 4.15843 | 199 |
| **TNFR: Fc1** | SQHTQPTPEPSTAPSTSFLLPMGPSPPAEGSTGDEPK | 4.10315 | 6.31254 | 0 | 202 |
| **TNFR: Fc1** | SMAPGAVHLPQPVSTR | 37.28126 | 55.31193 | 3.79571 | 199 |
| **TNFR: Fc1** | SQHTQPTPEPSTAPSTSFLLPMGPSPPAEGSTGDEPK | 5.77805 | 8.8893 | 0 | 205 |
| **TNFR: Fc1** | SMAPGAVHLPQPVSTR | 38.53949 | 57.1248 | 4.02393 | 199 |
| **TNFR: Fc1** | SQHTQPTPEPSTAPSTSFLLPMGPSPPAEGSTGDEPK | 9.21826 | 14.18194 | 0 | 202 |
| **TNFR: Fc1** | SQHTQPTPEPSTAPSTSFLLPMGPSPPAEGSTGDEPK | 4.15786 | 6.3967 | 0 | 202 |
| **TNFR: Fc1** | SQHTQPTPEPSTAPSTSFLLPMGPSPPAEGSTGDEPK | 4.46512 | 6.86941 | 0 | 205 |
| **TNFR: Fc1** | SQHTQPTPEPSTAPSTSFLLPMGPSPPAEGSTGDEPK | 5.71586 | 6.86932 | 3.57374 | 202 |
| **TNFR: Fc1** | AVHLPQPVSTR | 45.11714 | 64.08879 | 9.88408 | 199 |
| **TNFR: Fc1** | GAVHLPQPVSTR | 29.53914 | 42.51281 | 5.44519 | 199 |
| **TNFR: Fc1** | APGAVHLPQPVSTR | 43.56662 | 58.80999 | 15.25752 | 199 |
| **TNFR: Fc1** | SMAPGAVHLPQPVSTR | 49.99704 | 71.04368 | 10.91042 | 199 |
| **TNFR: Fc1** | SMAPGAVHLPQPVSTR | 51.60988 | 73.12766 | 11.6483 | 199 |
| **TNFR: Fc1** | APGAVHLPQPVSTR | 46.43835 | 68.46541 | 5.53096 | 199 |

| Sample | Peptide | TotalScore | PepScore | GlyScore | ProSites |
|---|---|---|---|---|---|
| **TNFR: Fc1** | APGAVHLPQPVSTR | 64.46638 | 96.21326 | 5.5079 | 199 |
| **TNFR: Fc1** | SMAPGAVHLPQPVSTR | 39.89314 | 58.43894 | 5.45095 | 200 |
| **TNFR: Fc1** | SMAPGAVHLPQPVSTR | 43.41676 | 63.85428 | 5.46137 | 200 |
| **TNFR: Fc1** | SMAPGAVHLPQPVSTR | 7.85472 | 12.08419 | 0 | 199 |
| **TNFR: Fc1** | SMAPGAVHLPQPVSTR | 22.14835 | 28.49576 | 10.36031 | 199 |
| **TNFR: Fc1** | SMAPGAVHLPQPVSTR | 36.64031 | 50.42216 | 11.04545 | 199 |
| **TNFR: Fc1** | SMAPGAVHLPQPVSTR | 30.37907 | 43.90584 | 5.25793 | 200 |
| **TNFR: Fc1** | SQHTQPTPEPSTAPSTSF | 17.13374 | 21.91744 | 8.24972 | 212 |
| **TNFR: Fc1** | SMAPGAVHLPQPVSTR | 39.98198 | 55.4551 | 11.2462 | 199 |
| **TNFR: Fc1** | SMAPGAVHLPQPVSTR | 50.91881 | 75.33429 | 5.57577 | 199 |
| **TNFR: Fc1** | SMAPGAVHLPQPVSTR | 46.06742 | 67.83678 | 5.63861 | 199 |
| **TNFR: Fc1** | SMAPGAVHLPQPVSTR | 98.42526 | 147.7389 | 6.84286 | 199 |
| **TNFR: Fc1** | SMAPGAVHLPQPVSTR | 65.61657 | 97.45631 | 6.4856 | 199 |
| **TNFR: Fc1** | SMAPGAVHLPQPVSTR | 44.36117 | 65.08861 | 5.86737 | 199 |
| **TNFR: Fc1** | SMAPGAVHLPQPVSTR | 28.20095 | 43.38608 | 0 | 199 |
| **TNFR: Fc1** | SMAPGAVHLPQPVSTR | 54.98428 | 81.36648 | 5.98878 | 199 |
| **TNFR: Fc1** | SMAPGAVHLPQPVSTR | 46.41557 | 68.2941 | 5.78402 | 199 |
| **TNFR: Fc1** | SMAPGAVHLPQPVSTR | 52.48058 | 79.905 | 1.54951 | 199 |
| **TNFR: Fc1** | PGAVHLPQPVSTR | 59.93533 | 76.55762 | 29.06537 | 199 |
| **TNFR: Fc1** | APGAVHLPQPVSTR | 53.63728 | 71.09948 | 21.20748 | 199 |
| **TNFR: Fc1** | SMAPGAVHLPQPVSTR | 51.05764 | 75.36748 | 5.9108 | 199 |
| **TNFR: Fc1** | APGAVHLPQPVSTR | 45.1857 | 58.68059 | 20.12374 | 199 |
| **TNFR: Fc1** | SMAPGAVHLPQPVSTR | 46.7646 | 68.90658 | 5.64378 | 200 |
| **TNFR: Fc1** | SMAPGAVHLPQPVSTR | 55.83147 | 82.6649 | 5.99796 | 199 |
| **TNFR: Fc1** | SMAPGAVHLPQPVSTR | 89.83094 | 134.6734 | 6.55215 | 200 |
| **TNFR: Fc1** | SMAPGAVHLPQPVSTR | 58.73703 | 81.29512 | 16.84343 | 199 |
| **TNFR: Fc1** | SMAPGAVHLPQPVSTR | 57.23905 | 84.98652 | 5.70805 | 200 |
| **TNFR: Fc1** | SMAPGAVHLPQPVSTR | 50.83735 | 74.96262 | 6.0333 | 199 |
| **TNFR: Fc1** | SMAPGAVHLPQPVSTR | 49.96196 | 71.64246 | 9.69816 | 199 |
| **TNFR: Fc1** | SMAPGAVHLPQPVSTR | 44.28664 | 65.16498 | 5.51258 | 199 |
| **TNFR: Fc1** | APGAVHLPQPVSTR | 55.41109 | 79.63708 | 10.41998 | 199 |
| **TNFR: Fc1** | SMAPGAVHLPQPVSTR | 54.40002 | 80.6837 | 5.58747 | 199 |
| **TNFR: Fc1** | APGAVHLPQPVSTR | 60.99909 | 90.73828 | 5.76917 | 199 |
| **TNFR: Fc1** | APGAVHLPQPVSTR | 63.41548 | 94.4828 | 5.71905 | 199 |
| **TNFR: Fc1** | SMAPGAVHLPQPVSTR | 38.67271 | 56.70892 | 5.17688 | 199 |
| **TNFR: Fc1** | SMAPGAVHLPQPVSTR | 95.26138 | 143.0753 | 6.46402 | 199 |
| **TNFR: Fc1** | QQGNVFSCSVMHEALHNHYTQK | 16.18494 | 24.6701 | 0.42678 | 444 |
| **TNFR: Fc1** | SMAPGAVHLPQPVSTR | 26.62258 | 35.90678 | 9.3805 | 200 |
| **TNFR: Fc1** | SMAPGAVHLPQPVSTR | 67.87548 | 101.0138 | 6.33285 | 199 |
| **TNFR: Fc1** | SMAPGAVHLPQPVSTR | 64.87573 | 96.688 | 5.79579 | 199 |
| **TNFR: Fc1** | SMAPGAVHLPQPVSTR | 24.57926 | 37.81425 | 0 | 199 |
| **TNFR: Fc1** | SMAPGAVHLPQPVSTR | 47.48408 | 69.79348 | 6.05233 | 199 |

| Sample | Peptide | TotalScore | PepScore | GlyScore | ProSites |
|---|---|---|---|---|---|
| **TNFR: Fc1** | SMAPGAVHLPQPVSTR | 57.05639 | 84.75659 | 5.61316 | 199 |
| **TNFR: Fc1** | SMAPGAVHLPQPVSTR | 64.79055 | 96.70773 | 5.5158 | 199 |
| **TNFR: Fc1** | SMAPGAVHLPQPVSTR | 53.26262 | 79.06778 | 5.33875 | 199 |
| **TNFR: Fc1** | SMAPGAVHLPQPVSTR | 51.71974 | 76.72291 | 5.28528 | 199 |
| **TNFR: Fc1** | SMAPGAVHLPQPVSTR | 43.04894 | 63.34409 | 5.35797 | 199 |
| **TNFR: Fc1** | SMAPGAVHLPQPVSTR | 42.73087 | 62.87297 | 5.3241 | 199 |
| **TNFR: Fc1** | SMAPGAVHLPQPVSTR | 48.12236 | 71.18423 | 5.29318 | 199 |
| **TNFR: Fc1** | SMAPGAVHLPQPVSTR | 24.43421 | 37.59109 | 0 | 199 |
| **TNFR: Fc1** | AQVAFTPYAPEPGSTCR | 9.7601 | 12.25487 | 5.12696 | 8 |
| **TNFR: Fc1** | SMAPGAVHLPQPVSTR | 37.89394 | 55.55105 | 5.10216 | 199 |
| **TNFR: Fc1** | SMAPGAVHLPQPVSTR | 36.416 | 53.2923 | 5.0743 | 199 |
| **TNFR: Fc1** | SMAPGAVHLPQPVSTR | 34.50846 | 50.41306 | 4.97136 | 199 |
| **TNFR: Fc1** | SMAPGAVHLPQPVSTR | 40.2389 | 59.20988 | 5.00709 | 199 |
| **TNFR: Fc1** | SQHTQPTPEPSTAPSTSFLLPMGPSPPAEGSTGDEPK | 76.10629 | 112.5612 | 8.40425 | 217 |
| **TNFR: Fc1** | SQHTQPTPEPSTAPSTSFLLPMGPSPPAEGSTGDEPK | 76.36693 | 114.4098 | 5.71595 | 216 |
| **TNFR: Fc1** | SQHTQPTPEPSTAPSTSFLLPMGPSPPAEGSTGDEPK | 94.70201 | 142.4512 | 6.02504 | 213 |
| **TNFR: Fc1** | SQHTQPTPEPSTAPSTSFLLPMGPSPPAEGSTGDEPK | 92.73354 | 139.3345 | 6.18891 | 217 |
| **TNFR: Fc1** | SQHTQPTPEPSTAPSTSFLLPMGPSPPAEGSTGDEPK | 55.71397 | 80.72059 | 9.2731 | 213 |
| **TNFR: Fc1** | SQHTQPTPEPSTAPSTSFLLPMGPSPPAEGSTGDEPK | 79.15294 | 118.5615 | 5.96557 | 213 |
| **TNFR: Fc1** | SQHTQPTPEPSTAPSTSFLLPMGPSPPAEGSTGDEPK | 78.56546 | 117.3415 | 6.55287 | 205 |
| **TNFR: Fc1** | SQHTQPTPEPSTAPSTSFLLPMGPSPPAEGSTGDEPK | 48.77056 | 72.45078 | 4.79302 | 205 |
| **TNFR: Fc1** | QPTPEPSTAPSTSFLLPMGPSPPAEGSTGDEPK | 12.47045 | 19.18531 | 0 | 208 |
| **TNFR: Fc1** | SQHTQPTPEPSTAPSTSFLLPMGPSPPAEGSTGDEPK | 60.93133 | 93.7405 | 0 | 205 |
| **TNFR: Fc1** | SQHTQPTPEPSTAPSTSFLLPMGPSPPAEGSTGDEPK | 98.95822 | 146.4155 | 10.82325 | 205 |
| **TNFR: Fc1** | QPTPEPSTAPSTSFLLPMGPSPPAEGSTGDEPK | 10.69627 | 16.45581 | 0 | 208 |
| **TNFR: Fc1** | SQHTQPTPEPSTAPSTSFLLPMGPSPPAEGSTGDEPK | 67.60618 | 100.7275 | 6.09508 | 205 |
| **TNFR: Fc1** | SQHTQPTPEPSTAPSTSFLLPMGPSPPAEGSTGDEP | 6.67571 | 10.27032 | 0 | 216 |
| **TNFR: Fc1** | SQHTQPTPEPSTAPSTSFLLPMGPSPPAEGSTGDEPK | 58.43966 | 86.59191 | 6.15691 | 205 |
| **TNFR: Fc1** | SQHTQPTPEPSTAPSTSFLLPMGPSPPAEGSTGDEPK | 81.58536 | 119.0464 | 12.01486 | 205 |
| **TNFR: Fc1** | SQHTQPTPEPSTAPSTSFLLPMGPSPPAEGSTGDEPK | 60.43306 | 89.9909 | 5.53993 | 205 |
| **TNFR: Fc1** | SQHTQPTPEPSTAPSTSFLLPMGPSPPAEGSTGDEPK | 92.45457 | 138.9086 | 6.1829 | 217 |
| **TNFR: Fc1** | LPAQVAFTPYAPEPGSTCR | 117.042 | 176.5909 | 6.45111 | 8 |
| **TNFR: Fc1** | SQHTQPTPEPSTAPSTSFLLPMGPSPPAEGSTGDEPK | 75.93556 | 113.7065 | 5.78955 | 217 |
| **TNFR: Fc1** | SQHTQPTPEPSTAPSTSFLLPMGPSPPAEGSTGDEPK | 61.81882 | 92.27731 | 5.25306 | 205 |
| **TNFR: Fc1** | SQHTQPTPEPSTAPSTSFLLPMGPSPPAEGSTGDEPK | 70.32527 | 105.3901 | 5.20478 | 205 |
| **TNFR: Fc1** | SQHTQPTPEPSTAPSTSFLLPMGPSPPAEGSTGDEPK | 74.70988 | 109.6224 | 9.87235 | 205 |
| **TNFR: Fc1** | SQHTQPTPEPSTAPSTSFLLPMGPSPPAEGSTGDEPK | 66.39885 | 96.85501 | 9.83741 | 213 |
| **TNFR: Fc1** | SQHTQPTPEPSTAPSTSFLLPMGPSPPAEGSTGDEPK | 50.11248 | 66.69607 | 19.31439 | 213 |
| **TNFR: Fc1** | SQHTQPTPEPSTAPSTSFLLPMGPSPPAEGSTGDEPK | 54.53433 | 81.14822 | 5.10854 | 213 |
| **TNFR: Fc1** | SQHTQPTPEPSTAPSTSFLLPMGPSPPAEGSTGDEPK | 71.17305 | 106.5241 | 5.52121 | 213 |
| **TNFR: Fc1** | SQHTQPTPEPSTAPSTSFLLPMGPSPPAEGSTGDEPK | 66.21713 | 99.03781 | 5.26444 | 213 |
| **TNFR: Fc1** | SQHTQPTPEPSTAPSTSFLLPMGPSPPAEGSTGDEPK | 95.60308 | 144.1694 | 5.40852 | 205 |

| Sample | Peptide | TotalScore | PepScore | GlyScore | ProSites |
|---|---|---|---|---|---|
| **TNFR: Fc1** | SQHTQPTPEPSTAPSTSFLLPMGPSPPAEGSTGDEPK | 75.57952 | 113.4491 | 5.25037 | 205 |
| **TNFR: Fc1** | SQHTQPTPEPSTAPSTSFLLPMGPSPPAEGSTGDEPK | 63.62128 | 95.10891 | 5.14426 | 213 |
| **TNFR: Fc1** | SQHTQPTPEPSTAPSTSFLLPMGPSPPAEGSTGDEPK | 49.92353 | 71.74732 | 9.39362 | 226 |
| **TNFR: Fc1** | SQHTQPTPEPSTAPST | 8.40024 | 12.92345 | 0 | 217 |
| **TNFR: Fc1** | SQHTQPTPEPSTAPSTSFLLPMGPSPPAEGSTGDEPK | 74.80734 | 115.0882 | 0 | 205 |
| **TNFR: Fc1** | SQHTQPTPEPSTAPSTSFLLPMGPSPPAEGSTGDEPK | 70.57637 | 105.8427 | 5.08182 | 213 |
| **TNFR: Fc1** | SQHTQPTPEPSTAPSTSFLLPMGPSPPAEGSTGDEPK | 54.41678 | 80.88363 | 5.26406 | 205 |
| **TNFR: Fc1** | SQHTQPTPEPSTAPSTSFLLPMGPSPPAEGSTGDEPK | 46.43334 | 69.25108 | 4.05753 | 213 |
| **TNFR: Fc1** | SQHTQPTPEPSTAPSTSFLLPMGPSPPAEGSTGDEPK | 42.46122 | 62.69937 | 4.87609 | 205 |
| **TNFR: Fc1** | SQHTQPTPEPSTAPSTSFLLPMGPSPPAEGSTGDEPK | 94.02689 | 137.8347 | 12.66952 | 226 |
| **TNFR: Fc1** | SQHTQPTPEPSTAPSTSFLLPMGPSPPAEGSTGDEPK | 87.90526 | 132.0735 | 5.87861 | 218 |
| **TNFR: Fc1** | SQHTQPTPEPSTAPSTSFLLPMGPSPPAEGSTGDEPK | 74.51523 | 111.4996 | 5.82994 | 216 |
| **TNFR: Fc1** | SQHTQPTPEPSTAPSTSFLLPMGPSPPAEGSTGDEPK | 114.2454 | 172.0814 | 6.8356 | 212 |
| **TNFR: Fc1** | SQHTQPTPEPSTAPSTSFLLPMGPSPPAEGSTGDEPK | 73.33146 | 109.6098 | 5.95749 | 213 |
| **TNFR: Fc1** | SQHTQPTPEPSTAPSTSFLLPMGPSPPAEGSTGDEPK | 50.99428 | 73.20624 | 9.74349 | 205 |
| **TNFR: Fc1** | SQHTQPTPEPSTAPSTSFLLPMGPSPPAEGSTGDEPK | 60.90713 | 90.6638 | 5.64476 | 205 |
| **TNFR: Fc1** | SQHTQPTPEPSTAPSTSFLLPMGPSPPAEGSTGDEPK | 30.698 | 47.2277 | 0 | 205 |
| **TNFR: Fc1** | SQHTQPTPEPSTAPSTSFLLPMGPSPPAEGSTGDEPK | 127.9021 | 196.7725 | 0 | 205 |
| **TNFR: Fc1** | SQHTQPTPEPSTAPSTSFLLPMGPSPPAEGSTGDEPK | 62.73068 | 92.78005 | 6.9247 | 205 |
| **TNFR: Fc1** | SQHTQPTPEPSTAPSTSFLLPMGPSPPAEGSTGDEPK | 134.5379 | 203.0549 | 7.29193 | 205 |
| **TNFR: Fc1** | QHTQPTPEPSTAPSTSFLLPMGPSPPAEGSTGDEPK | 12.5355 | 19.28539 | 0 | 217 |
| **TNFR: Fc1** | SQHTQPTPEPSTAPSTSFLLPMGPSPPAEGSTGDEPK | 81.97925 | 122.3305 | 7.04121 | 205 |
| **TNFR: Fc1** | SQHTQPTPEPSTAPSTSFLLPMGPSPPAEGSTGDEPK | 48.06535 | 73.9467 | 0 | 205 |
| **TNFR: Fc1** | QHTQPTPEPSTAPSTSFLLPMGPSPPAEGSTGDEPK | 10.75667 | 16.54872 | 0 | 217 |
| **TNFR: Fc1** | SQHTQPTPEPSTAPSTSFLLPMGPSPPAEGSTGDEPK | 89.66305 | 134.3374 | 6.69649 | 205 |
| **TNFR: Fc1** | SQHTQPTPEPSTAPSTSFLLPMGPSPPAEGSTGDEPK | 74.99992 | 109.0557 | 11.75353 | 218 |
| **TNFR: Fc1** | SQHTQPTPEPSTAPSTSFLLPMGPSPPAEGSTGDEPK | 86.84417 | 126.7213 | 12.78669 | 226 |
| **TNFR: Fc1** | SQHTQPTPEPSTAPSTSFLLPMGPSPPAEGSTGDEPK | 54.00572 | 80.76479 | 4.3103 | 218 |
| **TNFR: Fc1** | SQHTQPTPEPSTAPSTSFLLPMGPSPPAEGSTGDEPK | 75.82863 | 113.5173 | 5.83535 | 218 |
| **TNFR: Fc1** | HTQPTPEPSTAPSTSFLLPMGPSPPAEGSTGDEPK | 66.67794 | 99.9033 | 4.97369 | 208 |
| **TNFR: Fc1** | SQHTQPTPEPSTAPSTSFLLPMGPSPPAEGSTGDEPK | 89.57979 | 134.8306 | 5.54263 | 205 |
| **TNFR: Fc1** | HTQPTPEPSTAPSTSFLLPMGPSPPAEGSTGDEPK | 69.28341 | 104.0188 | 4.77481 | 205 |
| **TNFR: Fc1** | HTQPTPEPSTAPSTSFLLPMGPSPPAEGSTGDEPK | 7.17524 | 8.6519 | 4.43287 | 216 |
| **TNFR: Fc1** | SQHTQPTPEPSTAPSTSFLLPMGPSPPAEGSTGDEPK | 59.23458 | 88.18627 | 5.46716 | 205 |
| **TNFR: Fc1** | SQHTQPTPEPSTAPSTSFLLPMGPSPPAEGSTGDEPK | 56.0068 | 83.34359 | 5.23847 | 216 |
| **TNFR: Fc1** | HTQPTPEPSTAPSTSFLLPMGPSPPAEGSTGDEPK | 4.27809 | 6.58167 | 0 | 205 |
| **TNFR: Fc1** | THTCPPCPAPELLGGPSVFLFPPKPK | 41.9649 | 58.2383 | 11.74287 | 245 |
| **TNFR: Fc1** | THTCPPCPAPELLGGPSVFLFPPKPK | 43.21791 | 63.06396 | 6.36096 | 245 |
| **TNFR: Fc1** | THTCPPCPAPELLGGPSVFLFPPKPK | 91.74353 | 135.3862 | 10.69291 | 245 |
| **TNFR: Fc1** | THTCPPCPAPELLGGPSVFLFPPKPK | 67.86853 | 100.9156 | 6.49546 | 245 |
| **TNFR: Fc1** | THTCPPCPAPELLGGPSVFLFPPKPK | 68.56597 | 102.0365 | 6.40648 | 245 |
| **TNFR: Fc1** | THTCPPCPAPELLGGPSVFLFPPKPK | 37.47821 | 52.43248 | 9.70599 | 245 |

| Sample | Peptide | TotalScore | PepScore | GlyScore | ProSites |
|---|---|---|---|---|---|
| **TNFR: Fc1** | SMAPGAVHLPQPVSTR | 26.93774 | 39.11394 | 4.32478 | 199 |
| **TNFR: Fc1** | SMAPGAVHLPQPVSTR | 36.14504 | 53.55212 | 3.8176 | 199 |
| **TNFR: Fc1** | SQHTQPTPEPSTAPSTSFLLPMGPSPPAEGSTGDEPK | 9.80778 | 15.0889 | 0 | 208 |
| **TNFR: Fc1** | SMAPGAVHLPQPVSTR | 38.72211 | 57.4568 | 3.92912 | 199 |
| **TNFR: Fc1** | SQHTQPTPEPSTAPSTSFLLPMGPSPPAEGSTGDEPK | 4.58439 | 7.05291 | 0 | 202 |
| **TNFR: Fc1** | SMAPGAVHLPQPVSTR | 11.98748 | 16.587 | 3.44552 | 199 |
| **TNFR: Fc1** | AVHLPQPVSTR | 43.86795 | 62.26044 | 9.71048 | 199 |
| **TNFR: Fc1** | GAVHLPQPVSTR | 22.01563 | 30.93195 | 5.45674 | 199 |
| **TNFR: Fc1** | GAVHLPQPVSTR | 39.9645 | 56.45953 | 9.33087 | 199 |
| **TNFR: Fc1** | APGAVHLPQPVSTR | 49.96127 | 68.5518 | 15.43602 | 199 |
| **TNFR: Fc1** | SMAPGAVHLPQPVSTR | 59.87181 | 85.98805 | 11.37023 | 199 |
| **TNFR: Fc1** | APGAVHLPQPVSTR | 62.11389 | 92.41821 | 5.83444 | 199 |
| **TNFR: Fc1** | APGAVHLPQPVSTR | 70.15884 | 104.8335 | 5.763 | 199 |
| **TNFR: Fc1** | SMAPGAVHLPQPVSTR | 55.72087 | 82.63116 | 5.74461 | 200 |
| **TNFR: Fc1** | SMAPGAVHLPQPVSTR | 29.44364 | 42.41323 | 5.35725 | 200 |
| **TNFR: Fc1** | SQHTQPTPEPSTAPSTSF | 22.3898 | 27.45274 | 12.98722 | 212 |
| **TNFR: Fc1** | SMAPGAVHLPQPVSTR | 51.40991 | 73.07691 | 11.1712 | 199 |
| **TNFR: Fc1** | SMAPGAVHLPQPVSTR | 47.86817 | 67.63432 | 11.1596 | 199 |
| **TNFR: Fc1** | SMAPGAVHLPQPVSTR | 48.55347 | 71.64191 | 5.67492 | 199 |
| **TNFR: Fc1** | SMAPGAVHLPQPVSTR | 51.3898 | 75.83528 | 5.99103 | 199 |
| **TNFR: Fc1** | SMAPGAVHLPQPVSTR | 81.36832 | 121.6571 | 6.54625 | 199 |
| **TNFR: Fc1** | SMAPGAVHLPQPVSTR | 15.07289 | 17.63282 | 10.31873 | 199 |
| **TNFR: Fc1** | SMAPGAVHLPQPVSTR | 60.55252 | 89.69676 | 6.42751 | 199 |
| **TNFR: Fc1** | SMAPGAVHLPQPVSTR | 45.04497 | 66.10687 | 5.93003 | 199 |
| **TNFR: Fc1** | SMAPGAVHLPQPVSTR | 56.81414 | 84.09144 | 6.15629 | 199 |
| **TNFR: Fc1** | SMAPGAVHLPQPVSTR | 50.86757 | 75.01485 | 6.02261 | 199 |
| **TNFR: Fc1** | SMAPGAVHLPQPVSTR | 50.00429 | 73.79107 | 5.82885 | 199 |
| **TNFR: Fc1** | SMAPGAVHLPQPVSTR | 52.04257 | 73.81683 | 11.60466 | 199 |
| **TNFR: Fc1** | PGAVHLPQPVSTR | 68.97997 | 92.91258 | 24.53369 | 199 |
| **TNFR: Fc1** | PGAVHLPQPVSTR | 66.76925 | 86.71287 | 29.73112 | 199 |
| **TNFR: Fc1** | APGAVHLPQPVSTR | 61.18215 | 82.42517 | 21.73082 | 199 |
| **TNFR: Fc1** | SMAPGAVHLPQPVSTR | 51.22658 | 75.6086 | 5.94567 | 199 |
| **TNFR: Fc1** | SMAPGAVHLPQPVSTR | 52.5858 | 77.79718 | 5.76467 | 200 |
| **TNFR: Fc1** | SMAPGAVHLPQPVSTR | 46.37848 | 68.24477 | 5.76966 | 200 |
| **TNFR: Fc1** | SMAPGAVHLPQPVSTR | 58.16839 | 86.26253 | 5.99357 | 199 |
| **TNFR: Fc1** | REPQVYTLPPSR | 6.34425 | 9.0672 | 1.28733 | 374 |
| **TNFR: Fc1** | SMAPGAVHLPQPVSTR | 89.75703 | 134.4552 | 6.74624 | 200 |
| **TNFR: Fc1** | SMAPGAVHLPQPVSTR | 65.31477 | 97.35447 | 5.81248 | 199 |
| **TNFR: Fc1** | SMAPGAVHLPQPVSTR | 43.75075 | 64.04343 | 6.06434 | 199 |
| **TNFR: Fc1** | SMAPGAVHLPQPVSTR | 56.21965 | 83.26468 | 5.99318 | 199 |
| **TNFR: Fc1** | SMAPGAVHLPQPVSTR | 53.12827 | 75.03611 | 12.44228 | 199 |
| **TNFR: Fc1** | SMAPGAVHLPQPVSTR | 20.81765 | 29.48148 | 4.72767 | 199 |

| Sample | Peptide | TotalScore | PepScore | GlyScore | ProSites |
|---|---|---|---|---|---|
| **TNFR: Fc1** | SMAPGAVHLPQPVSTR | 46.501 | 68.40827 | 5.81607 | 199 |
| **TNFR: Fc1** | SMAPGAVHLPQPVSTR | 49.62582 | 70.12485 | 11.55618 | 199 |
| **TNFR: Fc1** | SMAPGAVHLPQPVSTR | 44.58722 | 62.53107 | 11.26293 | 199 |
| **TNFR: Fc1** | SMAPGAVHLPQPVSTR | 57.97639 | 85.9974 | 5.93737 | 199 |
| **TNFR: Fc1** | SMAPGAVHLPQPVSTR | 43.72201 | 64.48851 | 5.15565 | 199 |
| **TNFR: Fc1** | SMAPGAVHLPQPVSTR | 53.8007 | 79.86369 | 5.39799 | 199 |
| **TNFR: Fc1** | APGAVHLPQPVSTR | 64.16805 | 92.26331 | 11.99115 | 199 |
| **TNFR: Fc1** | APGAVHLPQPVSTR | 63.32024 | 90.86736 | 12.16131 | 199 |
| **TNFR: Fc1** | PGAVHLPQPVSTR | 52.97353 | 74.53328 | 12.934 | 199 |
| **TNFR: Fc1** | SMAPGAVHLPQPVSTR | 65.86693 | 92.69496 | 16.04343 | 199 |
| **TNFR: Fc1** | SMAPGAVHLPQPVSTR | 48.09859 | 70.93499 | 5.68813 | 199 |
| **TNFR: Fc1** | SMAPGAVHLPQPVSTR | 48.10886 | 70.97852 | 5.63664 | 199 |
| **TNFR: Fc1** | SMAPGAVHLPQPVSTR | 88.33404 | 131.8158 | 7.58216 | 199 |
| **TNFR: Fc1** | SMAPGAVHLPQPVSTR | 8.28499 | 12.74614 | 0 | 199 |
| **TNFR: Fc1** | APGAVHLPQPVSTR | 55.40158 | 82.26023 | 5.52124 | 199 |
| **TNFR: Fc1** | SMAPGAVHLPQPVSTR | 22.61524 | 29.61451 | 9.61661 | 199 |
| **TNFR: Fc1** | REPQVYTLPPSR | 4.1963 | 6.45585 | 0 | 374 |
| **TNFR: Fc1** | SMAPGAVHLPQPVSTR | 81.50497 | 121.9906 | 6.31732 | 199 |
| **TNFR: Fc1** | REPQVYTLPPSR | 3.58764 | 5.51944 | 0 | 374 |
| **TNFR: Fc1** | SMAPGAVHLPQPVSTR | 70.10976 | 104.4867 | 6.26684 | 199 |
| **TNFR: Fc1** | SMAPGAVHLPQPVSTR | 61.90018 | 91.966 | 6.06367 | 199 |
| **TNFR: Fc1** | SMAPGAVHLPQPVSTR | 59.72066 | 88.70049 | 5.90098 | 199 |
| **TNFR: Fc1** | SMAPGAVHLPQPVSTR | 19.81144 | 30.47913 | 0 | 199 |
| **TNFR: Fc1** | SMAPGAVHLPQPVSTR | 8.14958 | 12.53782 | 0 | 199 |
| **TNFR: Fc1** | SMAPGAVHLPQPVSTR | 58.1983 | 86.48776 | 5.66075 | 199 |
| **TNFR: Fc1** | NWYVDGVEVHNAKTK | 3.33607 | 5.13241 | 0 | 309 |
| **TNFR: Fc1** | EPQVYTLPPSREEMTK | 25.16169 | 30.26636 | 15.68158 | 370 |
| **TNFR: Fc1** | SMAPGAVHLPQPVSTR | 24.04414 | 34.52759 | 4.57488 | 199 |
| **TNFR: Fc1** | SMAPGAVHLPQPVSTR | 56.9075 | 84.56899 | 5.53616 | 199 |
| **TNFR: Fc1** | SMAPGAVHLPQPVSTR | 43.18262 | 63.5016 | 5.44737 | 199 |
| **TNFR: Fc1** | SMAPGAVHLPQPVSTR | 44.49209 | 65.62768 | 5.24028 | 199 |
| **TNFR: Fc1** | SMAPGAVHLPQPVSTR | 47.79645 | 70.73122 | 5.2033 | 199 |
| **TNFR: Fc1** | SMAPGAVHLPQPVSTR | 22.76743 | 35.02682 | 0 | 199 |
| **TNFR: Fc1** | AQVAFTPYAPEPGSTCR | 22.35261 | 34.38863 | 0 | 8 |
| **TNFR: Fc1** | SMAPGAVHLPQPVSTR | 42.79083 | 63.00843 | 5.24386 | 199 |
| **TNFR: Fc1** | SMAPGAVHLPQPVSTR | 37.78735 | 55.47724 | 4.93469 | 199 |
| **TNFR: Fc1** | SMAPGAVHLPQPVSTR | 33.95571 | 49.82037 | 4.49277 | 199 |
| **TNFR: Fc1** | SQHTQPTPEPSTAPSTSFLLPMGPSPPAEGSTGDEPK | 76.81314 | 113.2795 | 9.08994 | 217 |
| **TNFR: Fc1** | SQHTQPTPEPSTAPSTSFLLPMGPSPPAEGSTGDE | 6.86207 | 10.55704 | 0 | 216 |
| **TNFR: Fc1** | SQHTQPTPEPSTAPSTSFLLPMGPSPPAEGSTGDEPK | 74.92517 | 112.2582 | 5.59238 | 217 |
| **TNFR: Fc1** | SQHTQPTPEPSTAPSTSFLLPMGPSPPAEGSTGDEPK | 80.39286 | 120.6399 | 5.64837 | 216 |
| **TNFR: Fc1** | SQHTQPTPEPSTAPSTSFLLPMGPSPPAEGSTGDEPK | 97.80721 | 147.0254 | 6.40209 | 213 |

| Sample | Peptide | TotalScore | PepScore | GlyScore | ProSites |
|---|---|---|---|---|---|
| **TNFR: Fc1** | QPTPEPSTAPSTSFLLPMGPSPPAEGSTGDEPK | 12.58724 | 19.36498 | 0 | 208 |
| **TNFR: Fc1** | SQHTQPTPEPSTAPSTSFLLPMGPSPPAEGSTGDEPK | 86.28109 | 132.7401 | 0 | 213 |
| **TNFR: Fc1** | SQHTQPTPEPSTAPSTSFLLPMGPSPPAEGSTGDEPK | 80.71356 | 121.1418 | 5.63249 | 205 |
| **TNFR: Fc1** | SQHTQPTPEPSTAPSTSFLLPMGPSPPAEGSTGDEPK | 64.85681 | 99.77971 | 0 | 205 |
| **TNFR: Fc1** | SQHTQPTPEPSTAPSTSFLLPMGPSPPAEGSTGDEPK | 91.12161 | 133.9722 | 11.54191 | 205 |
| **TNFR: Fc1** | SQHTQPTPEPSTAPSTSFLLPMGPSPPAEGSTGDEPK | 75.49605 | 112.685 | 6.43088 | 205 |
| **TNFR: Fc1** | SQHTQPTPEPSTAPSTSFLLPMGPSPPAEGSTGDEPK | 77.40479 | 115.6457 | 6.38595 | 205 |
| **TNFR: Fc1** | SQHTQPTPEPSTAPSTSFLLPMGPSPPAEGSTGDEPK | 65.52855 | 97.40644 | 6.32674 | 205 |
| **TNFR: Fc1** | SQHTQPTPEPSTAPSTSFLLPMGPSPPAEGSTGDEPK | 83.22077 | 121.9873 | 11.22583 | 217 |
| **TNFR: Fc1** | SQHTQPTPEPSTAPSTSFLLPMGPSPPAEGSTGDEPK | 79.36705 | 116.1913 | 10.97924 | 217 |
| **TNFR: Fc1** | SQHTQPTPEPSTAPSTSFLLPMGPSPPAEGSTGDEPK | 91.48825 | 133.9593 | 12.61354 | 213 |
| **TNFR: Fc1** | SQHTQPTPEPSTAPSTSFLLPMGPSPPAEGSTGDEPK | 74.12363 | 110.995 | 5.64822 | 213 |
| **TNFR: Fc1** | SQHTQPTPEPSTAPSTSFLLPMGPSPPAEGSTGDEPK | 101.5522 | 152.9598 | 6.08082 | 218 |
| **TNFR: Fc1** | LPAQVAFTPYAPEPGSTCR | 118.2905 | 175.6788 | 11.71232 | 8 |
| **TNFR: Fc1** | SQHTQPTPEPSTAPSTSFLLPMGPSPPAEGSTGDEPK | 15.43019 | 20.53521 | 5.94944 | 205 |
| **TNFR: Fc1** | SQHTQPTPEPSTAPSTSFLLPMGPSPPAEGSTGDEPK | 63.97058 | 95.22545 | 5.92583 | 217 |
| **TNFR: Fc1** | SQHTQPTPEPSTAPSTSFLLPMGPSPPAEGSTGDEPK | 61.33819 | 92.23802 | 3.9528 | 205 |
| **TNFR: Fc1** | SQHTQPTPEPSTAPSTSFLLPMGPSPPAEGSTGDEPK | 76.73975 | 114.8699 | 5.92656 | 213 |
| **TNFR: Fc1** | SQHTQPTPEPSTAPSTSFLLPMGPSPPAEGSTGDEPK | 57.36063 | 85.33717 | 5.40421 | 213 |
| **TNFR: Fc1** | SQHTQPTPEPSTAPSTSFLLPMGPSPPAEGSTGDEPK | 63.50303 | 94.54533 | 5.85304 | 205 |
| **TNFR: Fc1** | SQHTQPTPEPSTAPSTSFLLPMGPSPPAEGSTGDEPK | 72.85711 | 106.8356 | 9.75411 | 205 |
| **TNFR: Fc1** | SQHTQPTPEPSTAPSTSFLLPMGPSPPAEGSTGDEPK | 77.00094 | 115.7444 | 5.04884 | 205 |
| **TNFR: Fc1** | SQHTQPTPEPSTAPSTSFLLPMGPSPPAEGSTGDEPK | 83.12354 | 124.7733 | 5.77406 | 205 |
| **TNFR: Fc1** | SQHTQPTPEPSTAPSTSFLLPMGPSPPAEGSTGDEPK | 70.09801 | 103.3058 | 8.42637 | 205 |
| **TNFR: Fc1** | SQHTQPTPEPSTAPSTSFLLPMGPSPPAEGSTGDEPK | 65.96785 | 98.68537 | 5.20673 | 205 |
| **TNFR: Fc1** | SQHTQPTPEPSTAPSTSFLLPMGPSPPAEGSTGDEPK | 29.54757 | 43.00725 | 4.55102 | 216 |
| **TNFR: Fc1** | SQHTQPTPEPSTAPSTSFLLPMGPSPPAEGSTGDEPK | 57.33674 | 85.30861 | 5.38898 | 205 |
| **TNFR: Fc1** | SQHTQPTPEPSTAPSTSFLLPMGPSPPAEGSTGDEPK | 61.78578 | 95.05504 | 0 | 205 |
| **TNFR: Fc1** | SQHTQPTPEPSTAPSTSFLLPMGPSPPAEGSTGDEPK | 67.40107 | 98.56841 | 9.51886 | 213 |
| **TNFR: Fc1** | SQHTQPTPEPSTAPSTSFLLPMGPSPPAEGSTGDEPK | 64.58057 | 94.50547 | 9.00575 | 218 |
| **TNFR: Fc1** | SQHTQPTPEPSTAPSTSFLLPMGPSPPAEGSTGDEPK | 49.2322 | 73.0253 | 5.04501 | 205 |
| **TNFR: Fc1** | SQHTQPTPEPSTAPSTSFLLPMGPSPPAEGSTGDEPK | 63.08241 | 94.31243 | 5.08379 | 205 |
| **TNFR: Fc1** | SQHTQPTPEPSTAPSTSFLLPMGPSPPAEGSTGDEPK | 52.07393 | 80.11374 | 0 | 205 |
| **TNFR: Fc1** | SQHTQPTPEPSTAPSTSFLLPMGPSPPAEGSTGDEPK | 53.1061 | 79.22103 | 4.60695 | 213 |
| **TNFR: Fc1** | SQHTQPTPEPSTAPSTSFLLPMGPSPPAEGSTGDEPK | 43.75777 | 67.31965 | 0 | 216 |
| **TNFR: Fc1** | LLPMGPSPPAEGSTGDEPK | 8.5534 | 13.15908 | 0 | 226 |
| **TNFR: Fc1** | SQHTQPTPEPSTAPSTSFLLPMGPSPPAEGSTGDEPK | 81.20479 | 121.8743 | 5.67574 | 213 |
| **TNFR: Fc1** | SQHTQPTPEPSTAPSTSFLLPMGPSPPAEGSTGDEPK | 112.102 | 165.2079 | 13.47677 | 226 |
| **TNFR: Fc1** | SQHTQPTPEPSTAPSTSFLLPMGPSPPAEGSTGDEPK | 99.93125 | 146.6614 | 13.14674 | 226 |
| **TNFR: Fc1** | SQHTQPTPEPSTAPSTSFLLPMGPSPPAEGSTGDEPK | 108.4212 | 159.9206 | 12.77934 | 218 |
| **TNFR: Fc1** | SQHTQPTPEPSTAPSTSFLLPMGPSPPAEGSTGDEPK | 54.97852 | 81.76475 | 5.23265 | 213 |
| **TNFR: Fc1** | SQHTQPTPEPSTAPSTSFLLPMGPSPPAEGSTGDEPK | 64.99767 | 97.2822 | 5.04068 | 216 |

| Sample | Peptide | TotalScore | PepScore | GlyScore | ProSites |
|---|---|---|---|---|---|
| **TNFR: Fc1** | SQHTQPTPEPSTAPSTSFLLPMGPSPPAEGSTGDEPK | 96.91805 | 145.5862 | 6.53442 | 216 |
| **TNFR: Fc1** | SQHTQPTPEPSTAPSTSFLLPMGPSPPAEGSTGDEPK | 68.36165 | 105.1718 | 0 | 217 |
| **TNFR: Fc1** | SQHTQPTPEPSTAPSTSFLLPMGPSPPAEGSTGDEPK | 25.70554 | 39.54699 | 0 | 216 |
| **TNFR: Fc1** | SQHTQPTPEPSTAPSTSFLLPMGPSPPAEGSTGDEPK | 60.4345 | 90.13816 | 5.27057 | 205 |
| **TNFR: Fc1** | SQHTQPTPEPSTAPSTSFLLPMGPSPPAEGSTGDEPK | 100.0355 | 147.4394 | 11.99977 | 212 |
| **TNFR: Fc1** | SQHTQPTPEPSTAPSTSFLLPMGPSPPAEGSTGDEPK | 81.91945 | 122.4658 | 6.61912 | 212 |
| **TNFR: Fc1** | SQHTQPTPEPSTAPSTSFLLPMGPSPPAEGSTGDEPK | 94.96505 | 142.8521 | 6.03197 | 213 |
| **TNFR: Fc1** | SQHTQPTPEPSTAPSTSFLLPMGPSPPAEGSTGDEPK | 103.8809 | 156.4445 | 6.26278 | 205 |
| **TNFR: Fc1** | SQHTQPTPEPSTAPSTSFLLPMGPSPPAEGSTGDEPK | 104.0827 | 156.3141 | 7.0814 | 205 |
| **TNFR: Fc1** | SQHTQPTPEPSTAPSTSFLLPMGPSPPAEGSTGDEPK | 51.45437 | 76.33604 | 5.24556 | 205 |
| **TNFR: Fc1** | SQHTQPTPEPSTAPSTSFLLPMGPSPPAEGSTGDEPK | 82.42736 | 126.8113 | 0 | 205 |
| **TNFR: Fc1** | SQHTQPTPEPSTAPSTSFLLPMGPSPPAEGSTGDEPK | 34.05492 | 52.39218 | 0 | 202 |
| **TNFR: Fc1** | SQHTQPTPEPSTAPSTSFLLPMGPSPPAEGSTGDEPK | 69.12926 | 102.6553 | 6.86653 | 205 |
| **TNFR: Fc1** | SQHTQPTPEPSTAPSTSFLLPMGPSPPAEGSTGDEPK | 74.05707 | 106.9091 | 13.0462 | 205 |
| **TNFR: Fc1** | SQHTQPTPEPSTAPSTSFLLPMGPSPPAEGSTGDEPK | 67.99173 | 97.67671 | 12.86248 | 217 |
| **TNFR: Fc1** | SQHTQPTPEPSTAPSTSFLLPMGPSPPAEGSTGDEPK | 77.40647 | 115.6343 | 6.41196 | 218 |
| **TNFR: Fc1** | SQHTQPTPEPSTAPSTSFLLPMGPSPPAEGSTGDEPK | 87.50445 | 131.2078 | 6.34114 | 226 |
| **TNFR: Fc1** | SQHTQPTPEPSTAPSTSFLLPMGPSPPAEGSTGDEPK | 79.746 | 115.8188 | 12.75371 | 226 |
| **TNFR: Fc1** | SQHTQPTPEPSTAPSTSFLLPMGPSPPAEGSTGDEPK | 94.30166 | 138.5284 | 12.16636 | 218 |
| **TNFR: Fc1** | HTQPTPEPSTAPSTSFLLPMGPSPPAEGSTGDEPK | 7.34339 | 11.29752 | 0 | 212 |
| **TNFR: Fc1** | SQHTQPTPEPSTAPSTSFLLPMGPSPPAEGSTGDEPK | 82.21621 | 120.1563 | 11.75609 | 217 |
| **TNFR: Fc1** | SQHTQPTPEPSTAPSTSFLLPMGPSPPAEGSTGDEPK | 63.70124 | 95.00479 | 5.56608 | 218 |
| **TNFR: Fc1** | SQHTQPTPEPSTAPSTSFLLPMGPSPPAEGSTGDEPK | 68.2216 | 101.9709 | 5.54427 | 218 |
| **TNFR: Fc1** | HTQPTPEPSTAPSTSFLLPMGPSPPAEGSTGDEPK | 35.51269 | 52.18391 | 4.55185 | 205 |
| **TNFR: Fc1** | HTQPTPEPSTAPSTSFLLPMGPSPPAEGSTGDEPK | 56.79254 | 84.77178 | 4.83111 | 205 |
| **TNFR: Fc1** | HTQPTPEPSTAPSTSFLLPMGPSPPAEGSTGDEPK | 58.45861 | 87.25979 | 4.97069 | 205 |
| **TNFR: Fc1** | SQHTQPTPEPSTAPSTSFLLPMGPSPPAEGSTGDEPK | 21.59915 | 33.22946 | 0 | 208 |
| **TNFR: Fc1** | HTQPTPEPSTAPSTSFLLPMGPSPPAEGSTGDEPK | 13.83782 | 18.69111 | 4.82457 | 216 |
| **TNFR: Fc1** | SQHTQPTPEPSTAPSTSFLLPMGPSPPAEGSTGDEPK | 54.32398 | 81.36512 | 4.10471 | 205 |
| **TNFR: Fc1** | SQHTQPTPEPSTAPSTSFLLPMGPSPPAEGSTGDEPK | 60.36372 | 90.10289 | 5.13383 | 205 |
| **TNFR: Fc1** | SQHTQPTPEPSTAPSTSFLLPMGPSPPAEGSTGDEPK | 76.37963 | 114.5146 | 5.55765 | 205 |
| **TNFR: Fc1** | SQHTQPTPEPSTAPSTSFLLPMGPSPPAEGSTGDEPK | 39.97536 | 58.87684 | 4.87263 | 205 |
| **TNFR: Fc1** | THTCPPCPAPELLGGPSVFLFPPKPK | 29.03609 | 38.46091 | 11.53285 | 245 |
| **TNFR: Fc1** | THTCPPCPAPELLGGPSVFLFPPKPK | 42.66828 | 56.35428 | 17.25142 | 245 |
| **TNFR: Fc1** | THTCPPCPAPELLGGPSVFLFPPKPK | 99.11952 | 146.7304 | 10.69934 | 245 |
| **TNFR: Fc1** | THTCPPCPAPELLGGPSVFLFPPKPK | 67.59219 | 98.42951 | 10.32289 | 245 |
| **TNFR: Fc1** | THTCPPCPAPELLGGPSVFLFPPKPK | 100.1551 | 148.2688 | 10.80115 | 245 |
| **TNFR: Fc1** | THTCPPCPAPELLGGPSVFLFPPKPK | 70.78588 | 105.3265 | 6.63896 | 245 |
| **TNFR: Fc1** | THTCPPCPAPELLGGPSVFLFPPKPK | 39.9232 | 55.2995 | 11.36722 | 245 |
| **TNFR: Fc1** | SMAPGAVHLPQPVSTR | 30.5008 | 44.59211 | 4.33123 | 199 |
| **TNFR: Fc1** | SMAPGAVHLPQPVSTR | 26.09587 | 38.21231 | 3.5939 | 199 |
| **TNFR: Fc1** | SQHTQPTPEPSTAPSTSFLLPMGPSPPAEGSTGDEPK | 6.29026 | 9.67733 | 0 | 205 |

| Sample | Peptide | TotalScore | PepScore | GlyScore | ProSites |
|---|---|---|---|---|---|
| **TNFR: Fc1** | SMAPGAVHLPQPVSTR | 38.69433 | 57.2886 | 4.1621 | 199 |
| **TNFR: Fc1** | SMAPGAVHLPQPVSTR | 14.25023 | 21.92343 | 0 | 199 |
| **TNFR: Fc1** | SMAPGAVHLPQPVSTR | 33.43682 | 49.30367 | 3.96982 | 199 |
| **TNFR: Fc1** | SQHTQPTPEPSTAPSTSFLLPMGPSPPAEGSTGDEPK | 3.21737 | 4.94979 | 0 | 205 |
| **TNFR: Fc1** | SQHTQPTPEPSTAPSTSFLLPMGPSPPAEGSTGDEPK | 6.70649 | 10.31767 | 0 | 205 |
| **TNFR: Fc1** | SQHTQPTPEPSTAPSTSFLLPMGPSPPAEGSTGDEPK | 5.01763 | 5.75264 | 3.65261 | 202 |
| **TNFR: Fc1** | SQHTQPTPEPSTAPSTSFLLPMGPSPPAEGSTGDEPK | 3.13563 | 4.82405 | 0 | 202 |
| **TNFR: Fc1** | SQHTQPTPEPSTAPSTSFLLPMGPSPPAEGSTGDEPK | 3.11557 | 4.79319 | 0 | 202 |
| **TNFR: Fc1** | SQHTQPTPEPSTAPSTSFLLPMGPSPPAEGSTGDEPK | 4.99466 | 5.96327 | 3.19582 | 202 |
| **TNFR: Fc2** | SQHTQPT | 3.14197 | 4.8338 | 0 | 205 |
| **TNFR: Fc2** | AVHLPQPVSTR | 27.83603 | 38.16024 | 8.66248 | 199 |
| **TNFR: Fc2** | SMAPGAVHLPQPVSTR | 44.39855 | 65.29959 | 5.58232 | 199 |
| **TNFR: Fc2** | SMAPGAVHLPQPVSTR | 50.20209 | 74.11999 | 5.78313 | 199 |
| **TNFR: Fc2** | SMAPGAVHLPQPVSTR | 32.11355 | 49.40546 | 0 | 199 |
| **TNFR: Fc2** | SMAPGAVHLPQPVSTR | 50.19945 | 74.02912 | 5.94436 | 199 |
| **TNFR: Fc2** | SMAPGAVHLPQPVSTR | 13.05 | 20.07693 | 0 | 199 |
| **TNFR: Fc2** | SMAPGAVHLPQPVSTR | 41.19513 | 60.43458 | 5.46471 | 199 |
| **TNFR: Fc2** | SMAPGAVHLPQPVSTR | 48.96745 | 72.32979 | 5.58025 | 199 |
| **TNFR: Fc2** | SMAPGAVHLPQPVSTR | 46.9573 | 69.21726 | 5.61737 | 199 |
| **TNFR: Fc2** | SMAPGAVHLPQPVSTR | 22.14077 | 31.85108 | 4.10733 | 199 |
| **TNFR: Fc2** | SMAPGAVHLPQPVSTR | 29.21545 | 39.73523 | 9.67871 | 199 |
| **TNFR: Fc2** | SMAPGAVHLPQPVSTR | 36.46959 | 50.61252 | 10.20414 | 199 |
| **TNFR: Fc2** | SMAPGAVHLPQPVSTR | 42.58047 | 62.79197 | 5.04482 | 200 |
| **TNFR: Fc2** | SMAPGAVHLPQPVSTR | 48.98215 | 72.81844 | 4.71475 | 200 |
| **TNFR: Fc2** | SMAPGAVHLPQPVSTR | 44.45233 | 65.46439 | 5.42993 | 199 |
| **TNFR: Fc2** | SMAPGAVHLPQPVSTR | 47.61701 | 70.40701 | 5.29273 | 199 |
| **TNFR: Fc2** | SMAPGAVHLPQPVSTR | 48.92826 | 72.24626 | 5.62341 | 199 |
| **TNFR: Fc2** | SMAPGAVHLPQPVSTR | 32.24813 | 46.99222 | 4.86625 | 199 |
| **TNFR: Fc2** | SMAPGAVHLPQPVSTR | 40.16352 | 58.82112 | 5.5137 | 199 |
| **TNFR: Fc2** | SMAPGAVHLPQPVSTR | 43.10916 | 63.33092 | 5.55447 | 199 |
| **TNFR: Fc2** | APGAVHLPQPVSTR | 47.13301 | 69.64732 | 5.32073 | 199 |
| **TNFR: Fc2** | SMAPGAVHLPQPVSTR | 9.7098 | 14.93816 | 0 | 199 |
| **TNFR: Fc2** | SMAPGAVHLPQPVSTR | 23.88871 | 36.75186 | 0 | 199 |
| **TNFR: Fc2** | REPQVYTLPPSR | 5.04287 | 7.12193 | 1.18176 | 374 |
| **TNFR: Fc2** | SMAPGAVHLPQPVSTR | 55.26383 | 81.82617 | 5.93377 | 199 |
| **TNFR: Fc2** | SMAPGAVHLPQPVSTR | 19.82794 | 30.50453 | 0 | 199 |
| **TNFR: Fc2** | SMAPGAVHLPQPVSTR | 55.68016 | 82.48075 | 5.90764 | 199 |
| **TNFR: Fc2** | SMAPGAVHLPQPVSTR | 18.27048 | 23.17219 | 9.1673 | 199 |
| **TNFR: Fc2** | SMAPGAVHLPQPVSTR | 17.30702 | 26.62619 | 0 | 199 |
| **TNFR: Fc2** | SMAPGAVHLPQPVSTR | 50.66966 | 74.97606 | 5.5292 | 199 |
| **TNFR: Fc2** | SMAPGAVHLPQPVSTR | 46.71048 | 68.89991 | 5.50155 | 199 |
| **TNFR: Fc2** | SMAPGAVHLPQPVSTR | 52.09955 | 77.43459 | 5.04876 | 199 |

| Sample | Peptide | TotalScore | PepScore | GlyScore | ProSites |
|---|---|---|---|---|---|
| **TNFR: Fc2** | SMAPGAVHLPQPVSTR | 31.45863 | 45.74288 | 4.93074 | 199 |
| **TNFR: Fc2** | SMAPGAVHLPQPVSTR | 21.82518 | 30.87798 | 5.01286 | 199 |
| **TNFR: Fc2** | SQHTQPTPEPSTAPSTSFLLPMGPSPPAEGSTGDEPK | 17.43821 | 21.87181 | 9.20438 | 226 |
| **TNFR: Fc2** | SQHTQPTPEPSTAPSTSFLLPMGPSPPAEGSTGD | 11.58968 | 17.83027 | 0 | 226 |
| **TNFR: Fc2** | SQHTQPTPEPSTAPSTSFLLPMGPSPPAEGSTGD | 11.94233 | 18.37282 | 0 | 226 |
| **TNFR: Fc2** | SQHTQPTPEPSTAPSTSFLLPMGPSPPAEGSTGDEPK | 22.1082 | 33.33337 | 1.26145 | 208 |
| **TNFR: Fc2** | SQHTQPTPEPSTAPSTSFLLPMGPSPPAEGSTGDEPK | 75.58169 | 113.176 | 5.7637 | 217 |
| **TNFR: Fc2** | SQHTQPTPEPSTAPSTSFLLPMGPSPPAEGSTGDEPK | 74.74312 | 112.3035 | 4.9882 | 217 |
| **TNFR: Fc2** | SQHTQPTPEPSTAPSTSFLLPMGPSPPAEGSTGDEPK | 77.16796 | 113.6519 | 9.41213 | 217 |
| **TNFR: Fc2** | SQHTQPTPEPSTAPSTSFLLPMGPSPPAEGSTGDEPK | 68.36572 | 102.0594 | 5.79171 | 213 |
| **TNFR: Fc2** | SQHTQPTPEPSTAPSTSFLLPMGPSPPAEGSTGDEPK | 62.32533 | 91.0361 | 9.00533 | 213 |
| **TNFR: Fc2** | SQHTQPTPEPSTAPSTSFLLPMGPSPPAEGSTGDEPK | 58.78668 | 87.20107 | 6.01711 | 213 |
| **TNFR: Fc2** | SQHTQPTPEPSTAPSTSFLLPMGPSPPAEGSTGDEPK | 69.65601 | 104.4236 | 5.08773 | 213 |
| **TNFR: Fc2** | SQHTQPTPEPSTAPSTSFLLPMGPSPPAEGSTGDEPK | 9.74104 | 14.98622 | 0 | 208 |
| **TNFR: Fc2** | SQHTQPTPEPSTAPSTSFLLPMGPSPPAEGSTGDEPK | 73.58455 | 109.939 | 6.06917 | 205 |
| **TNFR: Fc2** | SQHTQPTPEPSTAPSTSFLLPMGPSPPAEGSTGDEPK | 88.58362 | 133.0446 | 6.01331 | 205 |
| **TNFR: Fc2** | SQHTQPTPEPSTAPSTSFLLPMGPSPPAEGSTGDEPK | 71.13382 | 106.5186 | 5.41924 | 205 |
| **TNFR: Fc2** | SQHTQPTPEPSTAPSTSFLLPMGPSPPAEGSTGDEPK | 71.63277 | 107.5219 | 4.98147 | 205 |
| **TNFR: Fc2** | SQHTQPTPEPSTAPSTSFLLPMGPSPPAEGSTGDEPK | 47.30663 | 72.77943 | 0 | 213 |
| **TNFR: Fc2** | SQHTQPTPEPSTAPSTSFLLPMGPSPPAEGSTGDEPK | 85.22686 | 127.9411 | 5.90047 | 205 |
| **TNFR: Fc2** | SQHTQPTPEPSTAPSTSFLLPMGPSPPAEGSTGDEPK | 49.83369 | 74.66022 | 3.72729 | 205 |
| **TNFR: Fc2** | SQHTQPTPEPSTAPSTSFLLPMGPSPPAEGSTGDEPK | 74.89182 | 112.0503 | 5.88329 | 213 |
| **TNFR: Fc2** | SQHTQPTPEPSTAPSTSFLLPMGPSPPAEGSTGDEPK | 56.77927 | 87.35272 | 0 | 213 |
| **TNFR: Fc2** | SQHTQPTPEPSTAPSTSFLLPMGPSPPAEGSTGDEPK | 55.62862 | 81.01975 | 8.47366 | 213 |
| **TNFR: Fc2** | SQHTQPTPEPSTAPSTSFLLPMGPSPPAEGSTGDEPK | 67.84098 | 101.2326 | 5.82803 | 213 |
| **TNFR: Fc2** | SQHTQPTPEPSTAPSTSFLLPMGPSPPAEGSTGDEPK | 53.12901 | 78.62852 | 5.77279 | 213 |
| **TNFR: Fc2** | SQHTQPTPEPSTAPSTSFLLPMGPSPPAEGSTGDEPK | 58.20919 | 85.11332 | 8.24438 | 205 |
| **TNFR: Fc2** | SQHTQPTPEPSTAPSTSFLLPMGPSPPAEGSTGDEPK | 58.87112 | 87.84067 | 5.07053 | 205 |
| **TNFR: Fc2** | SQHTQPTPEPSTAPSTSFLLPMGPSPPAEGSTGDEPK | 43.31709 | 62.20698 | 8.23588 | 205 |
| **TNFR: Fc2** | SQHTQPTPEPSTAPSTSFLLPMGPSPPAEGSTGDEPK | 41.26263 | 61.18423 | 4.26539 | 205 |
| **TNFR: Fc2** | SQHTQPTPEPSTAPSTSFLLPMGPSPPAEGSTGDEPK | 54.12607 | 83.27088 | 0 | 205 |
| **TNFR: Fc2** | SQHTQPTPEPSTAPSTSFLLPMGPSPPAEGSTGDEPK | 64.9055 | 96.8053 | 5.66303 | 217 |
| **TNFR: Fc2** | SQHTQPTPEPSTAPSTSFLLPMGPSPPAEGSTGDEPK | 43.12486 | 64.46173 | 3.49926 | 205 |
| **TNFR: Fc2** | SQHTQPTPEPSTAPSTSFLLPMGPSPPAEGSTGDEPK | 29.64684 | 43.09374 | 4.67402 | 216 |
| **TNFR: Fc2** | SQHTQPTPEPSTAPSTSFLLPMGPSPPAEGSTGDEPK | 39.05259 | 57.50383 | 4.78599 | 205 |
| **TNFR: Fc2** | SQHTQPTPEPSTAPSTSFLLPMGPSPPAEGSTGDEPK | 16.92473 | 26.03805 | 0 | 202 |
| **TNFR: Fc2** | SQHTQPTPEPSTAPSTSFLLPMGPSPPAEGSTGDEPK | 46.83981 | 69.50177 | 4.75331 | 217 |
| **TNFR: Fc2** | SQHTQPTPEPSTAPSTSFLLPMGPSPPAEGSTGDEPK | 94.64013 | 138.3444 | 13.47502 | 226 |
| **TNFR: Fc2** | SQHTQPTPEPSTAPSTSFLLPMGPSPPAEGSTGDEPK | 105.9168 | 155.5121 | 13.81109 | 226 |
| **TNFR: Fc2** | SQHTQPTPEPSTAPSTSFLLPMGPSPPAEGSTGDEPK | 77.91777 | 116.8017 | 5.70483 | 216 |
| **TNFR: Fc2** | SQHTQPTPEPSTAPSTSFLLPMGPSPPAEGSTGDEPK | 81.52952 | 122.3829 | 5.65901 | 216 |
| **TNFR: Fc2** | SQHTQPTPEPSTAPSTSFLLPMGPSPPAEGSTGDEPK | 70.62282 | 108.6505 | 0 | 217 |

| Sample | Peptide | TotalScore | PepScore | GlyScore | ProSites |
|---|---|---|---|---|---|
| **TNFR: Fc2** | SQHTQPTPEPSTAPSTSFLLPMGPSPPAEGSTGDEPK | 71.35187 | 109.7721 | 0 | 213 |
| **TNFR: Fc2** | SQHTQPTPEPSTAPSTSFLLPMGPSPPAEGSTGDEPK | 105.3714 | 159.3143 | 5.19169 | 213 |
| **TNFR: Fc2** | SQHTQPTPEPSTAPSTSFLLPMGPSPPAEGSTGDEPK | 52.41907 | 76.44408 | 7.8012 | 205 |
| **TNFR: Fc2** | HTQPTPEPSTAPSTSFLLPMGPSPPAEGSTGDEPK | 35.60828 | 54.09906 | 1.26826 | 226 |
| **TNFR: Fc2** | SQHTQPTPEPSTAPSTSFLLPMGPSPPAEGSTGDEPK | 85.6728 | 128.6475 | 5.86266 | 213 |
| **TNFR: Fc2** | HTQPTPEPSTAPSTSFLLPMGPSPPAEGSTGDEPK | 30.41511 | 46.13149 | 1.22754 | 226 |
| **TNFR: Fc2** | SQHTQPTPEPSTAPSTSFLLPMGPSPPAEGSTGDEPK | 4.42926 | 6.81424 | 0 | 205 |
| **TNFR: Fc2** | HTQPTPEPSTAPSTSFLLPMGPSPPAEGSTGDEPK | 23.31897 | 35.76413 | 0.20655 | 212 |
| **TNFR: Fc2** | SQHTQPTPEPSTAPSTSFLLPMGPSPPAEGSTGDEPK | 8.32968 | 12.10105 | 1.32572 | 226 |
| **TNFR: Fc2** | SQHTQPTPEPSTAPSTSFLLPMGPSPPAEGSTGDEPK | 66.32072 | 99.47057 | 4.75671 | 205 |
| **TNFR: Fc2** | SQHTQPTPEPSTAPSTSFLLPMGPSPPAEGSTGDEPK | 5.9607 | 9.17031 | 0 | 208 |
| **TNFR: Fc2** | SQHTQPTPEPSTAPSTSFLLPMGPSPPAEGSTGDEPK | 133.5081 | 201.5628 | 7.1207 | 205 |
| **TNFR: Fc2** | SQHTQPTPEPSTAPSTSFLLPMGPSPPAEGSTGDEPK | 68.10666 | 102.2072 | 4.77716 | 205 |
| **TNFR: Fc2** | SQHTQPTPEPSTAPSTSFLLPMGPSPPAEGSTGDEPK | 86.7752 | 129.8532 | 6.77325 | 205 |
| **TNFR: Fc2** | SQHTQPTPEPSTAPSTSFLLPMGPSPPAEGSTGDEPK | 95.46656 | 143.2373 | 6.74942 | 205 |
| **TNFR: Fc2** | SQHTQPTPEPSTAPSTSFLLPMGPSPPAEGSTGDEPK | 87.18504 | 130.5662 | 6.62006 | 205 |
| **TNFR: Fc2** | SQHTQPTPEPSTAPSTSFLLPMGPSPPAEGSTGDEPK | 27.66099 | 42.55538 | 0 | 205 |
| **TNFR: Fc2** | SQHTQPTPEPSTAPSTSFLLPMGPSPPAEGSTGDEPK | 65.30588 | 97.50712 | 5.50357 | 208 |
| **TNFR: Fc2** | SQHTQPTPEPSTAPSTSFLLPMGPSPPAEGSTGDEPK | 68.57085 | 103.0063 | 4.61927 | 205 |
| **TNFR: Fc2** | SQHTQPTPEPSTAPSTSFLLPMGPSPPAEGSTGDEPK | 71.22726 | 106.7171 | 5.31766 | 205 |
| **TNFR: Fc2** | SQHTQPTPEPSTAPSTSFLLPMGPSPPAEGSTGDEPK | 43.80562 | 64.82139 | 4.77634 | 205 |
| **TNFR: Fc2** | SQHTQPTPEPSTAPSTSFLLPMGPSPPAEGSTGDEPK | 73.98272 | 111.3578 | 4.57195 | 205 |
| **TNFR: Fc2** | SQHTQPTPEPSTAPSTSFLLPMGPSPPAEGSTGDEPK | 65.26385 | 97.83873 | 4.76766 | 205 |
| **TNFR: Fc2** | SQHTQPTPEPSTAPSTSFLLPMGPSPPAEGSTGDEPK | 34.74304 | 51.13815 | 4.29497 | 205 |
| **TNFR: Fc2** | SQHTQPTPEPSTAPSTSFLLPMGPSPPAEGSTGDEPK | 27.29495 | 40.32656 | 3.09341 | 205 |
| **TNFR: Fc2** | SQHTQPTPEPSTAPSTSFLLPMGPSPPAEGSTGD | 17.59101 | 26.43987 | 1.15741 | 226 |
| **TNFR: Fc2** | TQPTPEPSTAPSTSFLLPMGPSPPAEGSTGDEPK | 14.02588 | 21.57828 | 0 | 205 |
| **TNFR: Fc2** | SQHTQPTPEPSTAPSTSFLLPMGPSPPAEGSTGDEP | 10.8227 | 16.65031 | 0 | 226 |
| **TNFR: Fc2** | SQHTQPTPEPSTAPSTSFLLPMG | 14.59621 | 22.45571 | 0 | 205 |
| **TNFR: Fc2** | SQHTQPTPEPSTAPSTSFLLPMG | 8.1899 | 12.59985 | 0 | 205 |
| **TNFR: Fc2** | SQHTQPTPEPSTAPSTSFLLPMGPSPPAEGSTGDEP | 5.68922 | 8.75264 | 0 | 216 |
| **TNFR: Fc2** | SQHTQPTPEPSTAPSTSFLLPMGPSPPAEGSTGD | 18.88526 | 26.44106 | 4.85306 | 202 |
| **TNFR: Fc2** | SQHTQPTPEPSTAPSTSFLLPM | 15.81508 | 24.33089 | 0 | 205 |
| **TNFR: Fc2** | SQHTQPTPEPSTAPSTSFLLPM | 16.62994 | 25.58453 | 0 | 205 |
| **TNFR: Fc2** | HTQPTPEPSTAPSTSFLLPMGPSPPAEGSTGDEPK | 3.00375 | 4.62115 | 0 | 205 |
| **TNFR: Fc2** | SQHTQPTPEPSTAPSTSFLLPMGPSPPAE | 6.30814 | 9.70483 | 0 | 226 |
| **TNFR: Fc2** | SQHTQPTPEPSTAPSTSFLLPMGPSPPAEGSTGD | 32.2019 | 43.8897 | 10.496 | 217 |
| **TNFR: Fc2** | SQHTQPTPEPSTAPSTSFLLPMGPSPPAEGSTGD | 32.60153 | 44.24754 | 10.97325 | 226 |
| **TNFR: Fc2** | SQHTQPTPEPSTAPSTSFLLPMGPSPPAEGSTGD | 35.36054 | 53.64652 | 1.40088 | 226 |
| **TNFR: Fc2** | SQHTQPTPEPSTAPSTSFLLPMGPSPPAEG | 55.3968 | 82.4171 | 5.21625 | 205 |
| **TNFR: Fc2** | SQHTQPTPEPSTAPSTSFLLPMGPSPPAEGSTGD | 14.79754 | 20.32461 | 4.53299 | 216 |
| **TNFR: Fc2** | SQHTQPTPEPSTAPSTSFLLPMGPSPPAEGSTGD | 9.45796 | 12.19528 | 4.37437 | 212 |

| Sample | Peptide | TotalScore | PepScore | GlyScore | ProSites |
|---|---|---|---|---|---|
| **TNFR: Fc2** | SQHTQPTPEPSTAPSTSFLLPMGPSPPAEGSTGDE | 6.90075 | 8.35189 | 4.20576 | 202 |
| **TNFR: Fc2** | SQHTQPTPEPSTAPSTSFLLPMGPSPPAEGSTGD | 45.61801 | 67.20275 | 5.53206 | 202 |
| **TNFR: Fc2** | SQHTQPTPEPSTAPSTSFLLPMGPSPPAEGSTGD | 2.48145 | 3.81761 | 0 | 202 |
| **TNFR: Fc2** | SMAPGAVHLPQPVSTR | 14.89419 | 21.56631 | 2.50311 | 199 |
| **TNFR: Fc2** | SMAPGAVHLPQPVSTR | 11.77298 | 18.11227 | 0 | 199 |
| **TNFR: Fc2** | SMAPGAVHLPQPVSTR | 5.80266 | 8.92717 | 0 | 199 |
| **TNFR: Fc2** | APGAVHLPQPVSTR | 44.13367 | 64.97962 | 5.41975 | 199 |
| **TNFR: Fc2** | SMAPGAVHLPQPVSTR | 53.61307 | 79.34953 | 5.81679 | 199 |
| **TNFR: Fc2** | SMAPGAVHLPQPVSTR | 49.08016 | 72.3852 | 5.79937 | 199 |
| **TNFR: Fc2** | SMAPGAVHLPQPVSTR | 51.18857 | 75.51242 | 6.01571 | 199 |
| **TNFR: Fc2** | SMAPGAVHLPQPVSTR | 25.0613 | 32.58321 | 11.09206 | 199 |
| **TNFR: Fc2** | SMAPGAVHLPQPVSTR | 23.47075 | 29.86928 | 11.58778 | 199 |
| **TNFR: Fc2** | SMAPGAVHLPQPVSTR | 44.48871 | 65.39796 | 5.65725 | 199 |
| **TNFR: Fc2** | SMAPGAVHLPQPVSTR | 41.90447 | 61.46714 | 5.5738 | 199 |
| **TNFR: Fc2** | SMAPGAVHLPQPVSTR | 51.97797 | 76.91273 | 5.67055 | 199 |
| **TNFR: Fc2** | SMAPGAVHLPQPVSTR | 32.57788 | 45.27295 | 9.00133 | 199 |
| **TNFR: Fc2** | MAPGAVHLPQPVSTR | 3.68286 | 5.66594 | 0 | 200 |
| **TNFR: Fc2** | MAPGAVHLPQPVSTR | 2.667 | 4.10308 | 0 | 200 |
| **TNFR: Fc2** | SMAPGAVHLPQPVSTR | 37.73985 | 55.32401 | 5.08355 | 200 |
| **TNFR: Fc2** | SMAPGAVHLPQPVSTR | 44.81009 | 65.99936 | 5.45858 | 199 |
| **TNFR: Fc2** | SMAPGAVHLPQPVSTR | 45.45802 | 67.00372 | 5.44457 | 199 |
| **TNFR: Fc2** | SMAPGAVHLPQPVSTR | 52.19809 | 77.47676 | 5.25198 | 200 |
| **TNFR: Fc2** | SMAPGAVHLPQPVSTR | 45.59194 | 67.12087 | 5.60963 | 199 |
| **TNFR: Fc2** | SMAPGAVHLPQPVSTR | 58.33545 | 86.82553 | 5.4253 | 199 |
| **TNFR: Fc2** | SMAPGAVHLPQPVSTR | 40.89659 | 60.04592 | 5.33353 | 199 |
| **TNFR: Fc2** | SMAPGAVHLPQPVSTR | 42.71632 | 62.82278 | 5.37574 | 199 |
| **TNFR: Fc2** | SMAPGAVHLPQPVSTR | 40.85591 | 59.87337 | 5.53777 | 199 |
| **TNFR: Fc2** | PGAVHLPQPVSTR | 41.49019 | 58.01743 | 10.79674 | 199 |
| **TNFR: Fc2** | PGAVHLPQPVSTR | 40.46392 | 56.48322 | 10.71379 | 199 |
| **TNFR: Fc2** | APGAVHLPQPVSTR | 44.11477 | 64.91396 | 5.48769 | 199 |
| **TNFR: Fc2** | SMAPGAVHLPQPVSTR | 24.41813 | 34.23559 | 6.18569 | 199 |
| **TNFR: Fc2** | APGAVHLPQPVSTR | 58.34569 | 86.55753 | 5.95226 | 199 |
| **TNFR: Fc2** | SMAPGAVHLPQPVSTR | 18.65313 | 28.69712 | 0 | 200 |
| **TNFR: Fc2** | REPQVYTLPPSR | 4.85934 | 6.82572 | 1.2075 | 370 |
| **TNFR: Fc2** | SMAPGAVHLPQPVSTR | 63.70129 | 94.84371 | 5.86538 | 199 |
| **TNFR: Fc2** | REPQVYTLPPSR | 4.5092 | 6.93723 | 0 | 374 |
| **TNFR: Fc2** | SMAPGAVHLPQPVSTR | 61.63873 | 91.62408 | 5.95166 | 199 |
| **TNFR: Fc2** | REPQVYTLPPSR | 3.81487 | 5.86904 | 0 | 370 |
| **TNFR: Fc2** | SMAPGAVHLPQPVSTR | 3.74321 | 5.75879 | 0 | 199 |
| **TNFR: Fc2** | SMAPGAVHLPQPVSTR | 33.86373 | 52.09805 | 0 | 199 |
| **TNFR: Fc2** | SMAPGAVHLPQPVSTR | 51.10981 | 75.6578 | 5.52067 | 199 |
| **TNFR: Fc2** | SMAPGAVHLPQPVSTR | 52.38226 | 77.66879 | 5.42156 | 199 |

| Sample | Peptide | TotalScore | PepScore | GlyScore | ProSites |
|---|---|---|---|---|---|
| **TNFR: Fc2** | SMAPGAVHLPQPVSTR | 54.09042 | 80.29067 | 5.43282 | 199 |
| **TNFR: Fc2** | SMAPGAVHLPQPVSTR | 11.23285 | 17.28131 | 0 | 199 |
| **TNFR: Fc2** | SMAPGAVHLPQPVSTR | 48.58357 | 71.99139 | 5.11192 | 199 |
| **TNFR: Fc2** | SMAPGAVHLPQPVSTR | 55.82168 | 83.13561 | 5.09579 | 199 |
| **TNFR: Fc2** | SMAPGAVHLPQPVSTR | 44.75899 | 66.18344 | 4.97072 | 199 |
| **TNFR: Fc2** | SMAPGAVHLPQPVSTR | 43.7279 | 64.81744 | 4.56162 | 199 |
| **TNFR: Fc2** | SMAPGAVHLPQPVSTR | 17.78966 | 24.78724 | 4.79415 | 199 |
| **TNFR: Fc2** | SMAPGAVHLPQPVSTR | 22.67674 | 32.49159 | 4.44915 | 199 |
| **TNFR: Fc2** | SQHTQPTPEPSTAPSTSFLLPMGPSPPAEGSTGDEPK | 7.50971 | 11.55341 | 0 | 226 |
| **TNFR: Fc2** | REPQVYTLPPSR | 2.87032 | 4.41587 | 0 | 374 |
| **TNFR: Fc2** | SMAPGAVHLPQPVSTR | 24.9531 | 35.99482 | 4.44705 | 199 |
| **TNFR: Fc2** | SMAPGAVHLPQPVSTR | 21.75818 | 31.05445 | 4.49368 | 199 |
| **TNFR: Fc2** | SQHTQPTPEPSTAPSTSFLLPMGPSPPAEGSTGD | 7.86417 | 12.09872 | 0 | 216 |
| **TNFR: Fc2** | SMAPGAVHLPQPVSTR | 21.14223 | 32.52651 | 0 | 199 |
| **TNFR: Fc2** | SQHTQPTPEPSTAPSTSFLLPMGPSPPAEGSTGDEPK | 17.75253 | 22.53399 | 8.87267 | 217 |
| **TNFR: Fc2** | SQHTQPTPEPSTAPSTSFLLPMGPSPPAEGSTGDEPK | 83.8077 | 125.7113 | 5.98667 | 217 |
| **TNFR: Fc2** | SQHTQPTPEPSTAPSTSFLLPMGPSPPAEGSTGDEPK | 66.93293 | 98.00644 | 9.22498 | 217 |
| **TNFR: Fc2** | SQHTQPTPEPSTAPSTSFLL | 20.02463 | 30.80712 | 0 | 205 |
| **TNFR: Fc2** | SQHTQPTPEPSTAPSTSFLLPMGPSPPAEGSTGDEPK | 31.5216 | 42.90685 | 10.37758 | 217 |
| **TNFR: Fc2** | SQHTQPTPEPSTAPSTSFLLPMGPSPPAEGSTGDEPK | 62.46235 | 92.75697 | 6.20093 | 213 |
| **TNFR: Fc2** | SQHTQPTPEPSTAPSTSFLLPMGPSPPAEGSTGDEPK | 61.58115 | 91.96854 | 5.14744 | 213 |
| **TNFR: Fc2** | SQHTQPTPEPSTAPSTSFLLPMGPSPPAEGSTGDEPK | 28.74953 | 42.21332 | 3.74535 | 205 |
| **TNFR: Fc2** | SQHTQPTPEPSTAPSTSFLLPMGPSPPAEGSTGDEPK | 80.80505 | 120.9523 | 6.24584 | 213 |
| **TNFR: Fc2** | SQHTQPTPEPSTAPSTSFLLPMGPSPPAEGSTGDEPK | 77.4596 | 115.815 | 6.22814 | 205 |
| **TNFR: Fc2** | SQHTQPTPEPSTAPSTSFLLPMGPSPPAEGSTGDEPK | 68.84682 | 100.5116 | 10.04074 | 205 |
| **TNFR: Fc2** | SQHTQPTPEPSTAPSTSFLLPMGPSPPAEGSTGDEPK | 64.53338 | 96.46109 | 5.23906 | 205 |
| **TNFR: Fc2** | SQHTQPTPEPSTAPSTSFLLPMGPSPPAEGSTGDEPK | 70.0422 | 104.8689 | 5.36405 | 205 |
| **TNFR: Fc2** | SQHTQPTPEPSTAPSTSFLLPMGPSPPAEGSTGDEPK | 59.54771 | 88.70048 | 5.40685 | 205 |
| **TNFR: Fc2** | SQHTQPTPEPSTAPSTSFLLPMGPSPPAEGSTGDEPK | 54.65945 | 81.2803 | 5.22073 | 205 |
| **TNFR: Fc2** | SQHTQPTPEPSTAPSTSFLLPMGPSPPAEGSTGDEPK | 63.41112 | 94.29 | 6.06462 | 205 |
| **TNFR: Fc2** | SQHTQPTPEPSTAPSTSFLLPMGPSPPAEGSTGDEPK | 76.83366 | 112.2913 | 10.98377 | 205 |
| **TNFR: Fc2** | SQHTQPTPEPSTAPSTSFLLPMGPSPPAEGSTGDEPK | 14.49999 | 17.63539 | 8.67709 | 205 |
| **TNFR: Fc2** | SQHTQPTPEPSTAPSTSFLLPMGPSPPAEGSTGDEPK | 56.5462 | 82.009 | 9.25813 | 213 |
| **TNFR: Fc2** | SQHTQPTPEPSTAPSTSFLLPMGPSPPAEGSTGDEPK | 55.12084 | 82.25091 | 4.73641 | 213 |
| **TNFR: Fc2** | SQHTQPTPEPSTAPSTSFLLPMGPSPPAEGSTGDEPK | 59.57291 | 88.64063 | 5.58999 | 213 |
| **TNFR: Fc2** | SQHTQPTPEPSTAPSTSFLLPMGPSPPAEGSTGDEPK | 53.25852 | 78.95959 | 5.52794 | 213 |
| **TNFR: Fc2** | SQHTQPTPEPSTAPSTSFLLPMGPSPPAEGSTGDEPK | 36.14808 | 53.31517 | 4.26633 | 213 |
| **TNFR: Fc2** | SQHTQPTPEPSTAPSTSFLLPMGPSPPAEGSTGDEPK | 55.62093 | 82.47172 | 5.75517 | 217 |
| **TNFR: Fc2** | SQHTQPTPEPSTAPSTSFLLPMGPSPPAEGSTGDEPK | 60.21148 | 90.06397 | 4.77114 | 205 |
| **TNFR: Fc2** | SQHTQPTPEPSTAPSTSFLLPMGPSPPAEGSTGDEPK | 62.59916 | 93.25157 | 5.67326 | 205 |
| **TNFR: Fc2** | QPTPEPSTAPSTSFLLPMGPSPPAEGSTGDEPK | 19.7235 | 30.34385 | 0 | 208 |
| **TNFR: Fc2** | SQHTQPTPEPSTAPSTSFLLPMGPSPPAEGSTGDEPK | 49.74019 | 73.90084 | 4.87042 | 213 |

| Sample | Peptide | TotalScore | PepScore | GlyScore | ProSites |
|---|---|---|---|---|---|
| **TNFR: Fc2** | SQHTQPTPEPSTAPSTSFLLPMGPSPPAEGSTGDEPK | 60.46078 | 88.68738 | 8.03995 | 213 |
| **TNFR: Fc2** | SQHTQPTPEPSTAPSTSFLLPMGPSPPAEGSTGDEPK | 55.05677 | 80.64705 | 7.53198 | 213 |
| **TNFR: Fc2** | SQHTQPTPEPSTAPSTSFLLPMGPSPPAEGSTGDEPK | 56.05243 | 83.33243 | 5.38956 | 213 |
| **TNFR: Fc2** | SQHTQPTPEPSTAPSTSFLLPMGPSPPAEGSTGDEPK | 9.1269 | 14.04139 | 0 | 213 |
| **TNFR: Fc2** | SQHTQPTPEPSTAPSTSFLLPMGPSPPAEGSTGDEPK | 25.12637 | 36.16556 | 4.625 | 205 |
| **TNFR: Fc2** | SQHTQPTPEPSTAPSTSFLLPMGPSPPAEGSTGDEPK | 38.99043 | 55.70837 | 7.94284 | 213 |
| **TNFR: Fc2** | PTPEPSTAPSTSFLLPMGPSPPAEGSTGDEPK | 6.93578 | 10.67044 | 0 | 208 |
| **TNFR: Fc2** | SQHTQPTPEPSTAPSTSFLLPMGPSPPAEGSTGDEPK | 55.39048 | 83.23345 | 3.68211 | 216 |
| **TNFR: Fc2** | SQHTQPTPEPSTAPSTSFLLPMGPSPPAEGSTGDEPK | 52.17116 | 80.26333 | 0 | 217 |
| **TNFR: Fc2** | SQHTQPTPEPSTAPSTSFLLPMGPSPPAEGSTGDEPK | 79.57177 | 118.9932 | 6.36061 | 216 |
| **TNFR: Fc2** | SQHTQPTPEPSTAPSTSFLLPMGPSPPAEGSTGDEPK | 53.38614 | 82.13253 | 0 | 217 |
| **TNFR: Fc2** | SQHTQPTPEPSTAPSTSFLLPMGPSPPAEGSTGDEPK | 111.0109 | 167.2965 | 6.48058 | 213 |
| **TNFR: Fc2** | SQHTQPTPEPSTAPSTSFLLPMGPSPPAEGSTGDEPK | 44.85417 | 67.34885 | 3.07832 | 205 |
| **TNFR: Fc2** | HTQPTPEPSTAPSTSFLLPMGPSPPAEGSTGDEPK | 31.97818 | 48.536 | 1.22794 | 226 |
| **TNFR: Fc2** | HTQPTPEPSTAPSTSFLLPMGPSPPAEGSTGDEPK | 27.36727 | 41.46259 | 1.19025 | 226 |
| **TNFR: Fc2** | SQHTQPTPEPSTAPSTSFLLPMGPSPPAEGSTGDEPK | 60.51587 | 90.44749 | 4.92858 | 213 |
| **TNFR: Fc2** | SQHTQPTPEPSTAPSTSFLLPMGPSPPAEGSTGDEPK | 53.83929 | 80.32093 | 4.65909 | 205 |
| **TNFR: Fc2** | SQHTQPTPEPSTAPSTSFLLPMGPSPPAEGSTGDEPK | 7.69625 | 11.84039 | 0 | 208 |
| **TNFR: Fc2** | SQHTQPTPEPSTAPSTSFLLPMGPSPPAEGSTGDEPK | 103.8415 | 159.7562 | 0 | 205 |
| **TNFR: Fc2** | SQHTQPTPEPSTAPSTSFLLPMGPSPPAEGSTGDEPK | 141.7747 | 214.2477 | 7.18204 | 205 |
| **TNFR: Fc2** | SQHTQPTPEPSTAPSTSFLLPMGPSPPAEGSTGDEPK | 89.96191 | 134.7608 | 6.76406 | 205 |
| **TNFR: Fc2** | SQHTQPTPEPSTAPSTSFLLPMGPSPPAEGSTGDEPK | 82.76916 | 123.6813 | 6.78948 | 205 |
| **TNFR: Fc2** | SQHTQPTPEPSTAPSTSFLLPMGPSPPAEGSTGDEPK | 70.30973 | 104.5605 | 6.70111 | 205 |
| **TNFR: Fc2** | SQHTQPTPEPSTAPSTSFLLPMGPSPPAEGSTGDEPK | 84.53319 | 126.5316 | 6.53619 | 205 |
| **TNFR: Fc2** | SQHTQPTPEPSTAPSTSFLLPMGPSPPAEGSTGDEPK | 89.71922 | 134.5027 | 6.54991 | 205 |
| **TNFR: Fc2** | SQHTQPTPEPSTAPSTSFLLPMGPSPPAEGSTGDEPK | 66.1536 | 101.7748 | 0 | 205 |
| **TNFR: Fc2** | SQHTQPTPEPSTAPSTSFLLPMGPSPPAEGSTGDEPK | 72.47793 | 108.4479 | 5.67662 | 213 |
| **TNFR: Fc2** | SQHTQPTPEPSTAPSTSFLLPMGPSPPAEGSTGDEPK | 80.23093 | 121.0087 | 4.50082 | 205 |
| **TNFR: Fc2** | SQHTQPTPEPSTAPSTSFLLPMGPSPPAEGSTGDEPK | 22.87606 | 32.63492 | 4.75247 | 205 |
| **TNFR: Fc2** | SQHTQPTPEPSTAPSTSFLLPMGPSPPAEGSTGDEPK | 57.64229 | 86.2484 | 4.51666 | 208 |
| **TNFR: Fc2** | HTQPTPEPSTAPSTSFLLPMGPSPPAEGSTGDEPK | 10.84712 | 16.68787 | 0 | 212 |
| **TNFR: Fc2** | HTQPTPEPSTAPSTSFLLPMGPSPPAEGSTGDEPK | 28.87427 | 42.35765 | 3.83372 | 205 |
| **TNFR: Fc2** | HTQPTPEPSTAPSTSFLLPMGPSPPAEGSTGDEPK | 24.64851 | 36.19051 | 3.21338 | 205 |
| **TNFR: Fc2** | SQHTQPTPEPSTAPSTSFLLPMGPSPPAEGSTGDEPK | 58.55353 | 85.56929 | 8.38139 | 205 |
| **TNFR: Fc2** | SQHTQPTPEPSTAPSTSFLLPMGPSPPAEGSTGD | 6.92069 | 10.64721 | 0 | 226 |
| **TNFR: Fc2** | SQHTQPTPEPSTAPSTSFLLPMGPSPPAEGSTGD | 18.01555 | 23.42604 | 7.9675 | 226 |
| **TNFR: Fc2** | SQHTQPTPEPSTAPSTSFLLPMG | 5.64953 | 8.69159 | 0 | 202 |
| **TNFR: Fc2** | SQHTQPTPEPSTAPSTSFLLPMGPS | 6.15516 | 9.46947 | 0 | 202 |
| **TNFR: Fc2** | SQHTQPTPEPSTAPSTSFLLPM | 6.63556 | 10.20856 | 0 | 202 |
| **TNFR: Fc2** | SQHTQPTPEPSTAPSTSFLLPMGPSPPAEGST | 14.85362 | 22.85173 | 0 | 226 |
| **TNFR: Fc2** | SQHTQPTPEPSTAPSTSFLLPMGPSPPAEGST | 4.01813 | 5.59986 | 1.08064 | 218 |
| **TNFR: Fc2** | SQHTQPTPEPSTAPSTSFLLPMGPSPP | 13.2794 | 20.42984 | 0 | 226 |

| Sample | Peptide | TotalScore | PepScore | GlyScore | ProSites |
|---|---|---|---|---|---|
| **TNFR: Fc2** | SQHTQPTPEPSTAPSTSFLLPMGPSPPAE | 9.21408 | 14.17551 | 0 | 226 |
| **TNFR: Fc2** | SQHTQPTPEPSTAPSTSFLLPMGPSPPAEGSTGDE | 10.93187 | 16.81826 | 0 | 226 |
| **TNFR: Fc2** | TQPTPEPSTAPSTSFLLPMGPSPPAEGSTGDEPK | 10.2124 | 15.71138 | 0 | 216 |
| **TNFR: Fc2** | SQHTQPTPEPSTAPSTSFLLPMGPSPPAEG | 16.58567 | 20.28569 | 9.71419 | 226 |
| **TNFR: Fc2** | SQHTQPTPEPSTAPSTSFLLPMGPSPPAEGSTGD | 22.46618 | 33.84447 | 1.33506 | 226 |
| **TNFR: Fc2** | SQHTQPTPEPSTAPSTSFLLPMGPSPPAEGSTGD | 35.58398 | 49.01798 | 10.63513 | 217 |
| **TNFR: Fc2** | SQHTQPTPEPSTAPSTSFLLPMGPSPPAEGSTGD | 35.22175 | 53.52315 | 1.23343 | 226 |
| **TNFR: Fc2** | STAPSTSFLLPMGPSPPAEGSTGDEPK | 5.23846 | 8.05916 | 0 | 212 |
| **TNFR: Fc2** | SQHTQPTPEPSTAPSTSFLLPMGPSPPAEG | 29.66519 | 42.82617 | 5.22335 | 202 |
| **TNFR: Fc2** | SQHTQPTPEPSTAPSTSFLLPMGPSPPAEGSTGD | 41.09838 | 60.7001 | 4.69519 | 216 |
| **TNFR: Fc2** | SQHTQPTPEPSTAPSTSFLLPMGPSPPAEGSTGD | 49.95726 | 73.85914 | 5.56803 | 202 |
| **TNFR: Fc2** | SQHTQPTPEPSTAPSTSFLLPMGPSPPAEGSTGD | 11.94646 | 15.68258 | 5.00795 | 202 |
| **TNFR: Fc2** | SQHTQPTPEPSTAPSTSFLLPMGPSPPAEGSTGD | 6.8857 | 8.14265 | 4.55136 | 202 |
| **TNFR: Fc2** | PEPSTAPSTSFLLPMGPSPPAEGSTGDEPK | 3.9773 | 5.58961 | 0.98302 | 212 |
| **TNFR: Fc2** | PTPEPSTAPSTSFLLPMGPSPPAEGSTGDEPK | 18.79439 | 28.43062 | 0.89854 | 212 |
| **TNFR: Fc2** | SMAPGAVHLPQPVSTR | 10.6531 | 16.38938 | 0 | 199 |
| **TNFR: Fc2** | SMAPGAVHLPQPVSTR | 13.0328 | 18.14825 | 3.53268 | 199 |
| **TNFR: Fc2** | AVHLPQPVSTR | 26.26552 | 37.76988 | 4.90029 | 199 |
| **TNFR: Fc2** | APGAVHLPQPVSTR | 45.66991 | 67.45547 | 5.21102 | 199 |
| **TNFR: Fc2** | SMAPGAVHLPQPVSTR | 53.05545 | 78.50039 | 5.80055 | 199 |
| **TNFR: Fc2** | SMAPGAVHLPQPVSTR | 50.60997 | 74.65715 | 5.95091 | 199 |
| **TNFR: Fc2** | SMAPGAVHLPQPVSTR | 54.2849 | 80.28616 | 5.99686 | 199 |
| **TNFR: Fc2** | SMAPGAVHLPQPVSTR | 24.57654 | 31.68018 | 11.38408 | 199 |
| **TNFR: Fc2** | SMAPGAVHLPQPVSTR | 18.90958 | 29.09166 | 0 | 199 |
| **TNFR: Fc2** | SMAPGAVHLPQPVSTR | 13.99234 | 21.52668 | 0 | 199 |
| **TNFR: Fc2** | SMAPGAVHLPQPVSTR | 40.68947 | 59.59208 | 5.58462 | 199 |
| **TNFR: Fc2** | SMAPGAVHLPQPVSTR | 46.43569 | 68.42449 | 5.59934 | 199 |
| **TNFR: Fc2** | SMAPGAVHLPQPVSTR | 50.07439 | 74.01528 | 5.61274 | 199 |
| **TNFR: Fc2** | SMAPGAVHLPQPVSTR | 46.88798 | 69.10573 | 5.62644 | 199 |
| **TNFR: Fc2** | MAPGAVHLPQPVSTR | 30.99613 | 44.96539 | 5.0532 | 199 |
| **TNFR: Fc2** | SMAPGAVHLPQPVSTR | 32.78418 | 47.78123 | 4.93251 | 200 |
| **TNFR: Fc2** | SMAPGAVHLPQPVSTR | 32.62197 | 47.59949 | 4.80659 | 200 |
| **TNFR: Fc2** | SMAPGAVHLPQPVSTR | 46.08715 | 67.9295 | 5.52278 | 199 |
| **TNFR: Fc2** | SMAPGAVHLPQPVSTR | 19.12466 | 24.02536 | 10.02336 | 199 |
| **TNFR: Fc2** | SMAPGAVHLPQPVSTR | 54.73377 | 81.38349 | 5.24143 | 199 |
| **TNFR: Fc2** | SMAPGAVHLPQPVSTR | 42.23409 | 61.98719 | 5.54978 | 199 |
| **TNFR: Fc2** | SMAPGAVHLPQPVSTR | 52.27822 | 77.38536 | 5.65067 | 199 |
| **TNFR: Fc2** | SMAPGAVHLPQPVSTR | 44.52587 | 65.6187 | 5.35349 | 199 |
| **TNFR: Fc2** | APGAVHLPQPVSTR | 37.6565 | 55.0587 | 5.33814 | 199 |
| **TNFR: Fc2** | SMAPGAVHLPQPVSTR | 14.96389 | 23.02137 | 0 | 199 |
| **TNFR: Fc2** | PGAVHLPQPVSTR | 68.30573 | 99.10769 | 11.1021 | 199 |
| **TNFR: Fc2** | PGAVHLPQPVSTR | 50.6632 | 72.39398 | 10.30603 | 199 |

| Sample | Peptide | TotalScore | PepScore | GlyScore | ProSites |
|---|---|---|---|---|---|
| **TNFR: Fc2** | SMAPGAVHLPQPVSTR | 88.6062 | 132.3665 | 7.33702 | 199 |
| **TNFR: Fc2** | APGAVHLPQPVSTR | 48.53007 | 71.91528 | 5.1004 | 199 |
| **TNFR: Fc2** | SMAPGAVHLPQPVSTR | 19.94819 | 30.68952 | 0 | 199 |
| **TNFR: Fc2** | SMAPGAVHLPQPVSTR | 67.48005 | 100.5609 | 6.0442 | 199 |
| **TNFR: Fc2** | SMAPGAVHLPQPVSTR | 60.21973 | 89.43101 | 5.97023 | 199 |
| **TNFR: Fc2** | SMAPGAVHLPQPVSTR | 47.81158 | 70.33734 | 5.97801 | 199 |
| **TNFR: Fc2** | SMAPGAVHLPQPVSTR | 57.35656 | 85.23627 | 5.57995 | 199 |
| **TNFR: Fc2** | SMAPGAVHLPQPVSTR | 39.48743 | 57.94776 | 5.20397 | 199 |
| **TNFR: Fc2** | SMAPGAVHLPQPVSTR | 49.97529 | 74.15431 | 5.07139 | 199 |
| **TNFR: Fc2** | SMAPGAVHLPQPVSTR | 25.1222 | 35.97326 | 4.97024 | 199 |
| **TNFR: Fc2** | SMAPGAVHLPQPVSTR | 39.51576 | 58.28089 | 4.66625 | 199 |
| **TNFR: Fc2** | SMAPGAVHLPQPVSTR | 19.27733 | 27.41179 | 4.17048 | 199 |
| **TNFR: Fc2** | SMAPGAVHLPQPVSTR | 27.65853 | 39.93642 | 4.85676 | 199 |
| **TNFR: Fc2** | SMAPGAVHLPQPVS | 45.17076 | 63.92118 | 10.34855 | 199 |
| **TNFR: Fc2** | SQHTQPTPEPSTAPSTSFLLPMGPSPPAEGSTGDEPK | 82.71209 | 120.8766 | 11.83506 | 217 |
| **TNFR: Fc2** | SQHTQPTPEPSTAPSTSFLLPMGPSPPAEGSTGDEPK | 74.30603 | 109.45 | 9.03862 | 217 |
| **TNFR: Fc2** | SQHTQPTPEPSTAPSTSFLL | 13.96406 | 21.48317 | 0 | 205 |
| **TNFR: Fc2** | QPTPEPSTAPSTSFLLPMGPSPPAEGSTGDEPK | 7.59202 | 11.68003 | 0 | 208 |
| **TNFR: Fc2** | SQHTQPTPEPSTAPSTSFLLPMGPSPPAEGSTGDE | 8.04606 | 12.37855 | 0 | 216 |
| **TNFR: Fc2** | SQHTQPTPEPSTAPSTSFLLPMGPSPPAEGSTGDEPK | 65.50302 | 94.39288 | 11.85043 | 213 |
| **TNFR: Fc2** | SQHTQPTPEPSTAPSTSFLLPMGPSPPAEGSTGDEPK | 50.9774 | 73.46748 | 9.21011 | 217 |
| **TNFR: Fc2** | SQHTQPTPEPSTAPSTSFLLPMGPSPPAEGSTGDEPK | 89.46388 | 134.3431 | 6.11674 | 205 |
| **TNFR: Fc2** | SQHTQPTPEPSTAPSTSFLLPMGPSPPAEGSTGDEPK | 70.60736 | 103.4303 | 9.6504 | 205 |
| **TNFR: Fc2** | SQHTQPTPEPSTAPSTSFLLPMGPSPPAEGSTGDEPK | 69.4665 | 103.6193 | 6.03986 | 205 |
| **TNFR: Fc2** | SQHTQPTPEPSTAPSTSFLLPMGPSPPAEGSTGDEPK | 64.53691 | 96.35088 | 5.45384 | 205 |
| **TNFR: Fc2** | SQHTQPTPEPSTAPSTSFLLPMGPSPPAEGSTGDEPK | 77.72811 | 116.6862 | 5.37728 | 213 |
| **TNFR: Fc2** | SQHTQPTPEPSTAPSTSFLLPMGPSPPAEGSTGDEPK | 91.6579 | 135.6603 | 9.93918 | 217 |
| **TNFR: Fc2** | SQHTQPTPEPSTAPSTSFLLPMGPSPPAEGSTGDEPK | 42.76557 | 65.79318 | 0 | 205 |
| **TNFR: Fc2** | SQHTQPTPEPSTAPSTSFLLPMGPSPPAEGSTGDEPK | 59.1539 | 87.82124 | 5.91456 | 213 |
| **TNFR: Fc2** | SQHTQPTPEPSTAPSTSFLLPMGPSPPAEGSTGDEPK | 57.95381 | 86.56827 | 4.81267 | 205 |
| **TNFR: Fc2** | SQHTQPTPEPSTAPSTSFLLPMGPSPPAEGSTGDEPK | 68.30439 | 105.0837 | 0 | 217 |
| **TNFR: Fc2** | SQHTQPTPEPSTAPSTSFLLPMGPSPPAEGSTGDEPK | 61.03994 | 91.44392 | 4.5754 | 213 |
| **TNFR: Fc2** | SQHTQPTPEPSTAPSTSFLLPMGPSPPAEGSTGDEPK | 65.23347 | 97.20181 | 5.86372 | 205 |
| **TNFR: Fc2** | SQHTQPTPEPSTAPSTSFLLPMGPSPPAEGSTGDEPK | 60.93366 | 91.046 | 5.01073 | 213 |
| **TNFR: Fc2** | SQHTQPTPEPSTAPSTSFLLPMGPSPPAEGSTGDEPK | 28.33743 | 40.99558 | 4.82945 | 213 |
| **TNFR: Fc2** | SQHTQPTPEPSTAPSTSFLLPMGPSPPAEGSTGDEPK | 43.08045 | 66.27761 | 0 | 205 |
| **TNFR: Fc2** | SQHTQPTPEPSTAPSTSFLLPMGPSPPAEGSTGDEPK | 59.58878 | 88.67132 | 5.57837 | 213 |
| **TNFR: Fc2** | SQHTQPTPEPSTAPSTSFLLPMGPSPPAEGSTGDEPK | 52.34653 | 78.1139 | 4.49285 | 217 |
| **TNFR: Fc2** | SQHTQPTPEPSTAPSTSFLLPMGPSPPAEGSTGDEPK | 61.31652 | 91.74483 | 4.8068 | 213 |
| **TNFR: Fc2** | SQHTQPTPEPSTAPSTSFLLPMGPSPPAEGSTGDEPK | 42.75376 | 63.21362 | 4.75688 | 205 |
| **TNFR: Fc2** | PTPEPSTAPSTSFLLPMGPSPPAEGSTGDEPK | 6.05422 | 9.31419 | 0 | 208 |
| **TNFR: Fc2** | SQHTQPTPEPSTAPSTSFLLPMGPSPPAEGSTGDEPK | 104.743 | 154.0046 | 13.25694 | 226 |

| Sample | Peptide | TotalScore | PepScore | GlyScore | ProSites |
|---|---|---|---|---|---|
| **TNFR: Fc2** | SQHTQPTPEPSTAPSTSFLLPMGPSPPAEGSTGDEPK | 90.04701 | 131.2452 | 13.53603 | 226 |
| **TNFR: Fc2** | SQHTQPTPEPSTAPSTSFLLPMGPSPPAEGSTGDEPK | 64.47643 | 96.23715 | 5.49223 | 216 |
| **TNFR: Fc2** | SQHTQPTPEPSTAPSTSFLLPMGPSPPAEGSTGDEPK | 69.68888 | 103.0011 | 7.82326 | 213 |
| **TNFR: Fc2** | SQHTQPTPEPSTAPSTSFLLPMGPSPPAEGSTGDEPK | 58.17923 | 87.02216 | 4.61378 | 213 |
| **TNFR: Fc2** | SQHTQPTPEPSTAPSTSFLLPMGPSPPAEGSTGDEPK | 82.10575 | 122.9004 | 6.34423 | 216 |
| **TNFR: Fc2** | SQHTQPTPEPSTAPSTSFLLPMGPSPPAEGSTGDEPK | 47.12543 | 72.50065 | 0 | 217 |
| **TNFR: Fc2** | SQHTQPTPEPSTAPSTSFLLPMGPSPPAEGSTGDEPK | 119.78 | 177.5361 | 12.51875 | 213 |
| **TNFR: Fc2** | SQHTQPTPEPSTAPSTSFLLPMGPSPPAEGSTGDEPK | 62.82076 | 94.06165 | 4.80198 | 212 |
| **TNFR: Fc2** | SQHTQPTPEPSTAPSTSFLLPMGPSPPAEGSTGDEPK | 6.85245 | 10.54222 | 0 | 216 |
| **TNFR: Fc2** | SQHTQPTPEPSTAPST | 3.30449 | 5.08383 | 0 | 216 |
| **TNFR: Fc2** | HTQPTPEPSTAPSTSFLLPMGPSPPAEGSTGDEPK | 22.50859 | 33.95395 | 1.25293 | 226 |
| **TNFR: Fc2** | SQHTQPTPEPSTAPSTSFLLPMGPSPPAEGSTGDEPK | 65.73363 | 98.56947 | 4.75278 | 213 |
| **TNFR: Fc2** | SQHTQPTPEPSTAPSTSFLLPMGPSPPAEGSTGDEPK | 3.64853 | 5.61313 | 0 | 205 |
| **TNFR: Fc2** | HTQPTPEPSTAPSTSFLLPMGPSPPAEGSTGDEPK | 14.90477 | 22.93041 | 0 | 216 |
| **TNFR: Fc2** | HTQPTPEPSTAPSTSFLLPMGPSPPAEGSTGDEPK | 7.43864 | 10.81035 | 1.17688 | 226 |
| **TNFR: Fc2** | SQHTQPTPEPSTAPSTSFLLPMGPSPPAEGSTGDEPK | 52.4289 | 78.22384 | 4.52401 | 205 |
| **TNFR: Fc2** | QPTPEPSTAPSTSFLLPMGPSPPAEGSTGDEPK | 8.73163 | 13.43327 | 0 | 208 |
| **TNFR: Fc2** | SQHTQPTPEPSTAPSTSFLLPMGPSPPAEGSTGDEPK | 87.39342 | 131.055 | 6.30768 | 205 |
| **TNFR: Fc2** | SQHTQPTPEPSTAPSTSFLLPMGPSPPAEGSTGDEPK | 55.44953 | 82.75999 | 4.7301 | 205 |
| **TNFR: Fc2** | SQHTQPTPEPSTAPSTSFLLPMGPSPPAEGSTGDEPK | 130.4139 | 196.7964 | 7.13207 | 205 |
| **TNFR: Fc2** | SQHTQPTPEPSTAPSTSFLLPMGPSPPAEGSTGDEPK | 104.2861 | 156.7503 | 6.85259 | 205 |
| **TNFR: Fc2** | SQHTQPTPEPSTAPSTSFLLPMGPSPPAEGSTGDEPK | 101.5804 | 152.5878 | 6.8523 | 205 |
| **TNFR: Fc2** | SQHTQPTPEPSTAPSTSFLLPMGPSPPAEGSTGDEPK | 78.97327 | 117.8587 | 6.75742 | 205 |
| **TNFR: Fc2** | SQHTQPTPEPSTAPSTSFLLPMGPSPPAEGSTGDEPK | 65.23919 | 100.368 | 0 | 205 |
| **TNFR: Fc2** | SQHTQPTPEPSTAPSTSFLLPMGPSPPAEGSTGDEPK | 53.3798 | 79.18032 | 5.46455 | 213 |
| **TNFR: Fc2** | SQHTQPTPEPSTAPSTSFLLPMGPSPPAEGSTGDEPK | 53.98808 | 80.95161 | 3.91296 | 218 |
| **TNFR: Fc2** | SQHTQPTPEPSTAPSTSFLLPMGPSPPAEGSTGDEPK | 62.56028 | 93.39185 | 5.30164 | 205 |
| **TNFR: Fc2** | SQHTQPTPEPSTAPSTSFLLPMGPSPPAEGSTGDEPK | 62.63026 | 93.8375 | 4.67394 | 213 |
| **TNFR: Fc2** | SQHTQPTPEPSTAPSTSFLLPMGPSPPAEGSTGDEPK | 97.89646 | 147.7463 | 5.31827 | 205 |
| **TNFR: Fc2** | SQHTQPTPEPSTAPSTSFLLPMGPSPPAEGSTGDEPK | 45.83205 | 68.01772 | 4.63008 | 208 |
| **TNFR: Fc2** | SQHTQPTPEPSTAPSTSFLLPMGPSPPAEGSTGDEPK | 55.14603 | 82.38571 | 4.55804 | 205 |
| **TNFR: Fc2** | SQHTQPTPEPSTAPSTSFLLPMGPSPPAEGSTGDEPK | 84.45522 | 127.2494 | 4.98037 | 213 |
| **TNFR: Fc2** | HTQPTPEPSTAPSTSFLLPMGPSPPAEGSTGDEPK | 25.18504 | 38.74622 | 0 | 205 |
| **TNFR: Fc2** | SQHTQPTPEPSTAPSTSFLLPMGPSPPAEGSTGDEPK | 58.50534 | 90.00821 | 0 | 205 |
| **TNFR: Fc2** | HTQPTPEPSTAPSTSFLLPMGPSPPAEGSTGDEPK | 5.83286 | 8.97363 | 0 | 205 |
| **TNFR: Fc2** | SQHTQPTPEPSTAPSTSFLLPMGPSPPAEGSTGD | 10.94668 | 16.18827 | 1.21231 | 226 |
| **TNFR: Fc2** | SQHTQPTPEPSTAPSTSFLLPMGPSPPAEGSTGD | 21.09497 | 32.4538 | 0 | 226 |
| **TNFR: Fc2** | TQPTPEPSTAPSTSFLLPMGPSPPAEGSTGDEPK | 13.71871 | 21.1057 | 0 | 217 |
| **TNFR: Fc2** | SQHTQPTPEPSTAPSTSFLLPMGPSPPAEGSTGD | 16.59956 | 24.89461 | 1.19446 | 226 |
| **TNFR: Fc2** | TQPTPEPSTAPSTSFLLPMGPSPPAEGSTGDEPK | 15.12591 | 23.27064 | 0 | 205 |
| **TNFR: Fc2** | SQHTQPTPEPSTAPSTSFLLPMGPSPPAEGSTGD | 45.36019 | 67.22931 | 4.74612 | 205 |
| **TNFR: Fc2** | SQHTQPTPEPSTAPSTSFLLPMGPSPPAEGST | 8.72508 | 12.82315 | 1.11437 | 226 |

| Sample | Peptide | TotalScore | PepScore | GlyScore | ProSites |
|---|---|---|---|---|---|
| **TNFR: Fc2** | SQHTQPTPEPSTAPSTSFLLPMGPSPPAEGSTGD | 6.59999 | 10.15383 | 0 | 233 |
| **TNFR: Fc2** | SQHTQPTPEPSTAPSTSFLLPMGPSPP | 4.3075 | 6.02327 | 1.12107 | 226 |
| **TNFR: Fc2** | SQHTQPTPEPSTAPSTSFLLPMGPSPP | 4.93458 | 7.59166 | 0 | 226 |
| **TNFR: Fc2** | SQHTQPTPEPSTAPSTSFLLPMGPSPPAEGSTGDE | 11.73654 | 18.05622 | 0 | 226 |
| **TNFR: Fc2** | SQHTQPTPEPSTAPSTSFLLPMGPSPPAEG | 5.10672 | 7.18606 | 1.24507 | 216 |
| **TNFR: Fc2** | SQHTQPTPEPSTAPSTSFLLPMGPSPPAEGSTGD | 31.33246 | 42.90729 | 9.83634 | 226 |
| **TNFR: Fc2** | SQHTQPTPEPSTAPSTSFLLPMGPSPPAEGSTGD | 31.66861 | 43.05286 | 10.52644 | 226 |
| **TNFR: Fc2** | SQHTQPTPEPSTAPSTSFLLPMGPSPPAEG | 47.08448 | 69.5915 | 5.28573 | 205 |
| **TNFR: Fc2** | SQHTQPTPEPSTAPSTSFLLPMGPSPPAEG | 46.39889 | 68.55122 | 5.25885 | 205 |
| **TNFR: Fc2** | SQHTQPTPEPSTAPSTSFLLPMGPSPPAEGSTGD | 15.63961 | 21.57746 | 4.61217 | 216 |
| **TNFR: Fc2** | SQHTQPTPEPSTAPSTSFLLPMGPSPPAEGSTGDE | 5.8569 | 6.7676 | 4.16558 | 202 |
| **TNFR: Fc2** | SQHTQPTPEPSTAPSTSFLLPMGPSPPAEGSTGD | 48.60295 | 71.94494 | 5.25352 | 202 |
| **TNFR: Fc2** | SQHTQPTPEPSTAPSTSFLLPMGPSPPAEGSTGD | 14.47341 | 22.26678 | 0 | 205 |
| **TNFR: Fc2** | SMAPGAVHLPQPVSTR | 13.19049 | 18.49139 | 3.34597 | 199 |
| **TNFR: Fc3** | AVHLPQPVSTR | 39.12644 | 54.90033 | 9.83208 | 199 |
| **TNFR: Fc3** | AVHLPQPVSTR | 44.17627 | 62.57526 | 10.00671 | 199 |
| **TNFR: Fc3** | AVHLPQPVSTR | 38.65573 | 53.99258 | 10.17301 | 199 |
| **TNFR: Fc3** | SMAPGAVHLPQPVSTR | 49.33053 | 69.69662 | 11.50781 | 199 |
| **TNFR: Fc3** | APGAVHLPQPVSTR | 51.48243 | 76.17728 | 5.62059 | 199 |
| **TNFR: Fc3** | APGAVHLPQPVSTR | 67.16105 | 100.2744 | 5.66479 | 199 |
| **TNFR: Fc3** | SMAPGAVHLPQPVSTR | 43.80703 | 64.52494 | 5.33091 | 200 |
| **TNFR: Fc3** | SMAPGAVHLPQPVSTR | 46.14626 | 68.10854 | 5.35916 | 200 |
| **TNFR: Fc3** | APGAVHLPQPVSTR | 63.85431 | 95.17203 | 5.69284 | 199 |
| **TNFR: Fc3** | SMAPGAVHLPQPVSTR | 46.25402 | 65.24285 | 10.98905 | 199 |
| **TNFR: Fc3** | SMAPGAVHLPQPVSTR | 37.54242 | 52.01633 | 10.66229 | 199 |
| **TNFR: Fc3** | SMAPGAVHLPQPVSTR | 46.93158 | 69.12666 | 5.71214 | 199 |
| **TNFR: Fc3** | SMAPGAVHLPQPVSTR | 55.68193 | 82.42252 | 6.02083 | 199 |
| **TNFR: Fc3** | SMAPGAVHLPQPVSTR | 67.55178 | 100.2529 | 6.82124 | 199 |
| **TNFR: Fc3** | SMAPGAVHLPQPVSTR | 76.44863 | 114.1233 | 6.48142 | 199 |
| **TNFR: Fc3** | SMAPGAVHLPQPVSTR | 61.25364 | 90.70799 | 6.55272 | 199 |
| **TNFR: Fc3** | SMAPGAVHLPQPVSTR | 55.36348 | 81.91422 | 6.05496 | 199 |
| **TNFR: Fc3** | SMAPGAVHLPQPVSTR | 46.8014 | 68.73179 | 6.07353 | 199 |
| **TNFR: Fc3** | APGAVHLPQPVSTR | 11.14775 | 9.36432 | 14.45984 | 199 |
| **TNFR: Fc3** | SMAPGAVHLPQPVSTR | 22.34986 | 29.06181 | 9.8848 | 199 |
| **TNFR: Fc3** | SMAPGAVHLPQPVSTR | 52.93048 | 78.16496 | 6.06644 | 199 |
| **TNFR: Fc3** | SMAPGAVHLPQPVSTR | 20.16273 | 25.74114 | 9.80283 | 199 |
| **TNFR: Fc3** | SMAPGAVHLPQPVSTR | 29.33498 | 39.09879 | 11.20221 | 199 |
| **TNFR: Fc3** | SMAPGAVHLPQPVSTR | 51.55662 | 73.05915 | 11.62335 | 199 |
| **TNFR: Fc3** | SMAPGAVHLPQPVSTR | 59.48043 | 85.18169 | 11.74951 | 199 |
| **TNFR: Fc3** | SMAPGAVHLPQPVSTR | 27.686 | 37.019 | 10.35329 | 199 |
| **TNFR: Fc3** | SMAPGAVHLPQPVSTR | 54.8487 | 81.13769 | 6.0263 | 199 |
| **TNFR: Fc3** | PGAVHLPQPVSTR | 61.90121 | 79.08704 | 29.98467 | 199 |

| Sample | Peptide | TotalScore | PepScore | GlyScore | ProSites |
|---|---|---|---|---|---|
| **TNFR: Fc3** | SMAPGAVHLPQPVSTR | 52.33881 | 77.29358 | 5.99423 | 199 |
| **TNFR: Fc3** | PGAVHLPQPVSTR | 56.79699 | 71.81725 | 28.90224 | 199 |
| **TNFR: Fc3** | APGAVHLPQPVSTR | 48.08104 | 66.1825 | 14.46404 | 199 |
| **TNFR: Fc3** | PGAVHLPQPVSTR | 55.09955 | 70.67396 | 26.17565 | 199 |
| **TNFR: Fc3** | SMAPGAVHLPQPVSTR | 63.36104 | 96.44972 | 1.91063 | 199 |
| **TNFR: Fc3** | SMAPGAVHLPQPVSTR | 15.347 | 23.61077 | 0 | 199 |
| **TNFR: Fc3** | SMAPGAVHLPQPVSTR | 53.89399 | 80.05012 | 5.31832 | 200 |
| **TNFR: Fc3** | SMAPGAVHLPQPVSTR | 58.70674 | 87.03231 | 6.10212 | 199 |
| **TNFR: Fc3** | SMAPGAVHLPQPVSTR | 56.65052 | 84.05539 | 5.75577 | 200 |
| **TNFR: Fc3** | SMAPGAVHLPQPVSTR | 58.98595 | 87.5287 | 5.97798 | 200 |
| **TNFR: Fc3** | SMAPGAVHLPQPVSTR | 60.45889 | 89.78172 | 6.00221 | 199 |
| **TNFR: Fc3** | SMAPGAVHLPQPVSTR | 83.29813 | 124.7356 | 6.34292 | 200 |
| **TNFR: Fc3** | SMAPGAVHLPQPVSTR | 11.23848 | 14.35697 | 5.447 | 199 |
| **TNFR: Fc3** | SMAPGAVHLPQPVSTR | 57.5861 | 85.44057 | 5.85637 | 200 |
| **TNFR: Fc3** | SMAPGAVHLPQPVSTR | 13.06732 | 17.10654 | 5.56591 | 199 |
| **TNFR: Fc3** | SMAPGAVHLPQPVSTR | 43.34498 | 65.86311 | 1.5256 | 199 |
| **TNFR: Fc3** | SMAPGAVHLPQPVSTR | 49.66336 | 70.45372 | 11.05267 | 199 |
| **TNFR: Fc3** | SMAPGAVHLPQPVSTR | 50.58642 | 74.53324 | 6.11374 | 199 |
| **TNFR: Fc3** | SMAPGAVHLPQPVSTR | 59.56685 | 88.81097 | 5.25633 | 199 |
| **TNFR: Fc3** | SMAPGAVHLPQPVSTR | 54.42636 | 80.73075 | 5.57537 | 199 |
| **TNFR: Fc3** | SMAPGAVHLPQPVSTR | 54.13722 | 80.29669 | 5.55536 | 199 |
| **TNFR: Fc3** | APGAVHLPQPVSTR | 64.87722 | 93.3766 | 11.9498 | 199 |
| **TNFR: Fc3** | PGAVHLPQPVSTR | 47.45232 | 69.39057 | 6.70985 | 199 |
| **TNFR: Fc3** | SMAPGAVHLPQPVSTR | 50.40217 | 68.90915 | 16.03207 | 199 |
| **TNFR: Fc3** | SMAPGAVHLPQPVSTR | 48.6761 | 71.77928 | 5.7702 | 199 |
| **TNFR: Fc3** | SMAPGAVHLPQPVSTR | 45.56972 | 67.14625 | 5.49904 | 199 |
| **TNFR: Fc3** | PGAVHLPQPVSTR | 49.66899 | 70.47131 | 11.0361 | 199 |
| **TNFR: Fc3** | APGAVHLPQPVSTR | 58.31878 | 86.61812 | 5.76287 | 199 |
| **TNFR: Fc3** | PGAVHLPQPVSTR | 31.65376 | 43.19278 | 10.22414 | 199 |
| **TNFR: Fc3** | SMAPGAVHLPQPVSTR | 90.64192 | 135.5784 | 7.18841 | 199 |
| **TNFR: Fc3** | APGAVHLPQPVSTR | 43.56807 | 64.19301 | 5.26461 | 199 |
| **TNFR: Fc3** | SMAPGAVHLPQPVSTR | 25.09044 | 38.60068 | 0 | 199 |
| **TNFR: Fc3** | SMAPGAVHLPQPVSTR | 63.7113 | 94.64127 | 6.26993 | 199 |
| **TNFR: Fc3** | SMAPGAVHLPQPVSTR | 56.34142 | 83.61037 | 5.69909 | 199 |
| **TNFR: Fc3** | SMAPGAVHLPQPVSTR | 47.63877 | 70.21953 | 5.70308 | 199 |
| **TNFR: Fc3** | SMAPGAVHLPQPVSTR | 62.46055 | 93.10739 | 5.54499 | 199 |
| **TNFR: Fc3** | SMAPGAVHLPQPVSTR | 47.75652 | 70.5643 | 5.39922 | 199 |
| **TNFR: Fc3** | SMAPGAVHLPQPVSTR | 51.81113 | 76.82534 | 5.35615 | 199 |
| **TNFR: Fc3** | REPQVYTLPPSR | 4.30795 | 5.62436 | 1.8632 | 374 |
| **TNFR: Fc3** | SMAPGAVHLPQPVSTR | 50.30828 | 74.46598 | 5.44397 | 199 |
| **TNFR: Fc3** | SMAPGAVHLPQPVSTR | 38.15281 | 55.93174 | 5.13481 | 199 |
| **TNFR: Fc3** | SMAPGAVHLPQPVSTR | 24.69604 | 35.3549 | 4.90103 | 199 |

| Sample | Peptide | TotalScore | PepScore | GlyScore | ProSites |
|---|---|---|---|---|---|
| **TNFR: Fc3** | SMAPGAVHLPQPVSTR | 29.06661 | 42.07051 | 4.91652 | 199 |
| **TNFR: Fc3** | SQHTQPTPEPSTAPSTSFLLPMGPSPPAEGSTGDEPK | 6.66942 | 8.33813 | 3.5704 | 208 |
| **TNFR: Fc3** | SQHTQPTPEPSTAPSTSFLLPMGPSPPAEGSTGDEPK | 33.71638 | 49.39315 | 4.60237 | 205 |
| **TNFR: Fc3** | SQHTQPTPEPSTAPSTSFLLPMGPSPPAEGSTGDEPK | 40.98218 | 60.49686 | 4.74062 | 205 |
| **TNFR: Fc3** | SQHTQPTPEPSTAPSTSFLLPMGPSPPAEGSTGDEPK | 48.19257 | 71.35548 | 5.17572 | 217 |
| **TNFR: Fc3** | SQHTQPTPEPSTAPSTSFLLPMGPSPPAEGSTGDEPK | 94.7225 | 142.3691 | 6.23603 | 212 |
| **TNFR: Fc3** | SQHTQPTPEPSTAPSTSFLLPMGPSPPAEGSTGDEPK | 80.98655 | 121.3104 | 6.09936 | 217 |
| **TNFR: Fc3** | SQHTQPTPEPSTAPSTSFLLPMGPSPPAEGSTGDEPK | 98.27159 | 147.7332 | 6.41426 | 213 |
| **TNFR: Fc3** | SQHTQPTPEPSTAPSTSFLLPMGPSPPAEGSTGDEPK | 89.4651 | 131.282 | 11.80506 | 205 |
| **TNFR: Fc3** | SQHTQPTPEPSTAPSTSFLLPMGPSPPAEGSTGDEPK | 99.5189 | 149.5369 | 6.62824 | 205 |
| **TNFR: Fc3** | SQHTQPTPEPSTAPSTSFLLPMGPSPPAEGSTGDEPK | 92.08455 | 138.3758 | 6.11513 | 205 |
| **TNFR: Fc3** | SQHTQPTPEPSTAPSTSFLLPMGPSPPAEGSTGDEPK | 91.73349 | 137.4858 | 6.765 | 205 |
| **TNFR: Fc3** | SQHTQPTPEPSTAPSTSFLLPMGPSPPAEGSTGDEPK | 84.71025 | 123.7142 | 12.27435 | 205 |
| **TNFR: Fc3** | SQHTQPTPEPSTAPSTSFLLPMGPSPPAEGSTGDEPK | 93.64733 | 140.5594 | 6.52489 | 205 |
| **TNFR: Fc3** | SQHTQPTPEPSTAPSTSFLLPMGPSPPAEGSTGDEPK | 104.3556 | 160.5471 | 0 | 205 |
| **TNFR: Fc3** | SQHTQPTPEPSTAPSTSFLLPMGPSPPAEGSTGDEPK | 77.04225 | 112.6528 | 10.90828 | 205 |
| **TNFR: Fc3** | SQHTQPTPEPSTAPSTSFLLPMGPSPPAEGSTGDEPK | 97.8637 | 147.2294 | 6.18449 | 205 |
| **TNFR: Fc3** | SQHTQPTPEPSTAPSTSFLLPMGPSPPAEGSTGDEPK | 67.73031 | 98.69957 | 10.21596 | 205 |
| **TNFR: Fc3** | SQHTQPTPEPSTAPSTSFLLPMGPSPPAEGSTGDEPK | 88.30223 | 135.8496 | 0 | 205 |
| **TNFR: Fc3** | SQHTQPTPEPSTAPSTSFLLPMGPSPPAEGSTGDEPK | 71.49416 | 107.6377 | 4.37037 | 205 |
| **TNFR: Fc3** | LPAQVAFTPYAPEPGSTCR | 90.53563 | 135.9867 | 6.12642 | 8 |
| **TNFR: Fc3** | LPAQVAFTPYAPEPGSTCR | 108.6875 | 163.8192 | 6.30007 | 8 |
| **TNFR: Fc3** | SQHTQPTPEPSTAPSTSFLLPMGPSPPAEGSTGDEPK | 68.89631 | 103.099 | 5.37696 | 205 |
| **TNFR: Fc3** | SQHTQPTPEPSTAPSTSFLLPMGPSPPAEGSTGDEPK | 75.22982 | 112.5226 | 5.97182 | 213 |
| **TNFR: Fc3** | SQHTQPTPEPSTAPSTSFLLPMGPSPPAEGSTGDEPK | 71.28238 | 106.8078 | 5.30665 | 213 |
| **TNFR: Fc3** | SQHTQPTPEPSTAPSTSFLLPMGPSPPAEGSTGDEPK | 57.25401 | 82.85419 | 9.7108 | 205 |
| **TNFR: Fc3** | SQHTQPTPEPSTAPSTSFLLPMGPSPPAEGSTGDEPK | 70.13876 | 102.7696 | 9.53863 | 205 |
| **TNFR: Fc3** | SQHTQPTPEPSTAPSTSFLLPMGPSPPAEGSTGDEPK | 71.20677 | 104.3529 | 9.64965 | 205 |
| **TNFR: Fc3** | SQHTQPTPEPSTAPSTSFLLPMGPSPPAEGSTGDEPK | 76.03271 | 111.7945 | 9.61799 | 205 |
| **TNFR: Fc3** | SQHTQPTPEPSTAPSTSFLLPMGPSPPAEGSTGDEPK | 67.4884 | 100.989 | 5.27302 | 205 |
| **TNFR: Fc3** | SQHTQPTPEPSTAPSTSFLLPMGPSPPAEGSTGDEPK | 64.72869 | 96.76542 | 5.2319 | 213 |
| **TNFR: Fc3** | SQHTQPTPEPSTAPSTSFLLPMGPSPPAEGSTGDEPK | 26.83205 | 41.28008 | 0 | 216 |
| **TNFR: Fc3** | SQHTQPTPEPSTAPSTSFLLPMGPSPPAEGSTGDEPK | 75.21042 | 112.6283 | 5.72013 | 205 |
| **TNFR: Fc3** | SQHTQPTPEPSTAPSTSFLLPMGPSPPAEGSTGDEPK | 80.39438 | 120.7318 | 5.48204 | 213 |
| **TNFR: Fc3** | SQHTQPTPEPSTAPSTSFLLPMGPSPPAEGSTGDEPK | 79.09608 | 118.7151 | 5.51786 | 205 |
| **TNFR: Fc3** | SQHTQPTPEPSTAPSTSFLLPMGPSPPAEGSTGDEPK | 103.6676 | 156.3023 | 5.91733 | 205 |
| **TNFR: Fc3** | SQHTQPTPEPSTAPSTSFLLPMGPSPPAEGSTGDEPK | 70.52244 | 103.4075 | 9.45027 | 213 |
| **TNFR: Fc3** | SQHTQPTPEPSTAPSTSFLLPMGPSPPAEGSTGDEPK | 91.05735 | 132.9789 | 13.20309 | 226 |
| **TNFR: Fc3** | SQHTQPTPEPSTAPSTSFLLPMGPSPPAEGSTGDEPK | 70.06021 | 104.7282 | 5.67675 | 205 |
| **TNFR: Fc3** | SQHTQPTPEPSTAPSTSFLLPMGPSPPAEGSTGDEPK | 72.43755 | 108.25 | 5.92867 | 218 |
| **TNFR: Fc3** | SQHTQPTPEPSTAPSTSFLLPMGPSPPAEGSTGDEPK | 78.05549 | 116.9507 | 5.82162 | 218 |
| **TNFR: Fc3** | SQHTQPTPEPSTAPSTSFLLPMGPSPPAEGSTGDEPK | 77.8865 | 116.3662 | 6.42419 | 216 |

| Sample | Peptide | TotalScore | PepScore | GlyScore | ProSites |
|---|---|---|---|---|---|
| **TNFR: Fc3** | SQHTQPTPEPSTAPSTSFLLPMGPSPPAEGSTGDEPK | 74.17621 | 111.1694 | 5.47468 | 217 |
| **TNFR: Fc3** | SQHTQPTPEPSTAPSTSFLLPMGPSPPAEGSTGDEPK | 116.4643 | 175.3654 | 7.07654 | 217 |
| **TNFR: Fc3** | SQHTQPTPEPSTAPSTSFLLPMGPSPPAEGSTGDEPK | 65.52614 | 97.93014 | 5.3473 | 205 |
| **TNFR: Fc3** | SQHTQPTPEPSTAPSTSFLLPMGPSPPAEGSTGDEPK | 90.74902 | 135.7653 | 7.14728 | 216 |
| **TNFR: Fc3** | SQHTQPTPEPSTAPSTSFLLPMGPSPPAEGSTGDEPK | 65.85769 | 98.49308 | 5.24912 | 213 |
| **TNFR: Fc3** | SQHTQPTPEPSTAPSTSFLLPMGPSPPAEGSTGDEPK | 89.41914 | 133.8103 | 6.97845 | 213 |
| **TNFR: Fc3** | HTQPTPEPSTAPSTSFLLPMGPSPPAEGSTGDEPK | 44.24472 | 67.30084 | 1.42621 | 226 |
| **TNFR: Fc3** | HTQPTPEPSTAPSTSFLLPMGPSPPAEGSTGDEPK | 48.79646 | 74.33724 | 1.36359 | 226 |
| **TNFR: Fc3** | SQHTQPTPEPSTAPSTSFLLPMGPSPPAEGSTGDEPK | 98.21165 | 147.4514 | 6.76649 | 213 |
| **TNFR: Fc3** | SQHTQPTPEPSTAPSTSFLLPMGPSPPAEGSTGDEPK | 130.0022 | 196.4251 | 6.64529 | 213 |
| **TNFR: Fc3** | HTQPTPEPSTAPSTSFLLPMGPSPPAEGSTGDEPK | 36.63385 | 55.61231 | 1.38815 | 226 |
| **TNFR: Fc3** | SQHTQPTPEPSTAPSTSFLLPMGPSPPAEGSTGDEPK | 71.83443 | 107.6555 | 5.30953 | 205 |
| **TNFR: Fc3** | SQHTQPTPEPSTAPSTSFLLPMGPSPPAEGSTGDEPK | 98.49567 | 147.6678 | 7.17601 | 205 |
| **TNFR: Fc3** | SQHTQPTPEPSTAPSTSFLLPMGPSPPAEGSTGDEPK | 66.37425 | 97.75727 | 8.09149 | 205 |
| **TNFR: Fc3** | SQHTQPTPEPSTAPSTSFLLPMGPSPPAEGSTGDEPK | 91.59019 | 136.9467 | 7.35667 | 205 |
| **TNFR: Fc3** | SQHTQPTPEPSTAPSTSFLLPMGPSPPAEGSTGDEPK | 134.6771 | 199.8948 | 13.55849 | 205 |
| **TNFR: Fc3** | SQHTQPTPEPSTAPSTSFLLPMGPSPPAEGSTGDEPK | 67.1533 | 103.3128 | 0 | 208 |
| **TNFR: Fc3** | SQHTQPTPEPSTAPSTSFLLPMGPSPPAEGSTGDEPK | 68.95523 | 106.085 | 0 | 205 |
| **TNFR: Fc3** | SQHTQPTPEPSTAPSTSFLLPMGPSPPAEGSTGDEPK | 74.74001 | 112.0262 | 5.4943 | 205 |
| **TNFR: Fc3** | QHTQPTPEPSTAPSTSFLLPMGPSPPAEGSTGDEPK | 15.19104 | 23.37082 | 0 | 217 |
| **TNFR: Fc3** | QHTQPTPEPSTAPSTSFLLPMGPSPPAEGSTGDEPK | 10.79165 | 16.60255 | 0 | 217 |
| **TNFR: Fc3** | SQHTQPTPEPSTAPSTSFLLPMGPSPPAEGSTGDEPK | 81.39571 | 119.6185 | 10.41051 | 205 |
| **TNFR: Fc3** | SQHTQPTPEPSTAPSTSFLLPMGPSPPAEGSTGDEPK | 75.90564 | 110.6669 | 11.34904 | 205 |
| **TNFR: Fc3** | SQHTQPTPEPSTAPSTSFLLPMGPSPPAEGSTGDEPK | 76.16446 | 114.0774 | 5.7547 | 216 |
| **TNFR: Fc3** | SQHTQPTPEPSTAPSTSFLLPMGPSPPAEGSTGDEPK | 38.50392 | 59.2368 | 0 | 218 |
| **TNFR: Fc3** | SQHTQPTPEPSTAPSTSFLLPMGPSPPAEGSTGDEPK | 65.4317 | 94.64953 | 11.17001 | 213 |
| **TNFR: Fc3** | SQHTQPTPEPSTAPSTSFLLPMGPSPPAEGSTGDEPK | 59.30356 | 91.23624 | 0 | 216 |
| **TNFR: Fc3** | SQHTQPTPEPSTAPSTSFLLPMGPSPPAEGSTGDEPK | 60.76563 | 93.48558 | 0 | 216 |
| **TNFR: Fc3** | SQHTQPTPEPSTAPSTSFLLPMGPSPPAEGSTGDEPK | 74.82705 | 112.2159 | 5.39069 | 205 |
| **TNFR: Fc3** | SQHTQPTPEPSTAPSTSFLLPMGPSPPAEGSTGDEPK | 85.25001 | 125.337 | 10.80277 | 205 |
| **TNFR: Fc3** | SQHTQPTPEPSTAPSTSFLLPMGPSPPAEGSTGDEPK | 49.22665 | 73.11059 | 4.87076 | 205 |
| **TNFR: Fc3** | HTQPTPEPSTAPSTSFLLPMGPSPPAEGSTGDEPK | 49.8954 | 74.24548 | 4.67381 | 205 |
| **TNFR: Fc3** | SQHTQPTPEPSTAPSTSFLLPMGPSPPAEGSTGDEPK | 73.55427 | 107.3655 | 10.76197 | 205 |
| **TNFR: Fc3** | SQHTQPTPEPSTAPSTSFLLPMGPSPPAEGSTGDEPK | 59.13527 | 88.82375 | 3.99952 | 205 |
| **TNFR: Fc3** | SQHTQPTPEPSTAPSTSFLLPMGPSPPAEGSTGDEPK | 50.64131 | 77.90971 | 0 | 202 |
| **TNFR: Fc3** | SQHTQPTPEPSTAPSTSFLLPMGPSPPAEGSTGDEPK | 39.5297 | 60.81492 | 0 | 205 |
| **TNFR: Fc3** | SQHTQPTPEPSTAPSTSFLLPMGPSPPAEGSTGDEPK | 49.82982 | 73.92253 | 5.08622 | 205 |
| **TNFR: Fc3** | SQHTQPTPEPSTAPSTSFLLPMGPSPPAEGSTGDEPK | 54.37153 | 80.94437 | 5.02196 | 205 |
| **TNFR: Fc3** | SQHTQPTPEPSTAPSTSFLLPMGPSPPAEGSTGDEPK | 50.08888 | 74.80054 | 4.19578 | 205 |
| **TNFR: Fc3** | THTCPPCPAPELLGGPSVFLFPPKPK | 52.92914 | 71.71189 | 18.04688 | 245 |
| **TNFR: Fc3** | THTCPPCPAPELLGGPSVFLFPPKPK | 49.7825 | 75.10632 | 2.75254 | 245 |
| **TNFR: Fc3** | THTCPPCPAPELLGGPSVFLFPPKPK | 48.20176 | 64.00851 | 18.84636 | 245 |

| Sample | Peptide | TotalScore | PepScore | GlyScore | ProSites |
|---|---|---|---|---|---|
| **TNFR: Fc3** | THTCPPCPAPELLGGPSVFLFPPKPK | 53.93693 | 76.46362 | 12.10165 | 245 |
| **TNFR: Fc3** | SMAPGAVHLPQPVSTR | 26.87822 | 38.97872 | 4.40585 | 199 |
| **TNFR: Fc3** | SMAPGAVHLPQPVSTR | 25.41281 | 36.76252 | 4.33479 | 199 |
| **TNFR: Fc3** | SMAPGAVHLPQPVSTR | 35.93572 | 52.96334 | 4.31299 | 199 |
| **TNFR: Fc3** | VCDSCEDSTYTQLWNWVPECLSCGSR | 21.12385 | 32.08693 | 0.76384 | 55 |
| **TNFR: Fc3** | SMAPGAVHLPQPVSTR | 23.95752 | 34.82619 | 3.77285 | 199 |
| **TNFR: Fc3** | SQHTQPTPEPSTAPSTSFLLPMGPSPPAEGSTGDEPK | 5.05302 | 7.77387 | 0 | 202 |
| **TNFR: Fc3** | SQHTQPTPEPSTAPSTSFLLPMGPSPPAEGSTGDEPK | 9.99633 | 15.37897 | 0 | 205 |
| **TNFR: Fc3** | SQHTQPTPEPSTAPSTSFLLPMGPSPPAEGSTGDEPK | 10.38649 | 15.97922 | 0 | 205 |
| **TNFR: Fc3** | SMAPGAVHLPQPVSTR | 8.70984 | 13.39976 | 0 | 199 |
| **TNFR: Fc3** | SMAPGAVHLPQPVSTR | 47.61564 | 71.00851 | 4.17175 | 199 |
| **TNFR: Fc3** | SMAPGAVHLPQPVSTR | 35.77584 | 52.83083 | 4.10228 | 199 |
| **TNFR: Fc3** | SQHTQPTPEPSTAPSTSFLLPMGPSPPAEGSTGDEPK | 4.1145 | 6.33001 | 0 | 202 |
| **TNFR: Fc3** | SQHTQPTPEPSTAPSTSFLLPMGPSPPAEGSTGDEPK | 13.74302 | 19.20788 | 3.59399 | 205 |
| **TNFR: Fc3** | SQHTQPTPEPSTAPSTSFLLPMGPSPPAEGSTGDEPK | 3.81774 | 5.87344 | 0 | 208 |
| **TNFR: Fc3** | SMAPGAVHLPQPVSTR | 34.18823 | 50.3668 | 4.14232 | 199 |
| **TNFR: Fc3** | SQHTQPTPEPSTAPSTSFLLPMGPSPPAEGSTGDEPK | 6.12725 | 7.42907 | 3.70957 | 202 |
| **TNFR: Fc3** | SQHTQPTPEPSTAPSTSFLLPMGPSPPAEGSTGDEPK | 8.01746 | 12.33455 | 0 | 202 |
| **TNFR: Fc3** | SQHTQPTPEPSTAPSTSFLLPMGPSPPAEGSTGDEPK | 6.99947 | 10.76842 | 0 | 208 |
| **TNFR: Fc3** | SQHTQPTPEPSTAPSTSFLLPMGPSPPAEGSTGDEPK | 5.18059 | 6.05268 | 3.56099 | 202 |
| **TNFR: Fc3** | SQHTQPTPEPSTAPSTSFLLPMGPSPPAEGSTGDEPK | 12.42143 | 19.1099 | 0 | 202 |
| **TNFR: Fc3** | SMAPGAVHLPQPVSTR | 16.98661 | 24.02544 | 3.91449 | 199 |
| **TNFR: Fc3** | SQHTQPTPEPSTAPSTSFLLPMGPSPPAEGSTGDEPK | 8.34873 | 12.8442 | 0 | 202 |
| **TNFR: Fc3** | SQHTQPTPEPSTAPSTSFLLPMGPSPPAEGSTGDEPK | 5.98182 | 7.4382 | 3.27711 | 208 |
| **TNFR: Fc3** | SMAPGAVHLPQPVSTR | 33.68461 | 49.59542 | 4.13597 | 199 |
| **TNFR: Fc3** | SQHTQPTPEPSTAPSTSFLLPMGPSPPAEGSTGDEPK | 4.14047 | 6.36995 | 0 | 202 |
| **TNFR: Fc3** | SQHTQPTPEPSTAPSTSFLLPMGPSPPAEGSTGDEPK | 7.94292 | 12.21987 | 0 | 202 |
| **TNFR: Fc3** | SMAPGAVHLPQPVSTR | 28.93184 | 42.35113 | 4.01029 | 199 |
| **TNFR: Fc3** | SMAPGAVHLPQPVSTR | 22.93371 | 33.23422 | 3.80418 | 199 |
| **TNFR: Fc3** | AVHLPQPVSTR | 46.78086 | 66.0945 | 10.91266 | 199 |
| **TNFR: Fc3** | QQGNVFSCSVMHEALHNHYTQK | 19.31622 | 27.96145 | 3.2608 | 444 |
| **TNFR: Fc3** | SMAPGAVHLPQPVSTR | 49.95628 | 70.99601 | 10.88249 | 199 |
| **TNFR: Fc3** | APGAVHLPQPVSTR | 42.61276 | 62.85197 | 5.02564 | 199 |
| **TNFR: Fc3** | APGAVHLPQPVSTR | 47.66806 | 70.37396 | 5.49996 | 199 |
| **TNFR: Fc3** | SMAPGAVHLPQPVSTR | 34.61181 | 50.35445 | 5.37549 | 200 |
| **TNFR: Fc3** | SMAPGAVHLPQPVSTR | 37.26198 | 51.67583 | 10.49341 | 199 |
| **TNFR: Fc3** | SMAPGAVHLPQPVSTR | 29.69935 | 39.9865 | 10.59465 | 199 |
| **TNFR: Fc3** | SMAPGAVHLPQPVSTR | 45.94959 | 65.09204 | 10.39931 | 199 |
| **TNFR: Fc3** | SMAPGAVHLPQPVSTR | 40.98721 | 57.33356 | 10.62971 | 199 |
| **TNFR: Fc3** | SMAPGAVHLPQPVSTR | 52.87138 | 78.21935 | 5.79657 | 199 |
| **TNFR: Fc3** | SMAPGAVHLPQPVSTR | 83.53335 | 124.9418 | 6.632 | 199 |
| **TNFR: Fc3** | SMAPGAVHLPQPVSTR | 81.76604 | 122.2168 | 6.64323 | 199 |

| Sample | Peptide | TotalScore | PepScore | GlyScore | ProSites |
|---|---|---|---|---|---|
| **TNFR: Fc3** | SMAPGAVHLPQPVSTR | 67.73458 | 100.6148 | 6.67125 | 199 |
| **TNFR: Fc3** | SMAPGAVHLPQPVSTR | 62.55236 | 92.73414 | 6.50049 | 199 |
| **TNFR: Fc3** | SMAPGAVHLPQPVSTR | 57.25537 | 84.61733 | 6.44031 | 199 |
| **TNFR: Fc3** | SMAPGAVHLPQPVSTR | 29.4656 | 45.33169 | 0 | 199 |
| **TNFR: Fc3** | SMAPGAVHLPQPVSTR | 41.6325 | 58.38422 | 10.52217 | 199 |
| **TNFR: Fc3** | SMAPGAVHLPQPVSTR | 58.99258 | 87.46464 | 6.11589 | 199 |
| **TNFR: Fc3** | APGAVHLPQPVSTR | 9.56819 | 7.0287 | 14.2844 | 199 |
| **TNFR: Fc3** | APGAVHLPQPVSTR | 7.86709 | 6.32652 | 10.72814 | 199 |
| **TNFR: Fc3** | SMAPGAVHLPQPVSTR | 56.13198 | 83.08231 | 6.08136 | 199 |
| **TNFR: Fc3** | SMAPGAVHLPQPVSTR | 55.46171 | 82.07844 | 6.03064 | 199 |
| **TNFR: Fc3** | SMAPGAVHLPQPVSTR | 21.22439 | 27.32184 | 9.90058 | 199 |
| **TNFR: Fc3** | SMAPGAVHLPQPVSTR | 18.9545 | 23.56172 | 10.39822 | 199 |
| **TNFR: Fc3** | SMAPGAVHLPQPVSTR | 49.76646 | 70.77957 | 10.74213 | 199 |
| **TNFR: Fc3** | SMAPGAVHLPQPVSTR | 49.51627 | 75.35463 | 1.53074 | 199 |
| **TNFR: Fc3** | SMAPGAVHLPQPVSTR | 14.01628 | 21.56351 | 0 | 199 |
| **TNFR: Fc3** | APGAVHLPQPVSTR | 67.80923 | 92.89787 | 21.21603 | 199 |
| **TNFR: Fc3** | PGAVHLPQPVSTR | 60.02381 | 76.11631 | 30.13775 | 199 |
| **TNFR: Fc3** | PGAVHLPQPVSTR | 66.04104 | 85.54637 | 29.81686 | 199 |
| **TNFR: Fc3** | APGAVHLPQPVSTR | 47.92432 | 62.92857 | 20.05928 | 199 |
| **TNFR: Fc3** | SMAPGAVHLPQPVSTR | 52.72968 | 77.86544 | 6.04898 | 199 |
| **TNFR: Fc3** | SMAPGAVHLPQPVSTR | 49.90426 | 73.61054 | 5.87831 | 200 |
| **TNFR: Fc3** | SMAPGAVHLPQPVSTR | 71.64514 | 107.1017 | 5.79728 | 199 |
| **TNFR: Fc3** | RICTCRPGWYCALSKQEGCR | 2.57048 | 3.95459 | 0 | 107 |
| **TNFR: Fc3** | SMAPGAVHLPQPVSTR | 13.43257 | 17.70118 | 5.50515 | 199 |
| **TNFR: Fc3** | SMAPGAVHLPQPVSTR | 46.86292 | 68.85411 | 6.02213 | 199 |
| **TNFR: Fc3** | SMAPGAVHLPQPVSTR | 24.1145 | 32.11899 | 9.249 | 199 |
| **TNFR: Fc3** | SMAPGAVHLPQPVSTR | 51.74517 | 73.39912 | 11.53069 | 199 |
| **TNFR: Fc3** | SMAPGAVHLPQPVSTR | 56.53588 | 80.8916 | 11.30382 | 199 |
| **TNFR: Fc3** | SMAPGAVHLPQPVSTR | 37.36523 | 54.67757 | 5.21374 | 199 |
| **TNFR: Fc3** | SMAPGAVHLPQPVSTR | 53.88769 | 79.77476 | 5.8117 | 199 |
| **TNFR: Fc3** | SMAPGAVHLPQPVSTR | 62.96105 | 93.69449 | 5.88468 | 199 |
| **TNFR: Fc3** | PGAVHLPQPVSTR | 49.29291 | 72.22624 | 6.70245 | 199 |
| **TNFR: Fc3** | APGAVHLPQPVSTR | 65.45004 | 94.15163 | 12.1471 | 199 |
| **TNFR: Fc3** | SMAPGAVHLPQPVSTR | 51.20754 | 75.72463 | 5.6758 | 199 |
| **TNFR: Fc3** | PGAVHLPQPVSTR | 70.49409 | 101.8031 | 12.34889 | 199 |
| **TNFR: Fc3** | SMAPGAVHLPQPVSTR | 54.79393 | 81.30856 | 5.55247 | 199 |
| **TNFR: Fc3** | APGAVHLPQPVSTR | 53.61229 | 79.15156 | 6.18223 | 199 |
| **TNFR: Fc3** | SMAPGAVHLPQPVSTR | 83.88581 | 125.031 | 7.47336 | 199 |
| **TNFR: Fc3** | SMAPGAVHLPQPVSTR | 40.49723 | 62.30343 | 0 | 199 |
| **TNFR: Fc3** | APGAVHLPQPVSTR | 44.26889 | 65.17901 | 5.43582 | 199 |
| **TNFR: Fc3** | SMAPGAVHLPQPVSTR | 52.79621 | 81.22495 | 0 | 199 |
| **TNFR: Fc3** | REPQVYTLPPSR | 3.02045 | 4.64684 | 0 | 374 |

| Sample | Peptide | TotalScore | PepScore | GlyScore | ProSites |
|---|---|---|---|---|---|
| **TNFR: Fc3** | SMAPGAVHLPQPVSTR | 70.69071 | 105.3889 | 6.25113 | 199 |
| **TNFR: Fc3** | SMAPGAVHLPQPVSTR | 48.67115 | 74.87869 | 0 | 199 |
| **TNFR: Fc3** | SMAPGAVHLPQPVSTR | 47.87547 | 70.33152 | 6.17137 | 199 |
| **TNFR: Fc3** | SMAPGAVHLPQPVSTR | 41.7747 | 64.26877 | 0 | 199 |
| **TNFR: Fc3** | SMAPGAVHLPQPVSTR | 57.6667 | 85.70719 | 5.59149 | 199 |
| **TNFR: Fc3** | SMAPGAVHLPQPVSTR | 31.84309 | 46.25653 | 5.07528 | 199 |
| **TNFR: Fc3** | SMAPGAVHLPQPVSTR | 47.19566 | 69.72124 | 5.36243 | 199 |
| **TNFR: Fc3** | SMAPGAVHLPQPVSTR | 50.85264 | 75.34955 | 5.35839 | 199 |
| **TNFR: Fc3** | SMAPGAVHLPQPVSTR | 51.8199 | 76.84958 | 5.33619 | 199 |
| **TNFR: Fc3** | SMAPGAVHLPQPVSTR | 54.82087 | 81.46453 | 5.33979 | 199 |
| **TNFR: Fc3** | KCRPGFGVARPGTETS | 3.98798 | 3.56351 | 4.7763 | 132 |
| **TNFR: Fc3** | SQHTQPTPEPSTAPSTSFLLPMGPSPPAEGSTGDEPK | 46.46105 | 68.94949 | 4.69681 | 205 |
| **TNFR: Fc3** | SQHTQPTPEPSTAPSTSFLLPMGPSPPAEGSTGDEPK | 89.59768 | 131.3434 | 12.06995 | 213 |
| **TNFR: Fc3** | SQHTQPTPEPSTAPSTSFLLPMGPSPPAEGSTGDEPK | 76.90483 | 114.989 | 6.1771 | 217 |
| **TNFR: Fc3** | LPMGPSPPAEGSTGDEPK | 3.63215 | 5.09699 | 0.91173 | 226 |
| **TNFR: Fc3** | SQHTQPTPEPSTAPSTSFLLPMGPSPPAEGSTGDEPK | 12.1595 | 11.42867 | 13.51676 | 208 |
| **TNFR: Fc3** | SQHTQPTPEPSTAPSTSFLLPMGPSPPAEGSTGDEPK | 83.07171 | 124.5436 | 6.0524 | 213 |
| **TNFR: Fc3** | SQHTQPTPEPSTAPSTSFLLPMGPSPPAEGSTGDEPK | 91.24828 | 136.8482 | 6.5627 | 205 |
| **TNFR: Fc3** | SQHTQPTPEPSTAPSTSFLLPMGPSPPAEGSTGDEPK | 85.54509 | 128.0729 | 6.56488 | 205 |
| **TNFR: Fc3** | SQHTQPTPEPSTAPSTSFLLPMGPSPPAEGSTGDEPK | 76.23986 | 114.0601 | 6.00222 | 205 |
| **TNFR: Fc3** | SQHTQPTPEPSTAPSTSFLLPMGPSPPAEGSTGDEPK | 82.21579 | 123.1125 | 6.26487 | 205 |
| **TNFR: Fc3** | SQHTQPTPEPSTAPSTSFLLPMGPSPPAEGSTGDEPK | 83.61426 | 125.4042 | 6.00437 | 205 |
| **TNFR: Fc3** | SQHTQPTPEPSTAPSTSFLLPMGPSPPAEGSTGDEPK | 78.36502 | 117.357 | 5.95133 | 213 |
| **TNFR: Fc3** | SQHTQPTPEPSTAPSTSFLLPMGPSPPAEGSTGDEPK | 78.03363 | 116.7087 | 6.20852 | 205 |
| **TNFR: Fc3** | SQHTQPTPEPSTAPSTSFLLPMGPSPPAEGSTGDEPK | 82.86606 | 124.2837 | 5.94755 | 205 |
| **TNFR: Fc3** | SQHTQPTPEPSTAPST | 7.85656 | 12.08702 | 0 | 216 |
| **TNFR: Fc3** | LPAQVAFTPYAPEPGSTCR | 110.495 | 166.576 | 6.34447 | 8 |
| **TNFR: Fc3** | SQHTQPTPEPSTAPSTSFLLPMGPSPPAEGSTGDEPK | 64.84431 | 94.4504 | 9.86157 | 205 |
| **TNFR: Fc3** | SQHTQPTPEPSTAPSTSFLLPMGPSPPAEGSTGDEPK | 75.90055 | 111.3462 | 10.07294 | 205 |
| **TNFR: Fc3** | LPAQVAFTPYAPEPGSTCR | 152.1612 | 230.171 | 7.28584 | 8 |
| **TNFR: Fc3** | SQHTQPTPEPSTAPSTSFLLPMGPSPPAEGSTGDEPK | 42.01324 | 64.63575 | 0 | 205 |
| **TNFR: Fc3** | TCVVVDVSHEDPEVKFNWYVDGVEVHNAK | 3.37607 | 2.9297 | 4.20505 | 280 |
| **TNFR: Fc3** | SQHTQPTPEPSTAPSTSFLLPMGPSPPAEGSTGDEPK | 73.68877 | 108.1424 | 9.70348 | 205 |
| **TNFR: Fc3** | SQHTQPTPEPSTAPSTSFLLPMGPSPPAEGSTGDEPK | 70.6634 | 103.6174 | 9.46319 | 213 |
| **TNFR: Fc3** | SQHTQPTPEPSTAPSTSFLLPMGPSPPAEGSTGDEPK | 72.56527 | 108.6879 | 5.48031 | 213 |
| **TNFR: Fc3** | SQHTQPTPEPSTAPSTSFLLPMGPSPPAEGSTGDEPK | 59.29424 | 85.82876 | 10.01584 | 205 |
| **TNFR: Fc3** | SQHTQPTPEPSTAPSTSFLLPMGPSPPAEGSTGDEPK | 71.64474 | 104.7778 | 10.11187 | 205 |
| **TNFR: Fc3** | SQHTQPTPEPSTAPSTSFLLPMGPSPPAEGSTGDEPK | 52.06928 | 80.10658 | 0 | 205 |
| **TNFR: Fc3** | SQHTQPTPEPSTAPSTSFLLPMGPSPPAEGSTGDEPK | 73.63485 | 110.1793 | 5.76653 | 205 |
| **TNFR: Fc3** | SQHTQPTPEPSTAPSTSFLLPMGPSPPAEGSTGDEPK | 69.72711 | 104.4077 | 5.32023 | 205 |
| **TNFR: Fc3** | SQHTQPTPEPSTAPSTSFLLPMGPSPPAEGSTGDEPK | 47.18213 | 70.00862 | 4.79008 | 216 |
| **TNFR: Fc3** | SQHTQPTPEPSTAPSTSFLLPMGPSPPAEGSTGDEPK | 45.28742 | 69.67295 | 0 | 216 |

| Sample | Peptide | TotalScore | PepScore | GlyScore | ProSites |
|---|---|---|---|---|---|
| **TNFR: Fc3** | SQHTQPTPEPSTAPSTSFLLPMGPSPPAEGSTGDEPK | 55.61071 | 82.89867 | 4.93308 | 216 |
| **TNFR: Fc3** | SQHTQPTPEPSTAPSTSFLLPMGPSPPAEGSTGDEPK | 78.5362 | 117.7377 | 5.73336 | 205 |
| **TNFR: Fc3** | SQHTQPTPEPSTAPSTSFLLPMGPSPPAEGSTGDEPK | 60.82576 | 90.80272 | 5.15425 | 213 |
| **TNFR: Fc3** | SQHTQPTPEPSTAPSTSFLLPMGPSPPAEGSTGDEPK | 38.05079 | 55.95737 | 4.79572 | 216 |
| **TNFR: Fc3** | SQHTQPTPEPSTAPSTSFLLPMGPSPPAEGSTGDEP | 2.00004 | 3.07699 | 0 | 226 |
| **TNFR: Fc3** | SQHTQPTPEPSTAPSTSFLLPMGPSPPAEGSTGDEPK | 47.09739 | 72.45753 | 0 | 213 |
| **TNFR: Fc3** | SQHTQPTPEPSTAPSTSFLLPMGPSPPAEGSTGDEPK | 87.2101 | 128.9111 | 9.76532 | 205 |
| **TNFR: Fc3** | SQHTQPTPEPSTAPSTSFLLPMGPSPPAEGSTGDEPK | 90.66626 | 136.4487 | 5.64175 | 205 |
| **TNFR: Fc3** | SQHTQPTPEPSTAPSTSFLLPMGPSPPAEGSTGDEPK | 77.18643 | 118.7484 | 0 | 217 |
| **TNFR: Fc3** | SQHTQPTPEPSTAPSTSFLLPMGPSPPAEGSTGDEPK | 92.98382 | 139.946 | 5.76838 | 205 |
| **TNFR: Fc3** | SQHTQPTPEPSTAPSTSFLLPMGPSPPAEGSTGDEPK | 74.24674 | 111.2934 | 5.44586 | 213 |
| **TNFR: Fc3** | SQHTQPTPEPSTAPSTSFLLPMGPSPPAEGSTGDEPK | 93.80187 | 141.5146 | 5.19247 | 213 |
| **TNFR: Fc3** | SQHTQPTPEPSTAPSTSFLLPMGPSPPAEGSTGDEPK | 78.82404 | 118.0875 | 5.90624 | 226 |
| **TNFR: Fc3** | SQHTQPTPEPSTAPSTSFLLPMGPSPPAEGSTGDEPK | 71.6228 | 107.2875 | 5.38843 | 213 |
| **TNFR: Fc3** | SQHTQPTPEPSTAPSTSFLLPMGPSPPAEGSTGDEPK | 106.4553 | 156.3658 | 13.76441 | 226 |
| **TNFR: Fc3** | SQHTQPTPEPSTAPSTSFLLPMGPSPPAEGSTGDEPK | 67.17616 | 100.3269 | 5.6105 | 213 |
| **TNFR: Fc3** | TRSMAPGAVHLPQPVSTR | 2.88084 | 4.43206 | 0 | 186 |
| **TNFR: Fc3** | SQHTQPTPEPSTAPSTSFLLPMGPSPPAEGSTGDEPK | 66.0733 | 101.6512 | 0 | 217 |
| **TNFR: Fc3** | SQHTQPTPEPSTAPSTSFLLPMGPSPPAEGSTGDEPK | 111.0878 | 167.0745 | 7.11236 | 216 |
| **TNFR: Fc3** | SQHTQPTPEPSTAPSTSFLLPMGPSPPAEGSTGDEPK | 51.78078 | 79.66274 | 0 | 217 |
| **TNFR: Fc3** | SQHTQPTPEPSTAPSTSFLLPMGPSPPAEGSTGDEPK | 108.4137 | 163.0001 | 7.03901 | 216 |
| **TNFR: Fc3** | SQHTQPTPEPSTAPSTSFLLPMGPSPPAEGSTGDEPK | 102.2903 | 153.6471 | 6.91343 | 213 |
| **TNFR: Fc3** | SQHTQPTPEPSTAPSTSFLLPMGPSPPAEGSTGDEPK | 114.5754 | 172.4992 | 7.00255 | 213 |
| **TNFR: Fc3** | SQHTQPTPEPSTAPSTSFLLPMGPSPPAEGSTGDEPK | 20.21279 | 31.0966 | 0 | 218 |
| **TNFR: Fc3** | SQHTQPTPEPSTAPSTSFLLPMGPSPPAEGSTGDEPK | 96.2707 | 144.3428 | 6.99386 | 212 |
| **TNFR: Fc3** | SQHTQPTPEPSTAPSTSFLLPMGPSPPAEGSTGDEPK | 61.36705 | 91.59465 | 5.23006 | 205 |
| **TNFR: Fc3** | SQHTQPTPEPSTAPSTSFLLPMGPSPPAEGSTGDEPK | 37.46361 | 57.63632 | 0 | 212 |
| **TNFR: Fc3** | SQHTQPTPEPSTAPSTSFLLPMGPSPPAEGSTGDEPK | 20.05362 | 30.85172 | 0 | 212 |
| **TNFR: Fc3** | SQHTQPTPEPSTAPSTSFLLPMGPSPPAEGSTGDEPK | 112.5095 | 166.7488 | 11.77929 | 213 |
| **TNFR: Fc3** | HTQPTPEPSTAPSTSFLLPMGPSPPAEGSTGDEPK | 47.41722 | 72.17431 | 1.43977 | 226 |
| **TNFR: Fc3** | SQHTQPTPEPSTAPSTSFLLPMGPSPPAEGSTGDEPK | 43.80315 | 67.38946 | 0 | 208 |
| **TNFR: Fc3** | SQHTQPTPEPSTAPSTSFLLPMGPSPPAEGSTGDEPK | 70.91127 | 106.0615 | 5.63231 | 213 |
| **TNFR: Fc3** | SQHTQPTPEPSTAPSTSFLLPMGPSPPAEGSTGDEP | 85.58191 | 127.9935 | 6.81758 | 213 |
| **TNFR: Fc3** | SQHTQPTPEPSTAPSTSFLLPMGPSPPAEGSTGDEPK | 70.04477 | 104.8399 | 5.42521 | 213 |
| **TNFR: Fc3** | SQHTQPTPEPSTAPSTSFLLPMGPSPPAEGSTGDEPK | 108.5002 | 163.1771 | 6.95728 | 213 |
| **TNFR: Fc3** | SQHTQPTPEPSTAPSTSFLLPMGPSPPAEGSTGDEPK | 125.0542 | 188.539 | 7.15375 | 205 |
| **TNFR: Fc3** | SQHTQPTPEPSTAPSTSFLLPMGPSPPAEGSTGDEPK | 55.68139 | 82.88263 | 5.16482 | 205 |
| **TNFR: Fc3** | SQHTQPTPEPSTAPSTSFLLPMGPSPPAEGSTGDEPK | 23.28492 | 35.82295 | 0 | 205 |
| **TNFR: Fc3** | SQHTQPTPEPSTAPSTSFLLPMGPSPPAEGSTGDEPK | 113.9976 | 175.3809 | 0 | 205 |
| **TNFR: Fc3** | SQHTQPTPEPSTAPSTSFLLPMGPSPPAEGSTGDEPK | 83.28744 | 128.1345 | 0 | 205 |
| **TNFR: Fc3** | SQHTQPTPEPSTAPSTSFLLPMGPSPPAEGSTGDEPK | 46.43056 | 71.43163 | 0 | 208 |
| **TNFR: Fc3** | SQHTQPTPEPSTAPSTSFLLPMGPSPPAEGSTGDEPK | 102.4617 | 153.8562 | 7.01474 | 205 |

| Sample | Peptide | TotalScore | PepScore | GlyScore | ProSites |
|---|---|---|---|---|---|
| **TNFR: Fc3** | SQHTQPTPEPSTAPSTSFLLPMGPSPPAEGSTGDEPK | 94.82602 | 142.1577 | 6.92427 | 205 |
| **TNFR: Fc3** | QHTQPTPEPSTAPSTSFLLPMGPSPPAEGSTGDEPK | 14.10786 | 21.7044 | 0 | 217 |
| **TNFR: Fc3** | SQHTQPTPEPSTAPSTSFLLPMGPSPPAEGSTGDEPK | 99.08813 | 146.0113 | 11.94505 | 205 |
| **TNFR: Fc3** | SQHTQPTPEPSTAPSTSFLLPMGPSPPAEGSTGDEPK | 43.6304 | 67.12369 | 0 | 205 |
| **TNFR: Fc3** | SQHTQPTPEPSTAPSTSFLLPMGPSPPAEGSTGDEPK | 72.70697 | 108.7871 | 5.70098 | 205 |
| **TNFR: Fc3** | SQHTQPTPEPSTAPSTSFLLPMGPSPPAEGSTGDEPK | 47.56039 | 70.02846 | 5.83396 | 216 |
| **TNFR: Fc3** | SQHTQPTPEPSTAPSTSFLLPMGPSPPAEGSTGDEPK | 52.41805 | 77.56288 | 5.72049 | 205 |
| **TNFR: Fc3** | SQHTQPTPEPSTAPSTSFLLPMGPSPPAEGSTGDEPK | 27.61478 | 42.48428 | 0 | 208 |
| **TNFR: Fc3** | HTQPTPEPSTAPSTSFLLPMGPSPPAEGSTGDEPK | 13.42971 | 20.6611 | 0 | 212 |
| **TNFR: Fc3** | SQHTQPTPEPSTAPSTSFLLPMGPSPPAEGSTGDEPK | 69.87505 | 100.1705 | 13.61209 | 213 |
| **TNFR: Fc3** | SQHTQPTPEPSTAPSTSFLLPMGPSPPAEGSTGDEPK | 65.02093 | 96.95478 | 5.71521 | 205 |
| **TNFR: Fc3** | SQHTQPTPEPSTAPSTSFLLPMGPSPPAEGSTGDEPK | 86.88034 | 127.8169 | 10.85536 | 205 |
| **TNFR: Fc3** | SQHTQPTPEPSTAPSTSFLLPMGPSPPAEGSTGDEPK | 75.34603 | 112.9899 | 5.43598 | 205 |
| **TNFR: Fc3** | NQVSLTCLVKGFYPSDIAVEWESNGQPENNY | 2.07917 | 3.19873 | 0 | 395 |
| **TNFR: Fc3** | HTQPTPEPSTAPSTSFLLPMGPSPPAEGSTGDEPK | 42.24469 | 62.49377 | 4.63926 | 205 |
| **TNFR: Fc3** | HTQPTPEPSTAPSTSFLLPMGPSPPAEGSTGDEPK | 58.61586 | 87.6359 | 4.7215 | 205 |
| **TNFR: Fc3** | HTQPTPEPSTAPSTSFLLPMGPSPPAEGSTGDEPK | 60.03625 | 92.36346 | 0 | 205 |
| **TNFR: Fc3** | HTQPTPEPSTAPSTSFLLPMGPSPPAEGSTGDEPK | 2.72655 | 4.19469 | 0 | 205 |
| **TNFR: Fc3** | SQHTQPTPEPSTAPSTSFLLPMGPSPPAEGSTGDEPK | 50.34695 | 74.42118 | 5.63767 | 208 |
| **TNFR: Fc3** | SQHTQPTPEPSTAPSTSFLLPMGPSPPAEGSTGDEPK | 30.50276 | 46.92732 | 0 | 205 |
| **TNFR: Fc3** | SQHTQPTPEPSTAPSTSFLLPMGPSPPAEGSTGDEPK | 63.61482 | 95.20094 | 4.95489 | 205 |
| **TNFR: Fc3** | SQHTQPTPEPSTAPSTSFLLPMGPSPPAEGSTGDEPK | 79.67714 | 117.5559 | 9.33086 | 205 |
| **TNFR: Fc3** | SQHTQPTPEPSTAPSTSFLLPMGPSPPAEGSTGDEPK | 65.98025 | 99.06135 | 4.54394 | 205 |
| **TNFR: Fc3** | THTCPPCPAPELLGGPSVFLFPPKPK | 58.78772 | 80.53714 | 18.39594 | 245 |
| **TNFR: Fc3** | THTCPPCPAPELLGGPSVFLFPPKPK | 51.41376 | 75.70832 | 6.2953 | 245 |
| **TNFR: Fc3** | THTCPPCPAPELLGGPSVFLFPPKPK | 89.84109 | 133.0838 | 9.53326 | 245 |
| **TNFR: Fc3** | REQNRICTCRPGWYCALSK | 1.95529 | 3.00814 | 0 | 97 |
| **TNFR: Fc3** | SMAPGAVHLPQPVSTR | 28.89178 | 42.23499 | 4.11153 | 199 |
| **TNFR: Fc3** | SMAPGAVHLPQPVSTR | 30.44782 | 44.63004 | 4.1094 | 199 |
| **TNFR: Fc3** | SMAPGAVHLPQPVSTR | 33.30336 | 48.85656 | 4.41884 | 199 |
| **TNFR: Fc3** | SMAPGAVHLPQPVSTR | 30.2099 | 44.15582 | 4.31035 | 199 |
| **TNFR: Fc3** | SMAPGAVHLPQPVSTR | 37.74054 | 55.80307 | 4.19584 | 199 |
| **TNFR: Fc3** | SMAPGAVHLPQPVSTR | 33.22305 | 48.76936 | 4.35134 | 199 |
| **TNFR: Fc3** | SMAPGAVHLPQPVSTR | 32.17145 | 47.2165 | 4.23066 | 199 |
| **TNFR: Fc3** | SMAPGAVHLPQPVSTR | 35.00993 | 51.60294 | 4.19435 | 199 |
| **TNFR: Fc3** | SMAPGAVHLPQPVSTR | 33.63213 | 49.55901 | 4.05364 | 199 |
| **TNFR: Fc3** | SQHTQPTPEPSTAPSTSFLLPMGPSPPAEGSTGDEPK | 6.05779 | 9.31967 | 0 | 202 |
| **TNFR: Fc3** | SMAPGAVHLPQPVSTR | 37.03272 | 54.83665 | 3.96827 | 199 |
| **TNFR: Fc3** | SQHTQPTPEPSTAPSTSFLLPMGPSPPAEGSTGDEPK | 3.81224 | 4.11368 | 3.25241 | 202 |
| **TNFR: Fc3** | SQHTQPTPEPSTAPSTSFLLPMGPSPPAEGSTGDEPK | 6.51003 | 10.01544 | 0 | 202 |
| **TNFR: Fc3** | SQHTQPTPEPSTAPSTSFLLPMGPSPPAEGSTGDEPK | 4.2566 | 4.81914 | 3.21186 | 202 |
| **TNFR: Fc3** | SMAPGAVHLPQPVSTR | 37.22317 | 55.00069 | 4.20777 | 199 |

| Sample | Peptide | TotalScore | PepScore | GlyScore | ProSites |
|---|---|---|---|---|---|
| **TNFR: Fc3** | SQHTQPTPEPSTAPSTSFLLPMGPSPPAEGSTGDEPK | 2.68196 | 4.12609 | 0 | 202 |
| **TNFR: Fc3** | SMAPGAVHLPQPVSTR | 38.53626 | 57.0583 | 4.13818 | 199 |
| **TNFR: Fc3** | SQHTQPTPEPSTAPSTSFLLPMGPSPPAEGSTGDEPK | 4.44526 | 6.83886 | 0 | 202 |
| **TNFR: Fc3** | SQHTQPTPEPSTAPSTSFLLPMGPSPPAEGSTGDEPK | 4.1971 | 6.45708 | 0 | 208 |
| **TNFR: Fc3** | SQHTQPTPEPSTAPSTSFLLPMGPSPPAEGSTGDEPK | 10.10454 | 15.54544 | 0 | 205 |
| **TNFR: Fc3** | SQHTQPTPEPSTAPSTSFLLPMGPSPPAEGSTGDEPK | 5.23775 | 8.05807 | 0 | 208 |
| **TNFR: Fc3** | SQHTQPTPEPSTAPSTSFLLPMGPSPPAEGSTGDEPK | 6.38531 | 9.82355 | 0 | 208 |
| **TNFR: Fc3** | SQHTQPTPEPSTAPSTSFLLPMGPSPPAEGSTGDEPK | 10.48142 | 16.12526 | 0 | 205 |
| **TNFR: Fc3** | SQHTQPTPEPSTAPSTSFLLPMGPSPPAEGSTGDEPK | 4.52282 | 6.95819 | 0 | 202 |
| **TNFR: Fc3** | SQHTQPTPEPSTAPSTSFLLPMGPSPPAEGSTGDEPK | 5.30982 | 8.16895 | 0 | 202 |
| **TNFR: Fc3** | SQHTQPTPEPSTAPSTSFLLPMGPSPPAEGSTGDEPK | 6.51023 | 8.12443 | 3.51243 | 212 |
| **TNFR: Fc3** | SMAPGAVHLPQPVSTR | 33.54651 | 49.399 | 4.10617 | 199 |
| **TNFR: Fc3** | SQHTQPTPEPSTAPSTSFLLPMGPSPPAEGSTGDEPK | 5.67347 | 8.72842 | 0 | 212 |
| **TNFR: Fc3** | SQHTQPTPEPSTAPSTSFLLPMGPSPPAEGSTGDEPK | 7.35254 | 9.36097 | 3.62259 | 202 |
| **TNFR: Fc3** | SMAPGAVHLPQPVSTR | 31.1252 | 45.74192 | 3.97985 | 199 |
| **TNFR: Fc3** | AVHLPQPVSTR | 44.66334 | 62.95021 | 10.702 | 199 |
| **TNFR: Fc3** | APGAVHLPQPVSTR | 41.08516 | 55.19862 | 14.87446 | 199 |
| **TNFR: Fc3** | QQGNVFSCSVMHEALHNHYTQK | 22.08422 | 33.97572 | 0 | 444 |
| **TNFR: Fc3** | SMAPGAVHLPQPVSTR | 50.48893 | 71.87684 | 10.76853 | 199 |
| **TNFR: Fc3** | APGAVHLPQPVSTR | 53.1884 | 78.95598 | 5.33432 | 199 |
| **TNFR: Fc3** | SMAPGAVHLPQPVSTR | 40.62065 | 59.62417 | 5.32841 | 200 |
| **TNFR: Fc3** | SMAPGAVHLPQPVSTR | 42.66303 | 62.77341 | 5.31519 | 200 |
| **TNFR: Fc3** | APGAVHLPQPVSTR | 75.13512 | 112.3594 | 6.00439 | 199 |
| **TNFR: Fc3** | SMAPGAVHLPQPVSTR | 43.93011 | 61.76062 | 10.8163 | 199 |
| **TNFR: Fc3** | SMAPGAVHLPQPVSTR | 40.78057 | 56.85751 | 10.92339 | 199 |
| **TNFR: Fc3** | SMAPGAVHLPQPVSTR | 48.84897 | 72.15325 | 5.56959 | 199 |
| **TNFR: Fc3** | SMAPGAVHLPQPVSTR | 55.42863 | 82.06144 | 5.9677 | 199 |
| **TNFR: Fc3** | SMAPGAVHLPQPVSTR | 59.26778 | 87.98464 | 5.93648 | 199 |
| **TNFR: Fc3** | SMAPGAVHLPQPVSTR | 74.4077 | 110.5545 | 7.27792 | 199 |
| **TNFR: Fc3** | SMAPGAVHLPQPVSTR | 74.49743 | 111.1277 | 6.46983 | 199 |
| **TNFR: Fc3** | SMAPGAVHLPQPVSTR | 58.43022 | 86.38375 | 6.51652 | 199 |
| **TNFR: Fc3** | SMAPGAVHLPQPVSTR | 56.54863 | 83.74774 | 6.036 | 199 |
| **TNFR: Fc3** | SMAPGAVHLPQPVSTR | 50.12178 | 73.8203 | 6.11023 | 199 |
| **TNFR: Fc3** | SMAPGAVHLPQPVSTR | 58.6954 | 87.07502 | 5.9904 | 199 |
| **TNFR: Fc3** | SMAPGAVHLPQPVSTR | 52.86399 | 78.13247 | 5.93681 | 199 |
| **TNFR: Fc3** | SMAPGAVHLPQPVSTR | 46.15381 | 65.29914 | 10.59821 | 199 |
| **TNFR: Fc3** | SMAPGAVHLPQPVSTR | 24.74638 | 32.69056 | 9.9929 | 199 |
| **TNFR: Fc3** | SMAPGAVHLPQPVSTR | 73.16563 | 104.6011 | 14.78545 | 199 |
| **TNFR: Fc3** | APGAVHLPQPVSTR | 38.11157 | 53.28722 | 9.92823 | 199 |
| **TNFR: Fc3** | SMAPGAVHLPQPVSTR | 14.8361 | 17.86177 | 9.21699 | 199 |
| **TNFR: Fc3** | SMAPGAVHLPQPVSTR | 51.29167 | 75.92783 | 5.5388 | 200 |
| **TNFR: Fc3** | SMAPGAVHLPQPVSTR | 70.16704 | 97.67144 | 19.08744 | 199 |

| Sample | Peptide | TotalScore | PepScore | GlyScore | ProSites |
|---|---|---|---|---|---|
| **TNFR: Fc3** | SMAPGAVHLPQPVSTR | 55.50309 | 82.31596 | 5.70776 | 200 |
| **TNFR: Fc3** | SMAPGAVHLPQPVSTR | 59.71728 | 88.55739 | 6.15708 | 199 |
| **TNFR: Fc3** | SMAPGAVHLPQPVSTR | 49.46387 | 72.95898 | 5.83011 | 200 |
| **TNFR: Fc3** | SMAPGAVHLPQPVSTR | 51.64564 | 76.15239 | 6.13311 | 199 |
| **TNFR: Fc3** | SMAPGAVHLPQPVSTR | 26.23401 | 34.23172 | 11.38114 | 199 |
| **TNFR: Fc3** | SMAPGAVHLPQPVSTR | 51.93697 | 76.81825 | 5.72887 | 199 |
| **TNFR: Fc3** | SMAPGAVHLPQPVSTR | 14.96773 | 17.30437 | 10.62827 | 199 |
| **TNFR: Fc3** | SMAPGAVHLPQPVSTR | 47.49995 | 66.61084 | 12.00831 | 199 |
| **TNFR: Fc3** | SMAPGAVHLPQPVSTR | 55.0462 | 78.35416 | 11.75999 | 199 |
| **TNFR: Fc3** | SMAPGAVHLPQPVSTR | 54.25019 | 77.04782 | 11.91172 | 199 |
| **TNFR: Fc3** | SMAPGAVHLPQPVSTR | 50.87477 | 75.01251 | 6.04755 | 199 |
| **TNFR: Fc3** | SMAPGAVHLPQPVSTR | 52.05388 | 76.86411 | 5.97774 | 199 |
| **TNFR: Fc3** | SMAPGAVHLPQPVSTR | 44.03978 | 64.93791 | 5.22897 | 199 |
| **TNFR: Fc3** | SMAPGAVHLPQPVSTR | 55.22579 | 81.85183 | 5.77743 | 199 |
| **TNFR: Fc3** | SMAPGAVHLPQPVSTR | 62.83906 | 93.54125 | 5.82072 | 199 |
| **TNFR: Fc3** | APGAVHLPQPVSTR | 61.37578 | 88.01528 | 11.90243 | 199 |
| **TNFR: Fc3** | SMAPGAVHLPQPVSTR | 55.64775 | 82.3973 | 5.97001 | 199 |
| **TNFR: Fc3** | SMAPGAVHLPQPVSTR | 115.4597 | 173.4251 | 7.80953 | 199 |
| **TNFR: Fc3** | PGAVHLPQPVSTR | 54.70438 | 74.73099 | 17.51209 | 199 |
| **TNFR: Fc3** | SMAPGAVHLPQPVSTR | 61.17691 | 91.0725 | 5.65653 | 199 |
| **TNFR: Fc3** | SMAPGAVHLPQPVSTR | 135.6979 | 204.6855 | 7.57802 | 199 |
| **TNFR: Fc3** | PGAVHLPQPVSTR | 40.39422 | 58.71561 | 6.36876 | 199 |
| **TNFR: Fc3** | APGAVHLPQPVSTR | 64.3074 | 95.6747 | 6.05384 | 199 |
| **TNFR: Fc3** | SMAPGAVHLPQPVSTR | 7.68715 | 11.82639 | 0 | 199 |
| **TNFR: Fc3** | SMAPGAVHLPQPVSTR | 54.42737 | 83.73442 | 0 | 199 |
| **TNFR: Fc3** | SMAPGAVHLPQPVSTR | 35.43883 | 47.63498 | 12.78882 | 199 |
| **TNFR: Fc3** | SMAPGAVHLPQPVSTR | 53.33208 | 78.62887 | 6.35233 | 199 |
| **TNFR: Fc3** | SMAPGAVHLPQPVSTR | 49.78512 | 73.48008 | 5.78019 | 199 |
| **TNFR: Fc3** | SMAPGAVHLPQPVSTR | 46.249 | 68.01507 | 5.82631 | 199 |
| **TNFR: Fc3** | SMAPGAVHLPQPVSTR | 47.94259 | 70.89333 | 5.31979 | 199 |
| **TNFR: Fc3** | SMAPGAVHLPQPVSTR | 25.94507 | 39.91549 | 0 | 199 |
| **TNFR: Fc3** | SMAPGAVHLPQPVSTR | 21.1582 | 32.55108 | 0 | 199 |
| **TNFR: Fc3** | SMAPGAVHLPQPVSTR | 41.84979 | 61.55481 | 5.25474 | 199 |
| **TNFR: Fc3** | SMAPGAVHLPQPVSTR | 39.2119 | 57.53177 | 5.1893 | 199 |
| **TNFR: Fc3** | SQHTQPTPEPSTAPSTSFLLPMGPSPPAEGSTGDEPK | 59.85305 | 87.42117 | 8.65513 | 213 |
| **TNFR: Fc3** | SQHTQPTPEPSTAPSTSFLLPMGPSPPAEGSTGDEPK | 24.8677 | 35.71322 | 4.726 | 205 |
| **TNFR: Fc3** | SQHTQPTPEPSTAPSTSFLLPMGPSPPAEGSTGDEPK | 34.82656 | 49.31876 | 7.91247 | 205 |
| **TNFR: Fc3** | SQHTQPTPEPSTAPSTSFLLPMGPSPPAEGSTGDEPK | 42.03335 | 64.6667 | 0 | 208 |
| **TNFR: Fc3** | SQHTQPTPEPSTAPSTSFLLPMGPSPPAEGSTGDEPK | 88.13757 | 129.7954 | 10.77306 | 213 |
| **TNFR: Fc3** | SQHTQPTPEPSTAPSTSFLLPMGPSPPAEGSTGDEPK | 85.62646 | 128.455 | 6.08769 | 213 |
| **TNFR: Fc3** | SQHTQPTPEPSTAPSTSFLLPMGPSPPAEGSTGDEPK | 57.59658 | 85.74526 | 5.32045 | 205 |
| **TNFR: Fc3** | SQHTQPTPEPSTAPSTSFLLPMGPSPPAEGSTGDEPK | 97.41494 | 146.2742 | 6.67632 | 205 |

| Sample | Peptide | TotalScore | PepScore | GlyScore | ProSites |
|---|---|---|---|---|---|
| **TNFR: Fc3** | SQHTQPTPEPSTAPSTSFLLPMGPSPPAEGSTGDEPK | 90.40361 | 135.6509 | 6.37302 | 205 |
| **TNFR: Fc3** | SQHTQPTPEPSTAPSTSFLLPMGPSPPAEGSTGDEPK | 82.93824 | 124.0806 | 6.53095 | 205 |
| **TNFR: Fc3** | SQHTQPTPEPSTAPSTSFLLPMGPSPPAEGSTGDEPK | 97.31541 | 143.104 | 12.27939 | 205 |
| **TNFR: Fc3** | SQHTQPTPEPSTAPSTSFLLPMGPSPPAEGSTGDEPK | 69.43353 | 103.4997 | 6.16785 | 205 |
| **TNFR: Fc3** | SQHTQPTPEPSTAPSTSFLLPMGPSPPAEGSTGDEPK | 99.99744 | 150.3061 | 6.56701 | 205 |
| **TNFR: Fc3** | SQHTQPTPEPSTAPSTSFLLPMGPSPPAEGSTGDEPK | 39.54816 | 60.84333 | 0 | 208 |
| **TNFR: Fc3** | SQHTQPTPEPSTAPSTSFLLPMGPSPPAEGSTGDEPK | 86.86867 | 127.7192 | 11.00339 | 205 |
| **TNFR: Fc3** | SQHTQPTPEPSTAPSTSFLLPMGPSPPAEGSTGDEPK | 90.75571 | 133.2279 | 11.87885 | 205 |
| **TNFR: Fc3** | SQHTQPTPEPSTAPSTSFLLPMGPSPPAEGSTGDEPK | 79.60043 | 119.2554 | 5.95543 | 205 |
| **TNFR: Fc3** | SQHTQPTPEPSTAPSTSFLLPMGPSPPAEGSTGDEPK | 92.09026 | 138.438 | 6.01583 | 205 |
| **TNFR: Fc3** | HTQPTPEPSTAPSTSFLLPMGPSPPAEGSTGDEPK | 35.33883 | 52.20885 | 4.0088 | 205 |
| **TNFR: Fc3** | LPAQVAFTPYAPEPGSTCR | 119.3277 | 180.1165 | 6.43435 | 8 |
| **TNFR: Fc3** | LPAQVAFTPYAPEPGSTCR | 121.1863 | 182.9626 | 6.4587 | 8 |
| **TNFR: Fc3** | SQHTQPTPEPSTAPSTSFLLPMGPSPPAEGSTGDEPK | 60.43099 | 90.6865 | 4.2422 | 205 |
| **TNFR: Fc3** | LPAQVAFTPYAPEPGSTCR | 158.1817 | 239.419 | 7.31254 | 8 |
| **TNFR: Fc3** | SQHTQPTPEPSTAPSTSFLLPMGPSPPAEGSTGDEPK | 80.61326 | 120.8696 | 5.85155 | 213 |
| **TNFR: Fc3** | SQHTQPTPEPSTAPSTSFLLPMGPSPPAEGSTGDEPK | 63.25502 | 94.46006 | 5.30279 | 205 |
| **TNFR: Fc3** | SQHTQPTPEPSTAPSTSFLLPMGPSPPAEGSTGDEPK | 69.46771 | 103.9141 | 5.49585 | 205 |
| **TNFR: Fc3** | SQHTQPTPEPSTAPSTSFLLPMGPSPPAEGSTGDEPK | 70.6964 | 105.8521 | 5.40731 | 213 |
| **TNFR: Fc3** | SQHTQPTPEPSTAPSTSFLLPMGPSPPAEGSTGDEPK | 63.68633 | 95.09177 | 5.36193 | 205 |
| **TNFR: Fc3** | SQHTQPTPEPSTAPSTSFLLPMGPSPPAEGSTGDEPK | 89.15774 | 134.0838 | 5.7237 | 205 |
| **TNFR: Fc3** | SQHTQPTPEPSTAPSTSFLLPMGPSPPAEGSTGDEPK | 74.20805 | 111.2373 | 5.4394 | 205 |
| **TNFR: Fc3** | SQHTQPTPEPSTAPSTSFLLPMGPSPPAEGSTGDEPK | 84.85512 | 127.4753 | 5.70334 | 213 |
| **TNFR: Fc3** | SQHTQPTPEPSTAPSTSFLLPMGPSPPAEGSTGDEPK | 48.07887 | 73.96749 | 0 | 208 |
| **TNFR: Fc3** | SQHTQPTPEPSTAPSTSFLLPMGPSPPAEGSTGDEPK | 52.55799 | 78.26901 | 4.80894 | 216 |
| **TNFR: Fc3** | SQHTQPTPEPSTAPSTSFLLPMGPSPPAEGSTGDEPK | 69.54575 | 104.0486 | 5.46901 | 213 |
| **TNFR: Fc3** | SQHTQPTPEPSTAPSTSFLLPMGPSPPAEGSTGDEPK | 62.22972 | 92.70826 | 5.62672 | 205 |
| **TNFR: Fc3** | SQHTQPTPEPSTAPSTSFLLPMGPSPPAEGSTGDEPK | 38.73831 | 56.90904 | 4.99265 | 216 |
| **TNFR: Fc3** | SQHTQPTPEPSTAPSTSFLLPMGPSPPAEGSTGDEPK | 84.60754 | 127.3821 | 5.16918 | 213 |
| **TNFR: Fc3** | SQHTQPTPEPSTAPSTSFLLPMGPSPPAEGSTGDEPK | 71.24936 | 109.6144 | 0 | 205 |
| **TNFR: Fc3** | SQHTQPTPEPSTAPSTSFLLPMGPSPPAEGSTGDEPK | 34.24946 | 52.69148 | 0 | 213 |
| **TNFR: Fc3** | SQHTQPTPEPSTAPSTSFLLPMGPSPPAEGSTGDEPK | 57.71108 | 85.98771 | 5.19734 | 213 |
| **TNFR: Fc3** | SQHTQPTPEPSTAPSTSFLLPMGPSPPAEGSTGDEPK | 55.28272 | 82.18002 | 5.33058 | 205 |
| **TNFR: Fc3** | SQHTQPTPEPSTAPSTSFLLPMGPSPPAEGSTGDEPK | 74.9385 | 112.5276 | 5.13017 | 213 |
| **TNFR: Fc3** | SQHTQPTPEPSTAPSTSFLLPMGPSPPAEGSTGDEPK | 80.48713 | 120.6717 | 5.85859 | 226 |
| **TNFR: Fc3** | SQHTQPTPEPSTAPSTSFLLPMGPSPPAEGSTGDEPK | 83.5693 | 125.3379 | 5.99908 | 218 |
| **TNFR: Fc3** | SQHTQPTPEPSTAPSTSFLLPMGPSPPAEGSTGDEPK | 58.19525 | 86.731 | 5.20029 | 213 |
| **TNFR: Fc3** | SQHTQPTPEPSTAPSTSFLLPMGPSPPAEGSTGDEPK | 113.309 | 170.5666 | 6.97362 | 216 |
| **TNFR: Fc3** | SQHTQPTPEPSTAPSTSFLLPMGPSPPAEGSTGDEPK | 68.31122 | 105.0942 | 0 | 213 |
| **TNFR: Fc3** | SQHTQPTPEPSTAPSTSFLLPMGPSPPAEGSTGDEPK | 48.56538 | 74.71597 | 0 | 216 |
| **TNFR: Fc3** | SQHTQPTPEPSTAPSTSFLLPMGPSPPAEGSTGDEPK | 96.24026 | 144.2052 | 7.16245 | 216 |
| **TNFR: Fc3** | SQHTQPTPEPSTAPSTSFLLPMGPSPPAEGSTGDEPK | 110.8319 | 166.7467 | 6.99023 | 216 |

| Sample | Peptide | TotalScore | PepScore | GlyScore | ProSites |
|---|---|---|---|---|---|
| **TNFR: Fc3** | LLPMGPSPPAEGSTGDEPK | 2.91869 | 4.49029 | 0 | 232 |
| **TNFR: Fc3** | SQHTQPTPEPSTAPSTSFLLPMGPSPPAEGSTGDEPK | 113.6462 | 171.2212 | 6.72128 | 212 |
| **TNFR: Fc3** | SQHTQPTPEPSTAPSTSFLLPMGPSPPAEGSTGDEPK | 65.37229 | 97.21253 | 6.24042 | 213 |
| **TNFR: Fc3** | HTQPTPEPSTAPSTSFLLPMGPSPPAEGSTGDEPK | 44.2234 | 67.28641 | 1.39211 | 226 |
| **TNFR: Fc3** | HTQPTPEPSTAPSTSFLLPMGPSPPAEGSTGDEPK | 51.27607 | 73.70602 | 9.62046 | 226 |
| **TNFR: Fc3** | SQHTQPTPEPSTAPSTSFLLPMGPSPPAEGSTGDEPK | 111.7437 | 168.5841 | 6.18287 | 213 |
| **TNFR: Fc3** | SQHTQPTPEPSTAPSTSFLLPMGPSPPAEGSTGDEPK | 7.50384 | 11.54437 | 0 | 202 |
| **TNFR: Fc3** | SQHTQPTPEPSTAPSTSFLLPMGPSPPAEGSTGDEPK | 113.7768 | 171.3362 | 6.88089 | 213 |
| **TNFR: Fc3** | SQHTQPTPEPSTAPSTSFLLPMGPSPPAEGSTGDEPK | 64.11369 | 96.23985 | 4.45082 | 205 |
| **TNFR: Fc3** | SQHTQPTPEPSTAPSTSFLLPMGPSPPAEGSTGDEPK | 59.41935 | 88.58391 | 5.2566 | 205 |
| **TNFR: Fc3** | SQHTQPTPEPSTAPSTSFLLPMGPSPPAEGSTGDEPK | 77.18256 | 118.7424 | 0 | 208 |
| **TNFR: Fc3** | SQHTQPTPEPSTAPSTSFLLPMGPSPPAEGSTGDEPK | 59.22774 | 91.1196 | 0 | 205 |
| **TNFR: Fc3** | SQHTQPTPEPSTAPSTSFLLPMGPSPPAEGSTGDEPK | 107.6743 | 161.6447 | 7.44359 | 205 |
| **TNFR: Fc3** | QHTQPTPEPSTAPSTSFLLPMGPSPPAEGSTGDEPK | 11.77378 | 18.1135 | 0 | 217 |
| **TNFR: Fc3** | SQHTQPTPEPSTAPSTSFLLPMGPSPPAEGSTGDEPK | 88.71813 | 132.9286 | 6.61303 | 205 |
| **TNFR: Fc3** | QHTQPTPEPSTAPSTSFLLPMGPSPPAEGSTGDEPK | 13.30833 | 20.47435 | 0 | 217 |
| **TNFR: Fc3** | SQHTQPTPEPSTAPSTSFLLPMGPSPPAEGSTGDEPK | 29.48775 | 45.36577 | 0 | 208 |
| **TNFR: Fc3** | SQHTQPTPEPSTAPSTSFLLPMGPSPPAEGSTGDEPK | 59.01751 | 87.64745 | 5.84763 | 213 |
| **TNFR: Fc3** | SQHTQPTPEPSTAPSTSFLLPMGPSPPAEGSTGDEPK | 76.10329 | 111.1064 | 11.09746 | 213 |
| **TNFR: Fc3** | SQHTQPTPEPSTAPSTSFLLPMGPSPPAEGSTGDEPK | 71.66615 | 104.9387 | 9.87427 | 216 |
| **TNFR: Fc3** | SQHTQPTPEPSTAPSTSFLLPMGPSPPAEGSTGDEPK | 74.84556 | 112.0515 | 5.74884 | 205 |
| **TNFR: Fc3** | SQHTQPTPEPSTAPSTSFLLPMGPSPPAEGSTGDEPK | 72.17068 | 107.9355 | 5.7503 | 205 |
| **TNFR: Fc3** | SQHTQPTPEPSTAPSTSFLLPMGPSPPAEGSTGDEPK | 74.64304 | 111.8814 | 5.48606 | 205 |
| **TNFR: Fc3** | SQHTQPTPEPSTAPSTSFLLPMGPSPPAEGSTGDEPK | 88.17869 | 132.6338 | 5.61924 | 205 |
| **TNFR: Fc3** | HTQPTPEPSTAPSTSFLLPMGPSPPAEGSTGDEPK | 61.96959 | 92.77573 | 4.7582 | 205 |
| **TNFR: Fc3** | SQHTQPTPEPSTAPSTSFLLPMGPSPPAEGSTGDEPK | 17.59925 | 24.76244 | 4.29618 | 205 |
| **TNFR: Fc3** | HTQPTPEPSTAPSTSFLLPMGPSPPAEGSTGDEPK | 44.93014 | 66.50044 | 4.871 | 205 |
| **TNFR: Fc3** | HTQPTPEPSTAPSTSFLLPMGPSPPAEGSTGDEPK | 10.00528 | 15.39274 | 0 | 205 |
| **TNFR: Fc3** | SQHTQPTPEPSTAPSTSFLLPMGPSPPAEGSTGDEPK | 69.01835 | 103.3518 | 5.25634 | 205 |
| **TNFR: Fc3** | SQHTQPTPEPSTAPSTSFLLPMGPSPPAEGSTGDEPK | 28.52967 | 41.64271 | 4.17689 | 205 |
| **TNFR: Fc3** | SQHTQPTPEPSTAPSTSFLLPMGPSPPAEGSTGDEPK | 60.20471 | 92.62263 | 0 | 208 |
| **TNFR: Fc3** | SQHTQPTPEPSTAPSTSFLLPMGPSPPAEGSTGDEPK | 63.04072 | 93.9974 | 5.54976 | 205 |
| **TNFR: Fc3** | SQHTQPTPEPSTAPSTSFLLPMGPSPPAEGSTGDEPK | 66.62327 | 102.4973 | 0 | 205 |
| **TNFR: Fc3** | SQHTQPTPEPSTAPSTSFLLPMGPSPPAEGSTGDEPK | 50.17131 | 74.49293 | 5.0026 | 205 |
| **TNFR: Fc3** | THTCPPCPAPELLGGPSVFLFPPKPK | 61.65132 | 85.01141 | 18.2683 | 245 |
| **TNFR: Fc3** | THTCPPCPAPELLGGPSVFLFPPKPK | 90.06036 | 130.7529 | 14.48852 | 245 |
| **TNFR: Fc3** | THTCPPCPAPELLGGPSVFLFPPKPK | 24.65882 | 34.8819 | 5.67311 | 245 |
| **TNFR: Fc3** | THTCPPCPAPELLGGPSVFLFPPK | 37.06339 | 56.37084 | 1.2067 | 245 |
| **TNFR: Fc3** | SMAPGAVHLPQPVSTR | 26.77649 | 38.84637 | 4.36101 | 199 |
| **TNFR: Fc3** | SMAPGAVHLPQPVSTR | 32.11518 | 47.1613 | 4.17239 | 199 |
| **TNFR: Fc3** | SMAPGAVHLPQPVSTR | 27.50777 | 40.03348 | 4.24574 | 199 |
| **TNFR: Fc3** | SMAPGAVHLPQPVSTR | 27.03142 | 39.43473 | 3.99669 | 199 |

| Sample | Peptide | TotalScore | PepScore | GlyScore | ProSites |
|---|---|---|---|---|---|
| **TNFR: Fc3** | SQHTQPTPEPSTAPSTSFLLPMGPSPPAEGSTGDEPK | 7.38755 | 11.36546 | 0 | 202 |
| **TNFR: Fc3** | SQHTQPTPEPSTAPSTSFLLPMGPSPPAEGSTGDEPK | 3.23695 | 4.97992 | 0 | 208 |
| **TNFR: Fc3** | SQHTQPTPEPSTAPSTSFLLPMGPSPPAEGSTGDEPK | 12.09442 | 18.6068 | 0 | 205 |
| **TNFR: Fc3** | SMAPGAVHLPQPVSTR | 33.80441 | 49.79168 | 4.11376 | 199 |
| **TNFR: Fc3** | SQHTQPTPEPSTAPSTSFLLPMGPSPPAEGSTGDEPK | 2.58709 | 3.98014 | 0 | 202 |
| **TNFR: Fc3** | SMAPGAVHLPQPVSTR | 39.00852 | 57.81393 | 4.08417 | 199 |
| **TNFR: Fc3** | SQHTQPTPEPSTAPSTSFLLPMGPSPPAEGSTGDEPK | 5.00935 | 7.7067 | 0 | 205 |
| **TNFR: Fc3** | SMAPGAVHLPQPVSTR | 30.84196 | 45.36669 | 3.86745 | 199 |
| **TNFR: Fc3** | SQHTQPTPEPSTAPSTSFLLPMGPSPPAEGSTGDEPK | 11.9697 | 16.50731 | 3.54272 | 202 |
| **TNFR: Fc3** | SQHTQPTPEPSTAPSTSFLLPMGPSPPAEGSTGDEPK | 3.84316 | 5.91256 | 0 | 202 |
| **TNFR: Fc3** | SQHTQPTPEPSTAPSTSFLLPMGPSPPAEGSTGDEPK | 3.38176 | 5.20271 | 0 | 205 |
| **TNFR: Fc3** | SQHTQPTPEPSTAPSTSFLLPMGPSPPAEGSTGDEPK | 7.40421 | 9.85993 | 2.84359 | 208 |
| **TNFR: Fc3** | SMAPGAVHLPQPVSTR | 30.32293 | 44.46735 | 4.05472 | 199 |
| **TNFR: Fc3** | SQHTQPTPEPSTAPSTSFLLPMGPSPPAEGSTGDEPK | 2.81226 | 4.32655 | 0 | 213 |
